# Body size shapes song in honeyeaters

**DOI:** 10.1101/2023.06.20.545811

**Authors:** Eleanor M. Hay, Matthew D. McGee, Craig R. White, Steven L. Chown

## Abstract

Birdsongs are among the most distinctive animal signals. Their evolution is thought to be shaped simultaneously by habitat structure and by the constraints of morphology. Habitat structure affects song transmission and detectability, thus influencing song (the acoustic adaptation hypothesis), while body size and beak size and shape necessarily constrain song characteristics (the morphological constraint hypothesis). Yet, support for the acoustic adaptation and morphological constraint hypotheses remains equivocal, and their simultaneous examination is infrequent. Using a phenotypically diverse Australasian bird clade, the honeyeaters (Aves: Meliphagidae), we compile a dataset consisting of song, environmental, and morphological variables for 163 species and jointly examine predictions of these two hypotheses. Overall, we find that body size constrains song frequency and pace in honeyeaters. Although habitat type and environmental temperature influence aspects of song, that influence is indirect, likely via effects of environmental variation on body size, with some evidence that elevation constrains the evolution of song peak frequency. Our results demonstrate that morphology has an overwhelming influence on birdsong, in support of the morphological constraint hypothesis, with the environment playing a secondary role generally via body size rather than habitat structure. These results suggest that changing body size, a consequence of both global effects such as climate change, and local effects such as habitat transformation, will substantially influence the nature of birdsong.

## INTRODUCTION

Birdsongs are acoustic signals influencing survival, reproduction, and reproductive isolation [1, 2], thus shaping the evolution of avian diversity [3–5]. Song is a complex trait and influenced by a range of factors but is commonly thought to be mediated by two major factors, one external, one internal. First, signal detectability, which is influenced by the physical properties of the signalling environment [6, 7] is thought to be a major external factor. Consequently, habitat characteristics are proposed to influence the acoustic properties of bird song. The acoustic adaptation hypothesis (AAH) [6] specifically makes the prediction that because low frequency sounds transmit further than high frequency sounds, birds (and other animals) residing in dense habitats, such as forests, will produce lower frequency sounds than their counterparts dwelling in more open environments [8, 9]. Additionally, it is also predicted that birds will sing at slower paces to limit reverberation and further aid signal transmission [9].

Second, birdsong is also subject to constraints imposed by morphology. Body size is one of the most well understood acoustic constraints [10–12] through its effect on the size of the vocal tract and sound producing structures, which places a limit on the lowest frequency an animal can effectively produce [13]. Allometric constraints of acoustic signals are clear: generally, larger animals produce lower frequency signals [1, 10]. In birds, beak morphology influences acoustic signals too. The avian bill is used in coordination with tongue and vocal tract movements to modify sounds [14] and is thought to act as a final modulator of song, influencing both its frequency and temporal components [15–19]. These effects are complicated, however, by the allometric relationship between body size and beak size [19, 20], and by interactions among the biotic and abiotic factors that are thought to influence acoustic signalling. Specifically, beak morphology and body size also affect thermoregulation and are influenced by environmental temperatures [21–24]. In consequence, what appears to be a straightforward morphological constraint hypothesis [10–12], is more complicated.

Such complexity, caused by the effects of ecological selection on signal production, characterises both the acoustic adaptation and morphological constraint hypotheses. For example, environmental conditions not only influence body size and beak shape, but they also affect habitat productivity, complexity, and food availability, which, because of dietary preferences, influence beak shape [14, 25] and, presumably, song characteristics [14, 26, 27]. This may account for mixed outcomes when studies focus on just one of the hypotheses, with particularly contradictory results for the AAH [28, 29]. Where morphology and habitat have been considered simultaneously, habitat was found to have little effect on song, body size influences frequency, and beak size influences the temporal and performance components of song [30], but understanding of the direct and indirect influences of body size, especially in relation to the environment, remains incomplete. Therefore, although birds are one of the most diverse vertebrate clades in which acoustic signalling plays such a conspicuous role in sexual signalling and speciation [4, 5], the manner in which multiple and sometimes non-independent factors influence their signals remains opaque [18, 30], and the avian songbook thus not fully interpretable.

Here, we set out to uncover the effects of both habitat and morphology on bird song at a macroevolutionary scale, while considering underlying interactions and correlations between variables. We address this complexity explicitly in a strong inference framework [31] and use a suite of phylogenetic comparative methods [32] to explore it with simultaneous investigation of the predictions of the major hypotheses. We focus on a large and diverse yet monophyletic group of birds to do so, recognising the power that such focus on a geographically restricted, diverse single clade can bring [33]. The honeyeaters (Aves:Meliphagidae) are a clade (∼190 species) of passerine birds which have been historically confined to Australasia [34]. They are oscine passerines that learn their songs and are therefore influenced by cultural transmission [35]. Learned songs are more subject to change over generations than innate songs and are also thought to be impacted substantially by the environment because they have greater plasticity [18, 36]. Honeyeaters exhibit exceptional diversity in characters thought to influence song: body sizes range from <10 g to 350 g [37], beaks vary from short for feeding on insects to long and decurved for feeding on nectar [38, 39], and species inhabit a wide variety of biomes from arid deserts to the wet tropics [34] (Fig. 1). Consequently, song in honeyeaters varies greatly among species, with some species exhibiting complex and variable songs such as the tūī [40], while others have simpler and more stereotyped songs, such as the noisy miner [41]. Like most bird species, intraspecific variation does exist in song for some species of honeyeater [e.g. 40, 41], however the extensive diversity that exists between species is far greater than any potential within-species variation (Fig. 2). Ultimately, this immense diversity in song, body size, beak shape, and environments are ideal for disentangling the factors that shape birdsong (Fig. 3).

**Figure 1.**
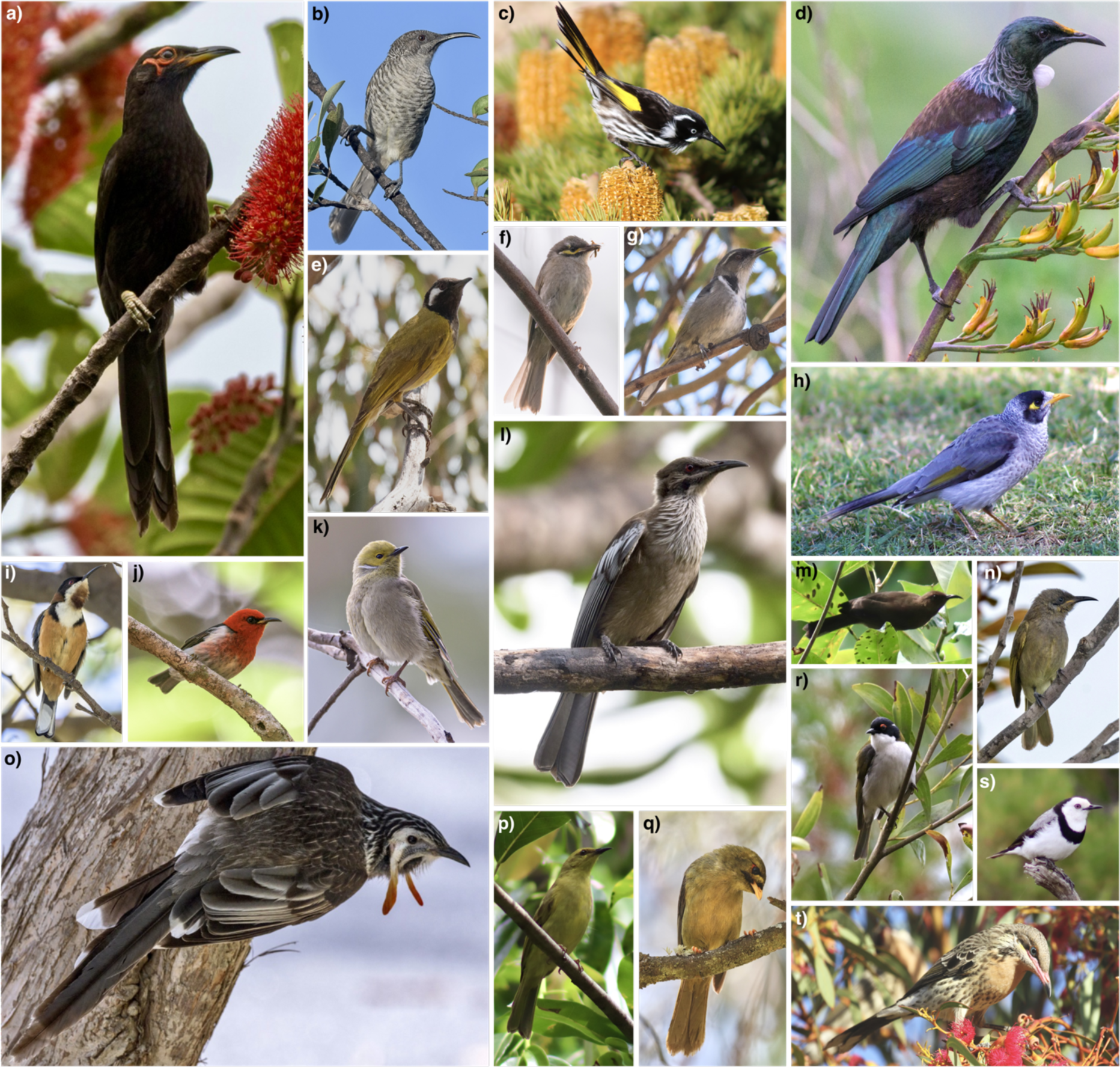
Morphological diversity of honeyeater body sizes and beak shapes. Images are of the: crow honeyeater^1^ (a; *Gymnomyza aubryana*), barred honeyeater^1^ (b; *Gliciphila undulata*), New Holland honeyeater^2^ (c; *Phylidonyris novaehollandiae*), tūī^1^ (d; *Prosthemadera novaeseelandiae*), white-eared honeyeater^3^ (e; *Nesoptilotis leucotis*), yellow- faced honeyeater^1^ (f; *Caligavis chrysops*), crescent honeyeater^1^ (g; *Phylidonyris pyrrhopterus*), noisy miner^2^ (h; *Manorina melanocephala*), eastern spinebill^1^ (i; *Acanthorhynchus tenuirostris*), New Caledonian myzomela^1^ (j; *Myzomela caledonica*), white-plumed honeyeater^1^ (k; *Ptilotula penicillata*), New Caledonian friarbird^1^ (l; *Philemon diemenensis*), dusky myzomela^2^ (m; *Myzomela obscura*), dark brown honeyeater^1^ (n; *Lichmera incana*), yellow wattlebird^1^ (o; *Anthochaera paradoxa*), yellow honeyeater^2^ (p; *Stomiopera flava*), bell miner^3^ (q; *Manorina melanophrys*), white-naped honeyeater^3^ (r;*Melithreptus lunatus*), white-fronted chat^2^ (s; *Epthianura albifrons*), spiny-cheeked honeyeater^3^ (t; *Acanthagenys rufogularis*). Image size for each species reflects relative size of the species. Image credits: ^1^Matthias Dehling, ^2^Eleanor Hay, ^3^Steven Chown.

**Figure 2.**
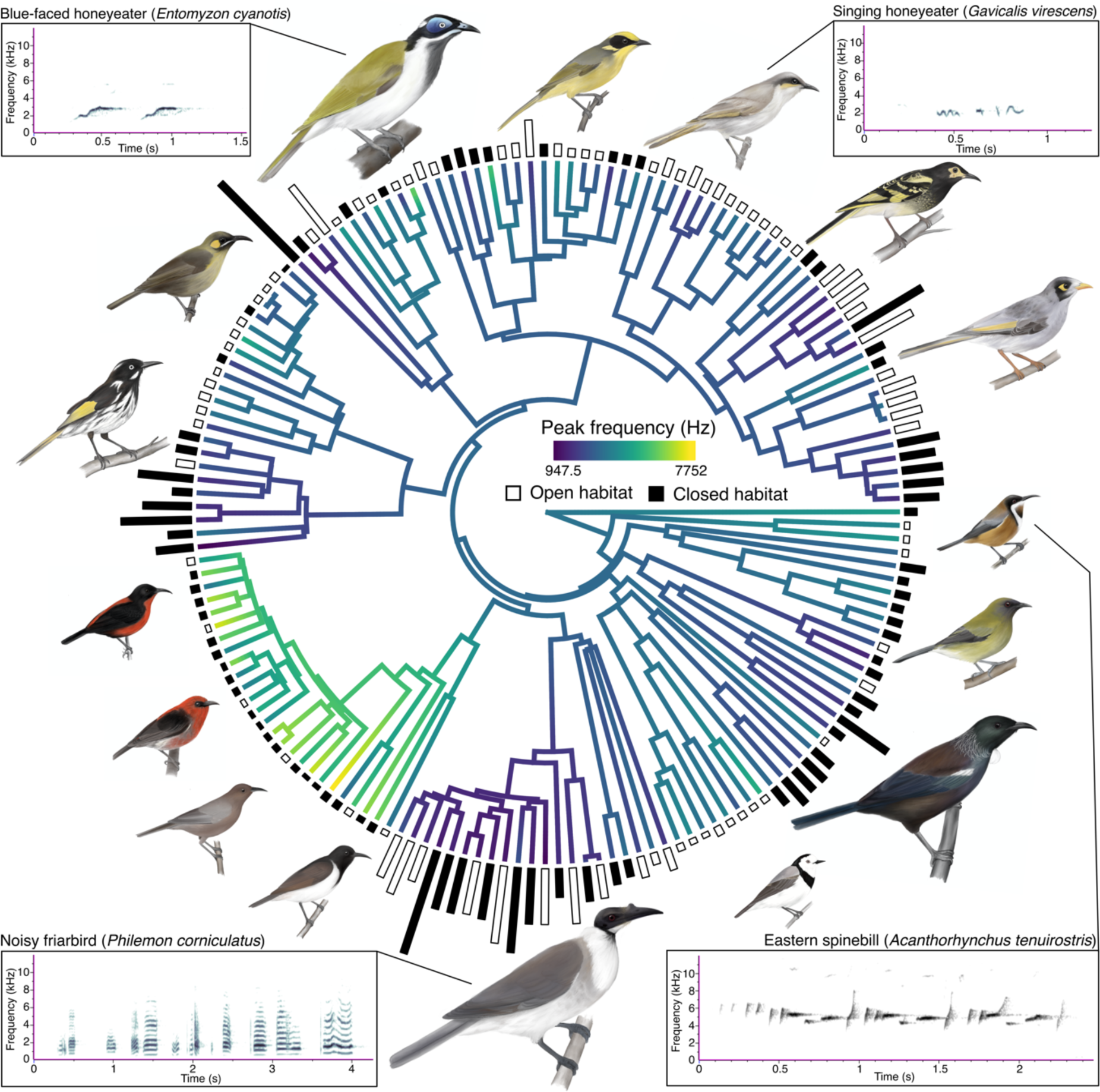
Phylogeny containing the 163 honeyeater species for which we obtained song variables for and used in analysis. Branches are coloured according to peak frequency. Size of bars at the tips of the phylogeny represent body size (mass in grams) and are coloured according to open (white) and closed (black) habitat types. Four examples of honeyeater song spectrograms are included: blue-faced honeyeater (*Entomyzon cyanotis*; xeno-canto accession number = XC439288; body size = 105.26 g), singing honeyeater (*Gavicalis virescens*; xeno- canto accession number = XC334251; body size = 23.33 g), noisy friarbird (*Philemon corniculatus*; xeno-canto accession number = XC287056; body size = 100.71 g), and eastern spinebill (*Acanthorhynchus tenuirostris;* xeno-canto accession number = XC490800; body size = 10.44 g). Birds are illustrations by Eleanor Hay.

**Figure 3.**
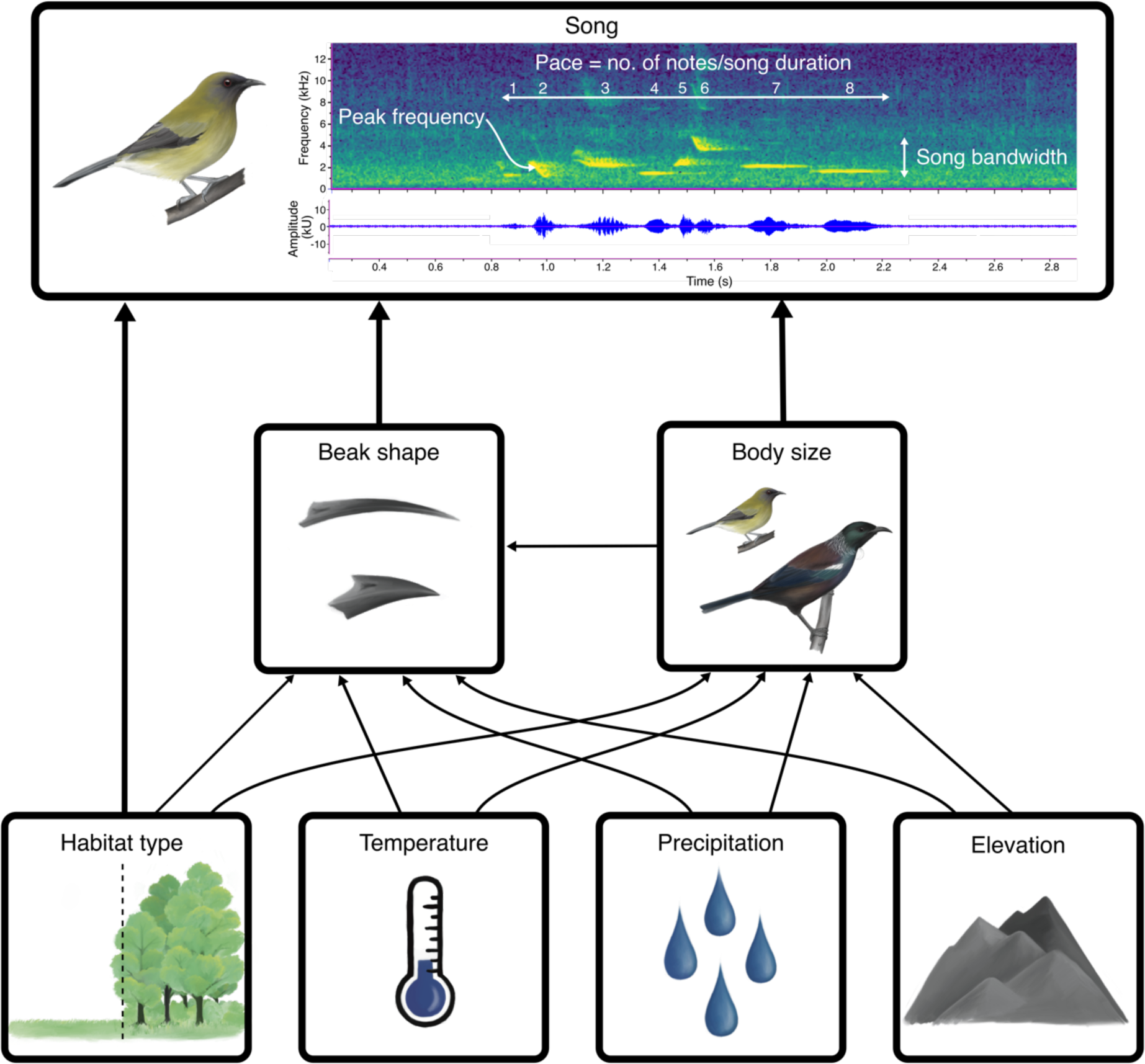
Conceptual diagram explaining the underlying interactions between environmental and phenotypic variables and their influence on song. Song spectrogram displays frequency (kHz) over time (s) and is an example from the New Zealand bellbird (*Anthornis melanura*; xeno-canto accession number = XC153005). The three song variables considered in this study are indicated with annotations; peak frequency represents the frequency of maximum amplitude across the entire song, bandwidth is the difference between maximum and minimum frequency across the song, and pace is the number of notes over song duration. Beak shape shows the beak variation in honeyeaters, the top is the species with the lowest value for beak PC2 (Arfak honeyeater; *Melipotes gymnops*) and the bottom is the species with the highest value for PC2 (Eastern spinebill; *Acanthorhynchus tenuirostris*). Illustrations by Eleanor Hay.

Specifically, we compile a dataset of song variables (peak frequency, song pace, and frequency bandwidth), morphological traits (body size and beak shape), and environmental variables (habitat type, temperature, elevation, and precipitation) for 163 species of honeyeater (Table S1). We first use a comparative phylogenetic approach consisting of phylogenetic multiple regression analyses to assess the factors influencing song traits. Models are also repeated in a Bayesian framework, and the influence of both phylogenetic and spatial covariance are considered. We next adopt an evolutionary approach, where we first examine the influence of open and closed habitats on song evolution. This is followed by phylogenetic Bayesian regression analysis in which we model the effect of continuous morphological and environmental variables on the phylogenetic variance of song, analogous to the Brownian rate parameter [42], to obtain further insight of constraints on rates of song evolution. Based on expectations of the morphological constraint hypothesis, we expect body size to constrain song frequency and beak morphology to impact the pace of which honeyeaters sing. For the acoustic adaptation hypothesis, we expect honeyeaters restricted to closed environments to sing at lower frequencies and slower paces to favour signal transmission.

## METHODS

### Honeyeater song data

Song recordings were sourced from xeno-canto (https://xeno-canto.org/), the Macaulay Library (Cornell Laboratory of Ornithology, https://www.macaulaylibrary.org/), the Australian National Wildlife Collection sound archive (ANWC), and the Avian Vocalization Center (AVoCet; https://avocet.integrativebiology.natsci.msu.edu/). We focussed exclusively on male song and did not consider recordings of female and juvenile song, or other types of vocalisations such as calls. Where available, expert classifications on recording type available in databases were followed (e.g. “song” or “call”). Following past studies which have explored song in honeyeaters, songs were defined as vocalisations including tonal elements, separated by gaps greater than 1 s [39]. After downloading, recordings were converted from.mp3 files to .wav files for analysis in Raven Pro v.1.6 (Cornell Laboratory of Ornithology). This was done in R [43] using the ‘warbleR’ package [44] and the ‘*mp32wav’* function with a sampling rate of 44.1 kHz. Here it is important to note that compression (i.e. MP3 format) of song recordings has the potential to influence precision of sound frequency measurements by up to 1 kHz in worst case scenarios [45]. In saying that, compression does not bias measurements [45], and this precision is likely to have minimal influence in macroevolutionary studies in which songs and frequency varies largely across species (Table S1; Fig. 2). To reduce any potential errors or bias cause by poor-quality recordings, we prioritised using the best quality recordings available for each species (recordings scored as “A” or “B” for xeno-canto, recordings ranked four stars or more for Macaulay Library).

To maximize the independence of our samples, only one bout of song was analysed from each recording. Maps were generated to visualise the number and location of song recordings in relation to each species breeding range using range maps from BirdLife International [46] (File S1). Recordings made at different geographic locations were assumed to be of different individual birds, and recordings that did not include a clear song example were discarded. Spectrograms were generated for 564 individuals from 163 honeyeater species (1-20 recordings per species; ∼85% of honeyeaters sampled; Table S2, File S1) using Raven Pro v.1.6 (Cornell Laboratory of Ornithology). Within Raven Pro, we used a Hann spectrogram with a window size of 256 samples. Following previous studies [e.g. 30, 39, 47], for each song up to 30 notes were manually selected and the following five song variables were extracted: Peak frequency (the frequency at maximum amplitude across the entire song recording, Hz), maximum frequency (upper frequency bound of the highest pitched note in the song, Hz), minimum frequency (lower frequency bound of the lowest pitched note in the song, Hz), song bandwidth (maximum frequency – minimum frequency), and song pace (number of notes/song duration). Peak frequency of each song was estimated using the peak frequency function available in the measurements of Raven Pro (Cornell Laboratory of Ornithology) and amplitude plots were used to measure values of maximum and minimum frequency. The mean value for each song variable was then calculated for each species. All song variables were then log_10_-transformed prior to statistical analysis to meet parametric assumptions of normality and homogeneity of variance, but also because logarithmic scales of sound frequency correspond to how birds perceive and modulate sound [48].

All five song variables are correlated to some extent, particularly the multiple measures of frequency (minimum frequency, maximum frequency, and peak frequency; Fig. S1). Thus, we chose to focus on one of the three frequency variables for simplicity. Out of the three frequency variables, peak frequency has the most evidence in support of the acoustic adaptation hypothesis [49]. Additionally peak frequency is thought to be less sensitive to background noise and recording conditions than minimum and maximum frequencies [50], and past studies considering only one frequency measurement have typically used peak frequency [e.g. 29]. For these reasons, we focus on peak frequency over maximum frequency and minimum frequency and assess the effect of predictor variables on peak frequency, pace, and bandwidth (Fig. 3). Because we do still include bandwidth, which is calculated from maximum and minimum frequency, caution needs to be taken with interpreting this variable due to the potential influence of compression and background noise on the accuracy of this measure as discussed previously.

### Trait and environmental data

To consider the influence of morphology on song, measures of body size [37, 51] and beak width, depth, and length [37, 51] were included (Table S1). As we are only using male song data, we only considered male beak measurements. Raw measurements of male beak length, width, and depth were extracted from AVONET [51] and Marki et al. [38] and the mean value was calculated for each species. All morphological variables were log_10_-transformed. From here, beak length, width, and depth were then subjected to principal component analysis (PCA) using the ‘*prcomp’* function in R [43]. The first principal component (PC1) accounted for 92.53% of the cumulative variation and describes bill size. Whereas PC2 accounted for 5.46% of the cumulative variation and describes a contrast between short and thick bills vs. long and narrow bills (visualised in Fig. 3; Table S3). A phylogenetic regression of beak PC1 on body size demonstrated that PC1 is strongly correlated with body size (PGLS regression λ = 0.66, β= 4.13 ± 0.22, *P* > 0.0001, Table S4; Fig. S2). Thus, PC2 was used to represent beak shape in analysis.

To investigate the AAH, habitat type was classified using a 2-point scale, and was classified as dense or open/semi-open habitats. We used the habitat density variable form the AVONET database which classifies birds globally into dense, semi-open, and open habitat types [51]. This defines open habitats as species primarily occurring in desert, grassland, low shrubs, rocky habitats, cities, and also applies to species living on top of forest canopy; semi- open as species primarily inhabiting open shrub, scattered bushes, parkland, dry or deciduous forest, and thorn forest; and closed habitats as species primarily living in tall evergreen forest with a closed canopy, or in the lower vegetation strata of dense thickets, shrubland or marshland [51, 52]. Because we are interested in comparing the differences between dense and open habitats, we grouped honeyeater species with open (N_open_ = 3) and semi-open (N_semi- open_ = 79) habitat types together (N_open+semi-open_ = 82) and compared them to species from dense habitats (N_dense_ = 81).

Body size and beak morphology are influenced by the environment [21–24, 39], therefore appropriate bioclimatic variables were extracted from the geographic range of each species and included in analysis. Because this study is conducted on a macroevolutionary scale, we focused on classifying these variables at the species-level, and honeyeater breeding range maps from BirdLife International [46] were used to sample values of mean annual temperature, annual precipitation, and elevation from the BioClim database of present-day climatic conditions [53]. These three bioclimatic variables were selected because they are all commonly expected to influence size and thermoregulation: mean annual temperature was included because temperature is expected to directly influence thermoregulation [24]; annual precipitation was selected because this is also strongly liked with thermoregulation and is especially important for areas with large variation in precipitation such as Australasia [54]; and elevation was included because lower elevations typically have warmer temperatures than higher elevations, and vary in other ways that have significant impacts on bird size [55, 56]. We used a resolution of 1 km^2^, and calculated the mean value for each bioclimatic variable across each species geographic range using the ‘raster’ package [57] in R [43].

### Phylogeny

Trait values of closely related species are expected to be more similar than those of distantly related species. Therefore, to comparatively analyse relationships between habitat, morphology, and song while accounting for phylogenetic history, we used a recently reconstructed honeyeater phylogeny from Hay et al. [58]. This phylogeny was constructed using a combination of nuclear genes, mitochondrial genes, and topological constraints from previous phylogenomic studies. The phylogeny was trimmed, using the ‘*drop.tip’* function from the ‘ape’ package [59], to the 163 species for which we had song data and was used as a phylogenetic effect in all analysis.

To visualise correlations between the morphological constraint hypothesis and the acoustic adaptation hypothesis, song variables, morphological traits, and habitat type were plotted on the phylogeny (Fig. 2) using the using the functions ‘*plotTree.wBars’* and ‘*contMap’* from the R package ‘phytools’ [60].

### Analyses

We used two approaches to assess the acoustic adaptation hypothesis and the morphological constraint hypothesis. The first is a typical comparative approach using regression analysis to identify relationships, whereas in the second an evolutionary constraint method is adopted.

When investigating hypotheses in a strong inference context, several biases might arise. To account for these potential biases a directed acyclic graph [DAG; 61] was created to visualise causal relationships and the backdoor criterion was applied to determine the covariates required to assess both hypotheses while considering interactions between the two hypotheses [62] (Fig. 3). In this case, two confounding biases exist in which a predictor variable affects both another predictor variable and the response variable of interest [62]. The first bias exists between body size and beak shape, which are two strongly correlated traits that have similar predictions for how they could each influence song frequency [19].

Statistically it can be difficult to disentangle effects such interrelated traits might have and there is a possibility that any effect of beak shape on song might not actually be attributed to that variable but could in fact be a statistical artefact [19, 20, 63]. Thus, to overcome this, the effect of body size needs to be controlled and body size must also be included as a predictor in analysis. The second bias exists between the interrelated effects of phenotypic traits and the environment. Environment has the potential to affect beak shape and body size, as well as song, and therefore this needs to be controlled by including environmental predictor variables in the model as well. Thus, in this scenario, to address both the acoustic adaptation hypothesis and the morphological constraint hypothesis body size, beak shape, habitat type, temperature, precipitation, and elevation must all be included as predictor variables in analysis (Fig. 3; DAG model available at <u>dagitty.net/mz3jqZU</u>). To further consider the direct effects of environmental variables on body size and beak shape we also use models where we explicitly test the effect of precipitation, elevation, and temperature on both body size and beak shape.

We first approach the acoustic adaptation hypothesis and the morphological constraint hypothesis from a typical comparative approach to determine the drivers of song.

Phylogenetic generalised least squares regression (PGLS; [64, 65] was used to determine the influence of predictor variables on song. PGLS regression tests were run using the ‘ape’ [v5.3; 59] and ‘nlme’ [66] packages in R [43]. An estimated lambda parameter was used to control for the amount of phylogenetic effect in the model residuals [67].

Spatiophylogenetic modelling [68] was then used to estimate the spatial and phylogenetic structure in honeyeater song, and to examine the contributions of body size, beak shape, and environment to variation in song. To model the spatial distributions of each species, we used honeyeater point occurrence records from Hay et al. [58] which were originally downloaded from eBird via GBIF [69, 70]. This dataset was reduced to match the number of species included in our study, and our analysis included a total of 253,742 honeyeater point occurrence records. The spatiophylogenetic model was implemented in the R package ‘INLA’ which uses an integrated nested Laplace approximation to estimate the joint posterior distribution of model parameters [71, 72]. INLA is an approach for latent Gaussian Markov random field models, providing a significantly faster alternative to Markov chain Monte Carlo [72]. Prior to analysis, all predictor variables were standardised by subtracting the mean and dividing by the standard deviation. For the phylogenetic effect, INLA requires a phylogenetic precision matrix which is the inverse of a phylogenetic covariance matrix. Prior to inverting, the phylogenetic covariance matrix was standardised by dividing by its determinant raised to the power of 1/N_species_ [68]. To model the spatial effect across the entire landscape, we followed a previous study that has used this approach to model speciation patterns in honeyeaters [58], and used a spatial mesh consisting of 859 vertices, which was constructed over honeyeater point occurrence records and averaged across the spatial random fields of each occurrence record for each species (see supplementary R script). This approach integrates a spatial random field across each species’ distribution, so each species contributes a single datapoint to the likelihood. This approach also reduces any bias which could arise from biased data in occurrence records due to differential sampling effort, and to some extent accounts for variation between wide ranging and narrowly distributed species. Further information on these models is available in Dinnage et al. [68]. Default priors were used for both the phylogenetic and spatial effects. The significance of effects in the models were based on whether the 95% credible intervals of the effect size overlapped with zero.

We then approach the acoustic adaptation hypothesis and the morphological constraint hypothesis from an evolutionary perspective to assess the influence of potential constraints by examining rates of trait evolution using Brownian motion and Ornstein-Uhlenbeck models [42, 73–75]. We fit multivariate Brownian motion (BM) and Ornstein-Uhlenbeck (OU) models in R [43] using mvMORPH [76] to assess whether open and closed habitat have constrained song evolution. We constructed multivariate Ornstein-Uhlenbeck models and Brownian motion models with either one rate (no evolutionary relationship between open and closed habitats) or two rates of evolution (open and closed habitats have evolved independently) and tested the effect on song pace, bandwidth, peak frequency. We performed stochastic character mapping of open and closed habitats on the phylogeny using the *make.simmap* function from the ‘phytools’ package [60] and generated 100 stochastic character histories using the tip states on the tree. Model fit was compared with AIC. Despite being limited to testing the influence of discrete predictors, the mvMORPH approach is beneficial here as a multivariate framework enables us to determine the influence of open and closed habitats on peak frequency, bandwidth, and pace altogether.

Finally, we apply phylogenetic Bayesian regression [77, 78] to model the effect of continuous morphological and environmental variables on the phylogenetic variance of song, analogous to the Brownian rate parameter, to obtain further insight on constraints from an evolutionary perspective. This approach enables us to examine how continuous predictor variables such as body size and beak shape may constrain rates of song evolution. Models were implemented in R with the package ‘brms’ [79], for Bayesian models using the probabilistic programming language STAN [77, 78], which is a Markov chain Monte Carlo (MCMC) sampler that uses techniques based on Hamiltonian Monte Carlo [80]. Here, we estimate the effect of predictor variables on both the mean to confirm earlier PGLS and INLA results, as well as the variance to examine constraints on rates of trait evolution. All continuous variables were transformed to have a mean of 0 and a standard deviation of 0.5.

Phylogenetic relatedness of species in our regression analysis was accounted for with a phylogenetic covariance matrix calculated using the *vcv* function from the R package ‘ape’ [59]. Models were run for at least 2,000 MCMC iterations with 4 chains and convergence was assessed using the estimated potential scale reduction statistic Rhat [78]. Effects were judged as significant if the 95% credible intervals of the effect size did not overlap with zero.

## RESULTS

Multiple PGLS regression models testing the effect of morphological and environmental variables on song revealed several relationships. Peak frequency (β = -0.27 ± 0.041, *P* < 0.0001; model λ = 0.44, *N* = 163) and pace (β = -0.31 ± 0.087, *P* < 0.001; model λ = 0.51, *N* = 163) are negatively related to body size, indicating that overall, smaller birds produce higher frequency sounds and faster paced songs (Fig. 4a; Table S4). Beak shape and habitat type had no impact on any of the song variables (Fig. 4a; Table S4). By contrast, bandwidth (β = -0. 020 ± 0.0071, *P* = 0.005; model λ = 0.44, *N* = 163) is negatively related to temperature, in which cooler mean annual temperatures are associated with broader bandwidths (Fig. 4a; Table S4). Body size in honeyeaters is positively related to precipitation suggesting that honeyeaters are larger in wetter areas (β = 0.0001 ± 0.000017, *P* = 0.0021; model λ = 0.99, *N* = 163; Table S4). A bivariate PGLS model of habitat type on body size identified honeyeaters to be larger in closed than open environments (β = 0.056 ± 0.024, *P* = 0.021; model λ = 1.00, *N* = 163; Table S4).

**Figure 4.**
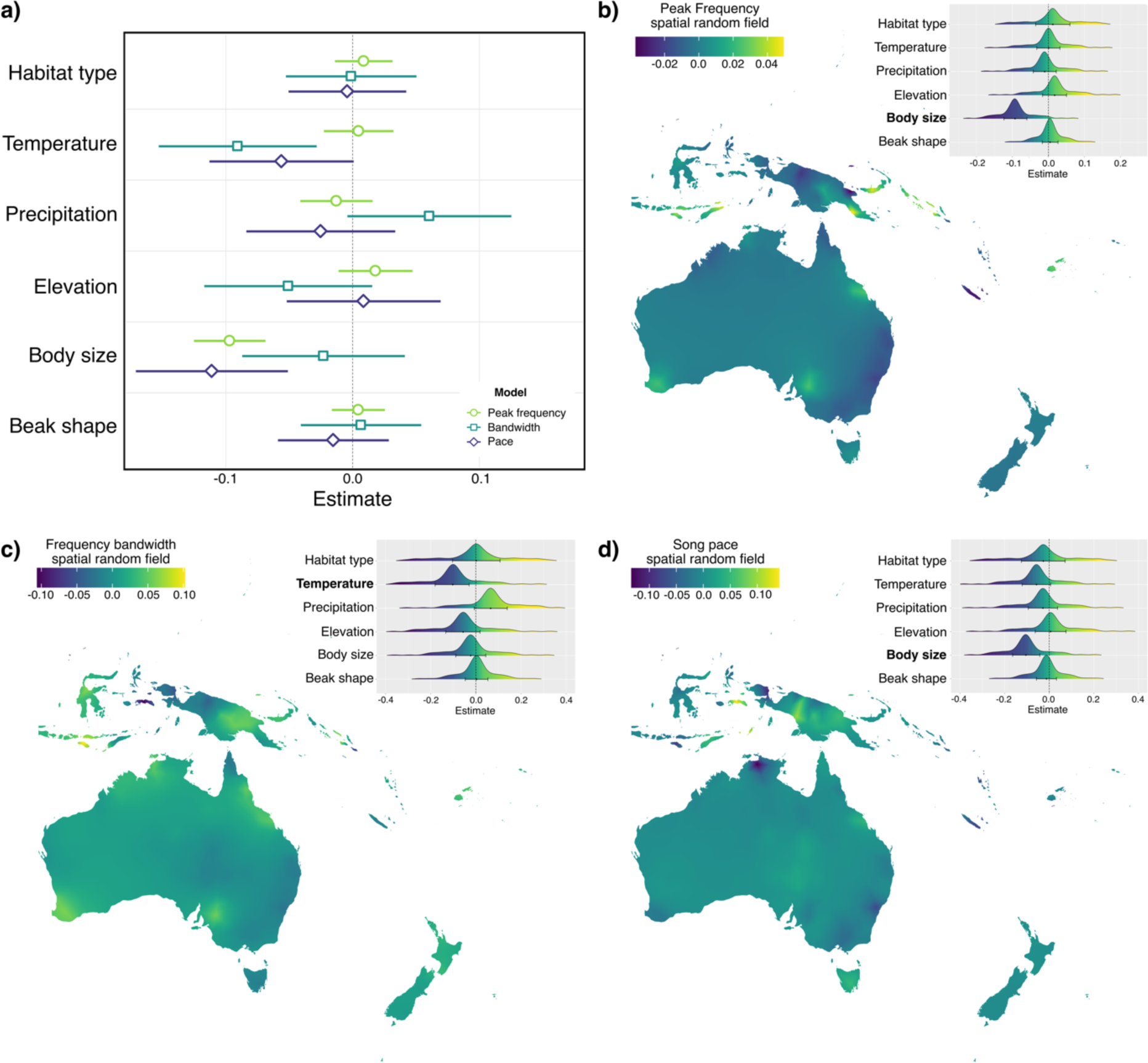
Results from phylogenetic generalised least squares regression models (a) and the spatiophylogenetic models (c-d), testing the influence of environmental and morphological traits on song variables. Separate models were run to test the influence on peak frequency, bandwidth, and pace, and are indicated in the legends. For the PGLS results (a), the predictor variables have been scaled so the effect sizes are comparable, full model results are available in Table S4. Maps in panels b-c display the mean spatial random effect for each song variable which was extracted from the final spatiophylogenetic model. Density plots show the marginals of the fixed effects in the model, with the mean and 95% credible intervals. Predictor variables which were identified to have significant effects are highlighted in bold, and full model results are available in Table S5.

Spatiophylogenetic analysis revealed considerable spatial and phylogenetic structure to the variation in honeyeater songs and show that song traits are variable across the distribution of honeyeaters (Fig. 4b-d). Consistent with PGLS models, frequency and pace are negatively related to body size (Fig. 4b; Fig. 4d; Table S5). Models further confirm the effects of temperature on frequency bandwidth, indicating that species in warmer areas sing with narrower bandwidths (Fig. 4c). Congruent with PGLS results, a positive relationship exists between precipitation and body size (Table S5).

Model comparisons of multivariate Ornstein-Uhlenbeck models and Brownian motion models indicate that open and closed habitat types have influenced the evolution of song in honeyeaters. The best model based on AIC was an Ornstein-Uhlenbeck model with two rates of evolution (Fig. 5a; Table S6), implying that song has evolved toward different evolutionary optima in open and closed habitats. Estimated theta values extracted from these models show that values tend to be lower for closed habitats than for open habitats, suggesting that lower values of frequency, bandwidth, and pace are selected for in closed habitats.

**Figure 5.**
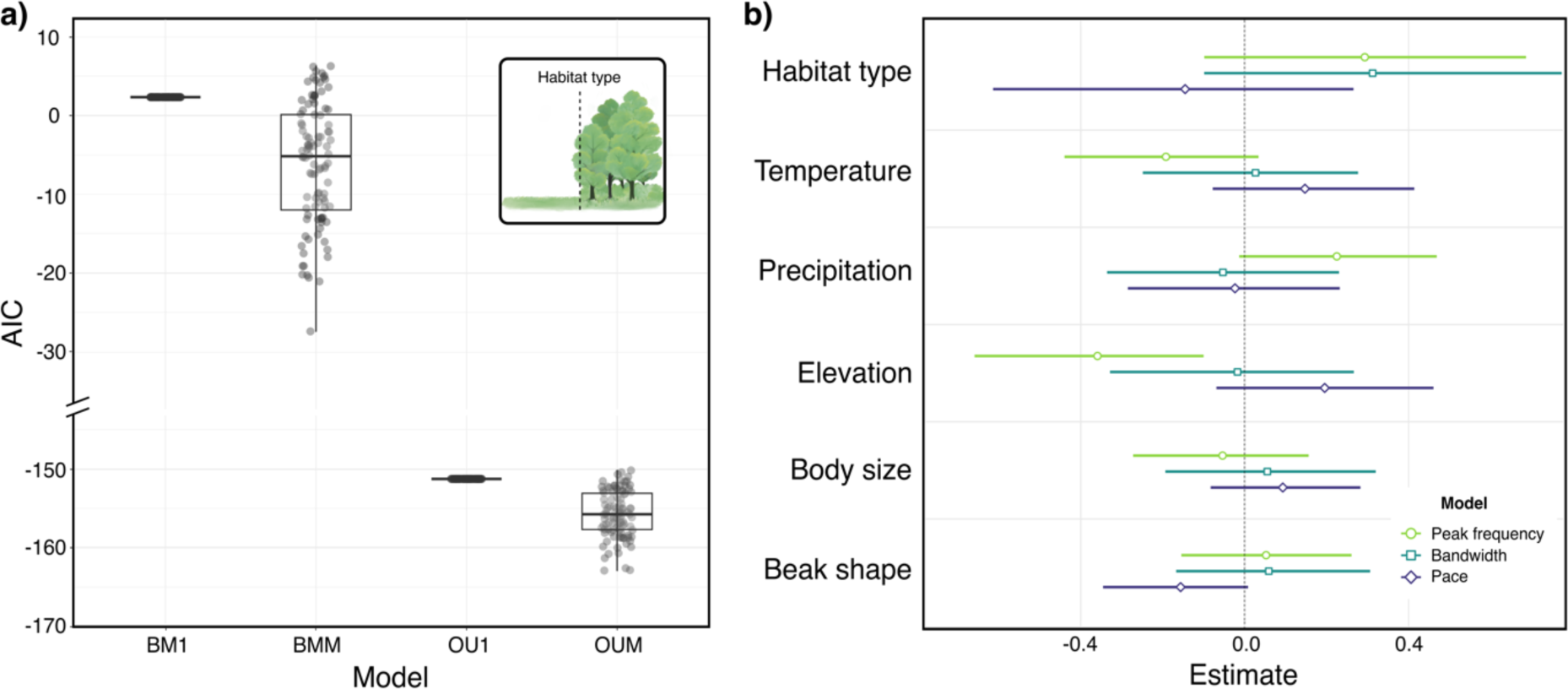
Distribution of AIC values from mvMORPH models (a) when testing the effect of open and closed habitat types on peak frequency, bandwidth, and pace. Note that models were conducted over 100 stochastic character mapping of open and closed habitats on the phylogeny. Models compared were single rate Brownian motion (BM1, n = 100), two rate Brownian motion (BMM, n = 100), single rate Ornstein-Uhlenbeck (OU1, n = 100) and two rate Ornstein Uhlenbeck (OUM, n = 100). Full model results are available in Table S6. Panel b displays outcomes of Bayesian phylogenetic regression models (a), testing the influence of environmental and morphological traits on the variance of peak frequency, bandwidth, and pace. Models are indicated with the legend and full model results are available in Table S7.

Bayesian phylogenetic regressions confirm all results of PGLS models (Table S7). When modelling the influence of predictor variables on the phylogenetic variance of song, we find peak frequency to be constrained by elevation. This indicates that honeyeaters with at lower elevations have higher rates of evolution, whereas higher elevations constrain the rate of peak frequency evolution (Fig. 5b; Table S7).

## DISCUSSION

In this study we used honeyeaters to test the predictions of two key hypotheses thought to influence acoustic signals: the morphological constraint hypothesis and the acoustic adaptation hypothesis. We found evidence to support the predictions of the morphological constraint hypothesis: body size constrains frequency and pace. We found no influence of beak shape on song, however. The predictions of the acoustic adaptation hypothesis were not fully supported, but temperature was found to influence bandwidth, and we found some evidence habitat type has influenced the evolution of song. However, these findings are likely mediated by environmental effects on body size. Nonetheless, our evolutionary approach showed that elevation constrains the evolution of peak frequency. Overall, we found that the main constraint of song is body size, and this is ultimately modulated by underlying environmental effects of precipitation on body size, which has consequences for song evolution.

### Morphological constraints on song

In the case of the morphological constraint hypothesis, we found overwhelming evidence that body size constrains peak frequency: smaller-sized honeyeaters produce higher frequency songs. This outcome is congruent with previous findings for other bird groups [18, 28–30, 47, 81], and likely the outcome of a proximal mechanitic relationship where larger birds have larger sound-producing organs, such as the syrinx [10–12]. In turn this raises the question of what factors influence body size. In African tinkerbirds, temperature and latitude account for variation in body size, ultimately having consequences on song frequency [82].

Among honeyeater species, no evidence exists for latitudinal variation in body size (Bergmann’s rule) [58], but the current analyses show that that honeyeater body size is negatively related to annual precipitation, indicating that species from wetter areas tend to be larger than those from drier areas. This could be in part driven by some form of tropical temperature effect but is also likely the outcome of aridity gradients and honeyeater responses to them across Australasia. Body size is fundamental to an organism’s ability to maintain homeostasis [23, 24], and precipitation is a key factor in thermoregulation [54, 83]. Australia has a particularly strong precipitation gradient from the coast to the interior [84]. The influence of such precipitation gradients on body size in honeyeaters is apparent in arid zone honeyeaters which comprise chats (genus *Epithanura*) and the gibberbird (*Ashbyia lovensis*). These honeyeater species are relatively small, weighing 9-13.5 g (*Epithanura* spp.) and 18 g (*Ashbyia lovensis*), respectively [37] (Table S1), and occupy the driest areas across the study region. Furthermore, this interplay between body size, song frequency, and the environment could suggest that indirect effects via ecological selection, in this case precipitation, on body size in honeyeaters can have consequences for signal production.

The morphological constraint from beak shape is not as strong as that for body size and no evidence was found for an influence of beak shape on song. This is contrary to expectations as a previous study including 88 species of honeyeater found both beak size and shape to negatively influence pace and minimum frequency, respectively [39]. However, that study did not consider influences of habitat in these models, which was acknowledged as a limitation because of the complex relationships between beak size and shape, foraging ecology, and the environment [39]. Beak morphology in general is understood to act as a mechanical constraint on song and influence movement [14, 16], the extent of which is underpinned by environmental conditions and beak specialisation for dietary preferences. For example, mechanics behind small and slender beaks of insectivores prioritise closing velocity for catching insects, whereas curved and narrow beaks of nectarivores prioritise nectar transfer [14], and these factors can influence song. Ultimately, this complex relationship between beak morphology, the environment, and song highlights the importance of exploring effects of habitat and morphology altogether.

Numerous past studies have identified beak attributes, prominently beak size, to impact song frequency [15, 16, 18, 39], but we find no evidence for an impact of beak shape on peak frequency. Disentangling the effects of beak size and shape on song is complicated by allometric effects of body size on beak size which could confound the relationship between beak size and many song features. In the present study we accounted for this allometry by using the second principal component of beak variables, representing shape, and controlled for body size in all analysis. The discrepancy between the present study and previous work is not overly surprising given that we focus on beak shape rather than size.

However, only a few past studies have explicitly acknowledged this statistical concern [19, 30] and caution is required when interpreting findings which might not have fully accounted for body size because any effect of beak shape on song variables could be mistakenly attributed [19, 20, 63]. Beyond statistical concerns, fully understanding the influence of beak shape on song is not entirely straightforward and requires more attention because behaviours such as adjusting gape and coordinated movements of the beak with the tongue also alter amplification and resonance of song [14].

Body size also constrains song pace: larger honeyeaters sing at slower paces. This negative effect of body size on song pace is consistent with previous studies of birds [28, 30], however, it has generally been attributed to beak size, and little discussion of the relationship between body size and pace exists throughout the literature. Certainly, beak shape does constrain pace in honeyeaters, but there is clearly an effect of body size too. One potential explanation is that body size influences air intake and output which could determine song pace. Larger birds have larger lungs and vocal producing organs, but also have greater oxygen consumption, and take larger and fewer breaths than smaller birds [85] which could explain differences in pace. Morphological structures such as the glottis which acts as a valve between the lungs and mouth, and the trachea which connects the syrinx in birds to the lungs, determine air movement. In most birds the trachea narrows towards the lungs [86], potentially limiting air movement, and could cause larger lungs take longer to fill and empty, perhaps placing a constraint on the pace at which birds can sing.

### Environmental constraints on song

Empirical evidence for the acoustic adaptation hypothesis has remained mixed and its predictions are generally not supported [9, 29, 36, 49]. In the present study, no direct evidence is uncovered for an effect of habitat type on song at a macroevolutionary scale in honeyeaters. The hypothesised selection imposed by closed habitats is likely not strong enough to have a macroevolutionary influence, or perhaps open habitats are also subject to their own constraints on the transmission and reception of signals. Signals produced in open habitats can experience degradation from wind and environmental attenuation through the atmosphere, ground, and vegetation [7]. Conversely, any constraint of closed habitats could possibly be overcome through behaviours such as movement. Honeyeaters in general have high dispersal ability [38, 58] and smaller sized birds could implement behaviours such as increased movement to compensate if acoustic signals are not being detected [87].

Our evolutionary approach did, however, indicate that habitat type influences the evolution of song. Here, model comparisons of multivariate Ornstein-Uhlenbeck models and Brownian motion models indicate song in open and closed habitats has evolved toward different optima. This was generally found to support the predictions of the acoustic adaptation hypothesis: lower values for peak frequency, bandwidth, and pace tended to be preferred in closed habitats. However, analysis revealed that honeyeaters tend to be larger in closed than open environments. Such underlying relationships suggest that any effect we uncover for habitat type influencing song is more likely to be driven by underlying variation in body size, and ultimately provides further support for body size being the primary constraint of song.

Although little support is uncovered for the AAH, we find some evidence that other environmental variables influence song in honeyeaters. In particular, bandwidth is consistently found to be negatively related to mean annual temperature, indicating that honeyeater species in warmer areas sing with narrower bandwidths or less frequency variation. Although the impacts of temperature on bandwidth could be related to habitat type, body size is more likely the main influence. As we have shown, body size in honeyeaters is structured by precipitation, however body size in birds is also commonly structured by temperature [24, 55] and there is likely an underlying relationship between temperature and body size in honeyeaters. Indeed, both temperature and precipitation jointly influence thermoregulation and desiccation, and can affect body size [54]. Body size is understood to place a limit on the lowest pitched sound a bird can produce, and larger species use wider bandwidths than smaller species [13].

Lastly, our evolutionary approach also found that elevation constrains the rate of peak frequency evolution, and that species occurring at lower elevations tended to have faster rates of peak frequency evolution than those at higher elevations. One of the most notable areas for elevation in honeyeaters is New Guinea, and interestingly our spatial maps (Fig. 4b-c) indicate that New Guinea is associated with higher frequencies, greater bandwidths, and faster paces. New Guinea also contains the greatest proportion of species richness for honeyeaters, where up to 41 species co-occur [38]. Past studies have found that niche partitioning along elevational zones in New Guinea has enabled many divergent species to co-exist here [88], and perhaps the constraints of elevation on peak frequency evolution in this area has contributed to this co-existence of species. Nevertheless, it is also likely that niche partitioning has resulted in divergent body sizes which has then influenced peak frequency, and the result of elevation constraining peak frequency evolution could simply be an artefact of this.

## Conclusion

This study provides support for the morphological constraint hypothesis and firmly establishes body size as the main constraint of song in honeyeaters. Effects of habitat type and any apparent environmental drivers on song have stronger associations with body size than with habitat structure *per se*.

These findings have implications for the future of birdsong in a rapidly changing world. Changing climates are resulting in a reduction in avian body size [23, 89], and habitat loss and transformation include declines of larger sized birds as part of overall changes to local assemblage composition [90, 91]. Our results therefore forecast an increase in higher pitched songs as body sizes decline. Overall, we demonstrate that morphology and habitat are deeply intertwined and can jointly influence song. Thus, climate change and habitat loss will not only have consequences for acoustic signalling and song evolution, but will also transform the avian songbook.

## Acknowledgements

We thank Leo Joseph and Margaret Cawsey at ANWC for assistance with accessing recordings and acknowledge receipt of media from The Macaulay Library at the Cornell Lab of Ornithology. Members of the Chown Lab, Melodie McGeoch, and members of the McGeoch lab provided feedback on aspects of this work. Two anonymous reviewers provided useful comments on a previous version of the manuscript.

## Funding

EMH was supported by an Australian Government Research Training Program (RTP) Scholarship, a Monash Graduate Excellence Scholarship (MGES), and a Faculty of Science Dean’s Postgraduate Research Scholarship. SLC was funded by Australian Research Council Discovery Project DP170101046. MDM was funded by Australian Research Council Discovery Early Career Research Award DE180101558.

## Authors’ contributions

EMH and SLC conceptualised the study. Trait and environmental variables were gathered by EMH. Processing of song recordings and song measurements were completed by EMH. Analysis was completed by EMH with supervision and input from MDM, CRW, and SLC. EMH wrote the manuscript with input from SLC, MDM, and CRW. All authors read and approved the final manuscript.

## Notes

### Competing Interest Statement

The authors have declared no competing interest.

### Summary of Updates

Just wanted to update the distribution/reuse option, there have been no other changes!

