## Supplementary material for "Body size shapes song in honeyeaters": Figures S1-S2

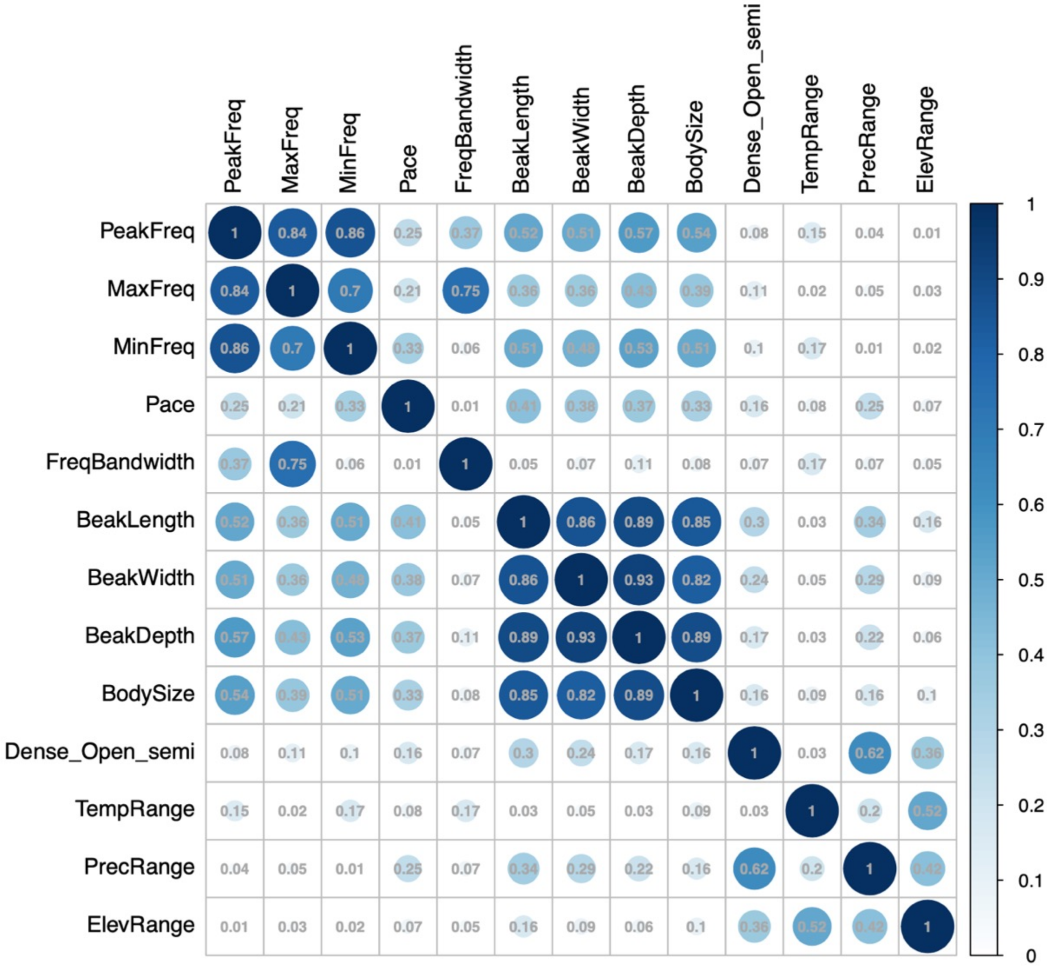


**Figure S1.** Correlation plot showing the absolute correlation between all song variables, morphological traits, and environmental variables included in this study. PeakFreq = mean song peak frequency (Hz), MaxFreq = mean song maximum frequency, MinFreq = mean song minimum frequency, Pace = mean song pace, FreqBandwidth = mean song frequency bandwidth, BeakLength = beak length in mm from AVONET, BeakWidth = beak width in mm from AVONET, BeakDepth = beak depth in mm from AVONET, BodySize = body mass in grams, Dense_Open_semi = Binary classification of habitat type for this study, TempRange = mean annual temperature extracted from each species range, PrecRange = mean annual total precipitation across species range, ElevRange = mean elevation across species range. See methods for further details.

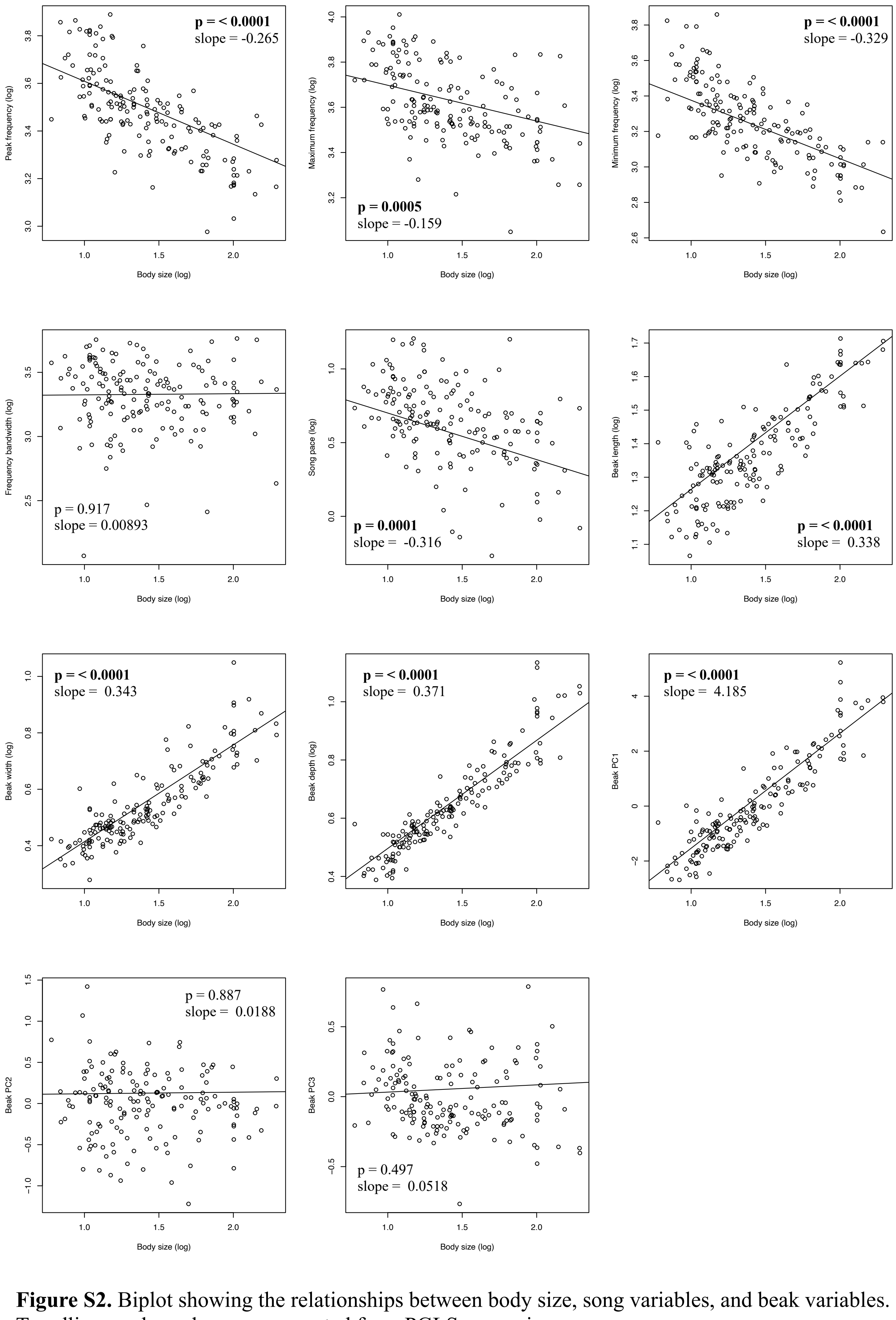


**Figure S2.** Biplot showing the relationships between body size, song variables, and beak variables. Trendlines and p-values are generated from PGLS regressions.
