## Supplementary material for "Body size shapes song in honeyeaters": File S1

Acanthagenys_rufogularis (n = 6)

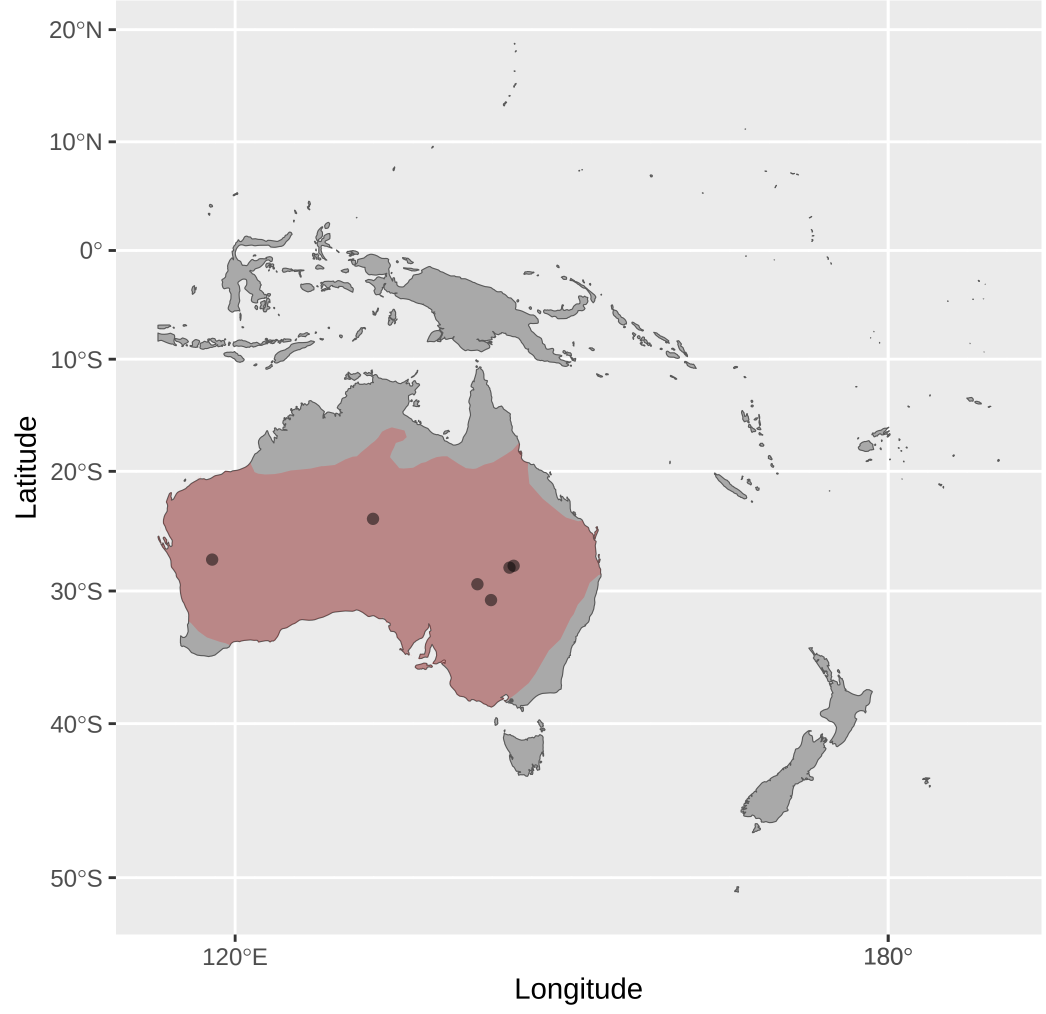


Anthochaera_chrysoptera (n = 4)

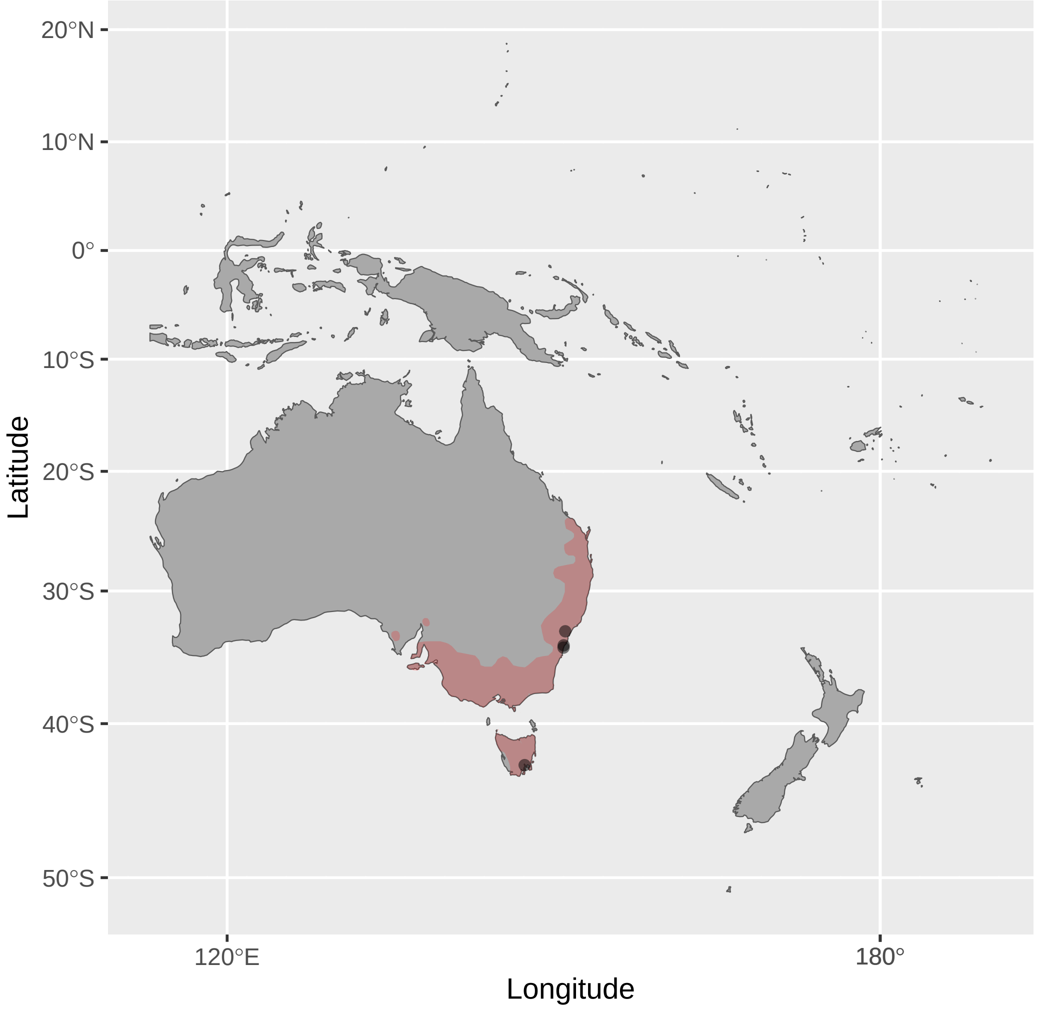


Anthornis_melanura (n = 14)

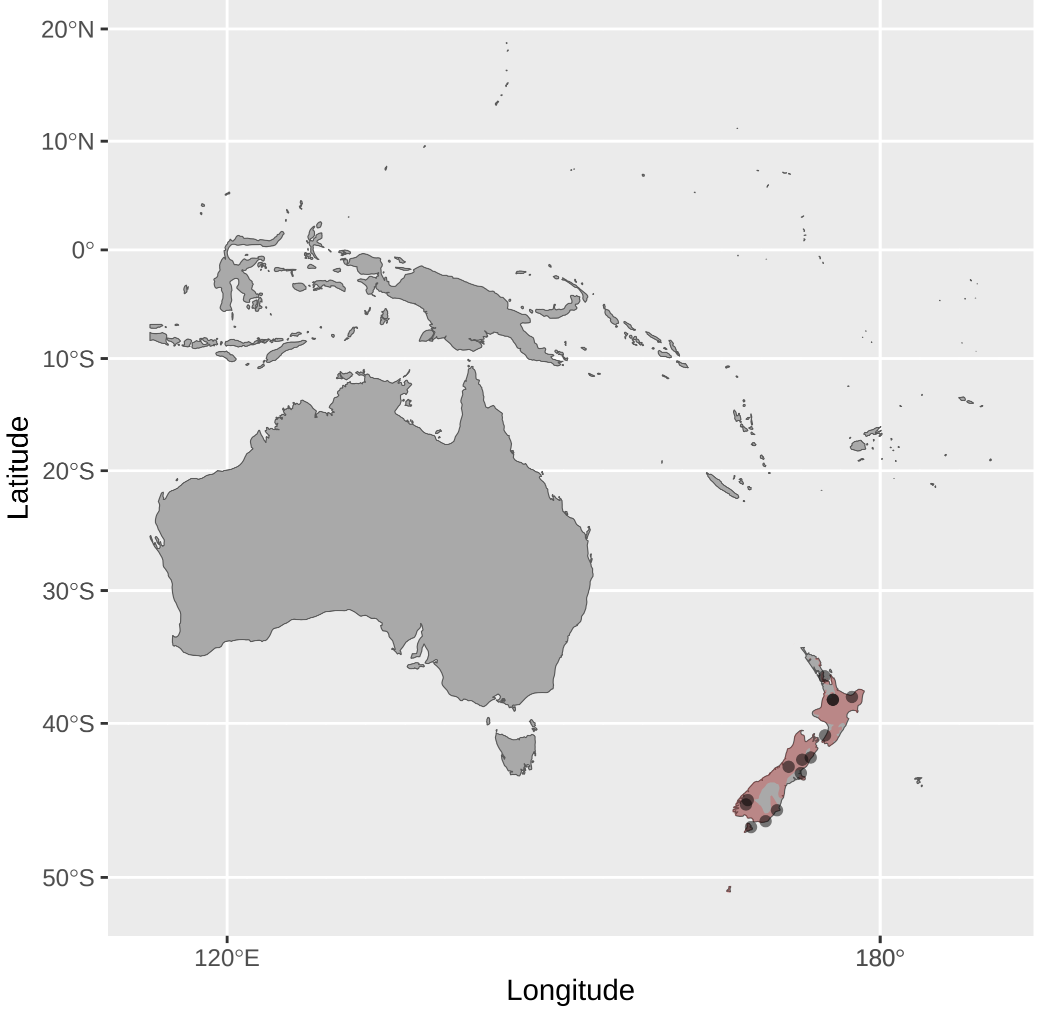


Acanthorhynchus_superciliosus (n = 2)

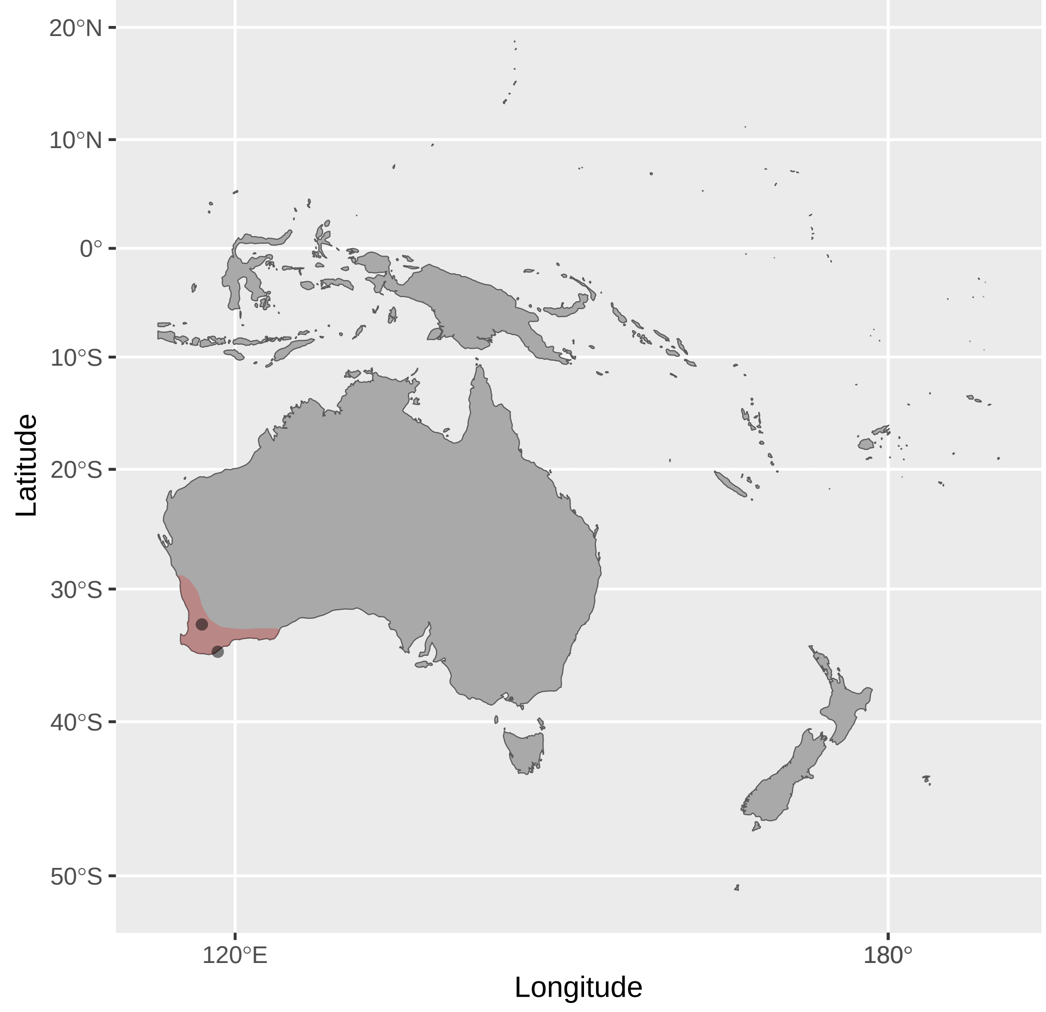


Anthochaera_lunulata (n = 4)

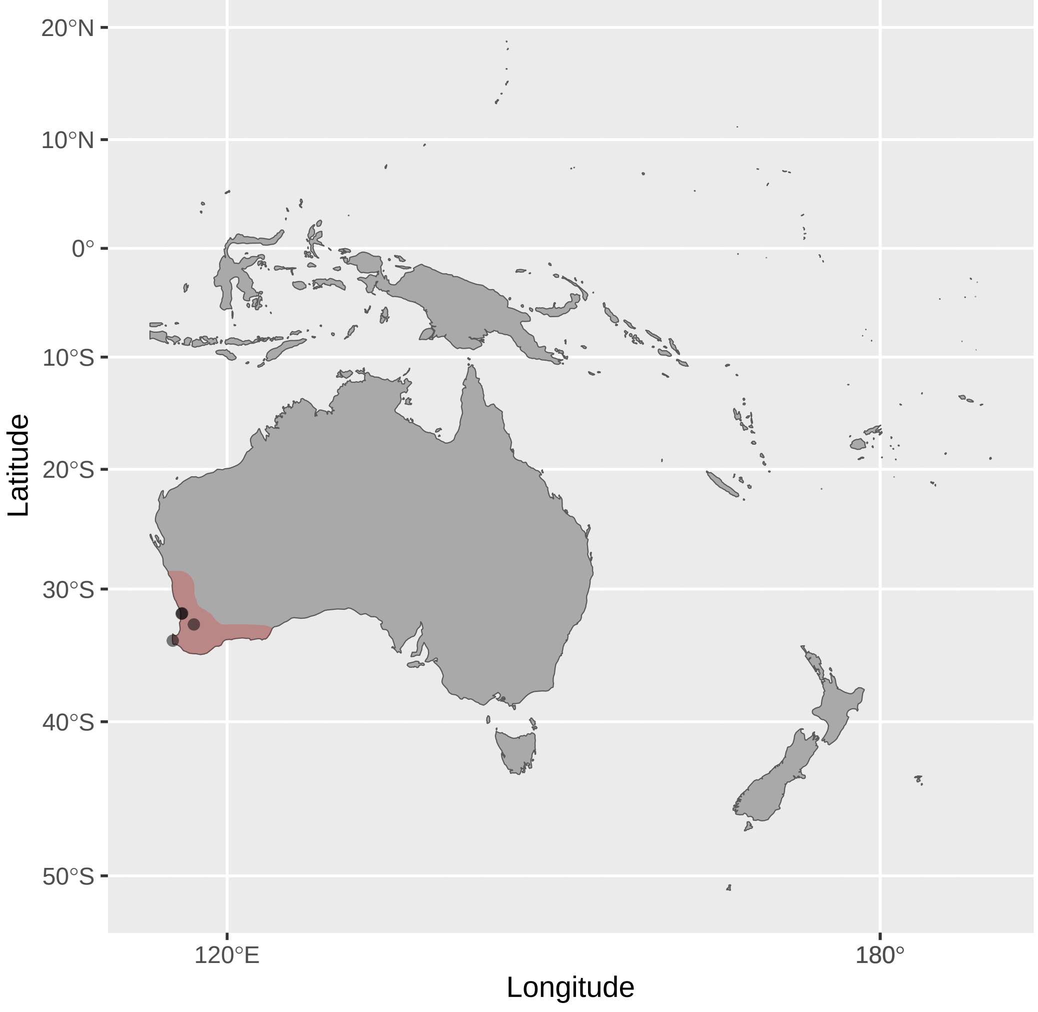


Ashbyia_lovensis (n = 1)

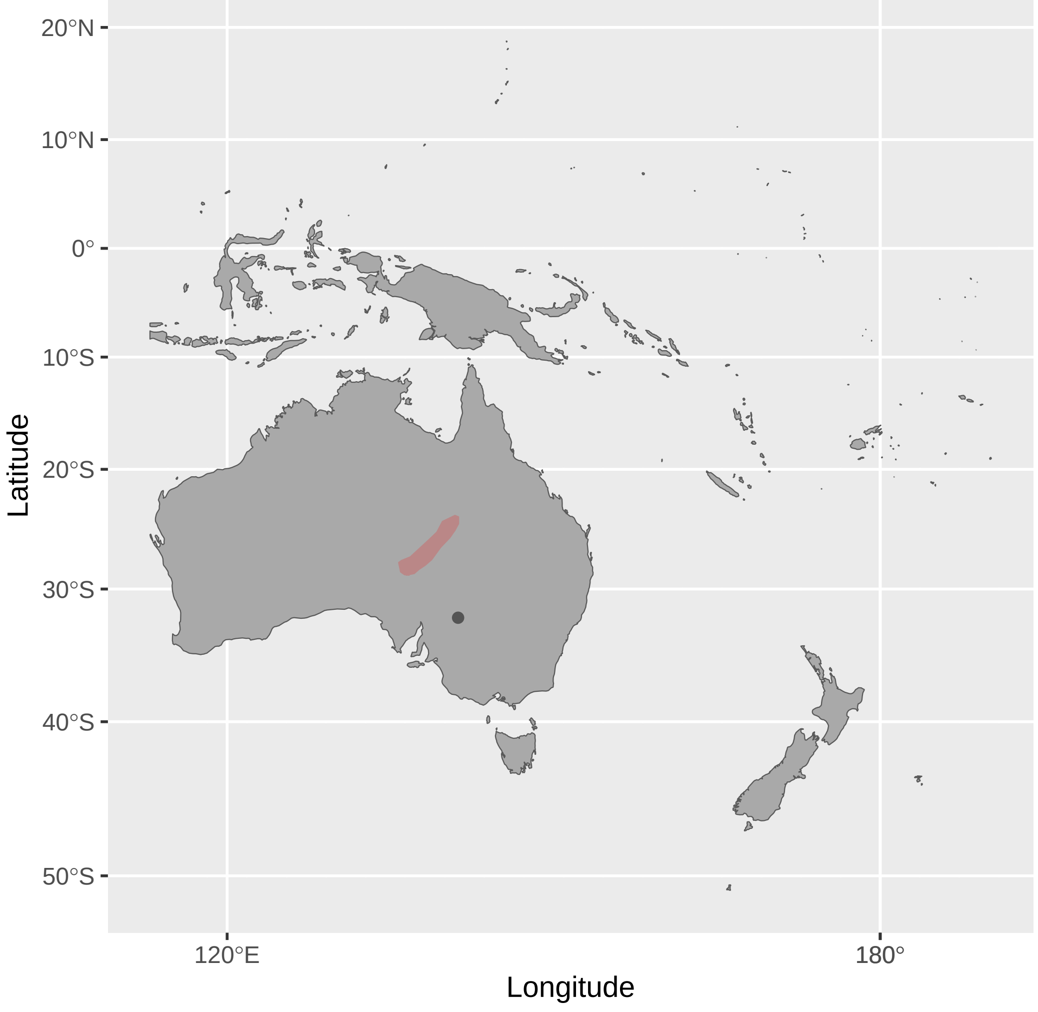


Acanthorhynchus_tenuirostris (n = 9)

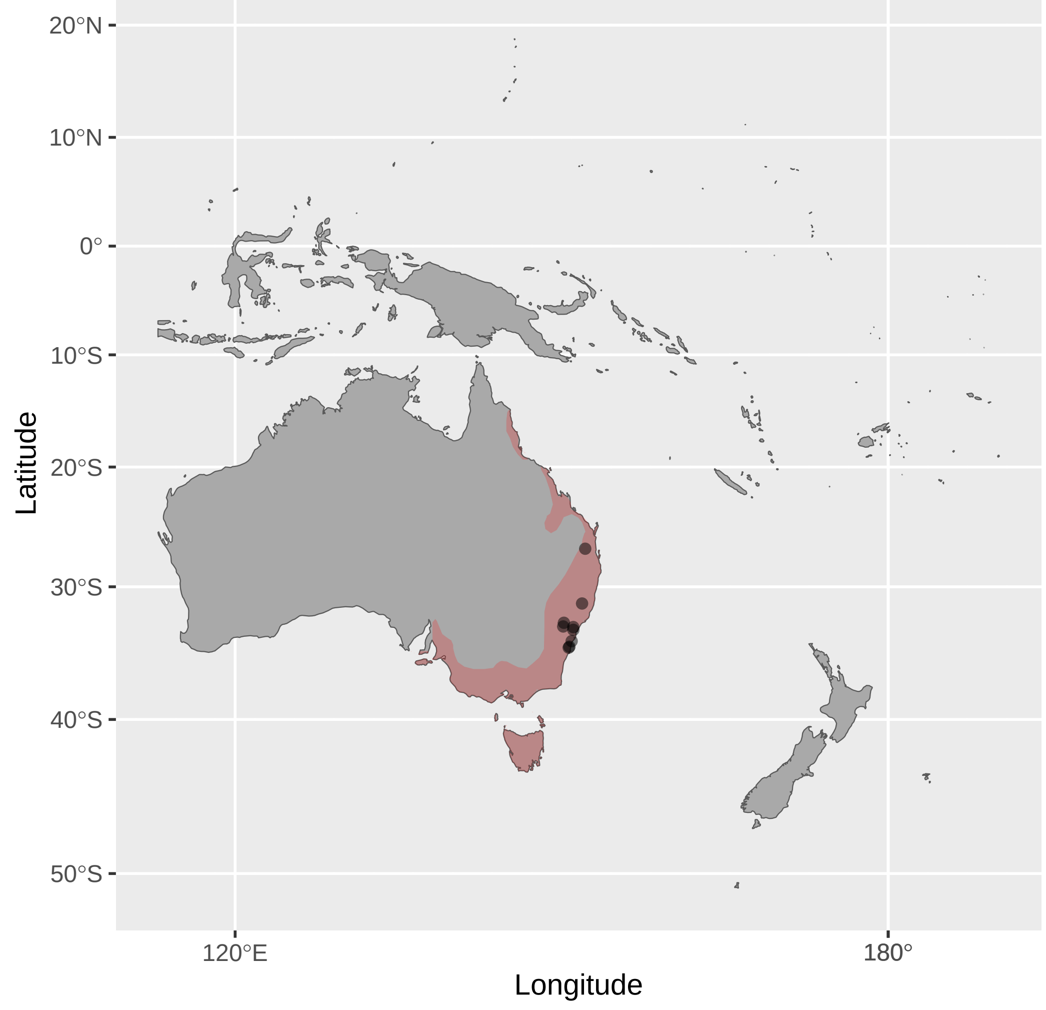


Anthochaera_paradoxa (n = 2)

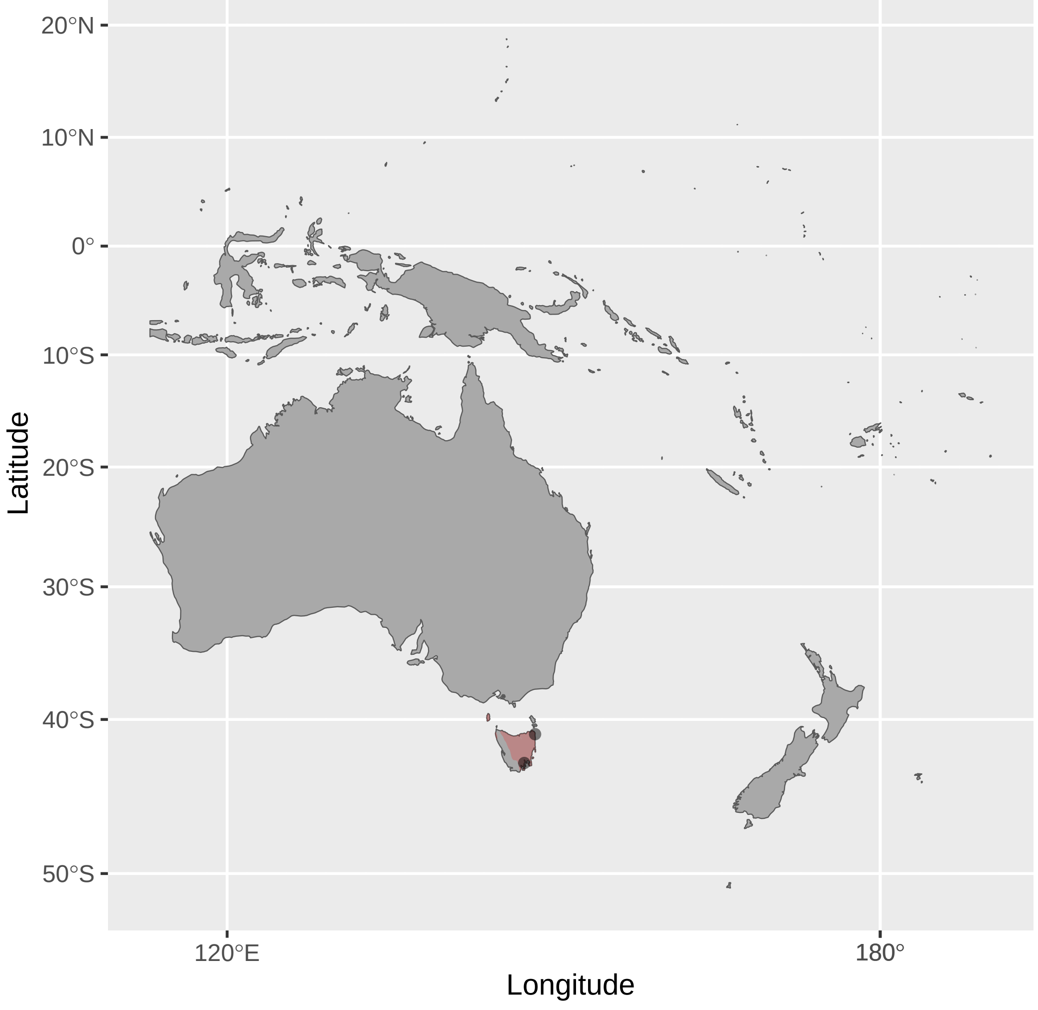


Bolemoreus_frenatus (n = 7)

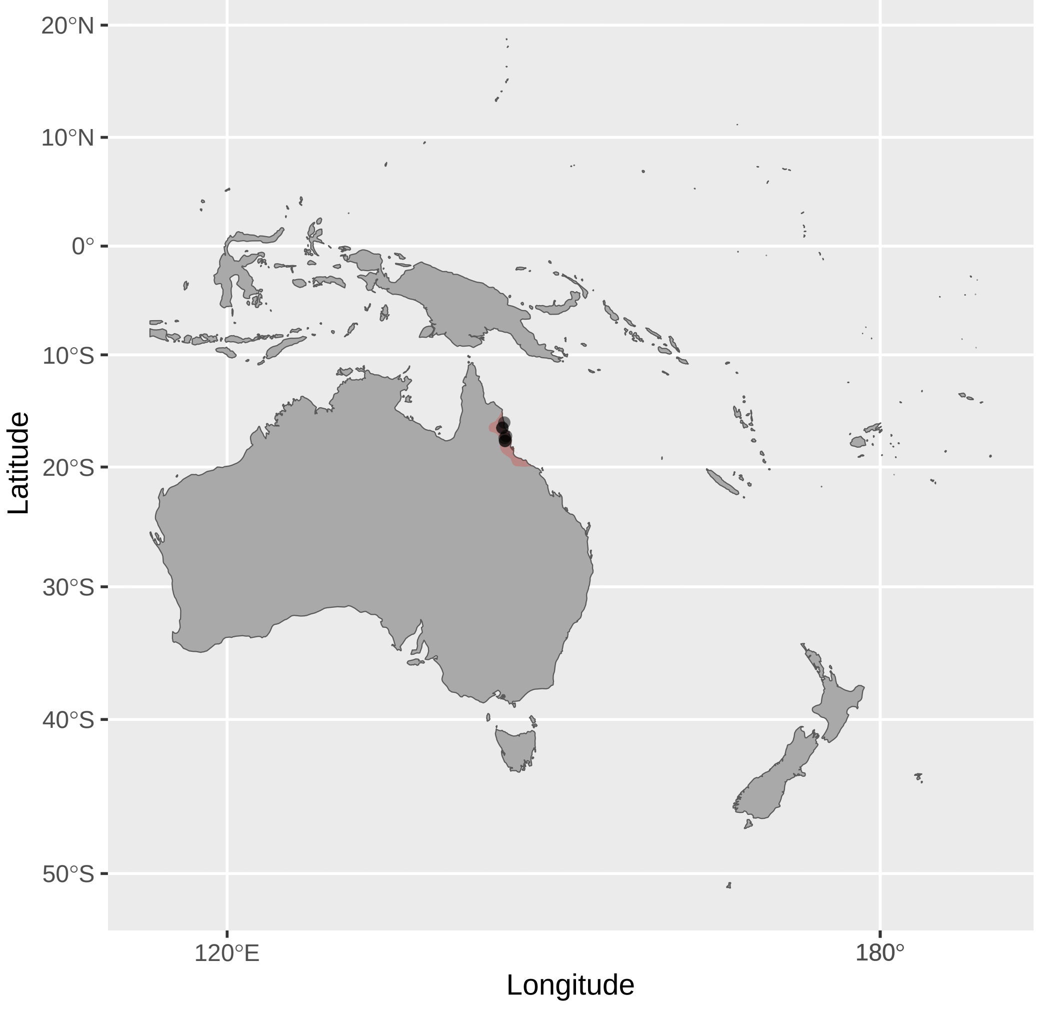


Anthochaera_carunculata (n = 4)

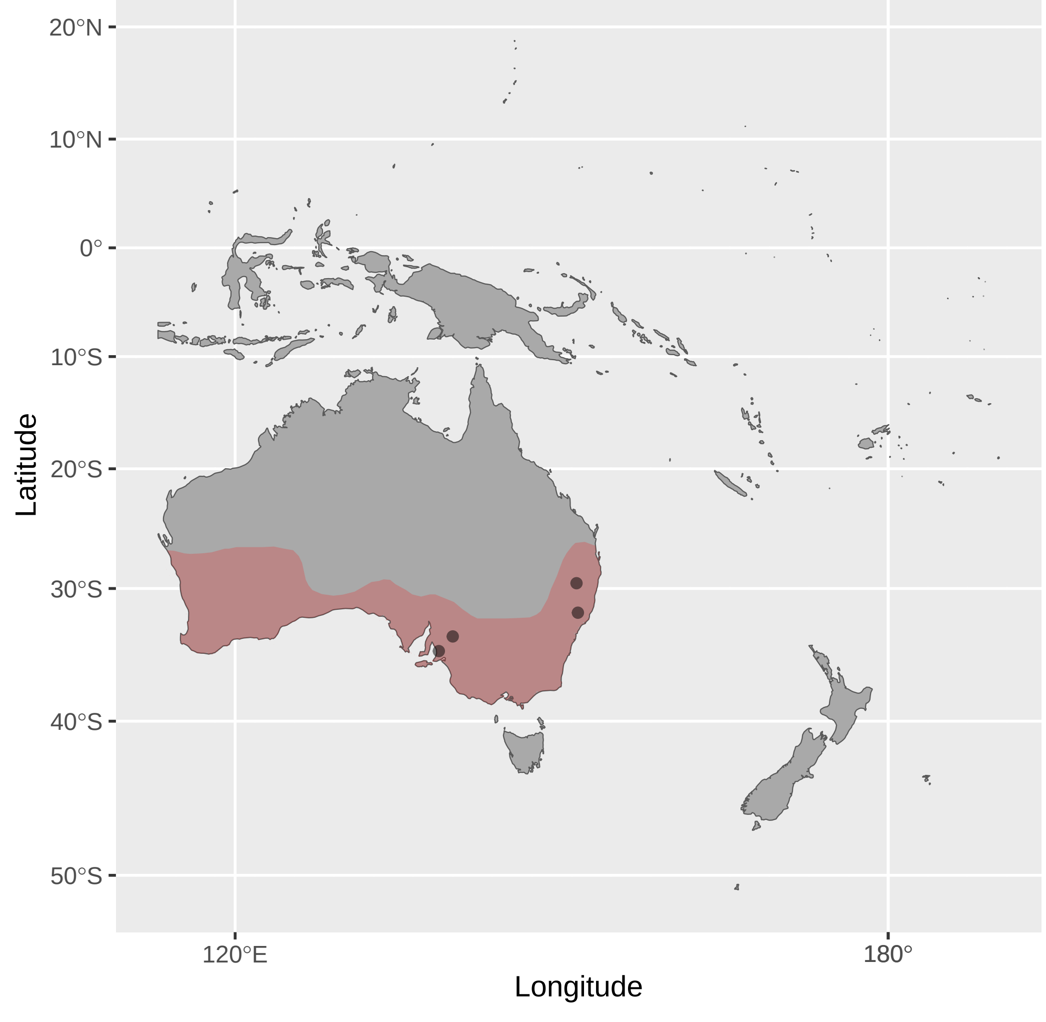


Anthochaera_phrygia (n = 2)

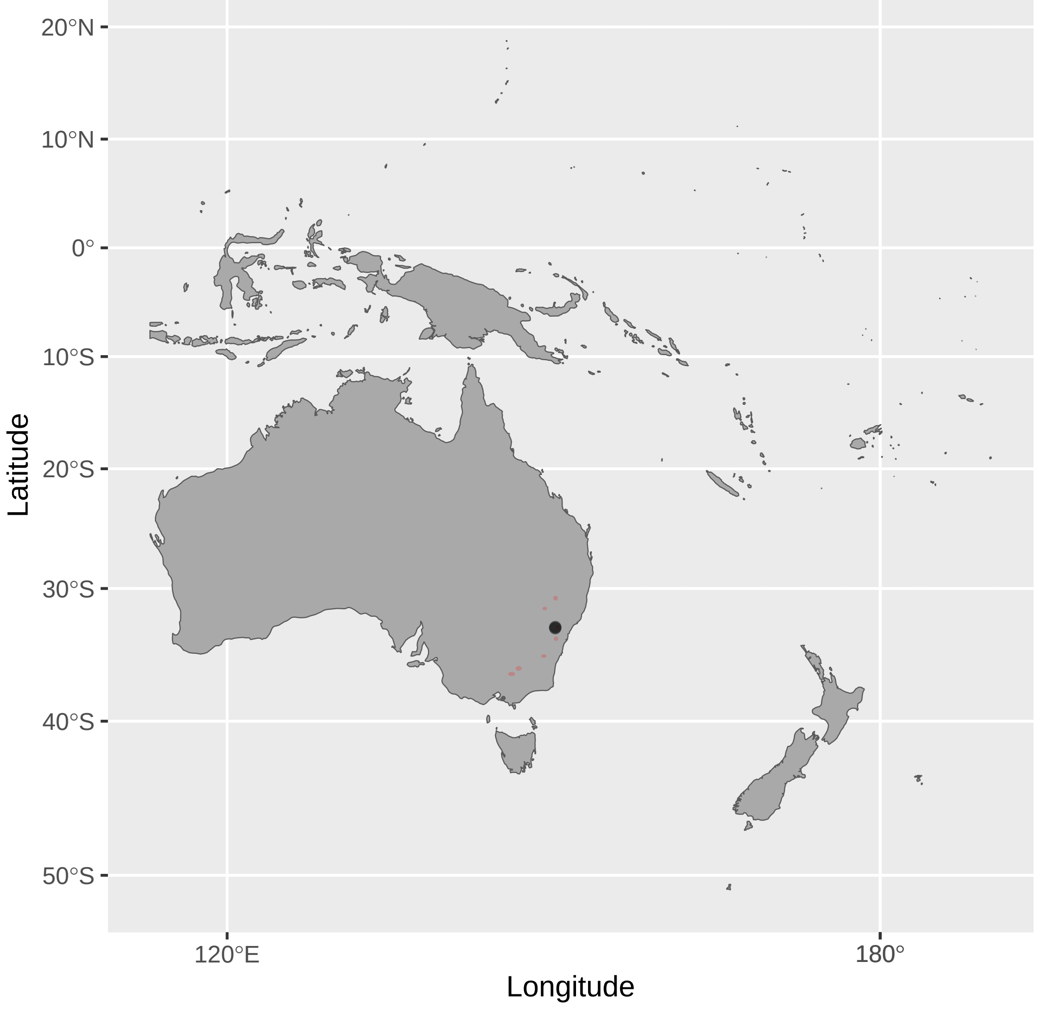


Bolemoreus_hindwoodi (n = 2)

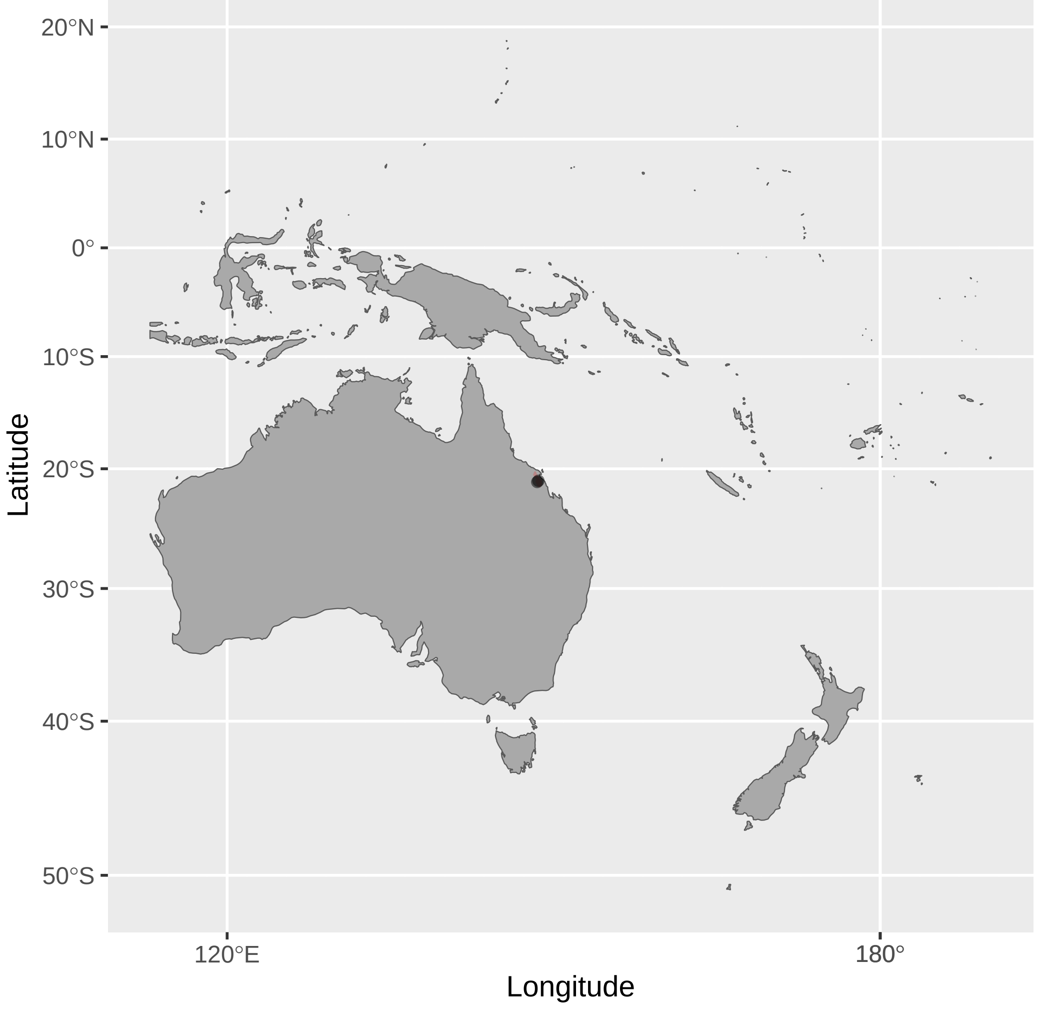

Caligavis_chrysops (n = 16)

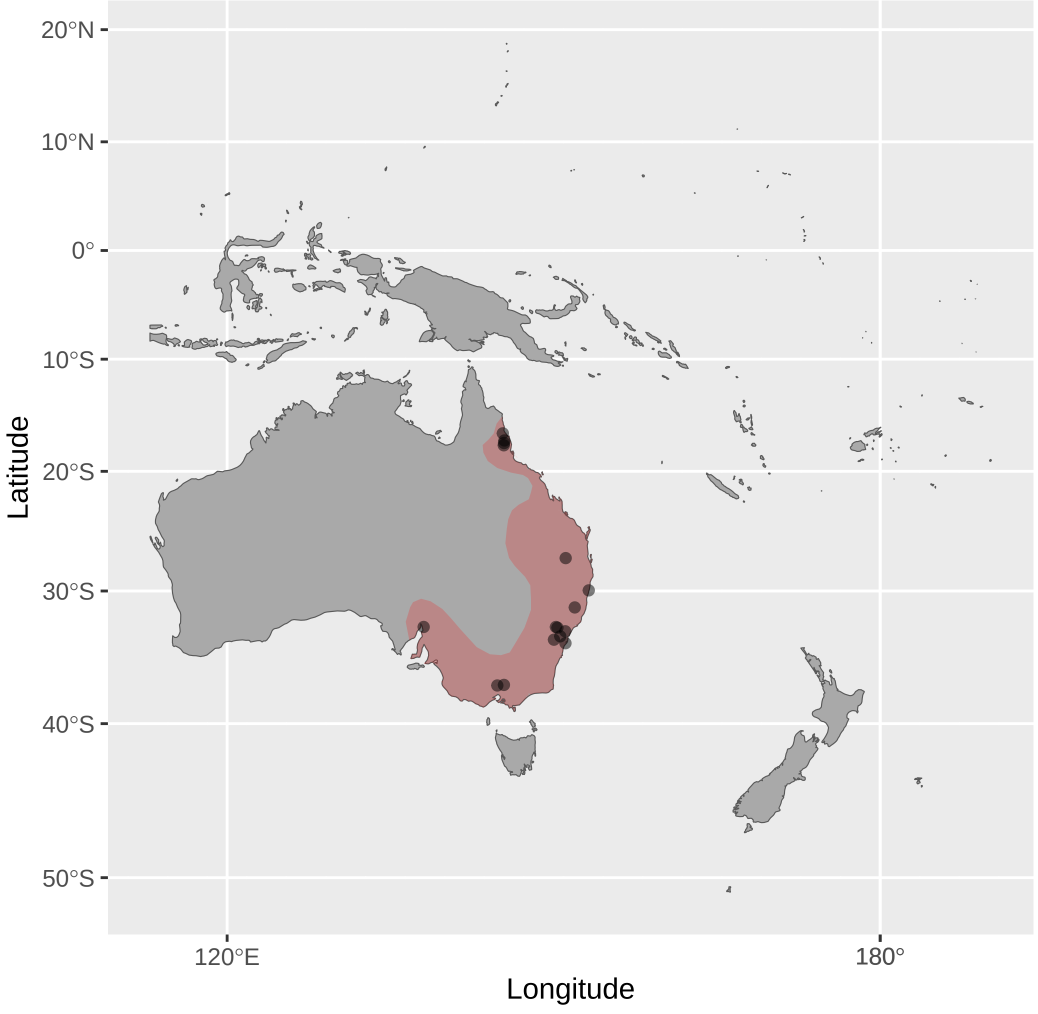


Caligavis_obscura (n = 2)

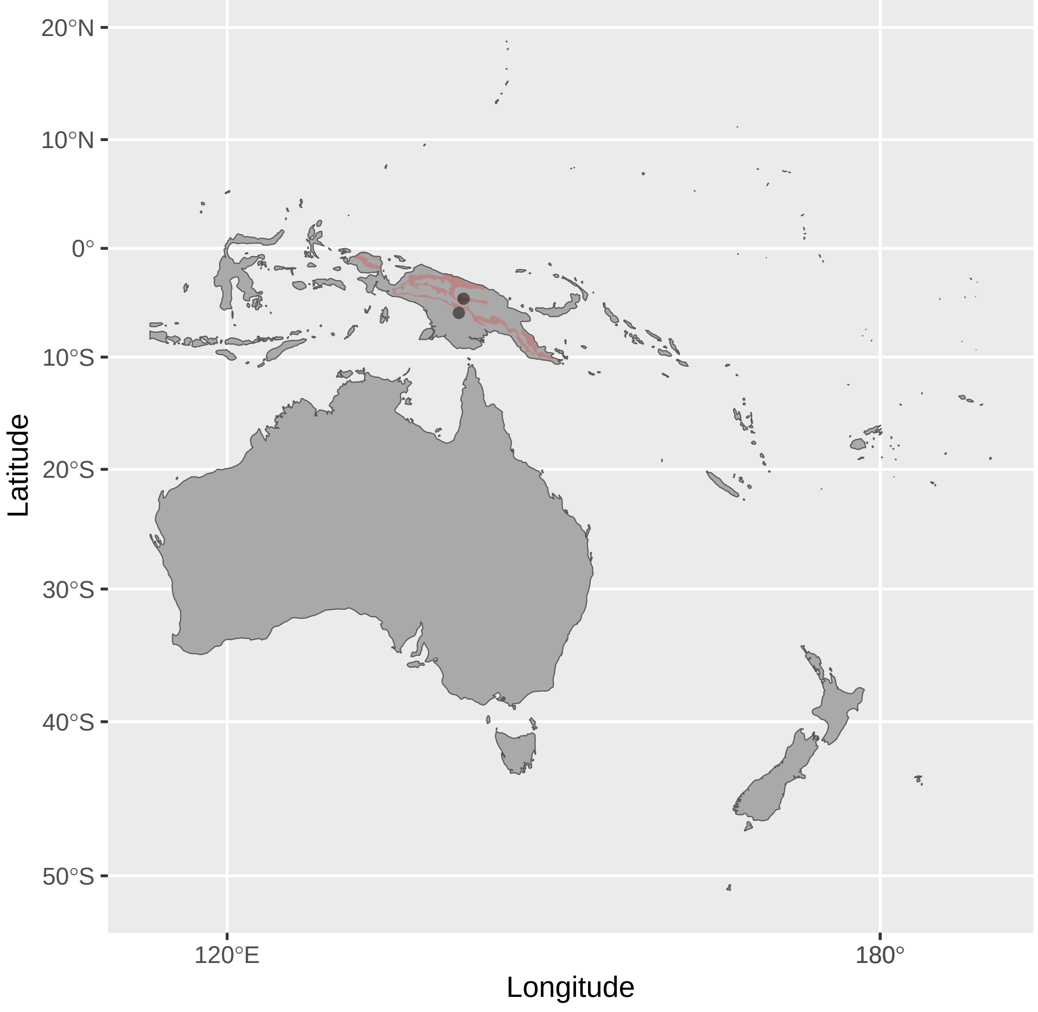


Caligavis_subfrenata (n = 5)

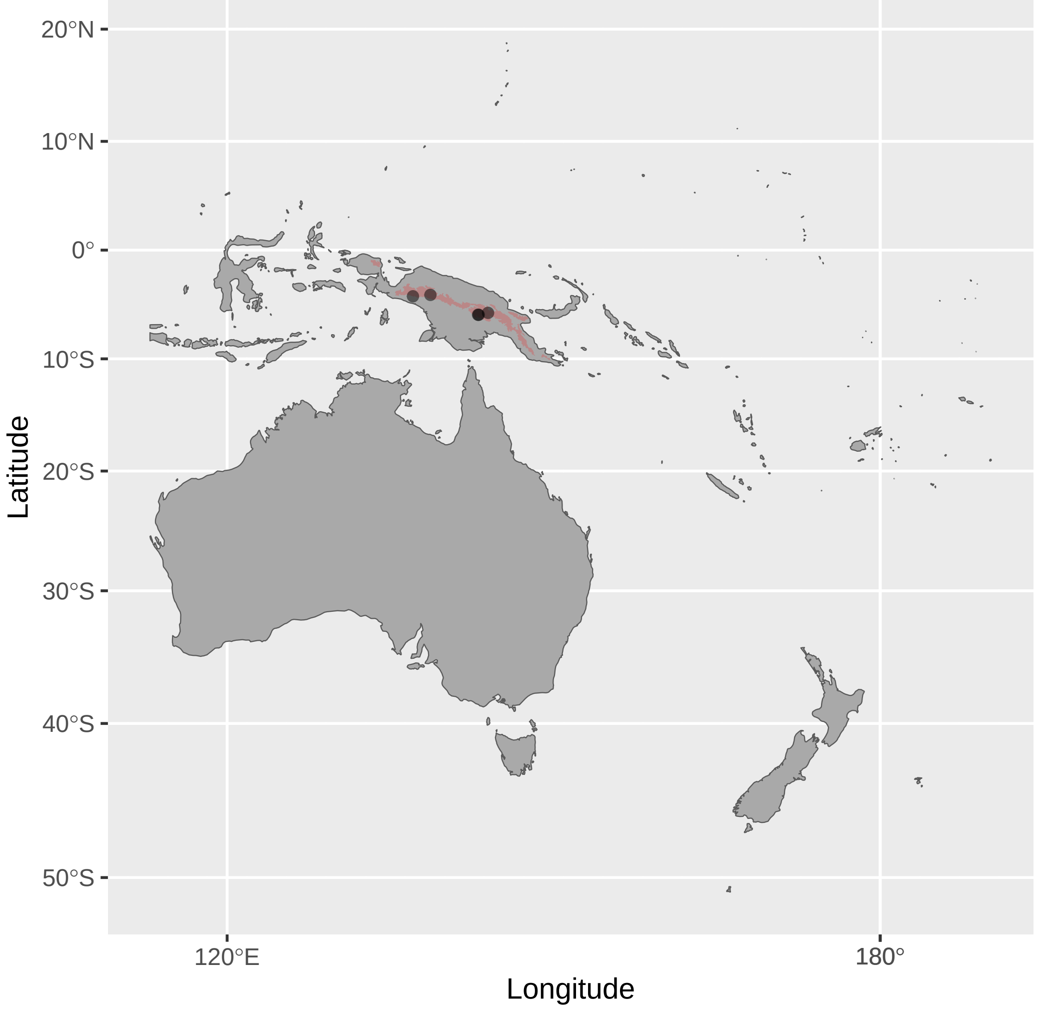


Certhionyx_variegatus (n = 6)

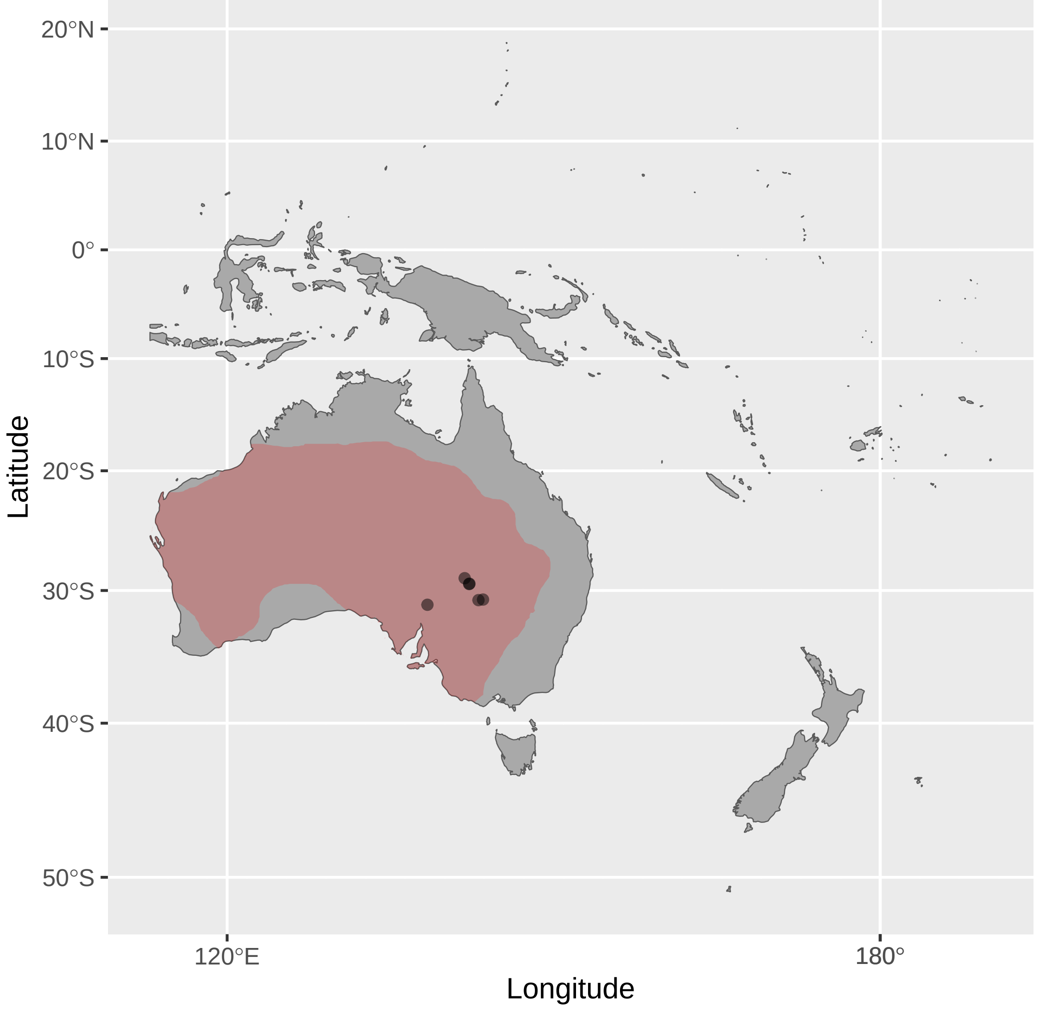


Cissomela_pectoralis (n = 1)

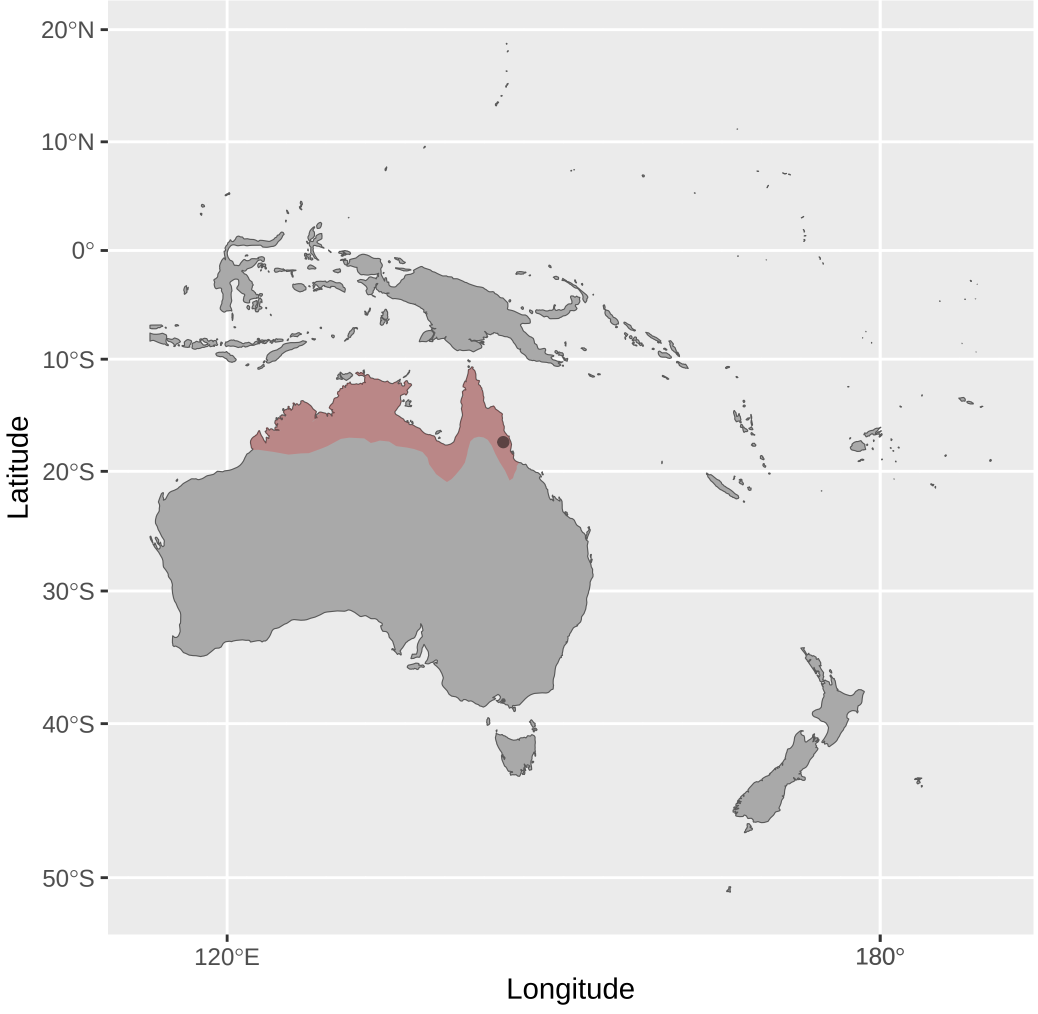


Conopophila_albogularis (n = 2)

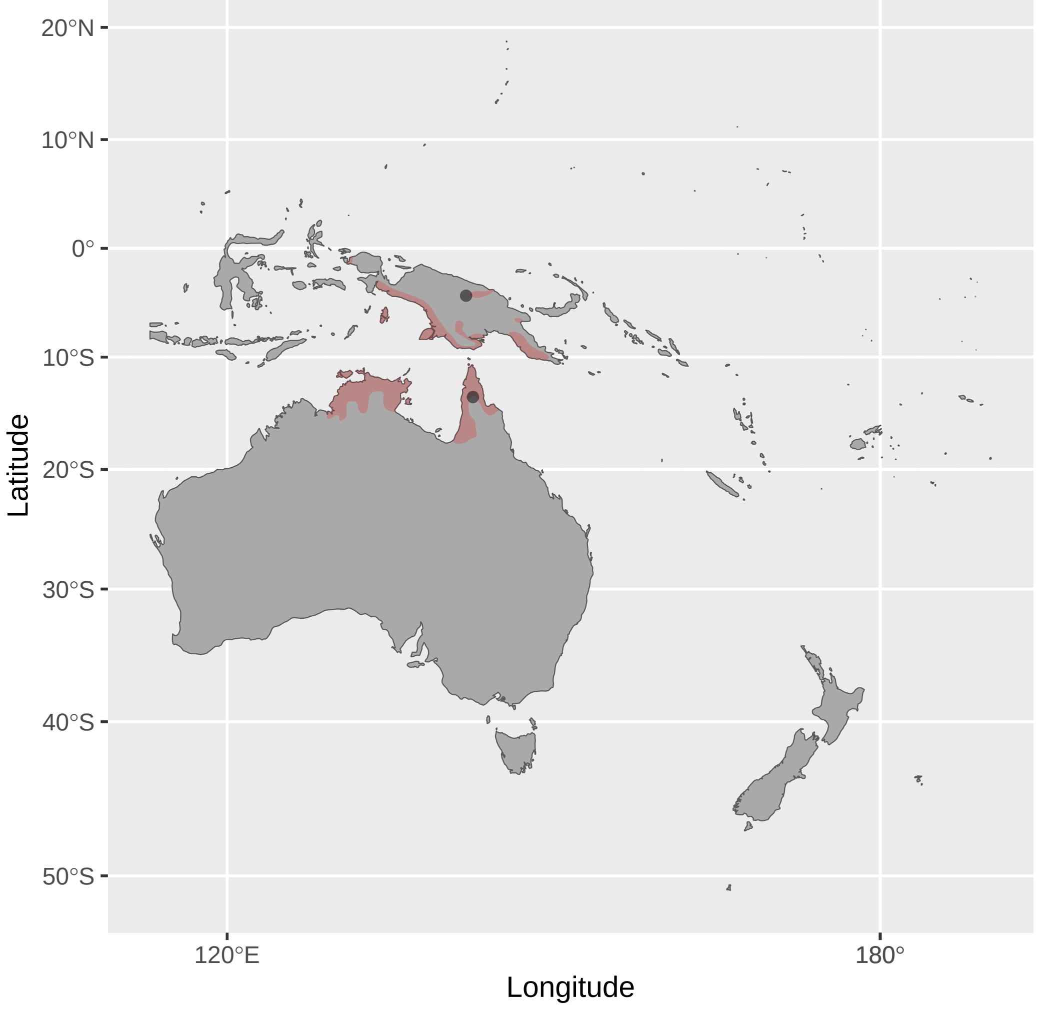


Conopophila_rufogularis (n = 1)

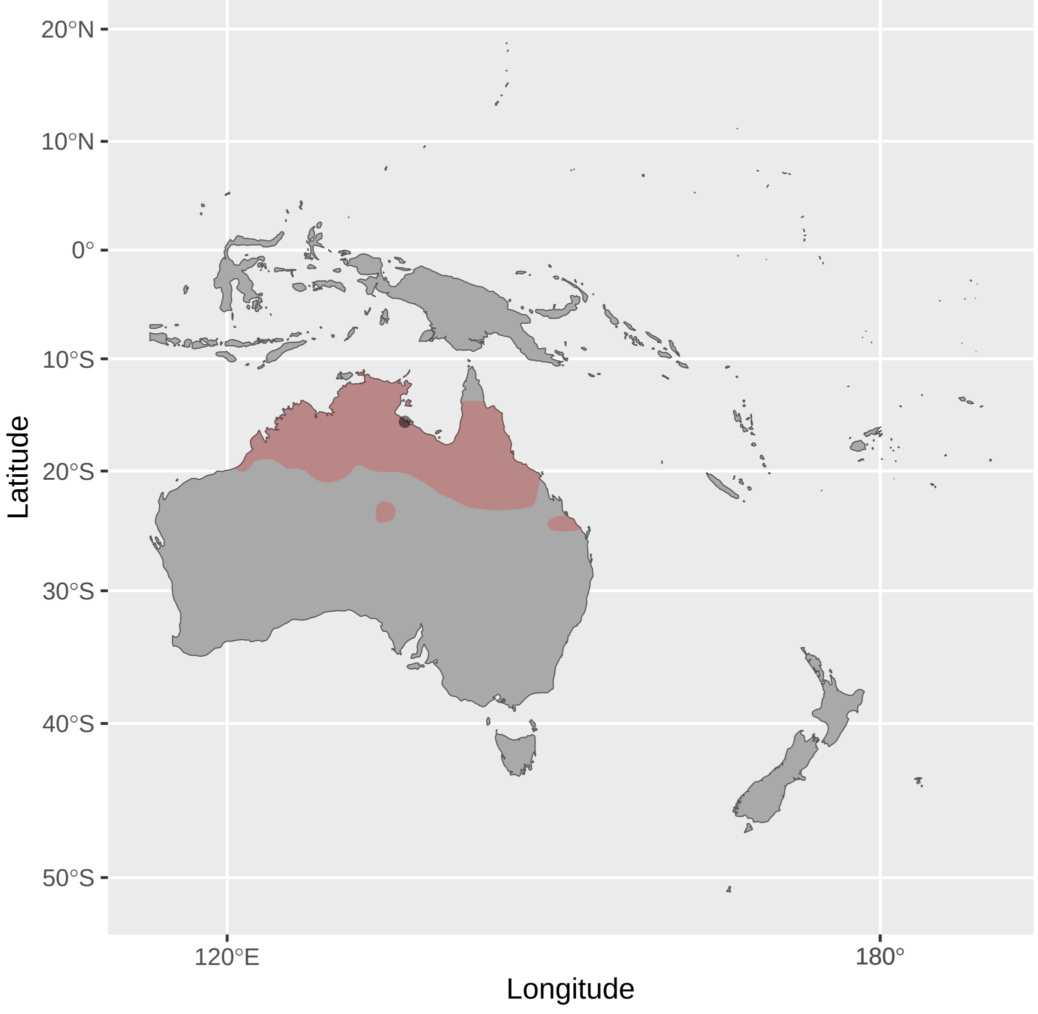


Conopophila_whitei (n = 1)

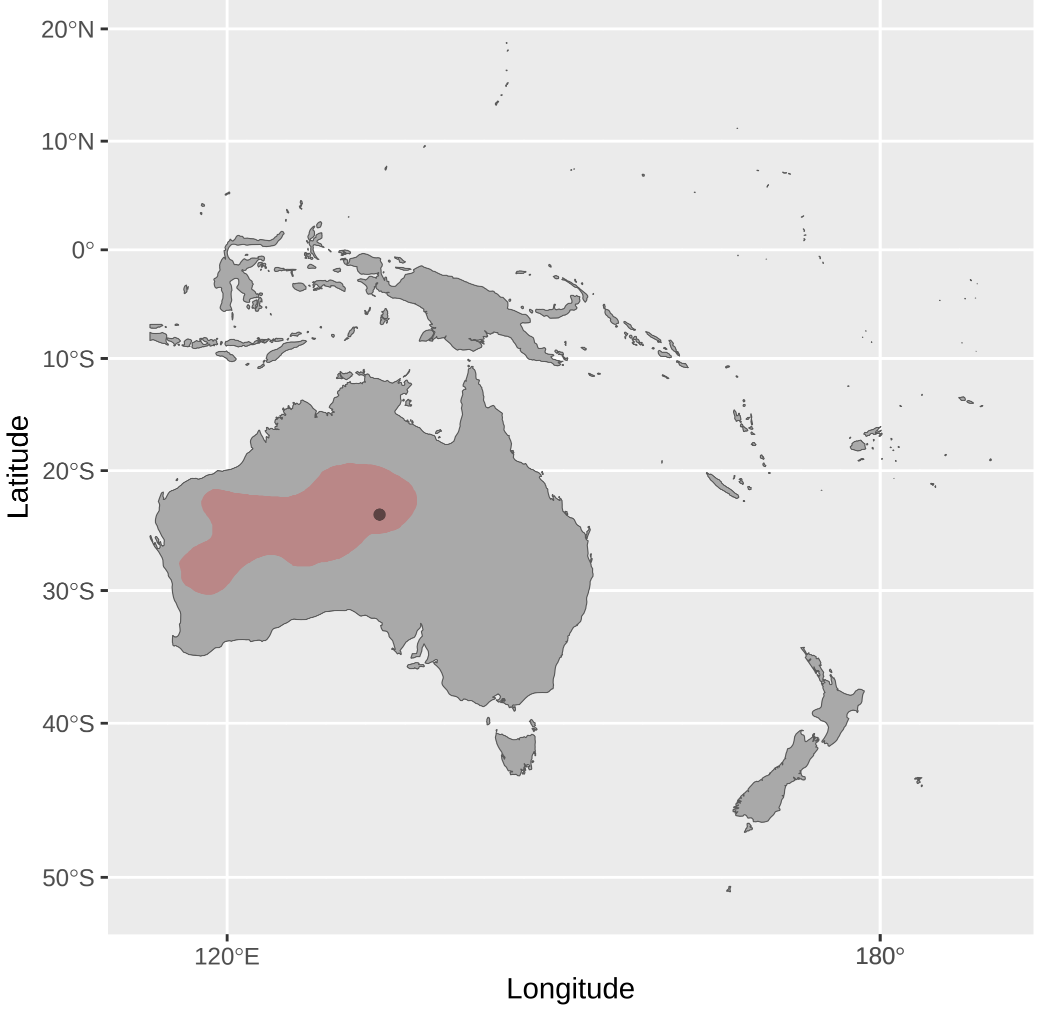


Entomyzon_cyanotis (n = 7)

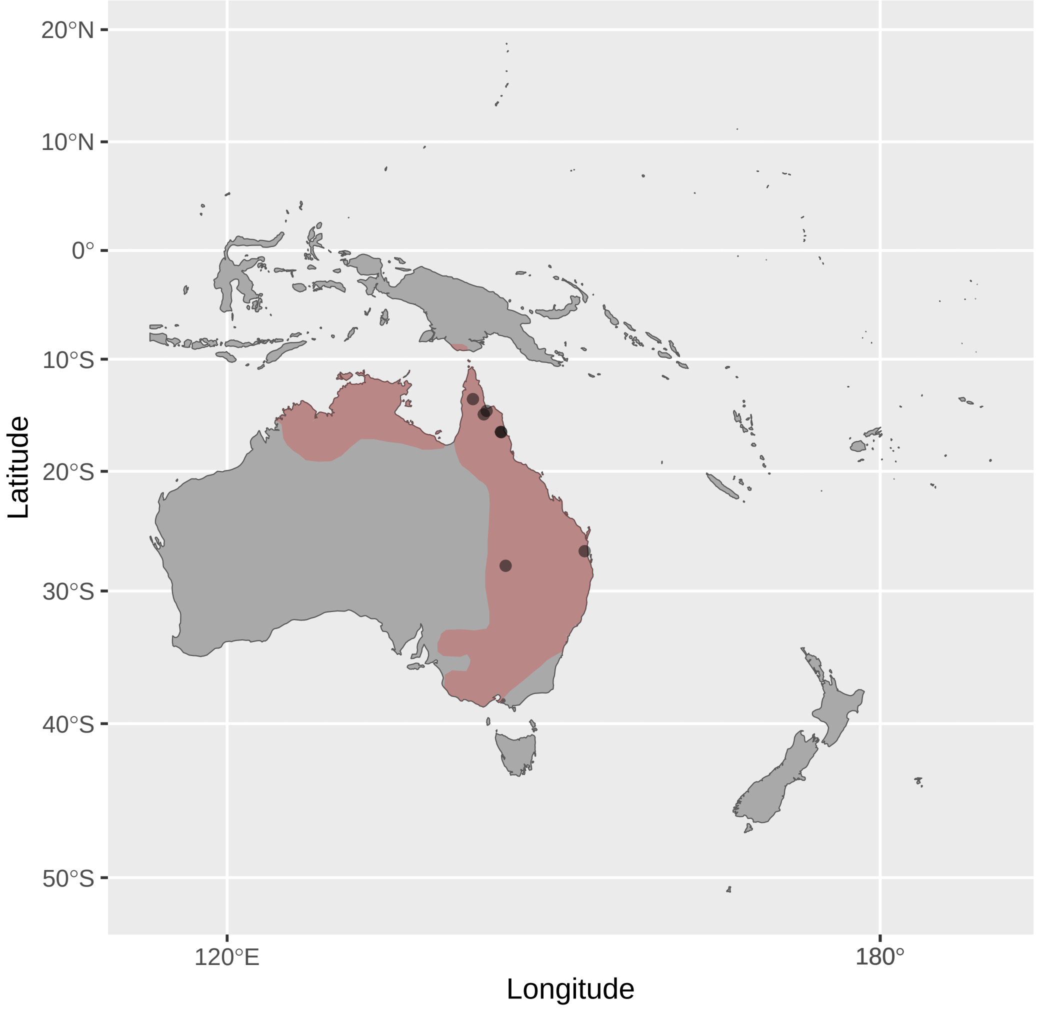


Epthianura_albifrons (n = 3)

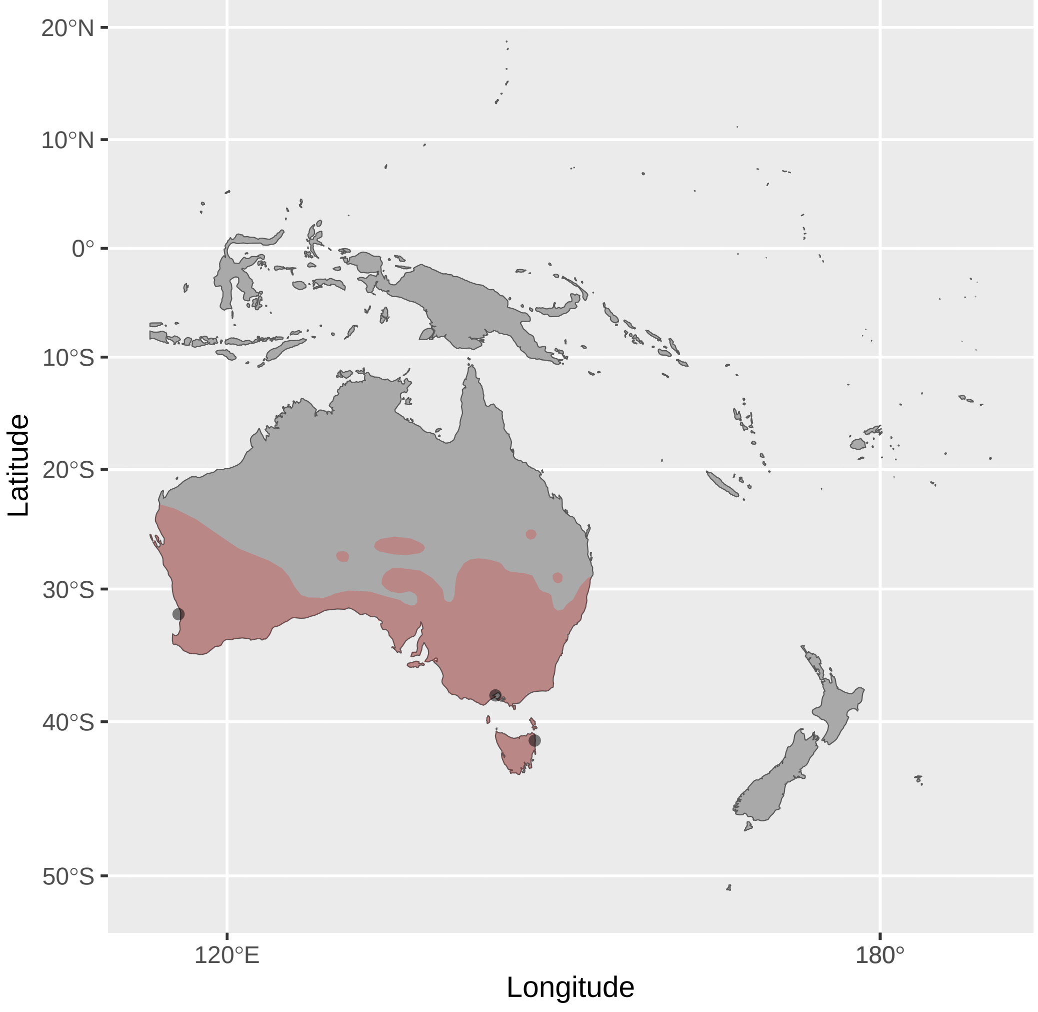


Epthianura_crocea (n = 2)

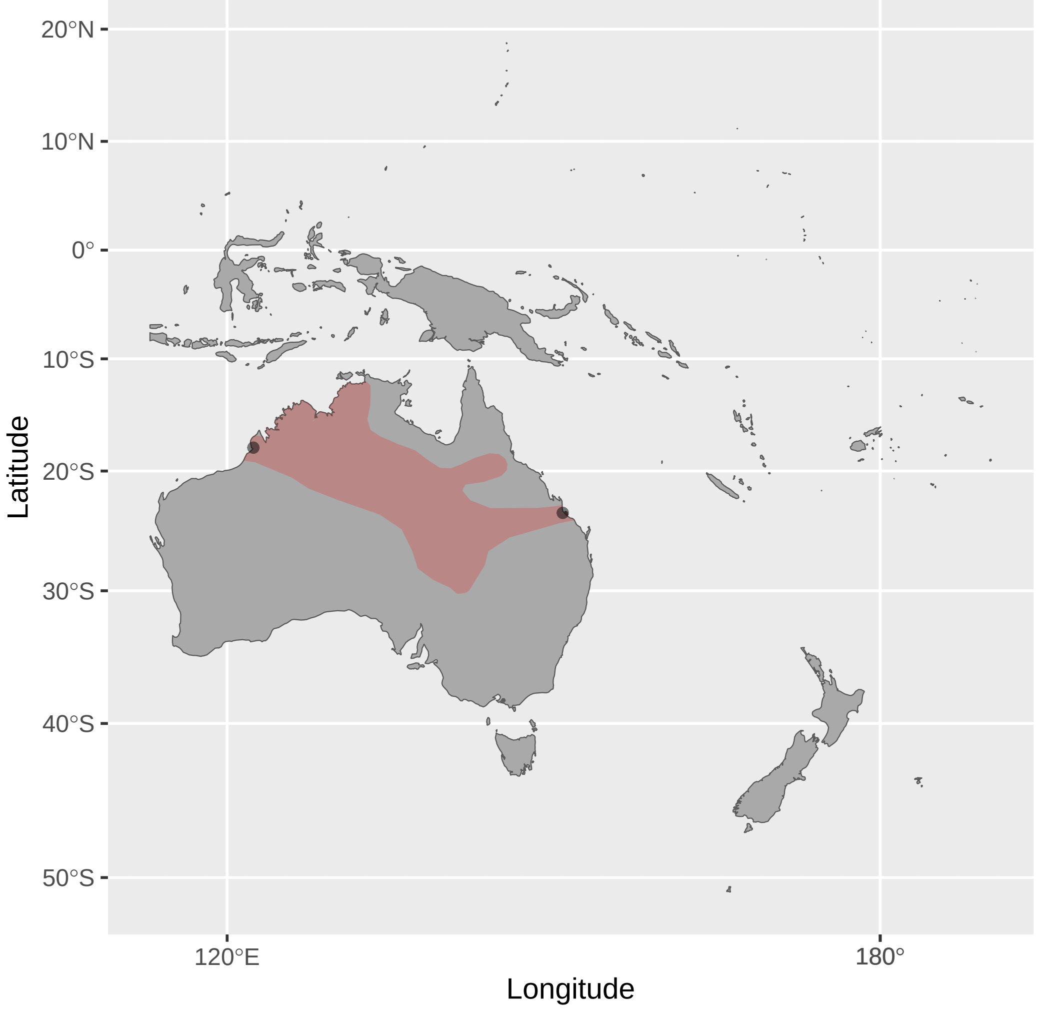


Epthianura_tricolor (n = 4)

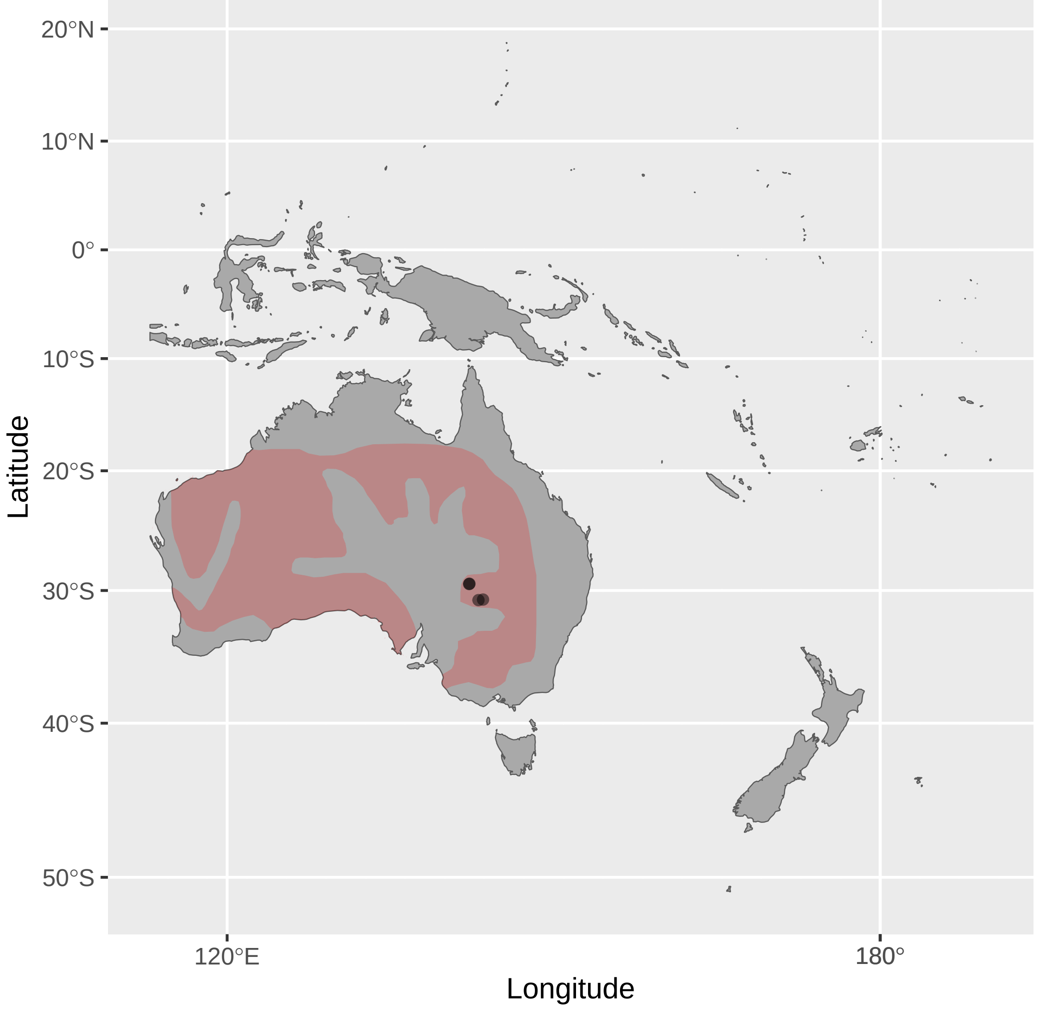

Foulehaio_carunculatus (n = 4)

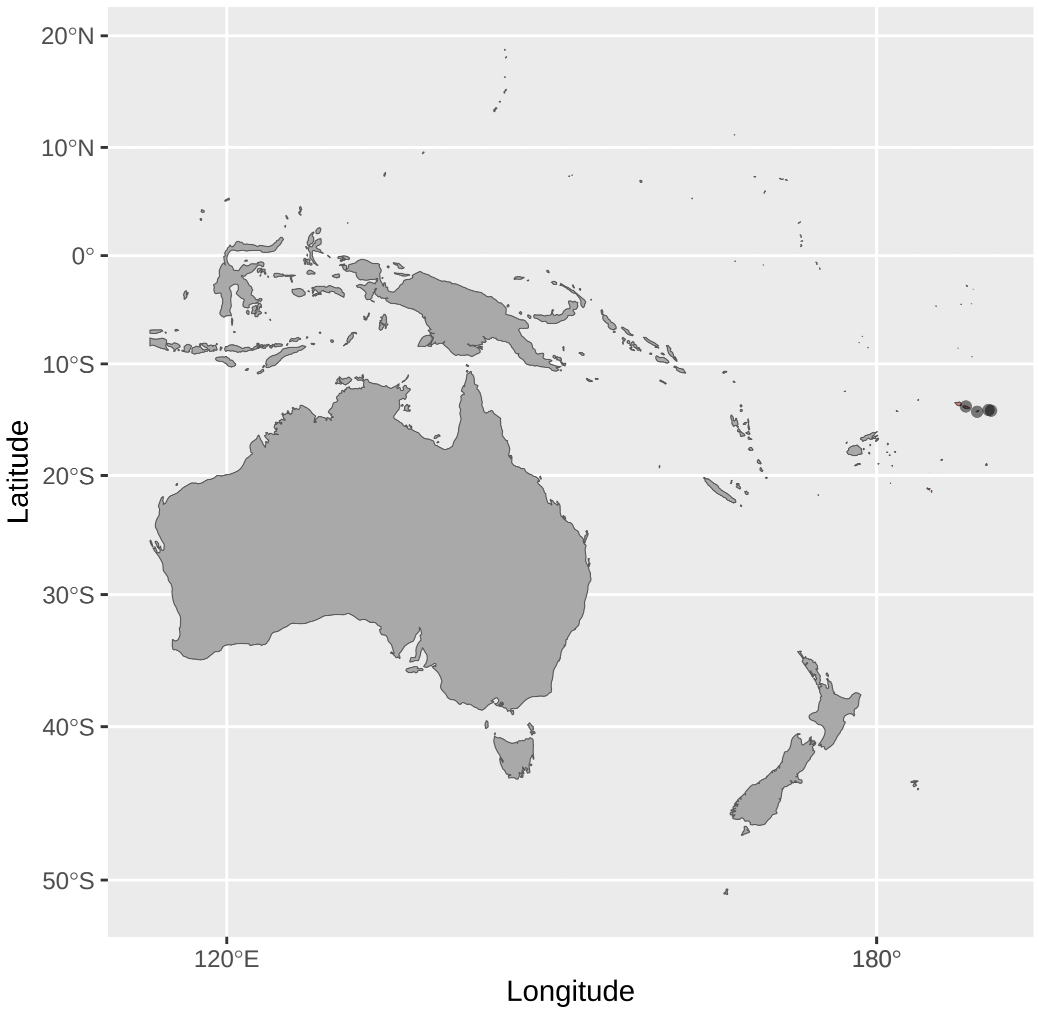


Gavicalis_versicolor (n = 5)

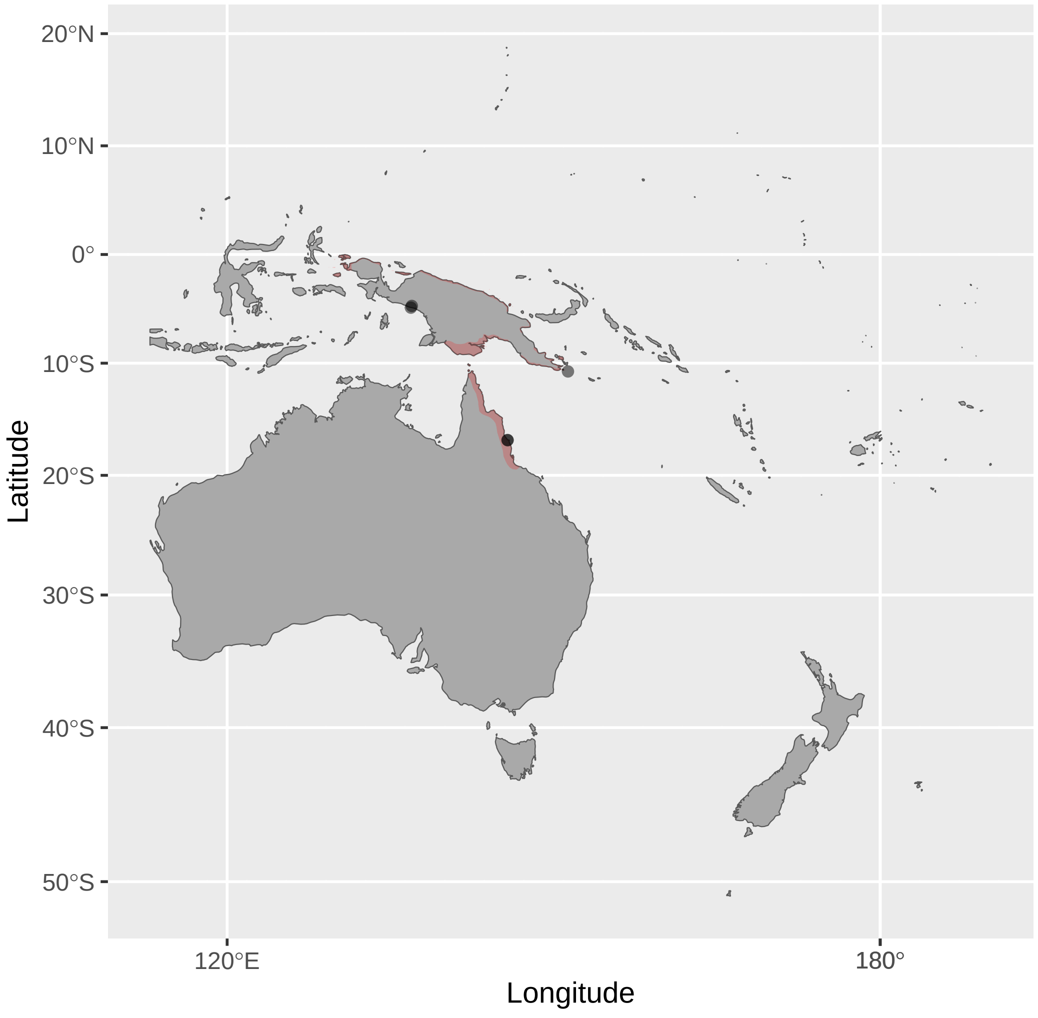


Glycifohia_notabilis (n = 1)

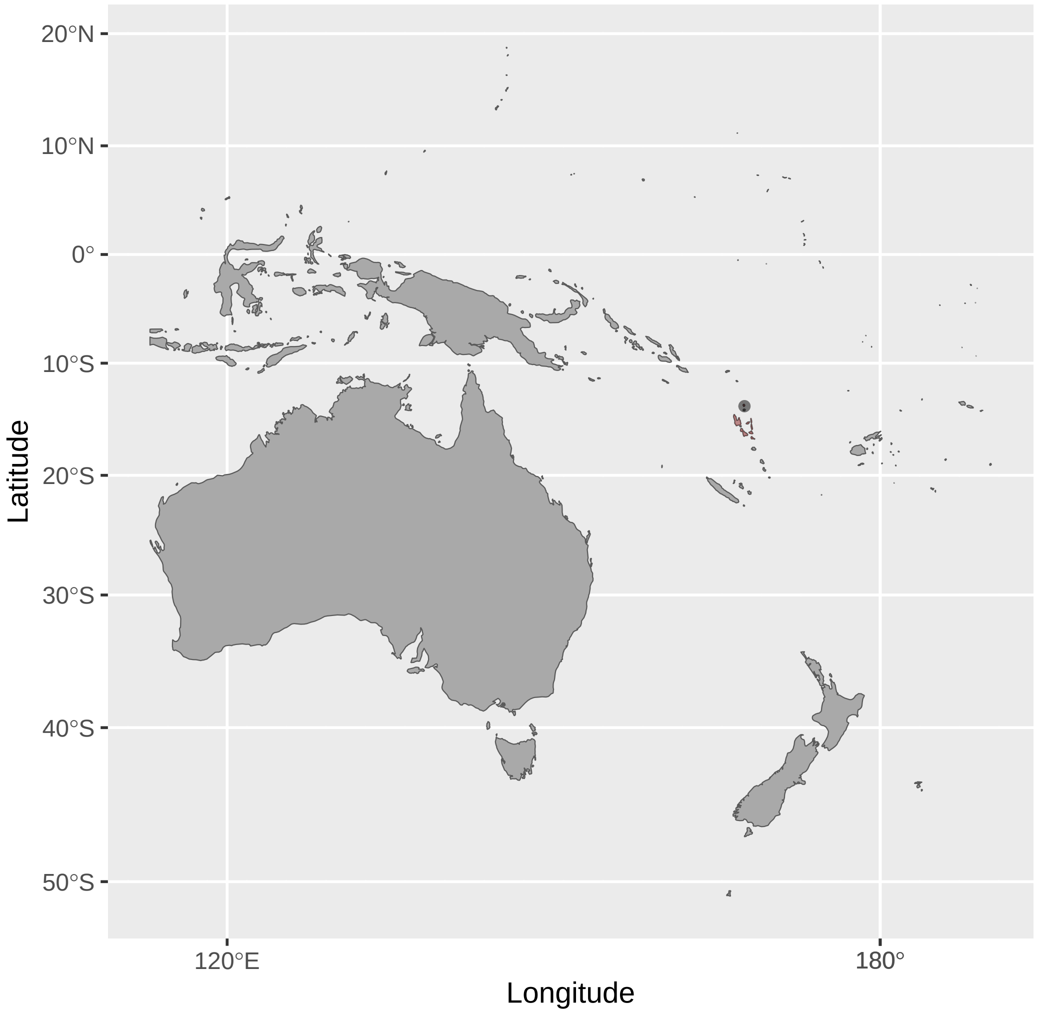


Foulehaio_procerior (n = 5)

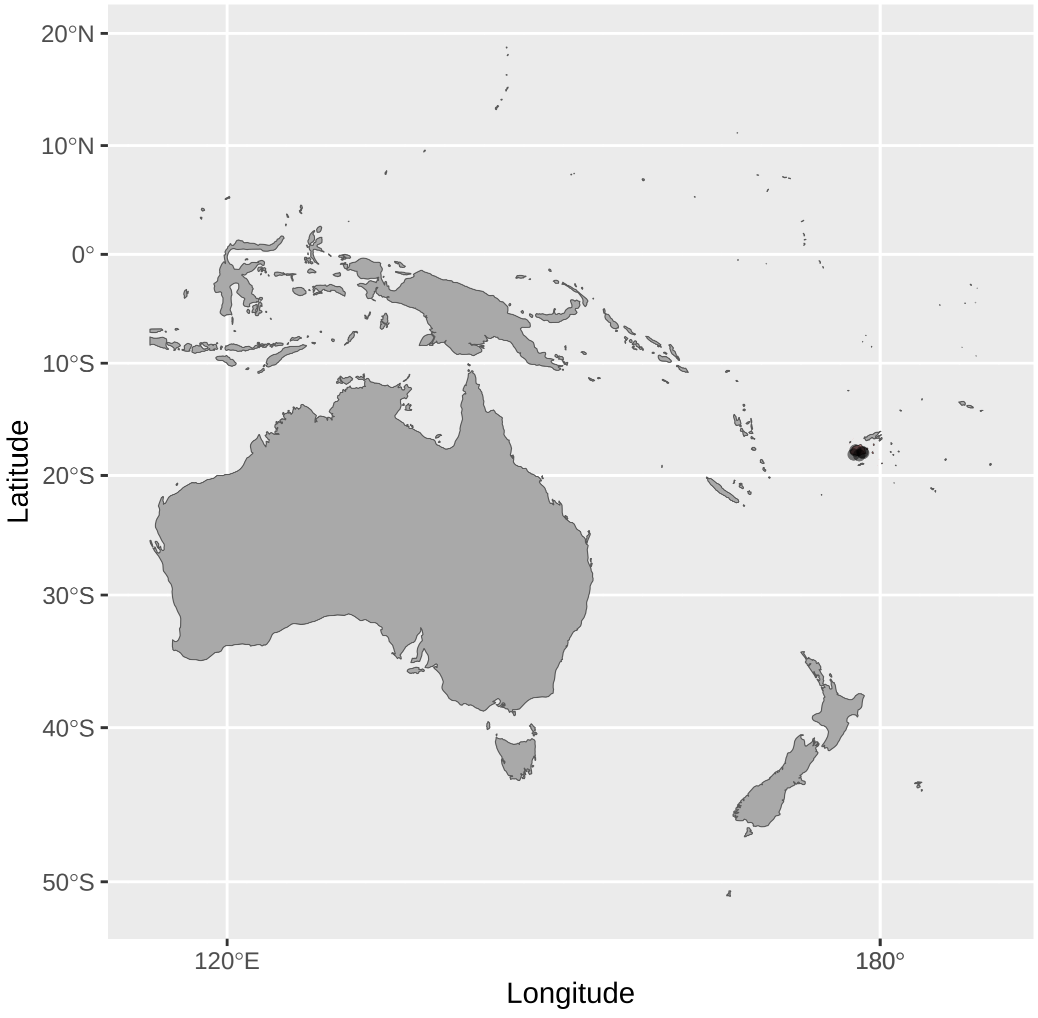


Gavicalis_virescens (n = 5)

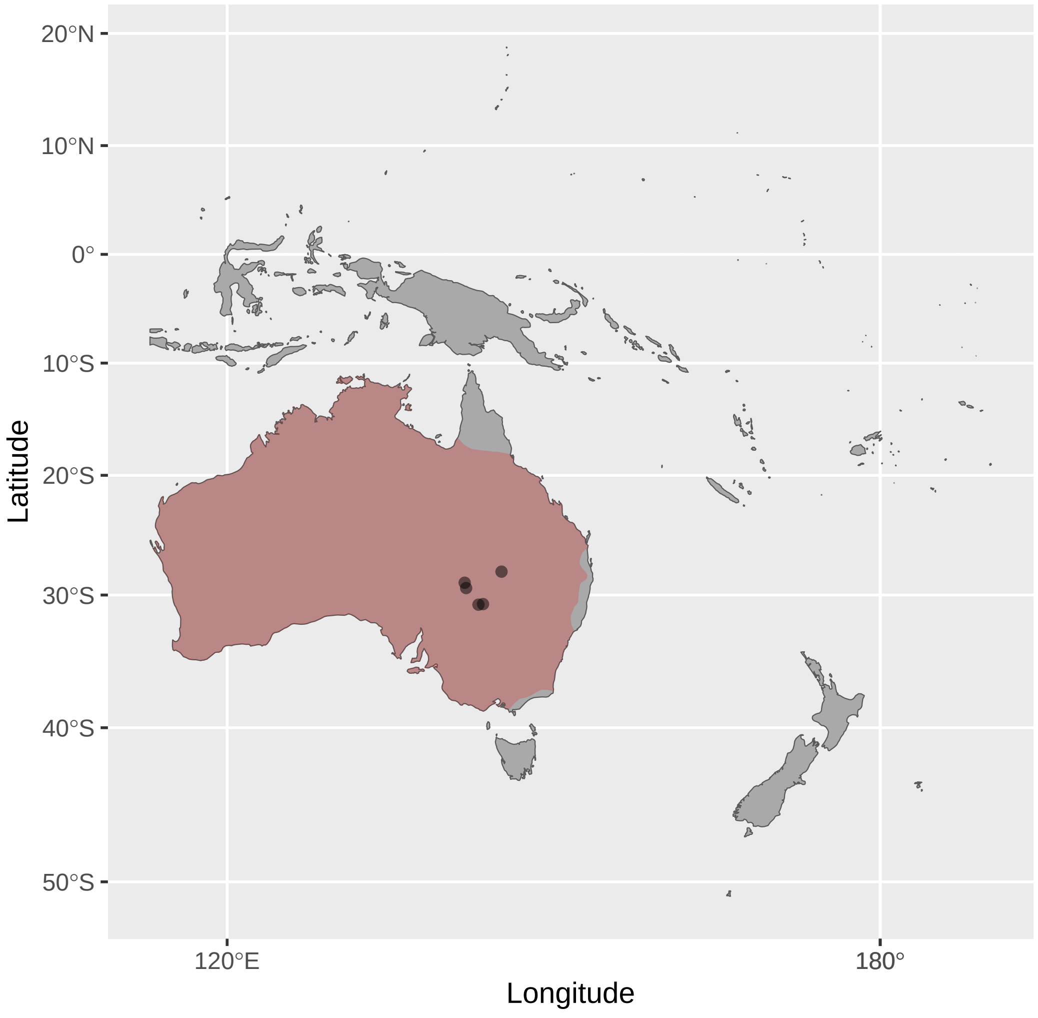


Glycifohia_undulata (n = 4)

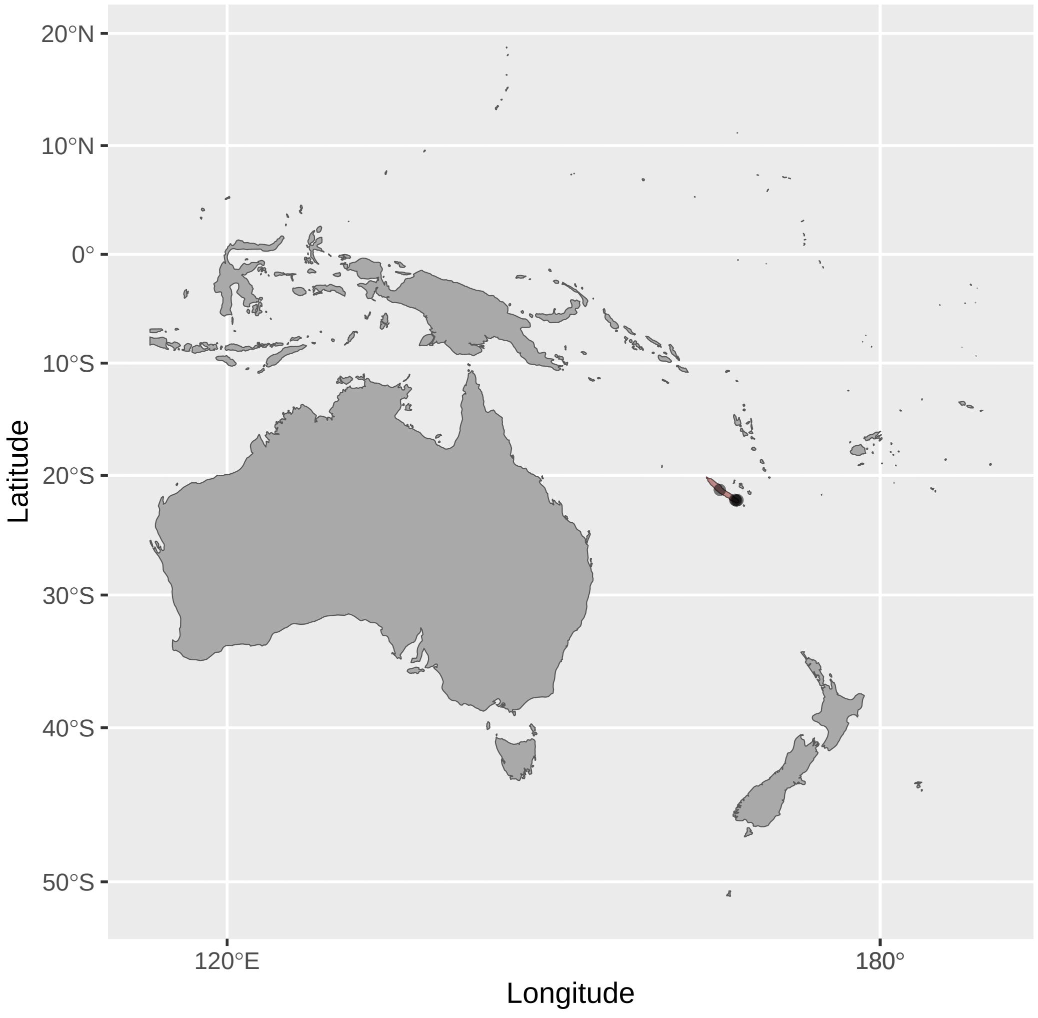


Foulehaio_taviunensis (n = 1)

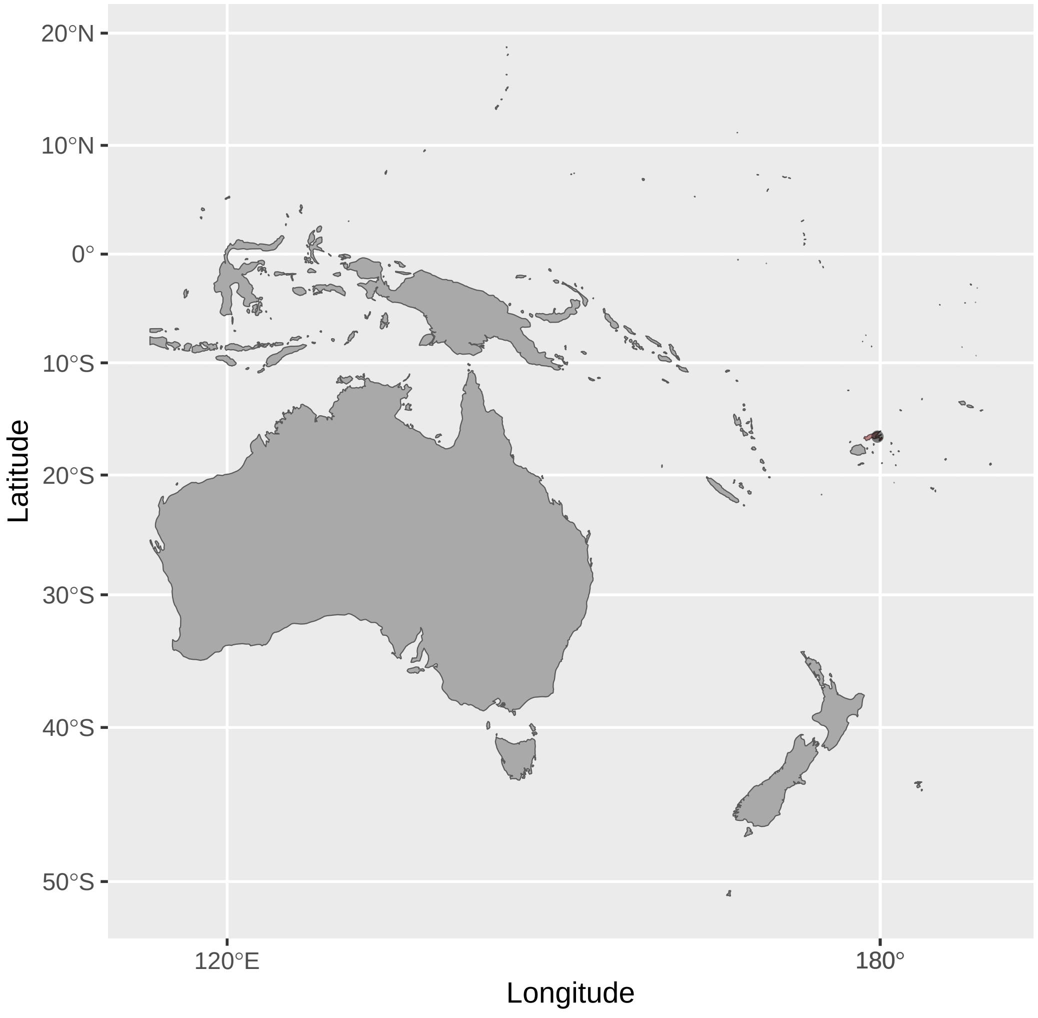


Gliciphila_melanops (n = 4)

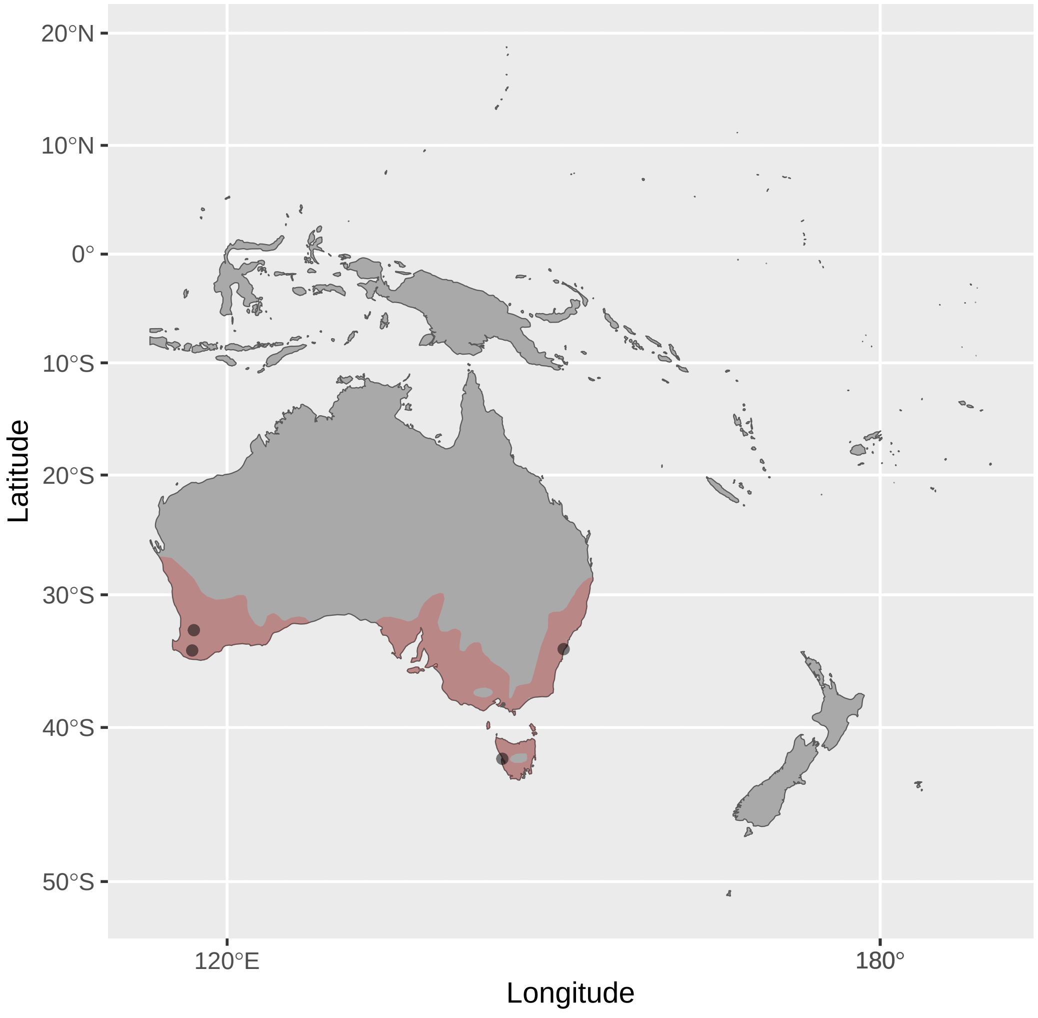


Grantiella_picta (n = 4)

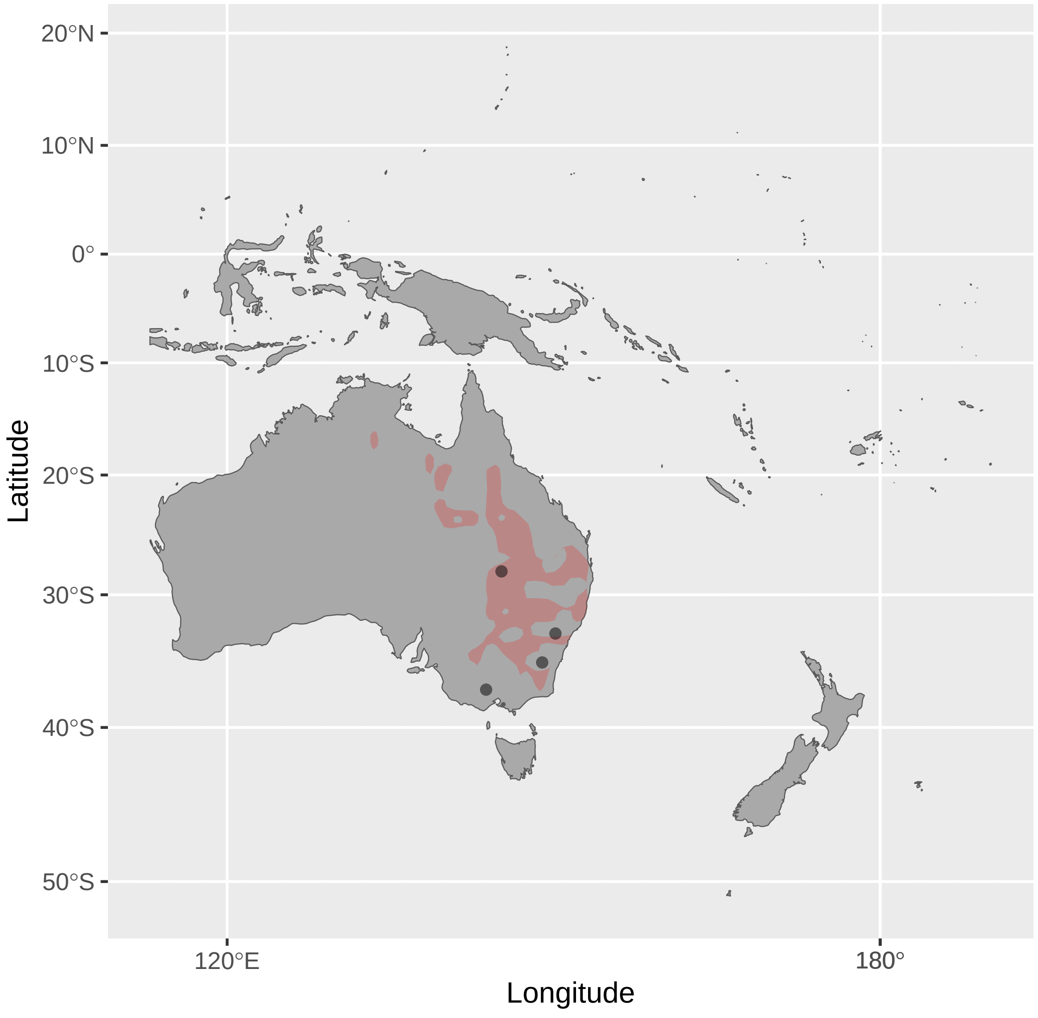


Gavicalis_fasciogularis (n = 2)

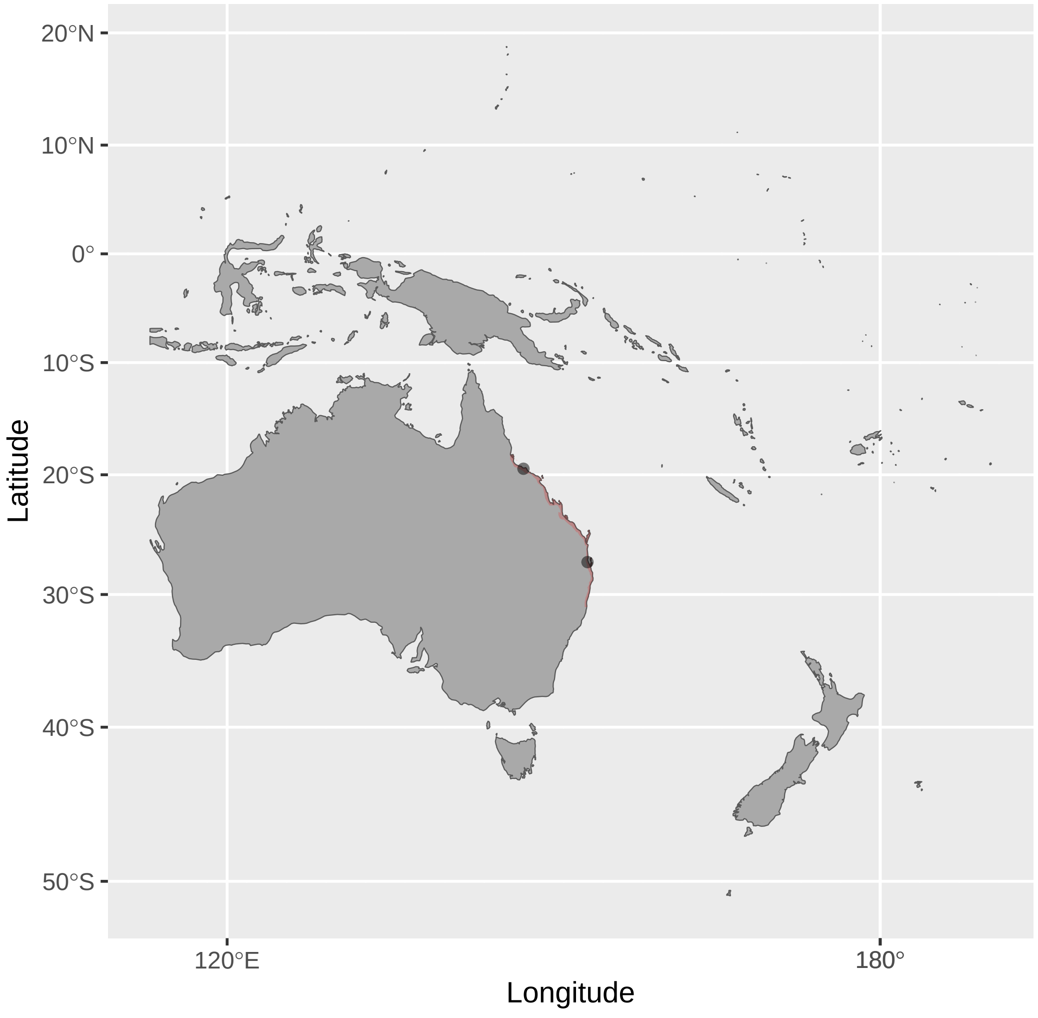


Glycichaera_fallax (n = 2)

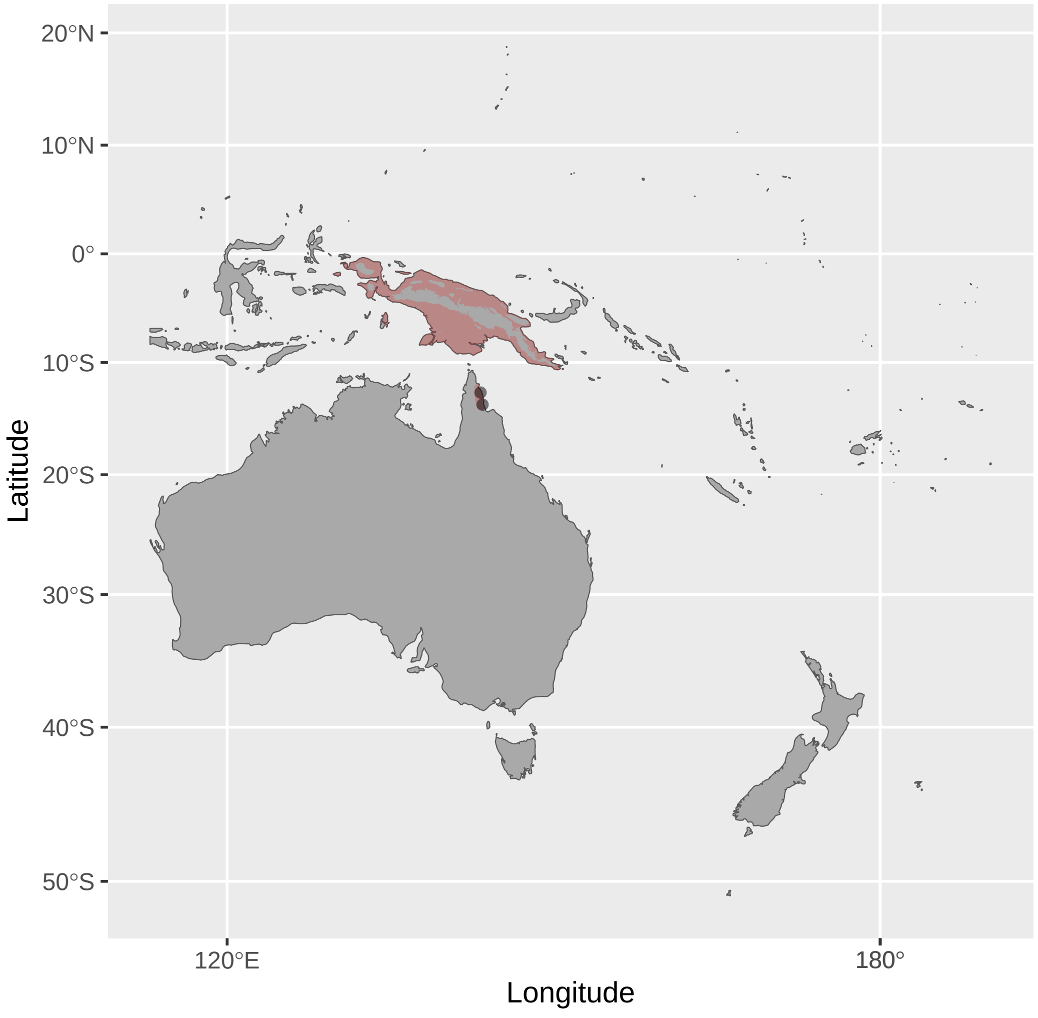


Guadalcanaria_inexpectata (n = 1)

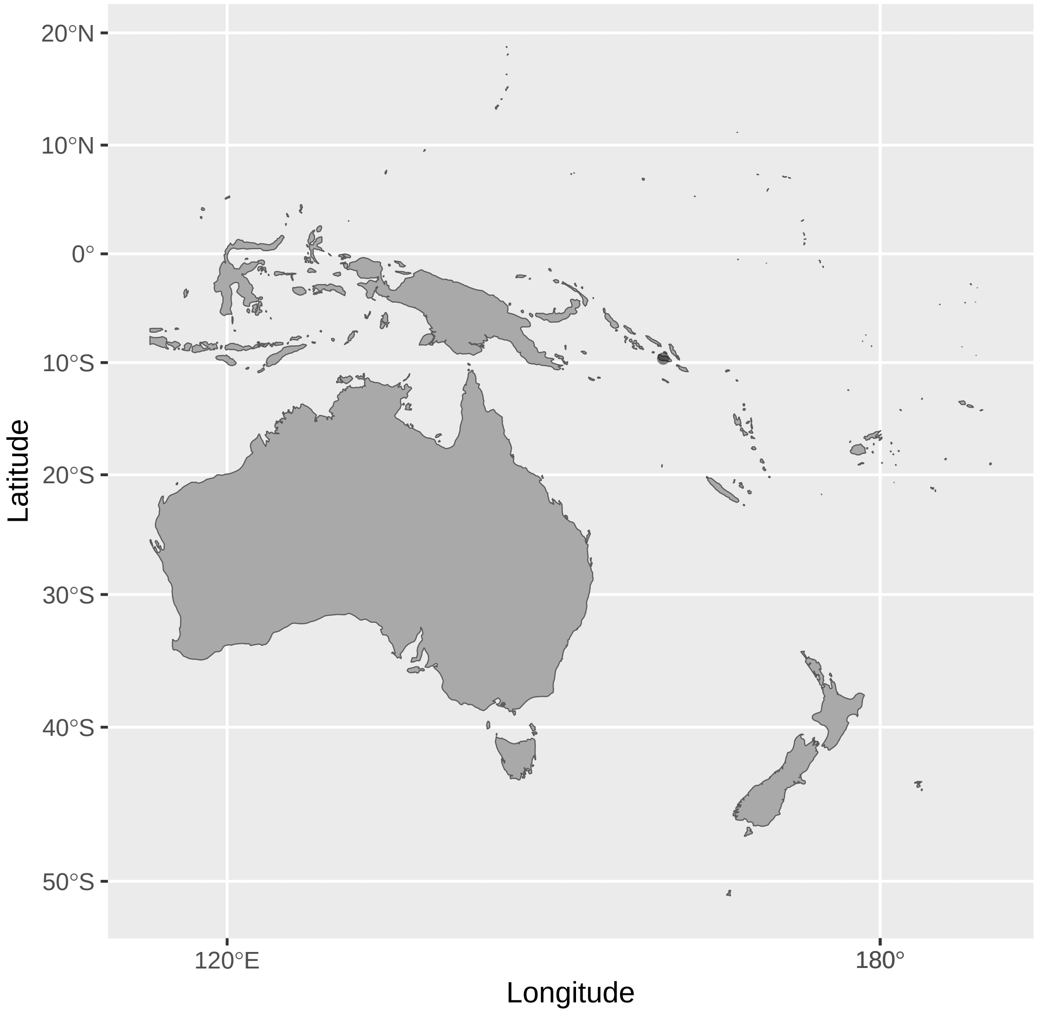

Gymnomyza_aubryana (n = 1)

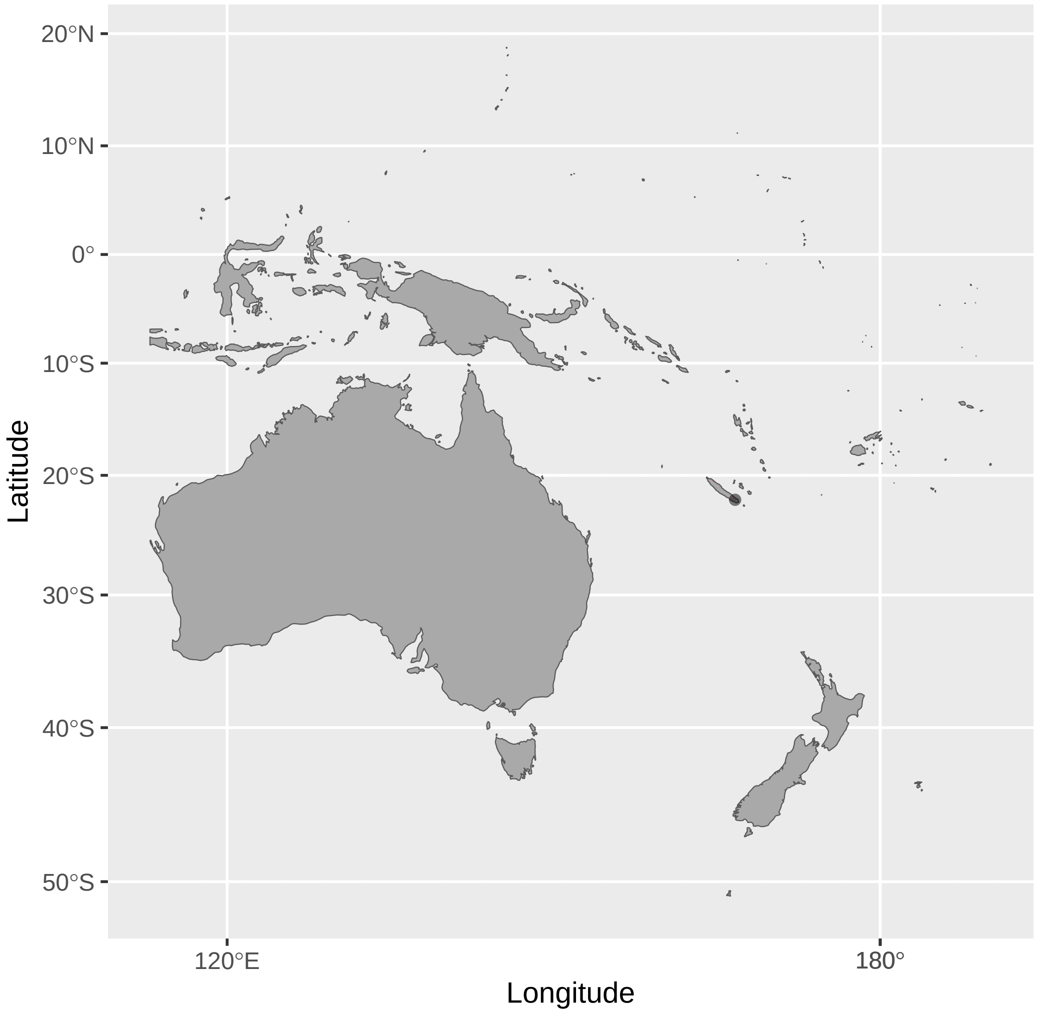


Gymnomyza_brunneirostris (n = 2)

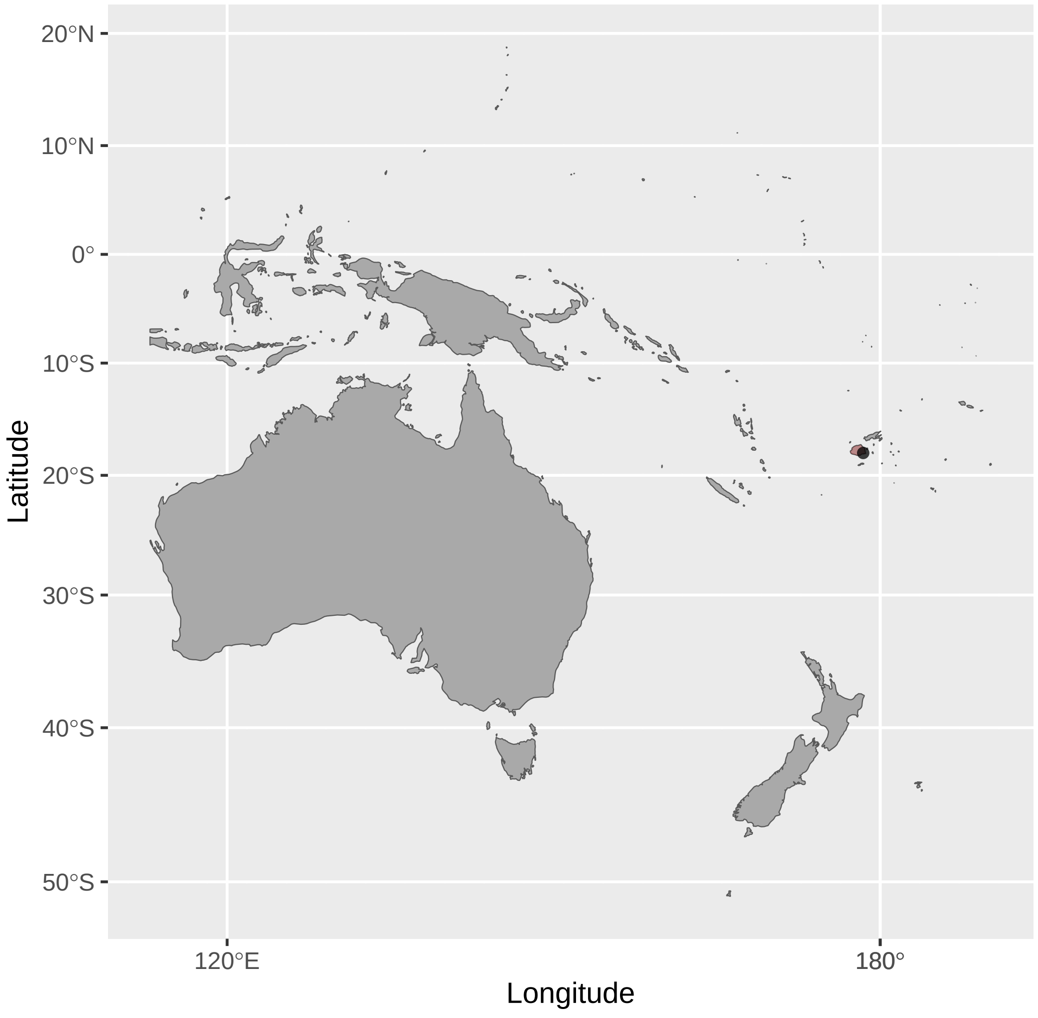


Gymnomyza_samoensis (n = 3)

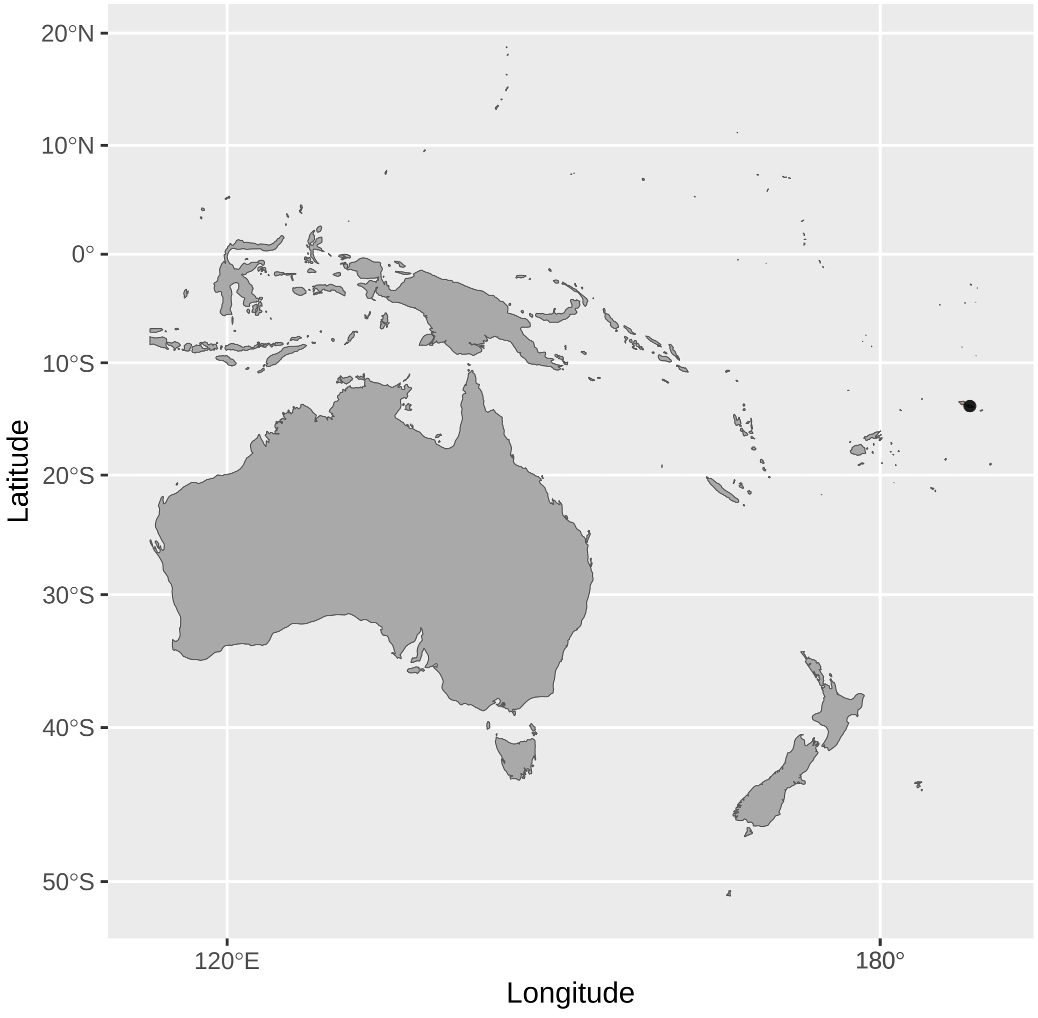


Gymnomyza_viridis (n = 1)

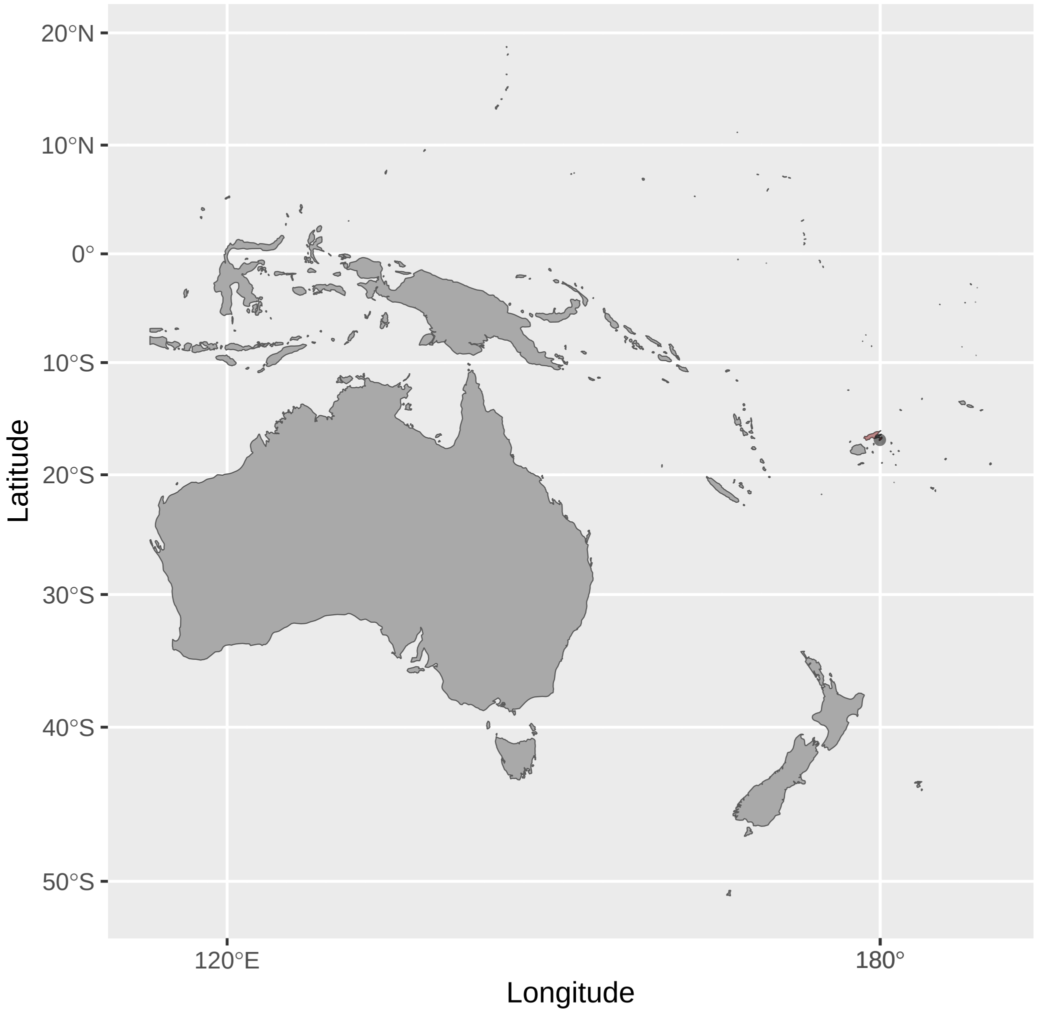


Lichenostomus_cratitius (n = 2)

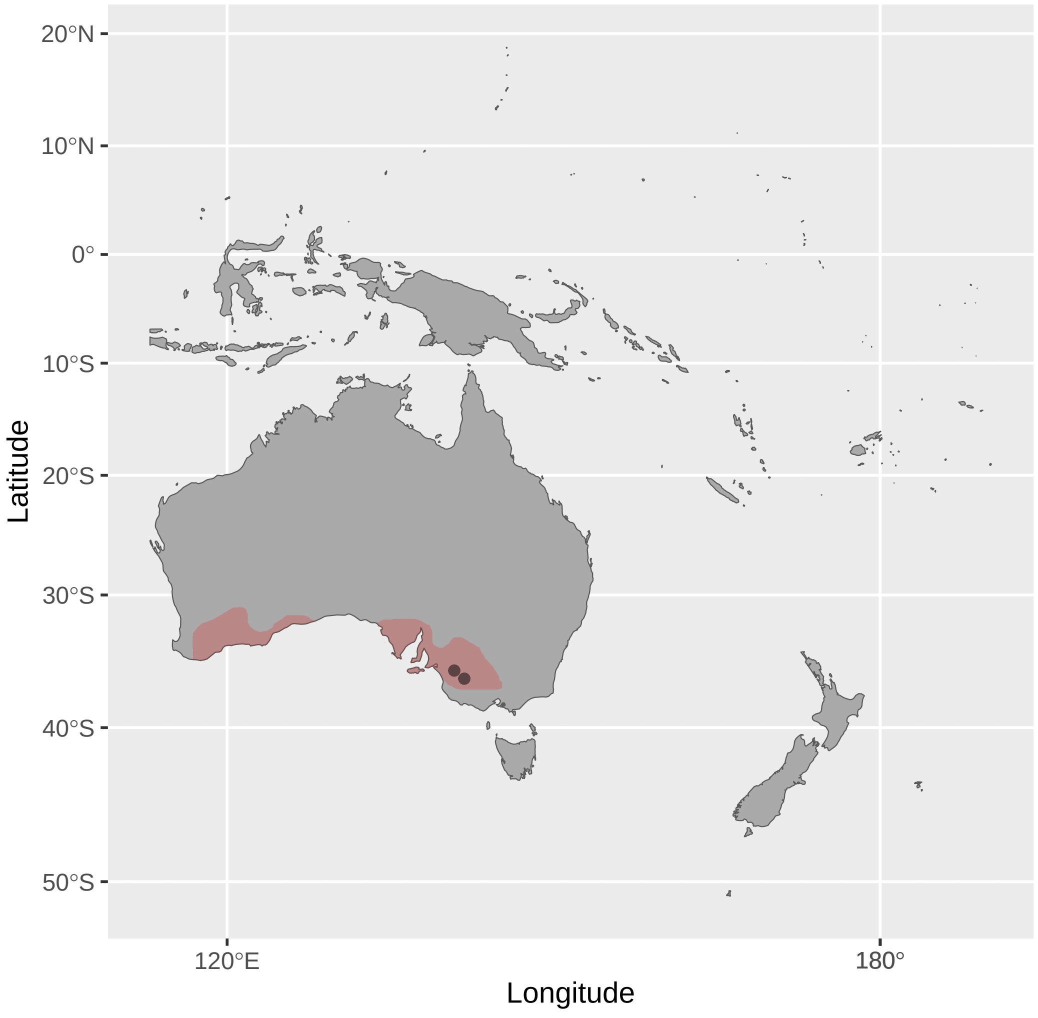


Lichenostomus_melanops (n = 1)

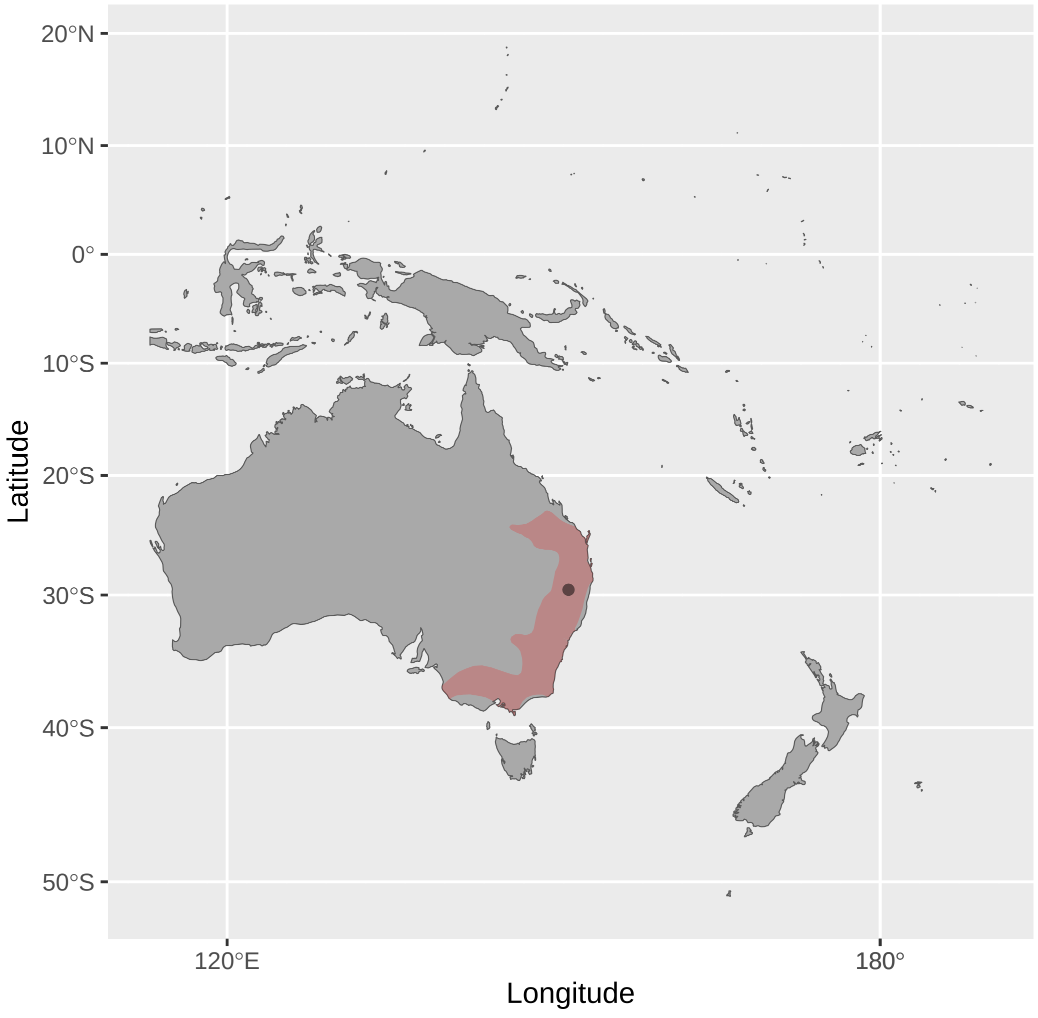


Lichmera_alboauricularis (n = 1)

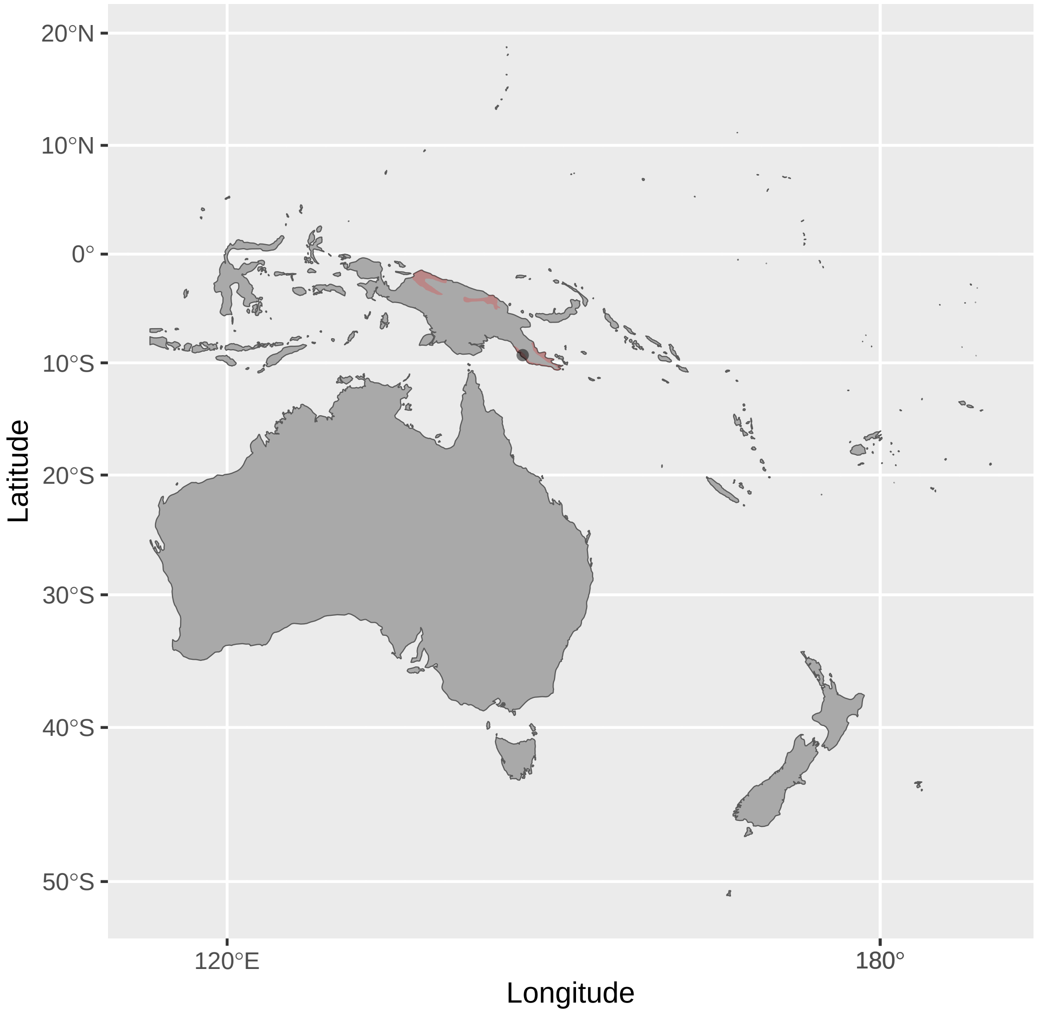


Lichmera_argentauris (n = 1)

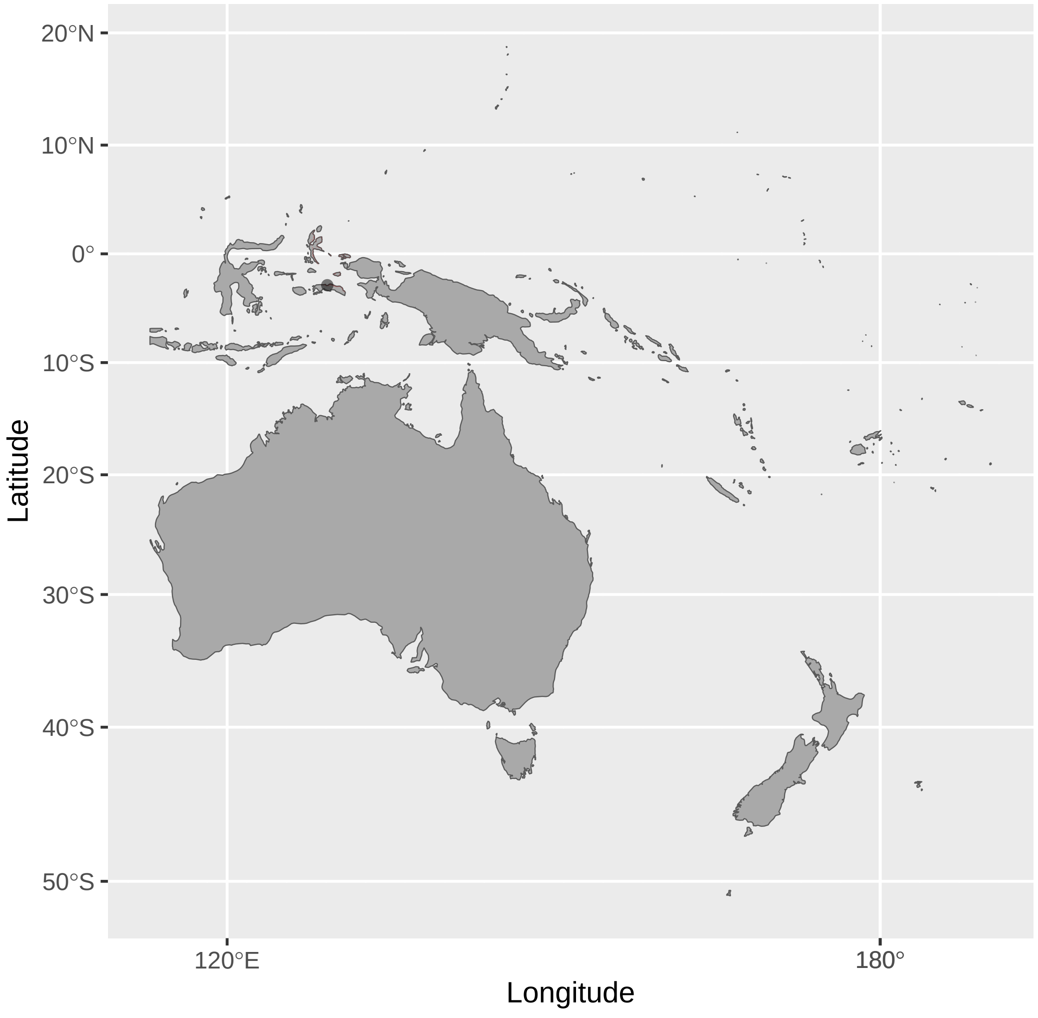


Lichmera_flavicans (n = 1)

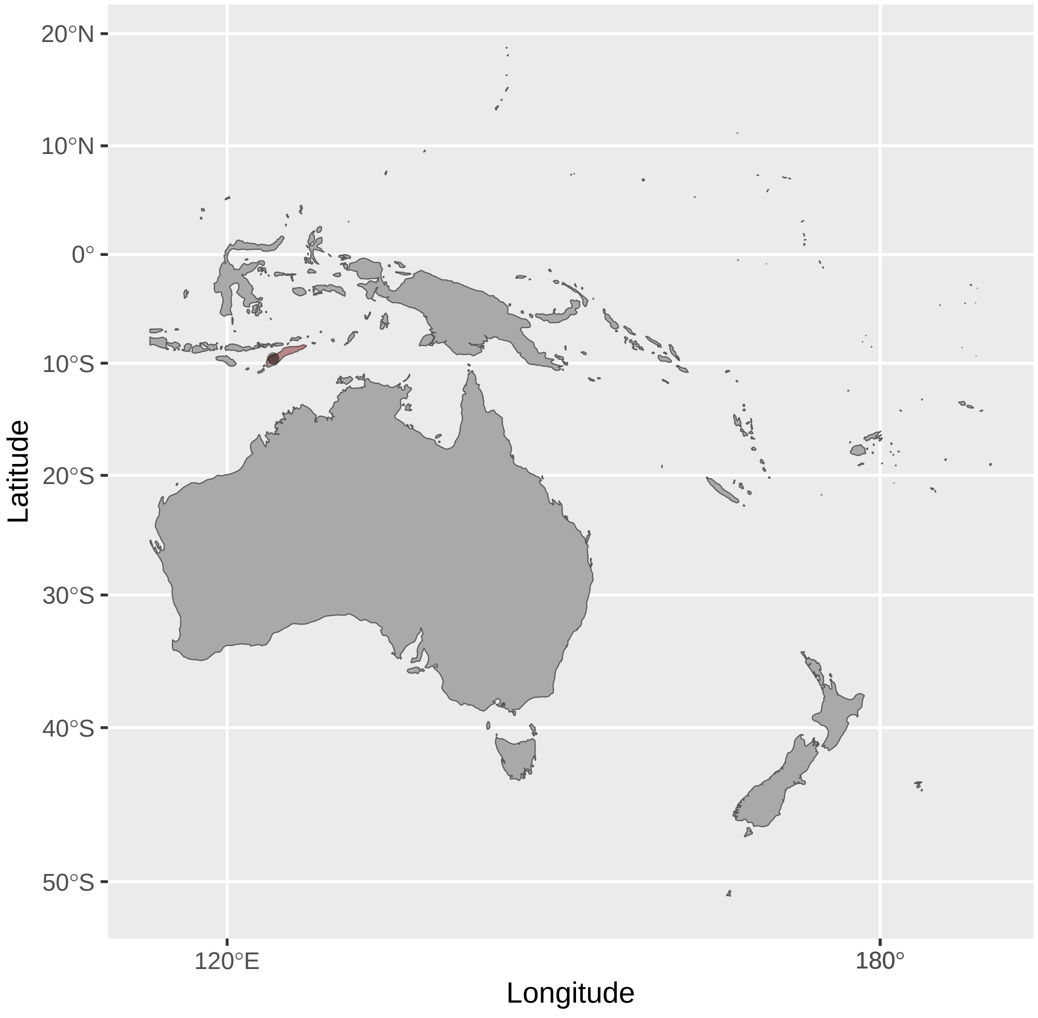


Lichmera_incana (n = 3)

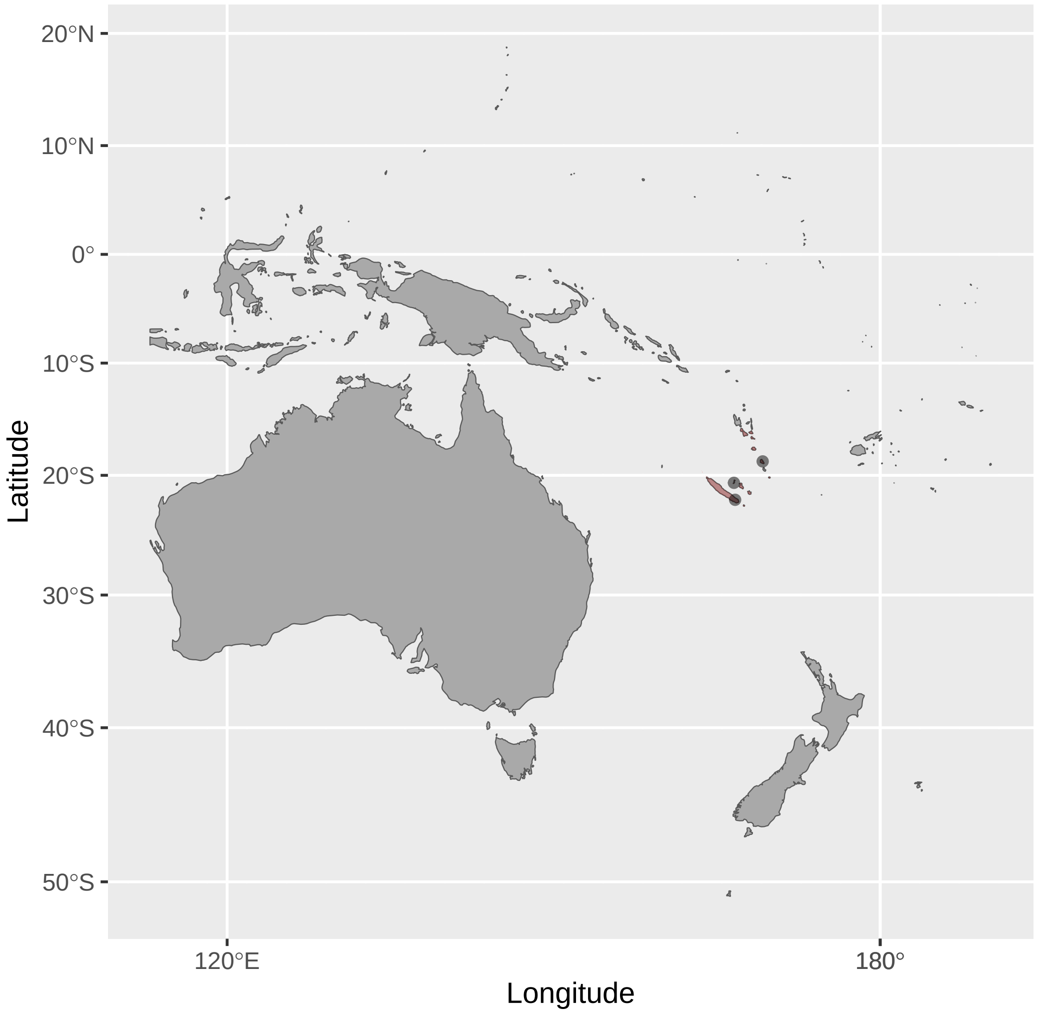


Lichmera_indistincta (n = 20)

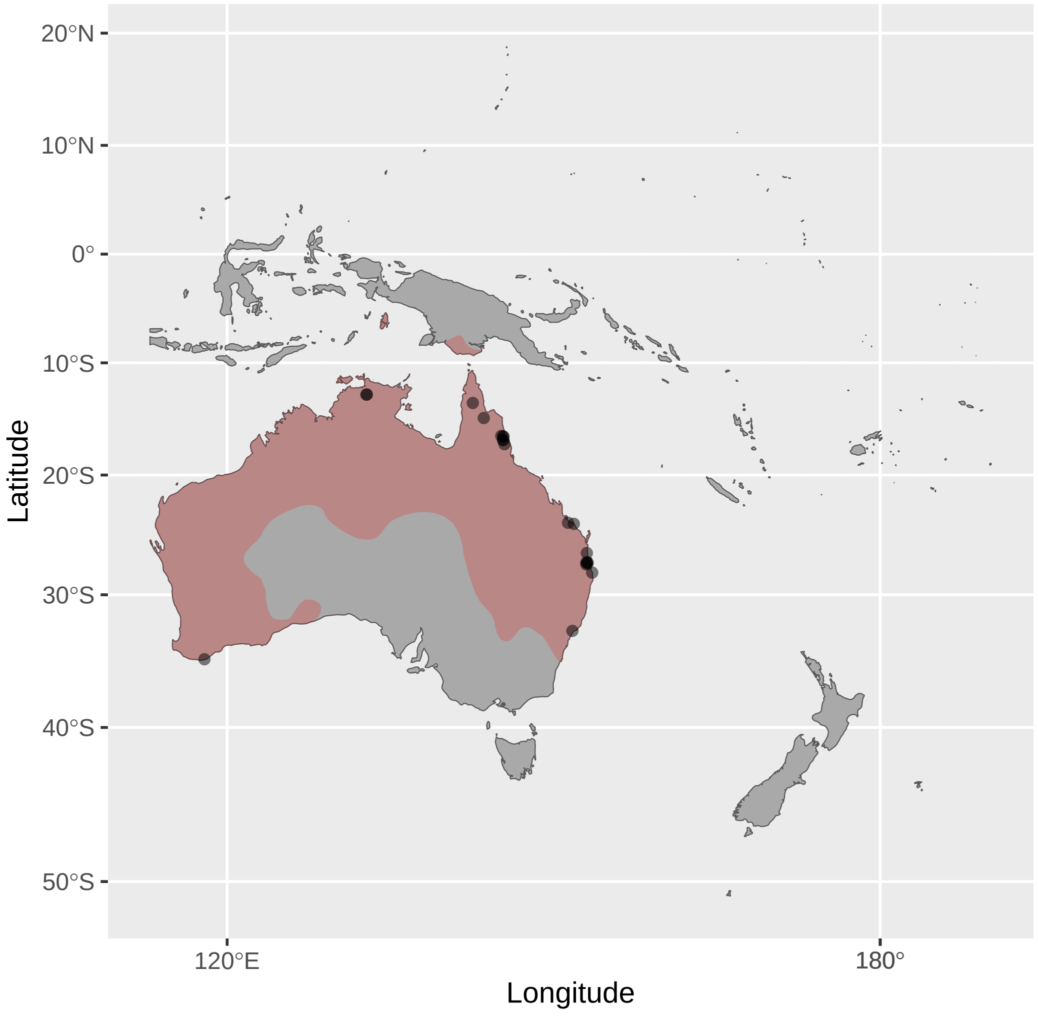


Lichmera_limbata (n = 3)

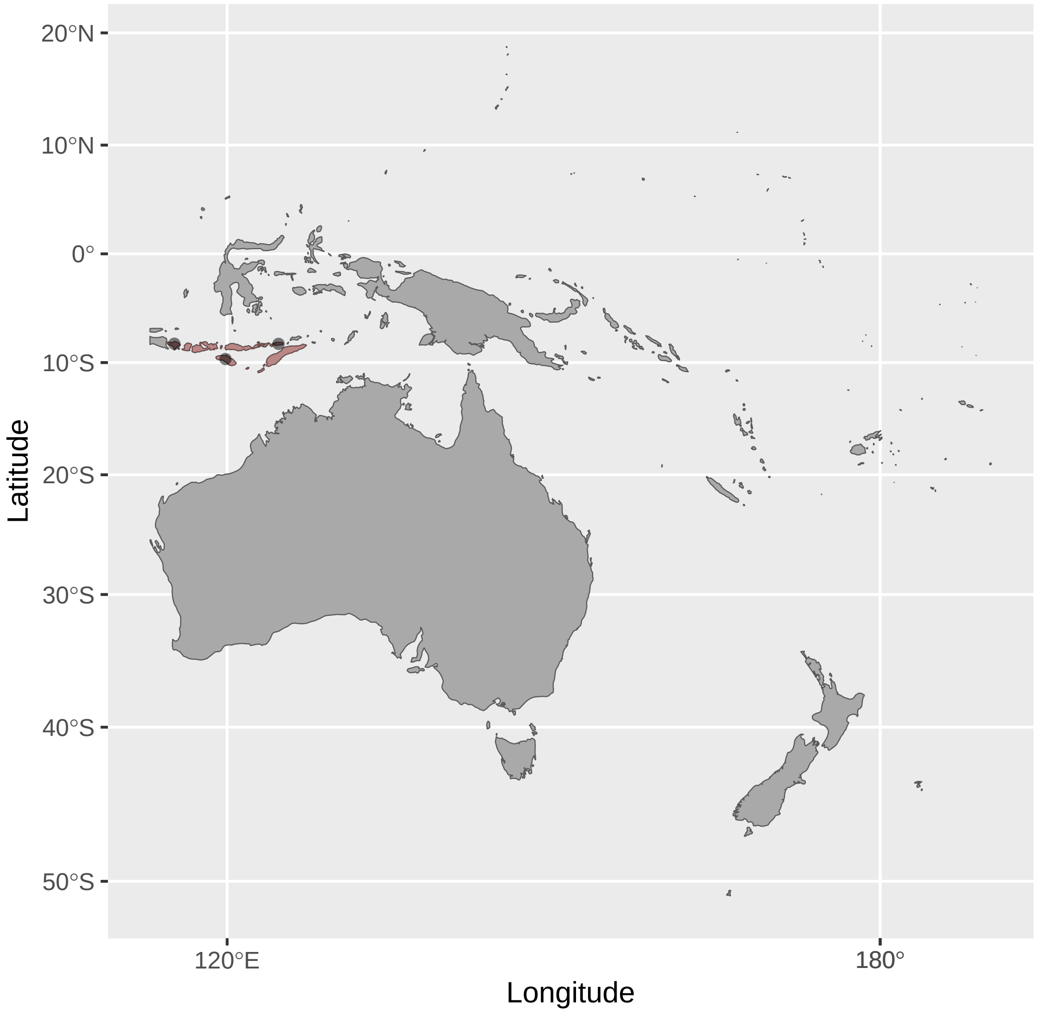

Lichmera_lombokia (n = 1)

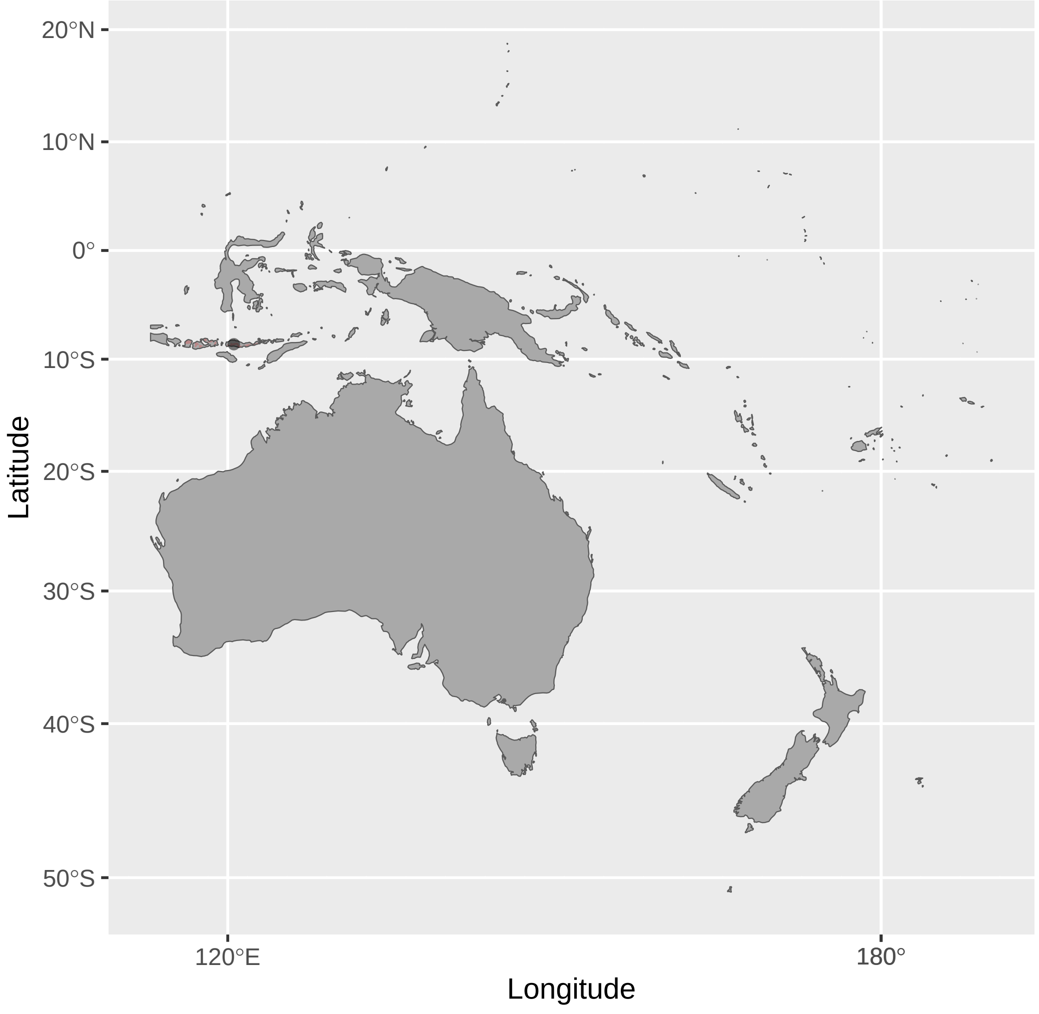


Manorina_flavigula (n = 2)

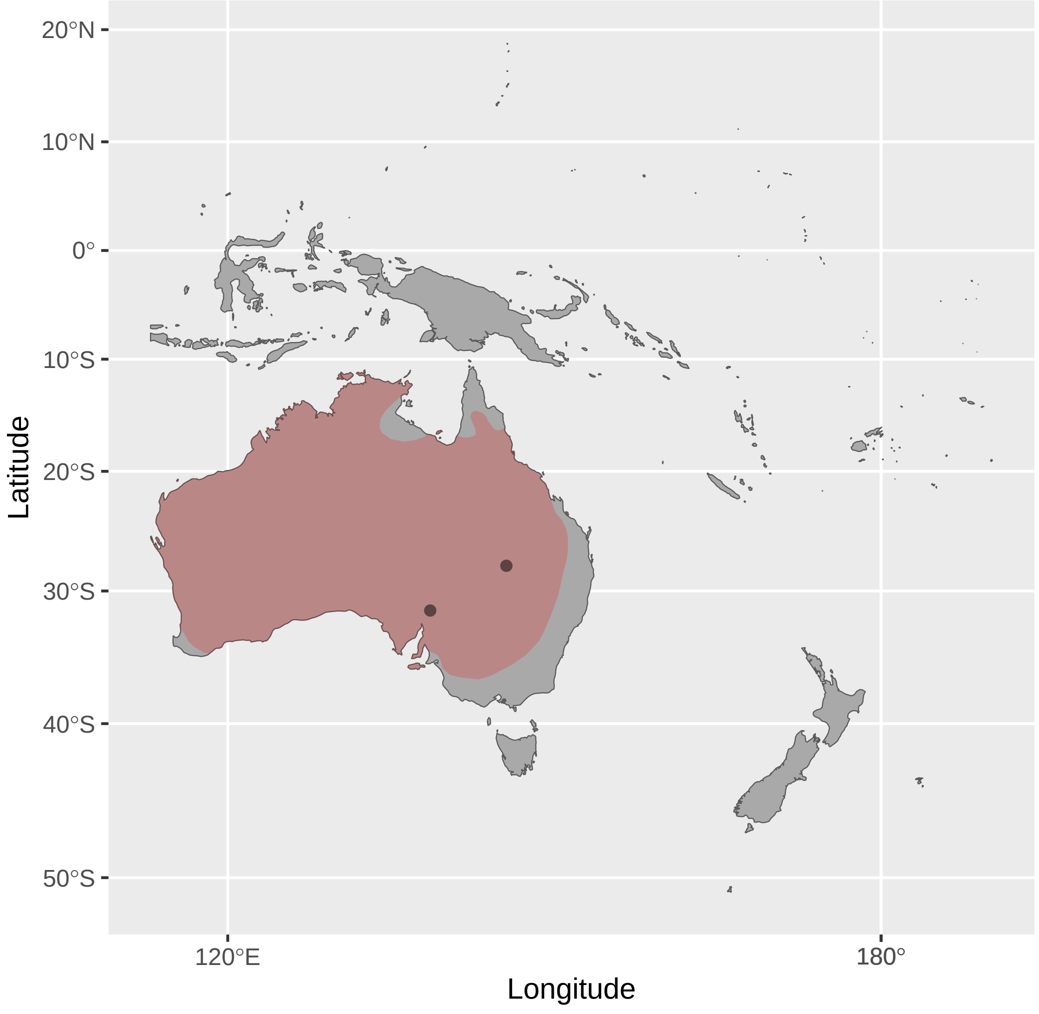


Meliarchus_sclateri (n = 1)

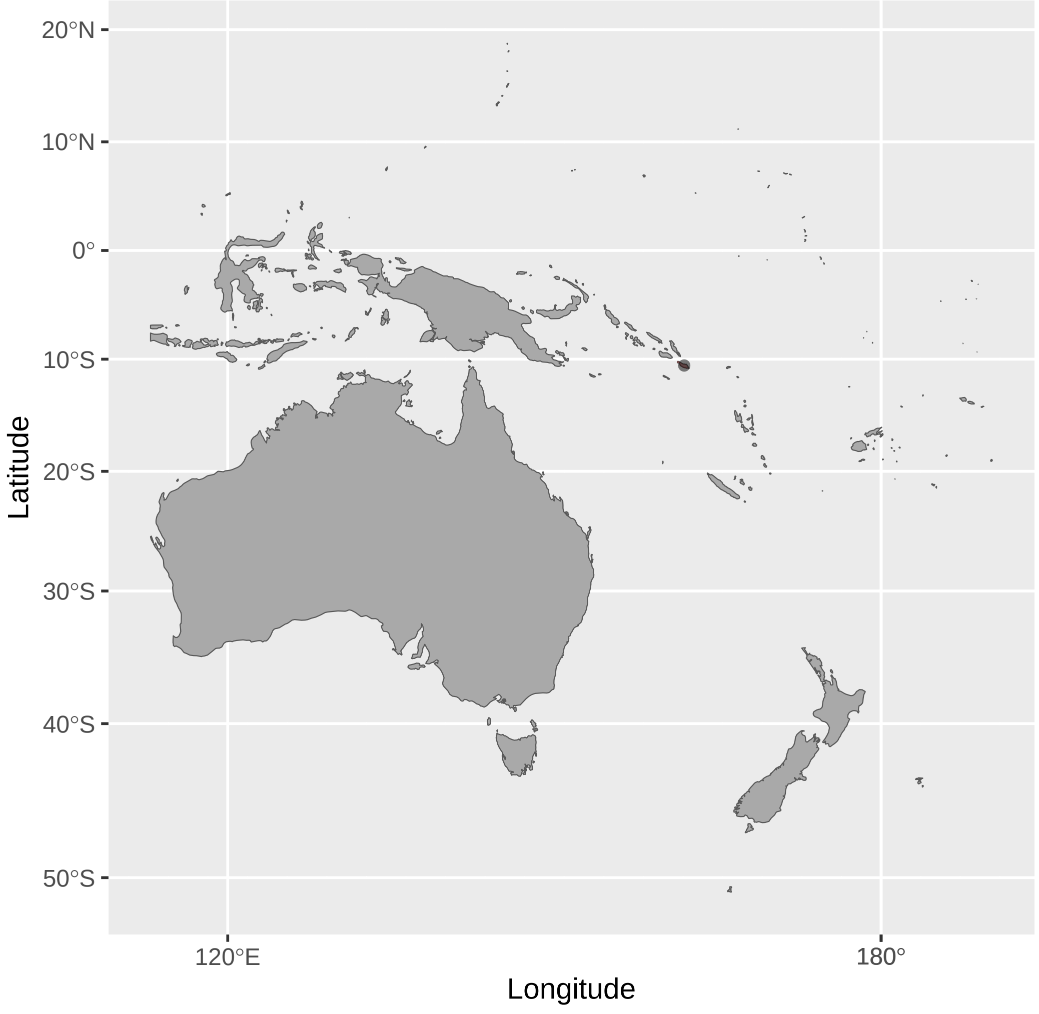


Lichmera_monticola (n = 1)

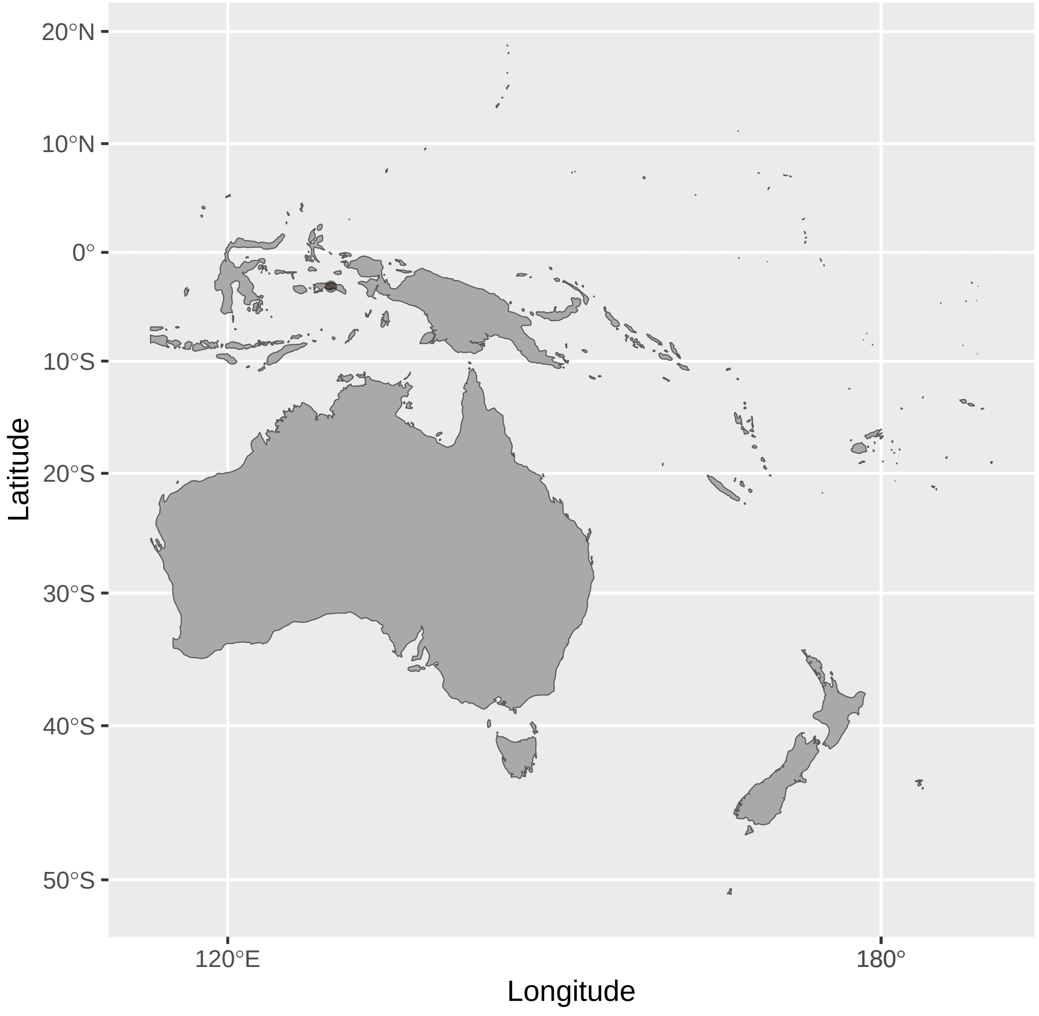


Manorina_melanocephala (n = 3)

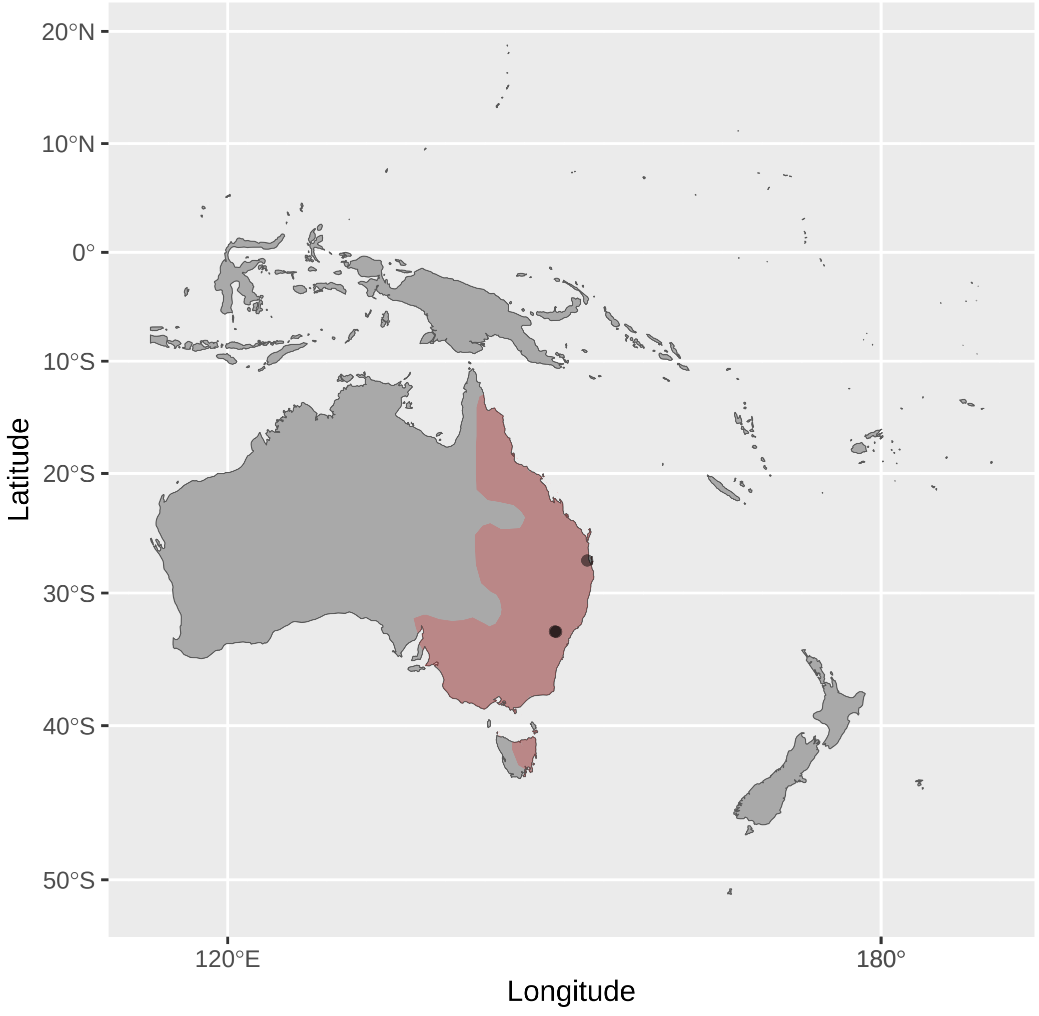


Melidectes_belfordi (n = 4)

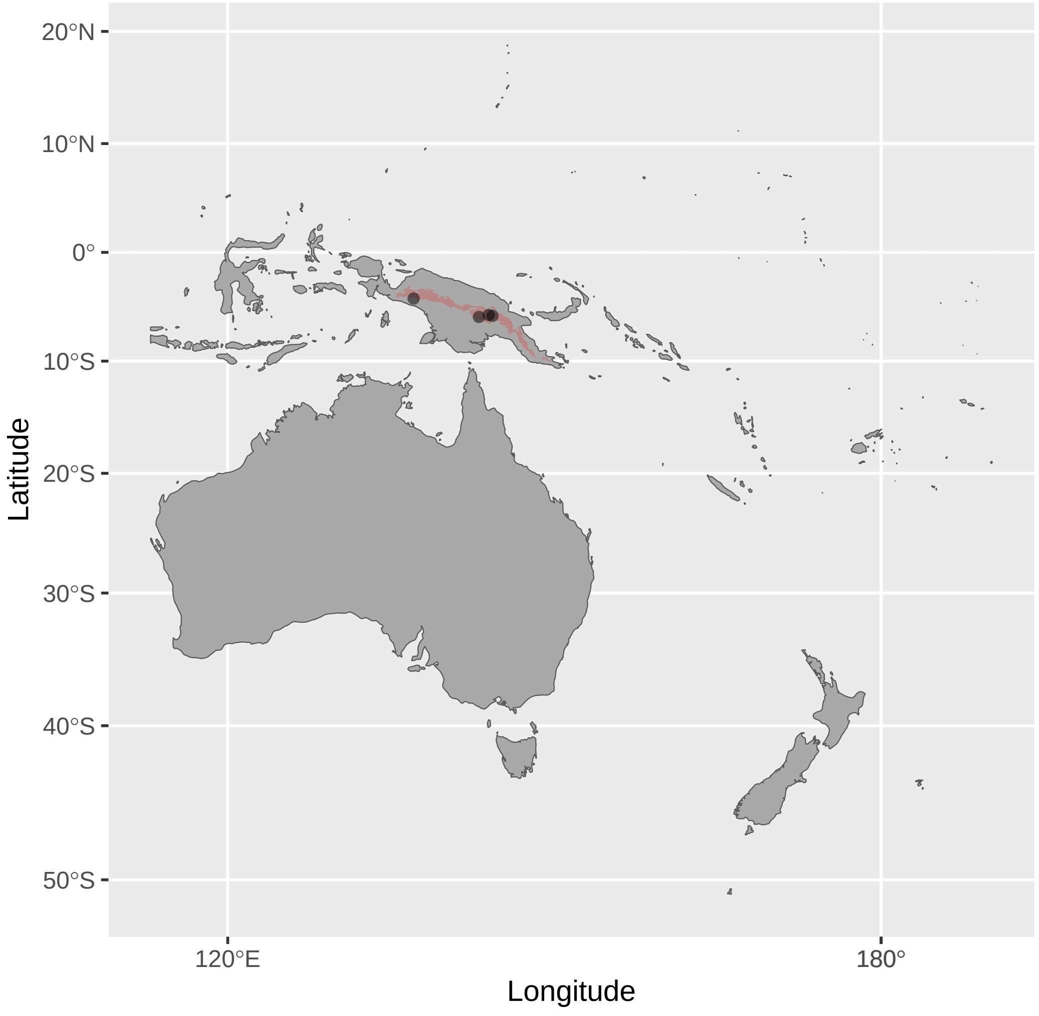


Lichmera_notabilis (n = 1)

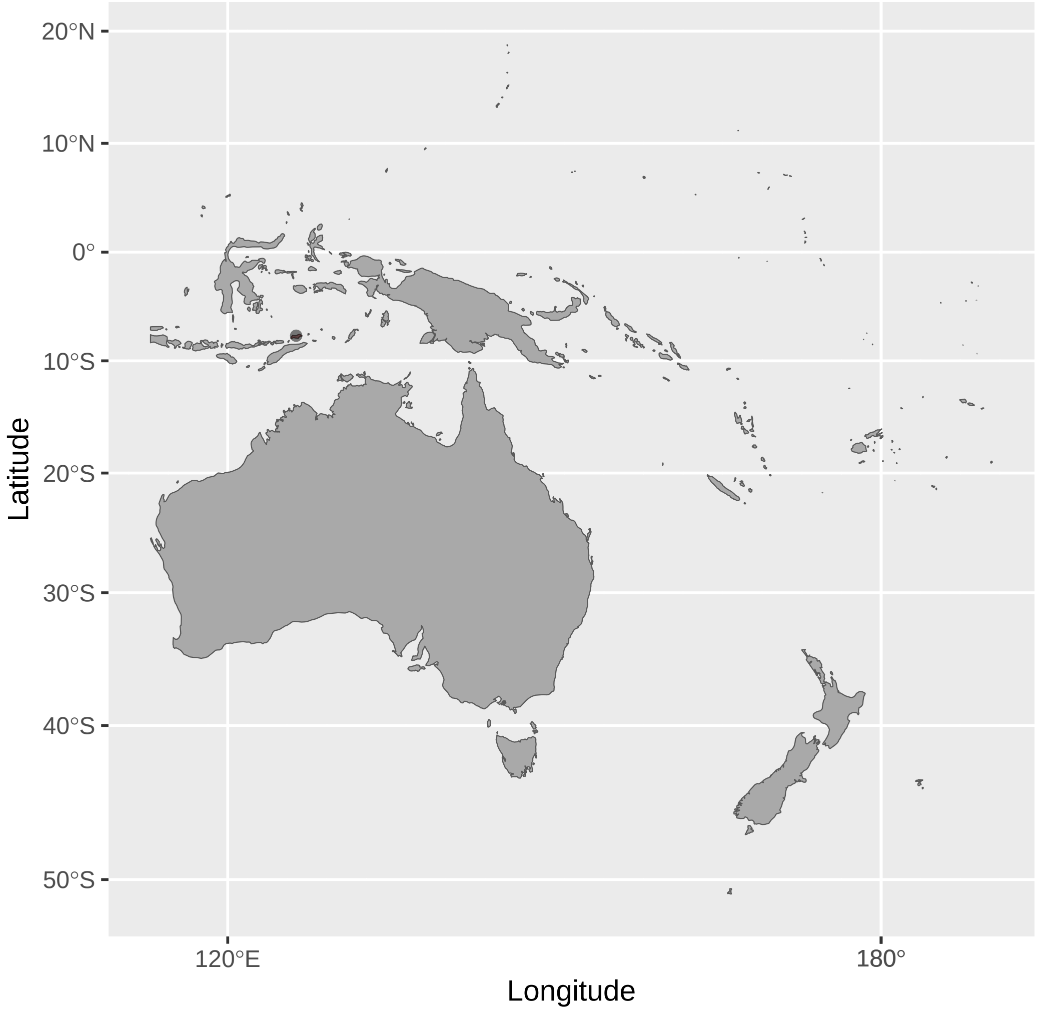


Manorina_melanophrys (n = 3)

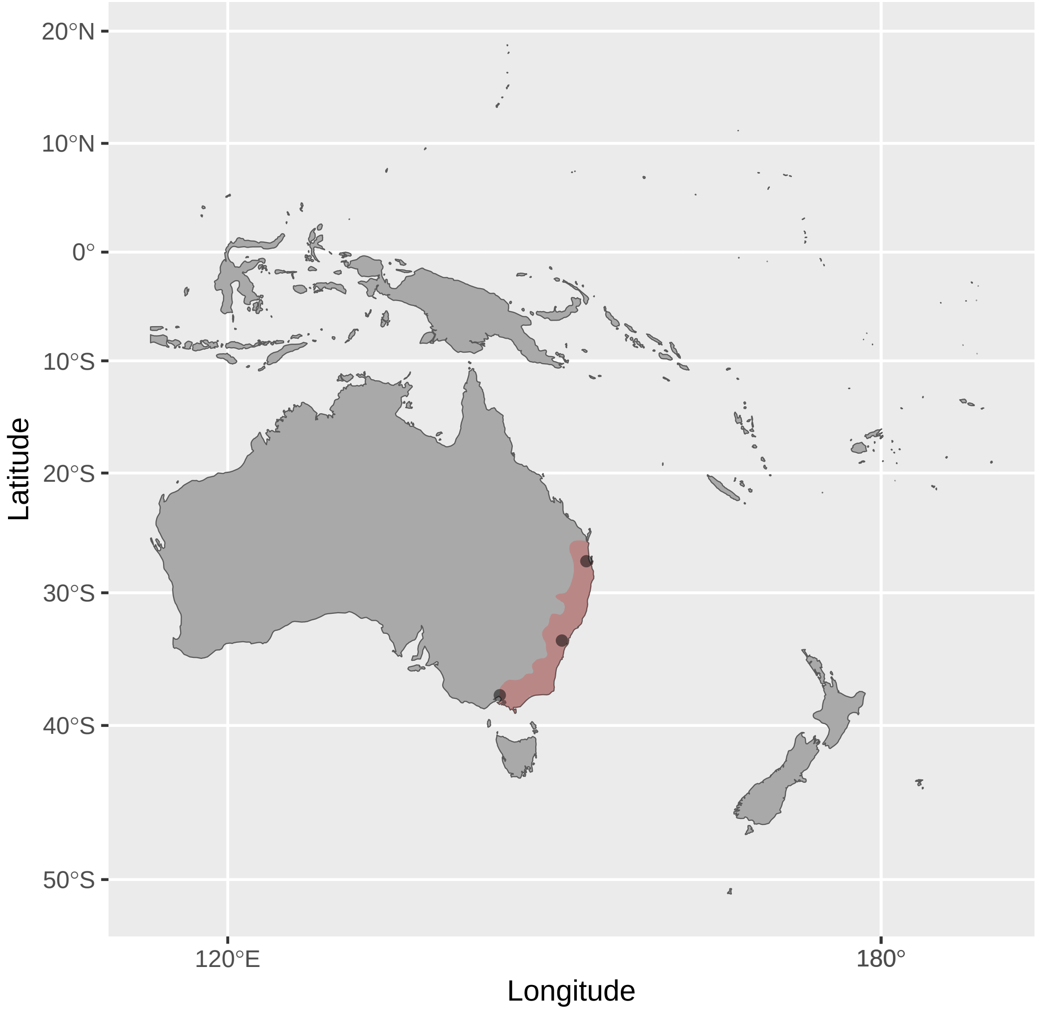


Melidectes_foersteri (n = 1)

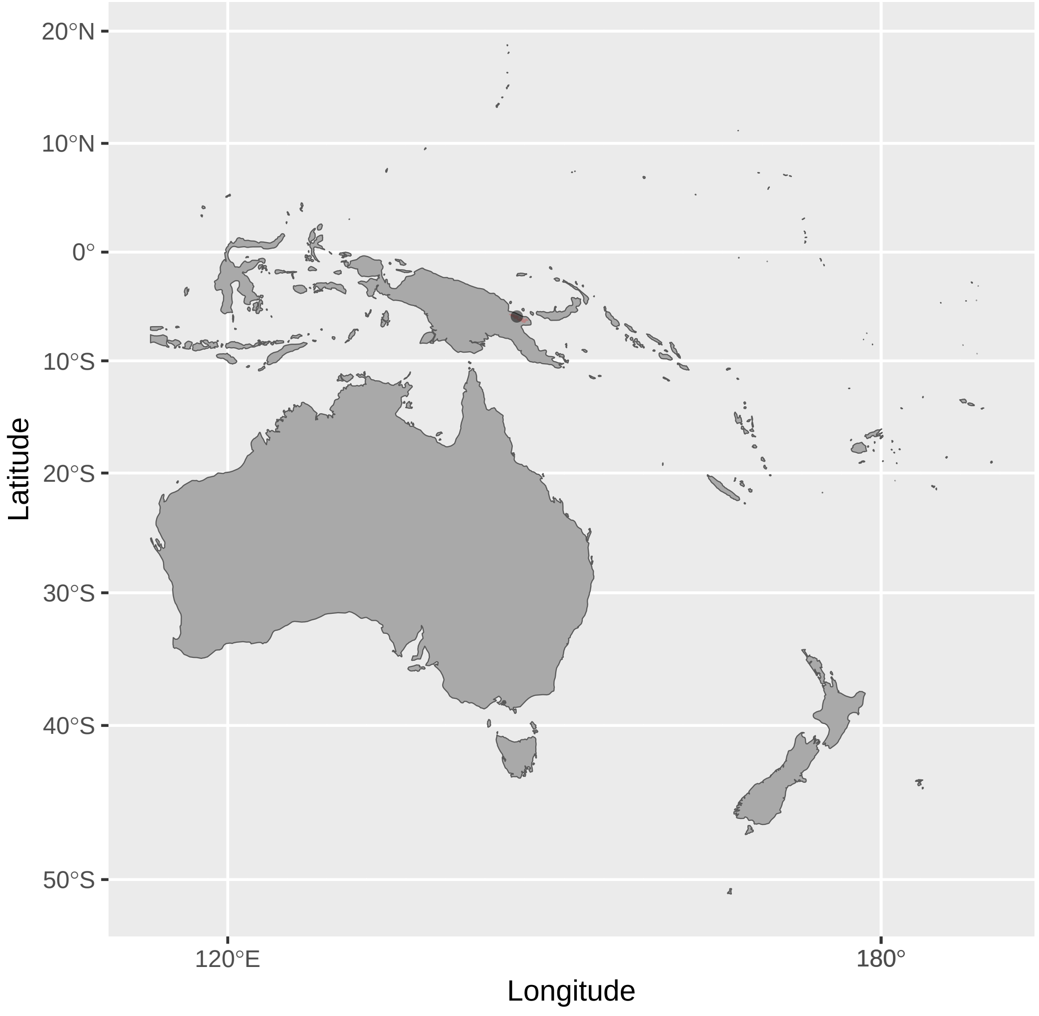


Lichmera_squamata (n = 2)

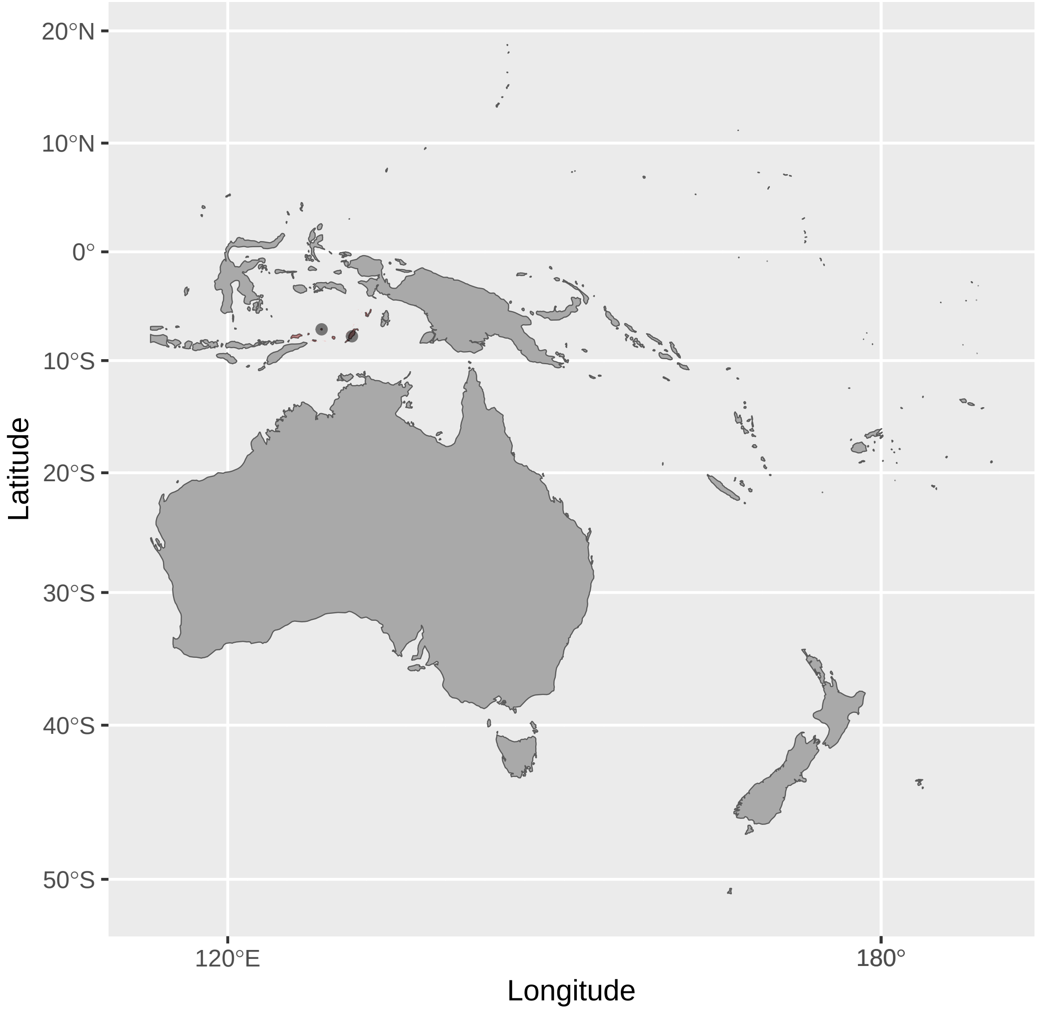


Manorina_melanotis (n = 1)

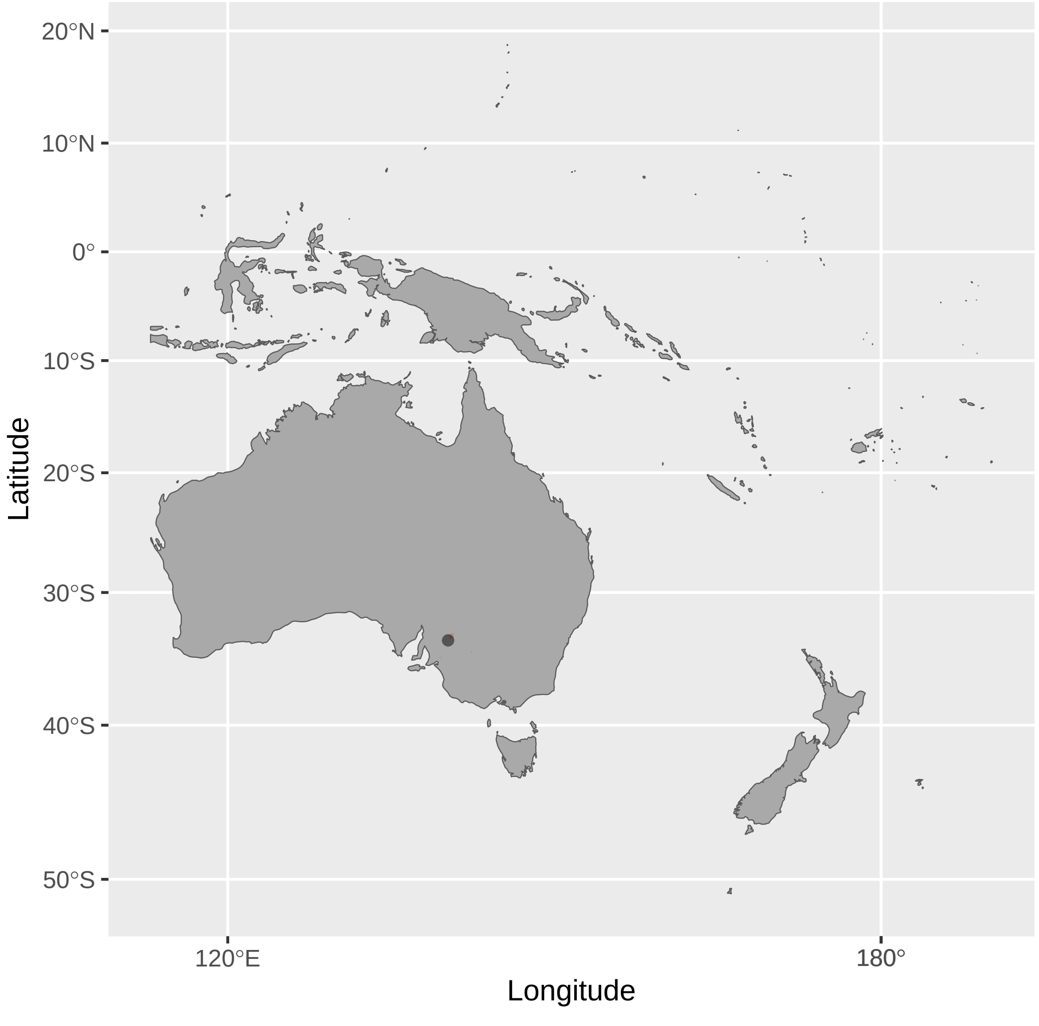


Melidectes_leucostephes (n = 1)

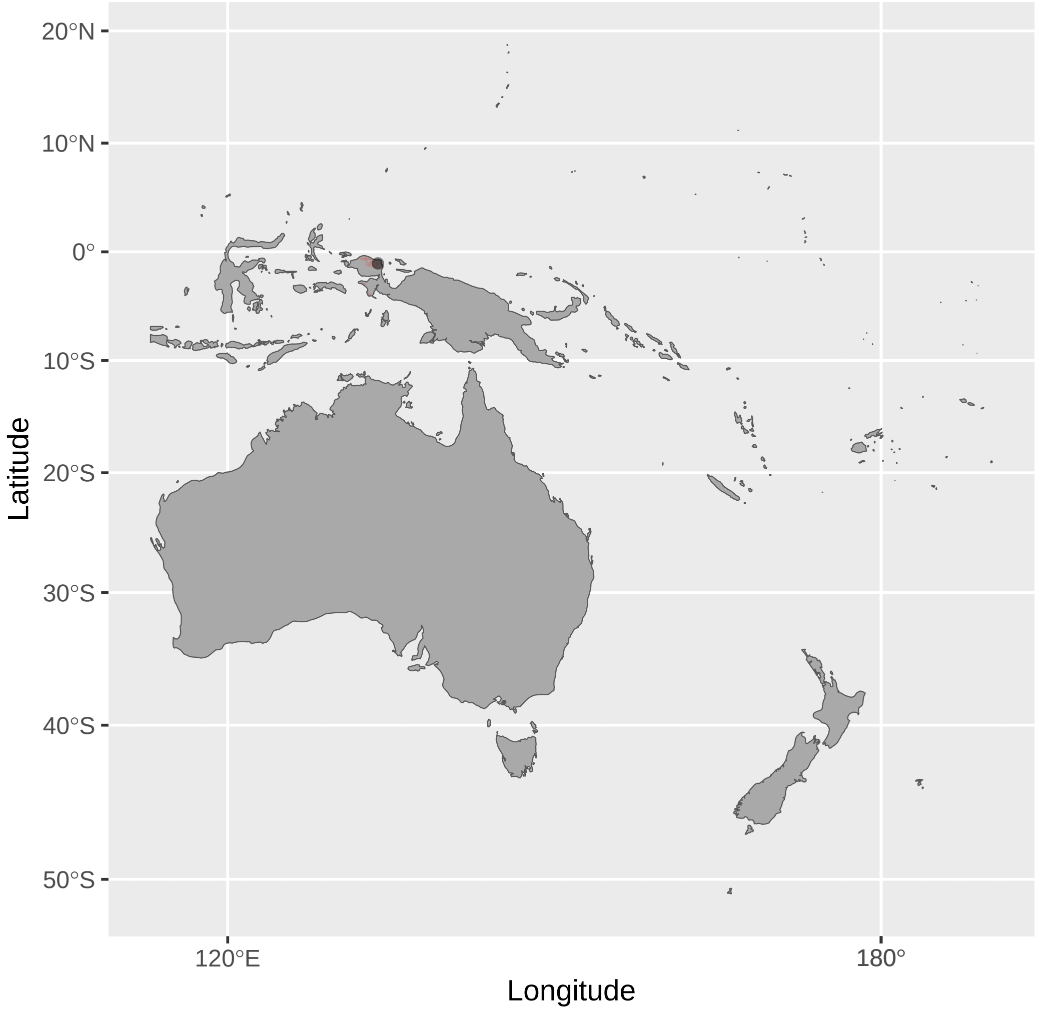

Melidectes_ochromelas (n = 2)

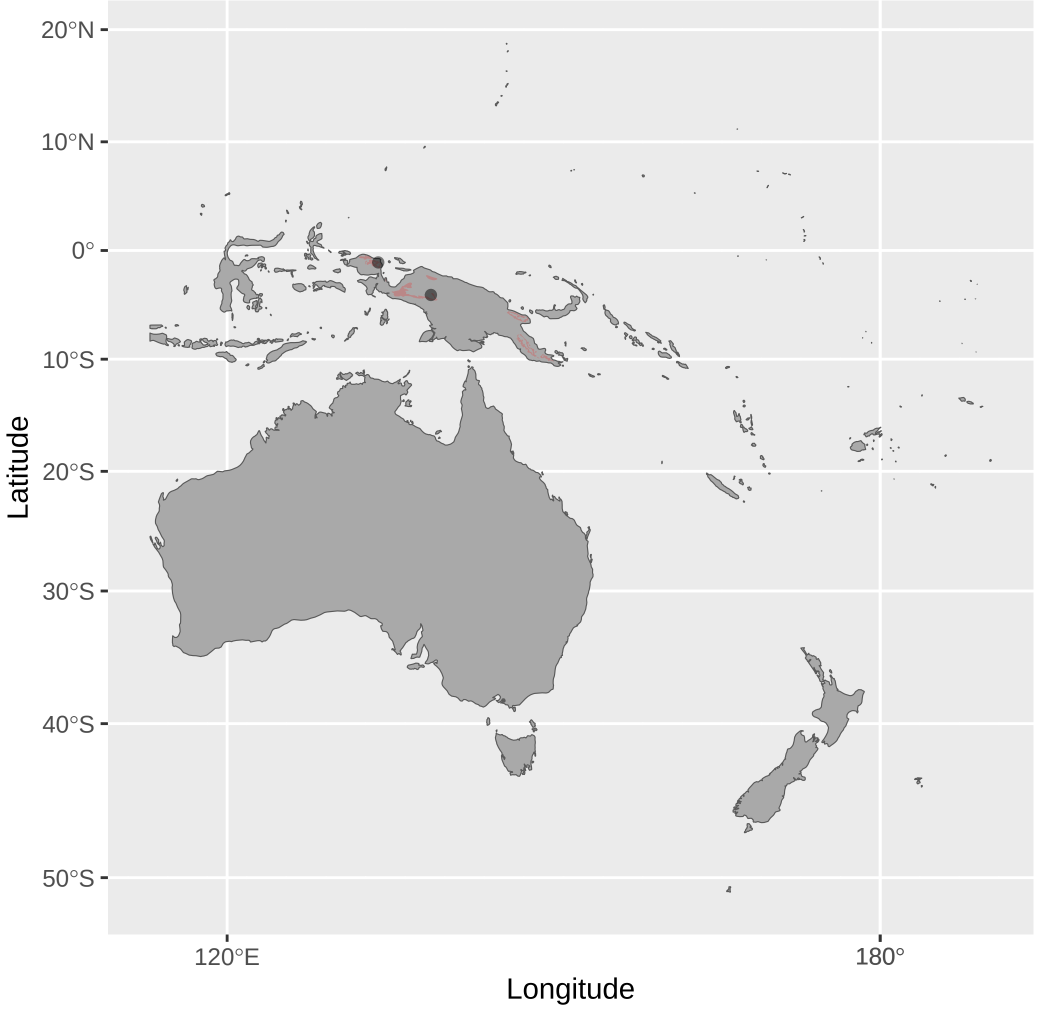


Melionyx_fuscus (n = 2)

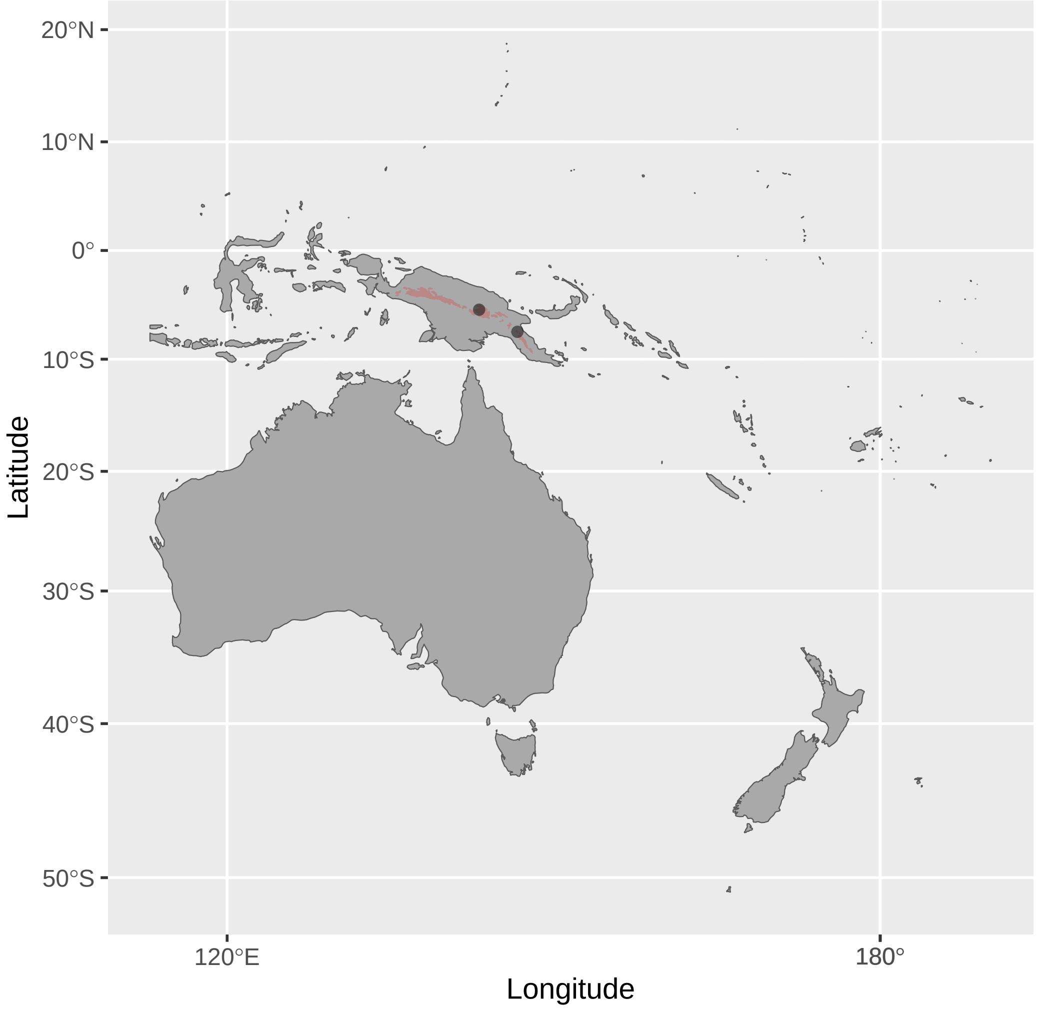


Meliphaga_notata (n = 13)

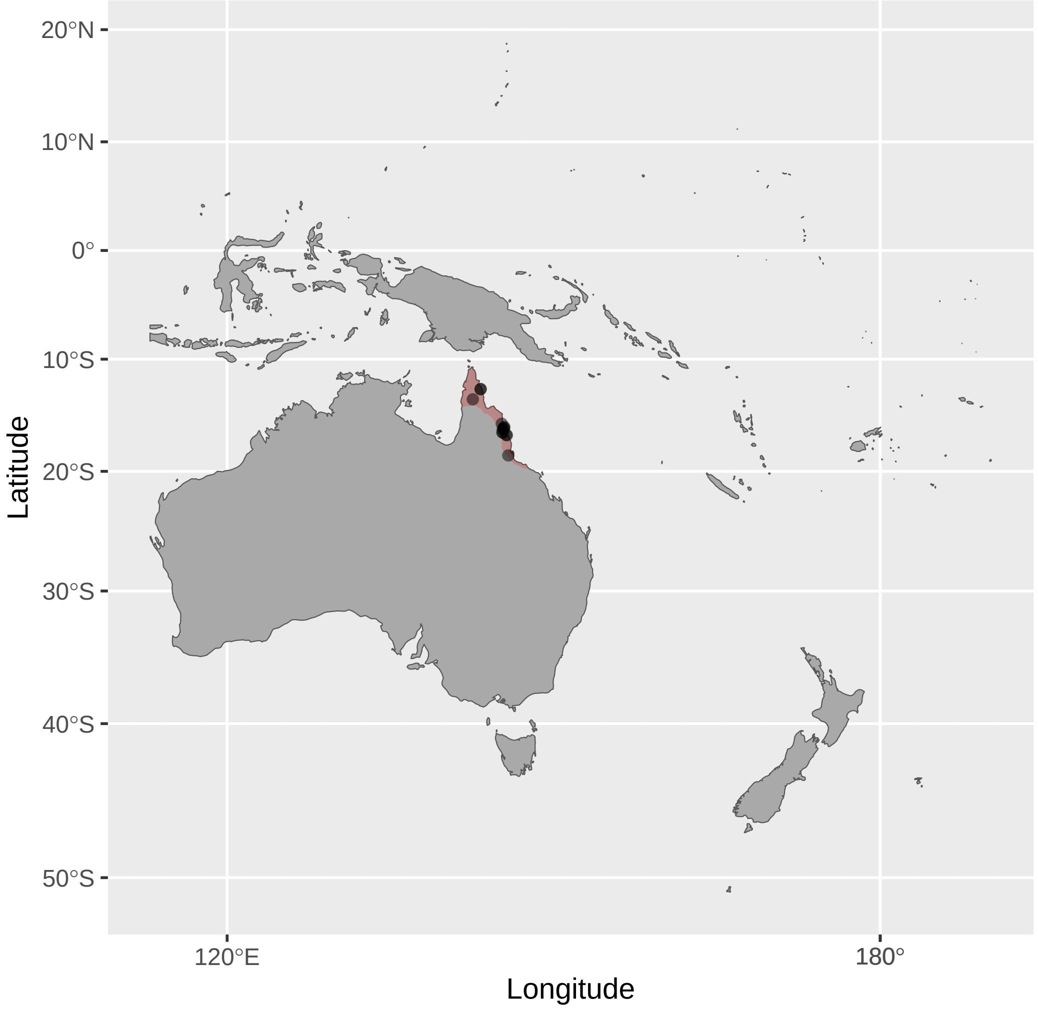


Melidectes_rufocrissalis (n = 2)

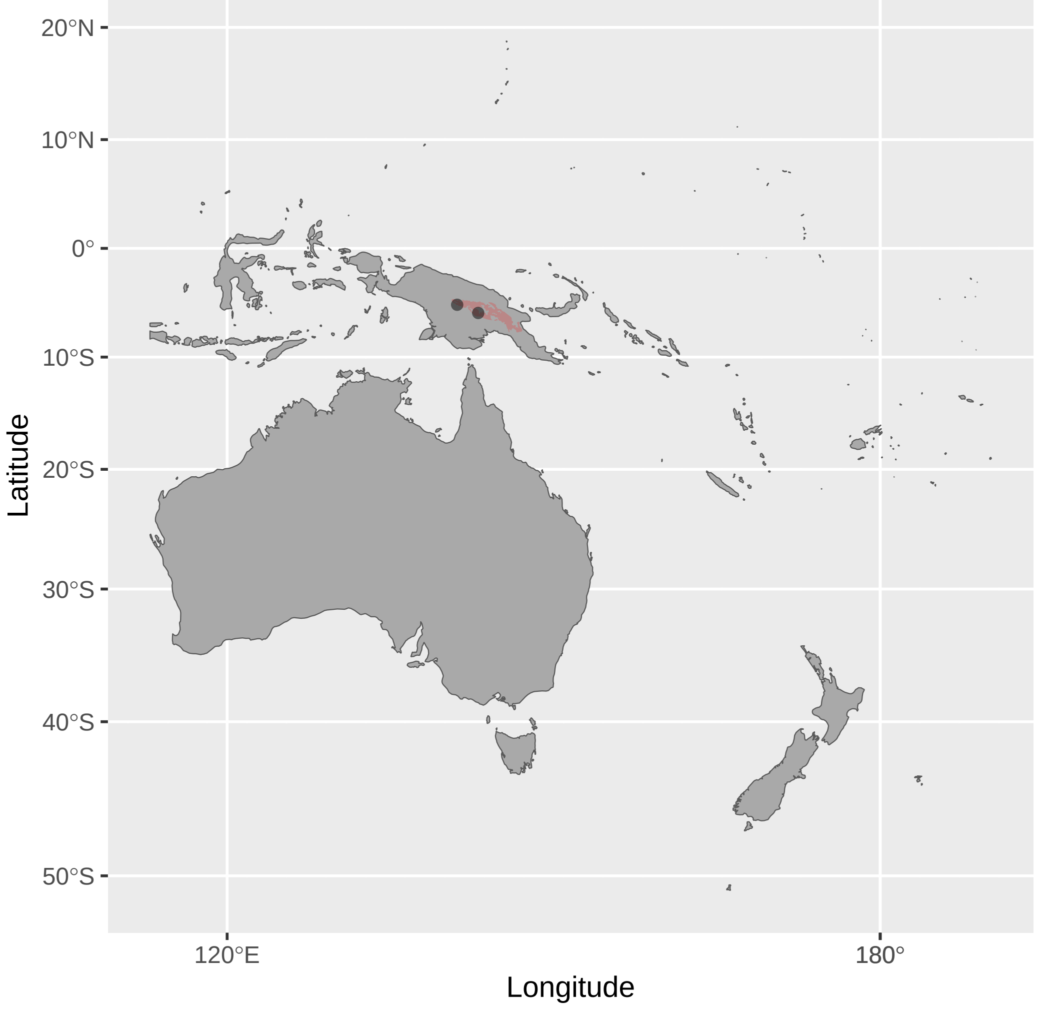


Meliphacator_provocator (n = 1)

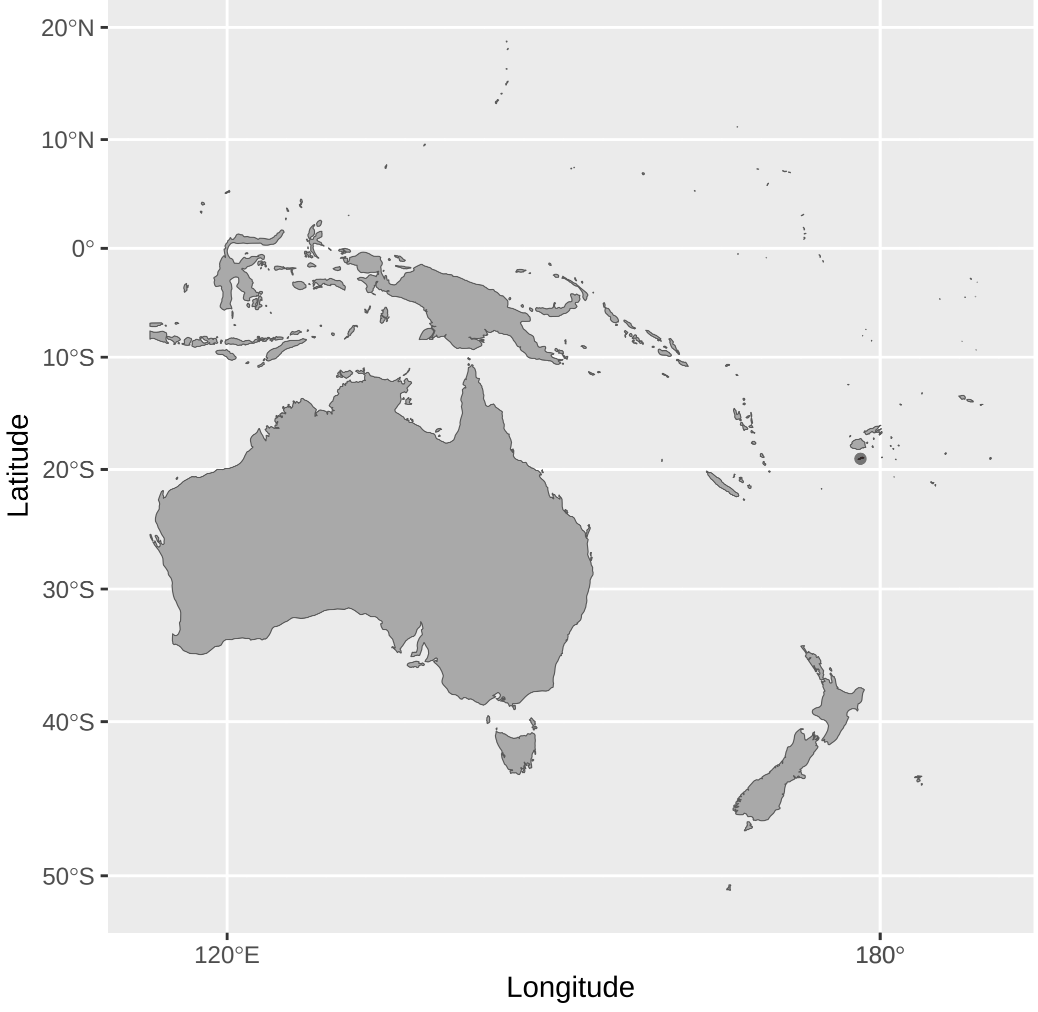


Melipotes_ater (n = 1)

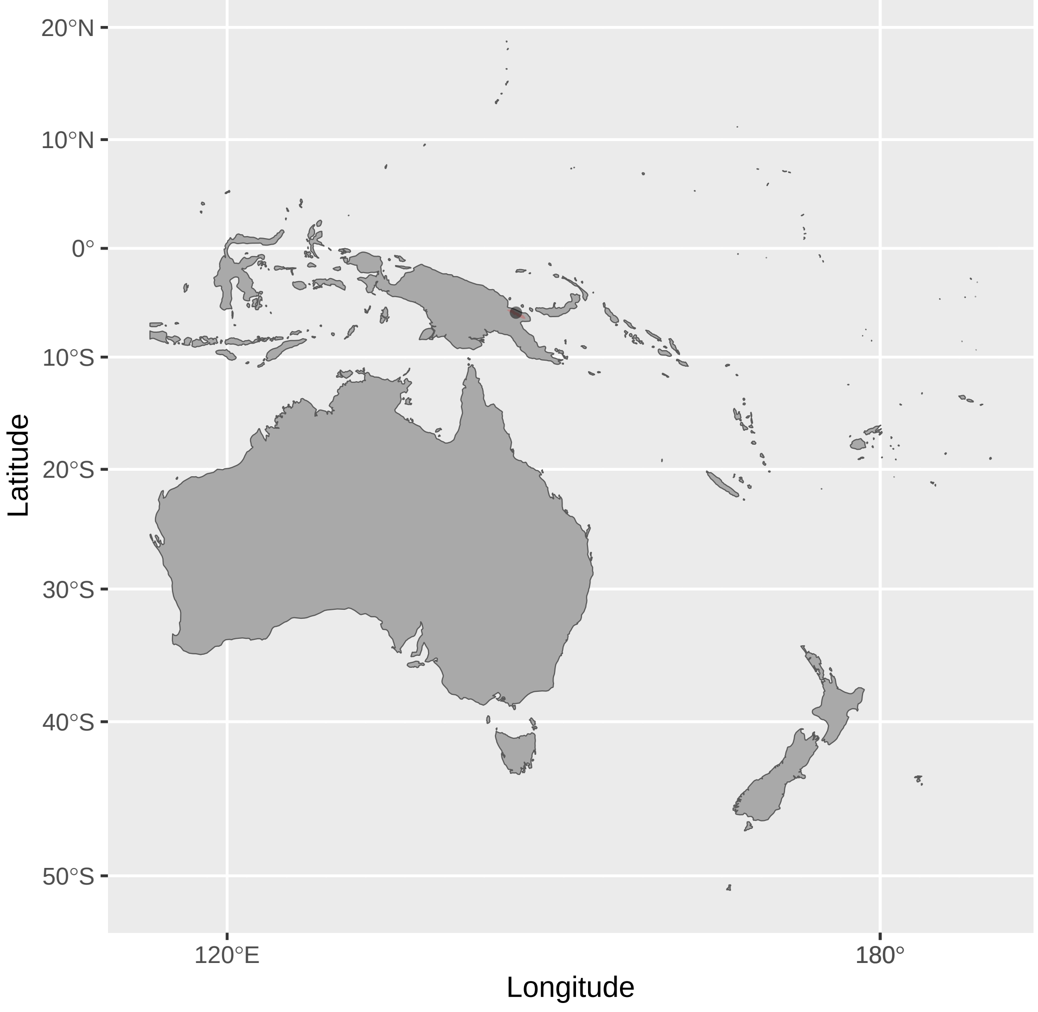


Melidectes_torquatus (n = 2)

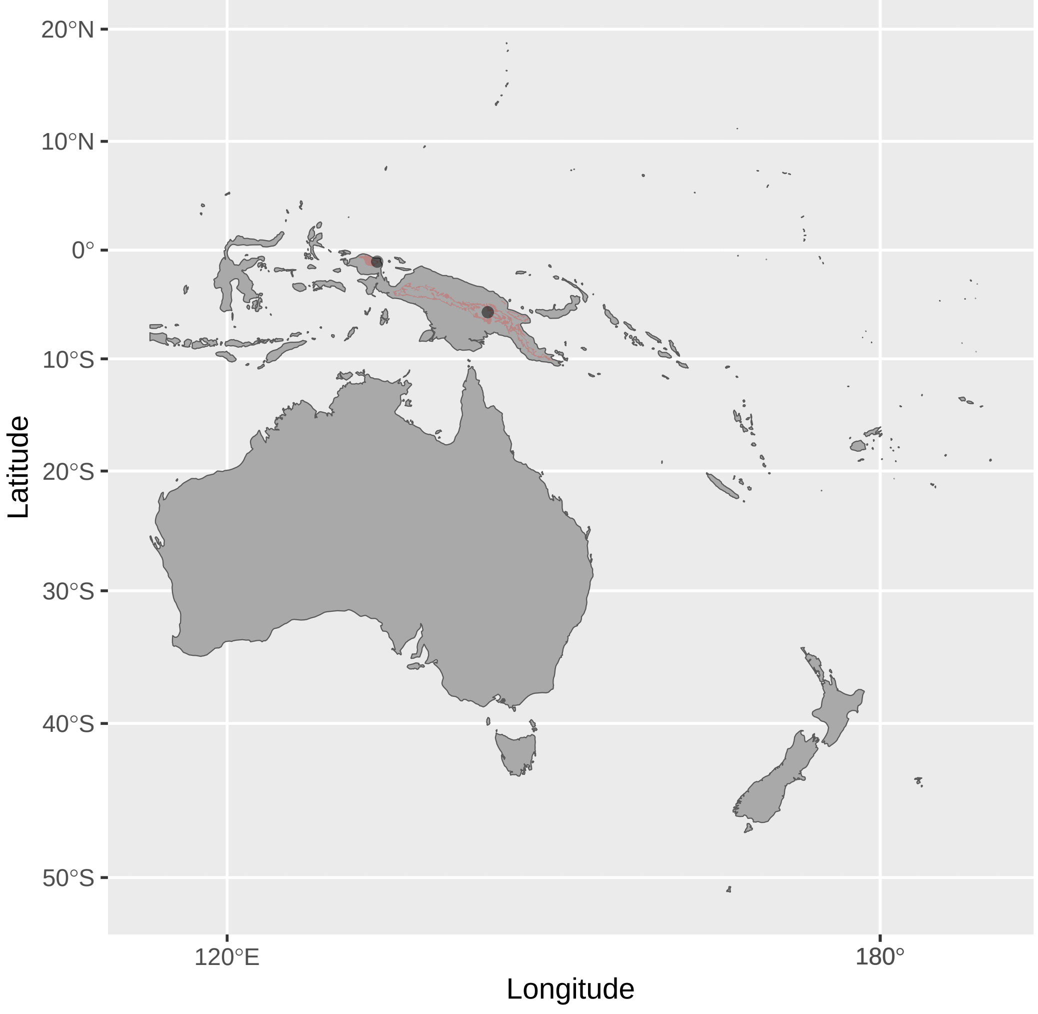


Meliphaga_aruensis (n = 1)

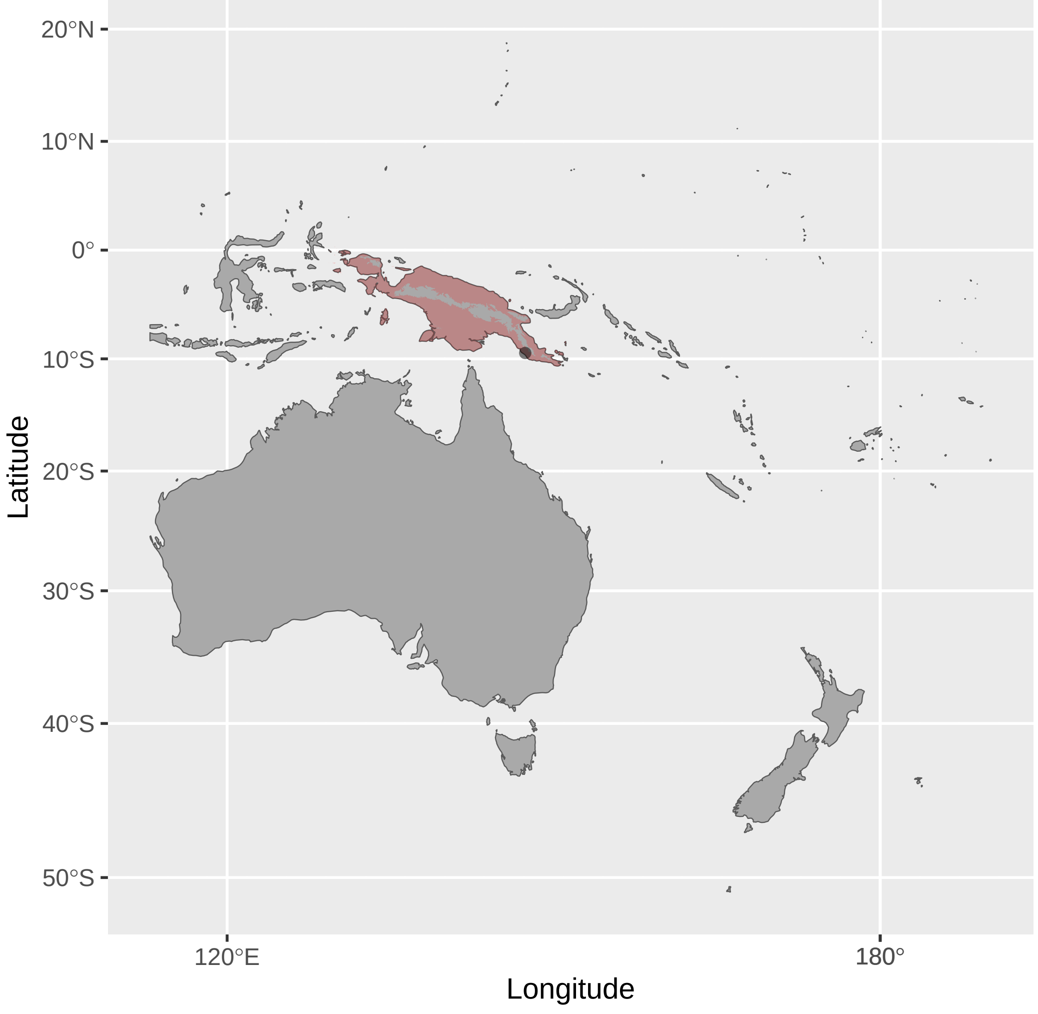


Melipotes_fumigatus (n = 1)

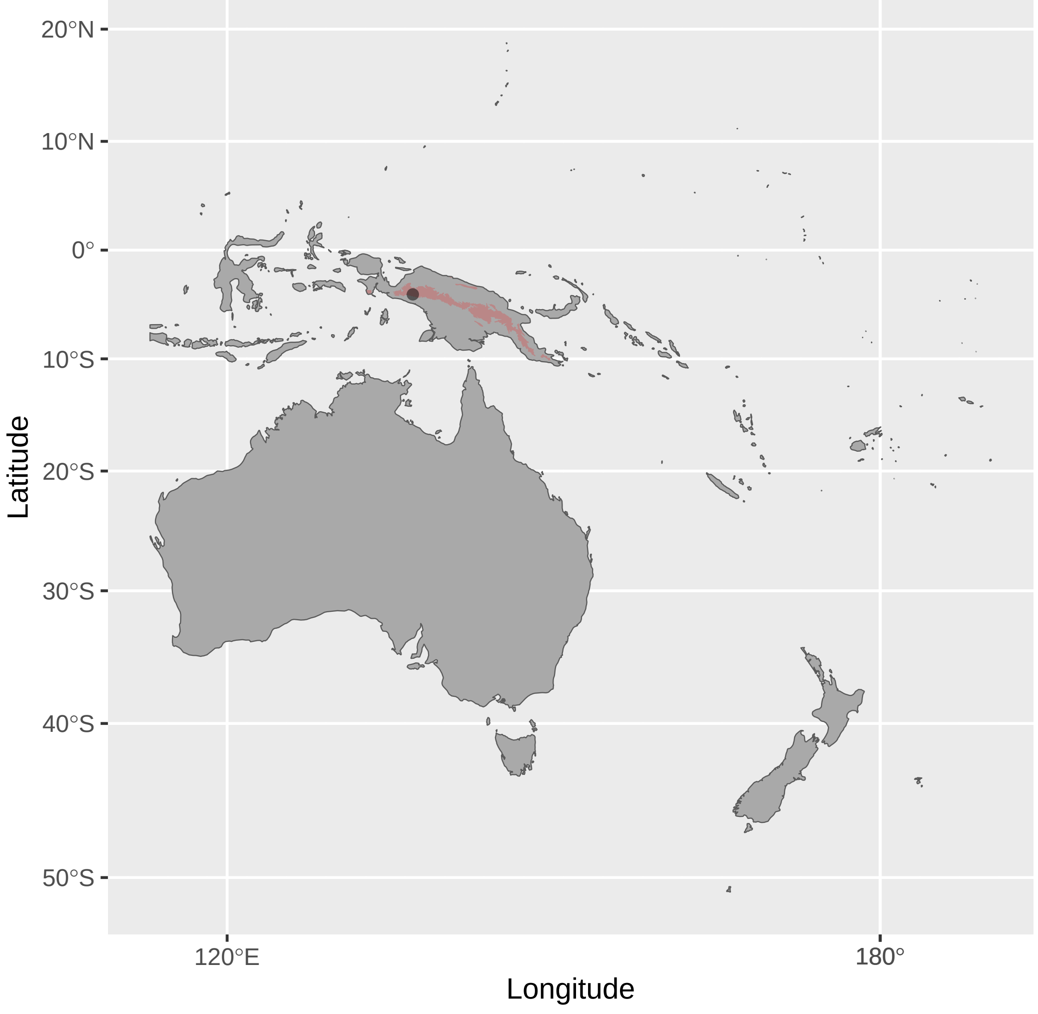


Melilestes_megarhynchus (n = 2)

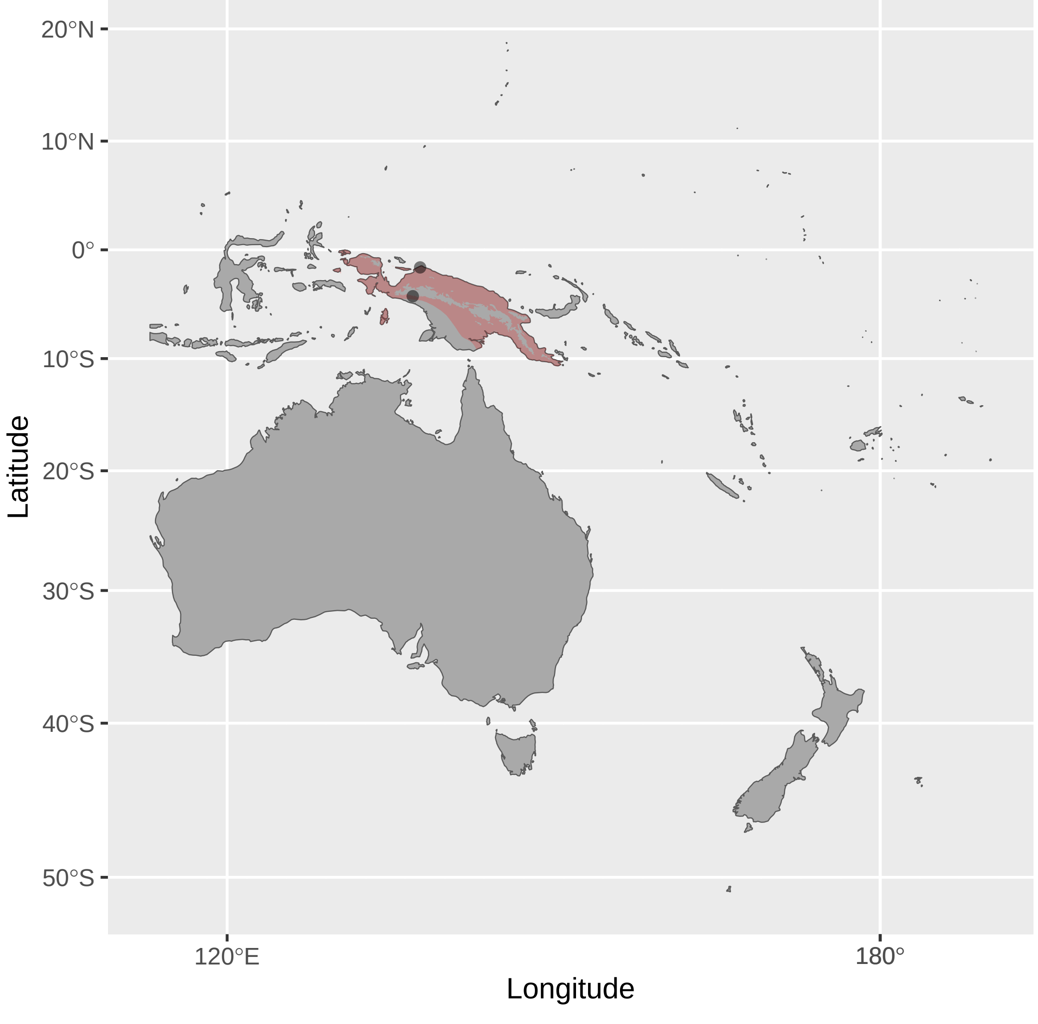


Meliphaga_lewinii (n = 20)

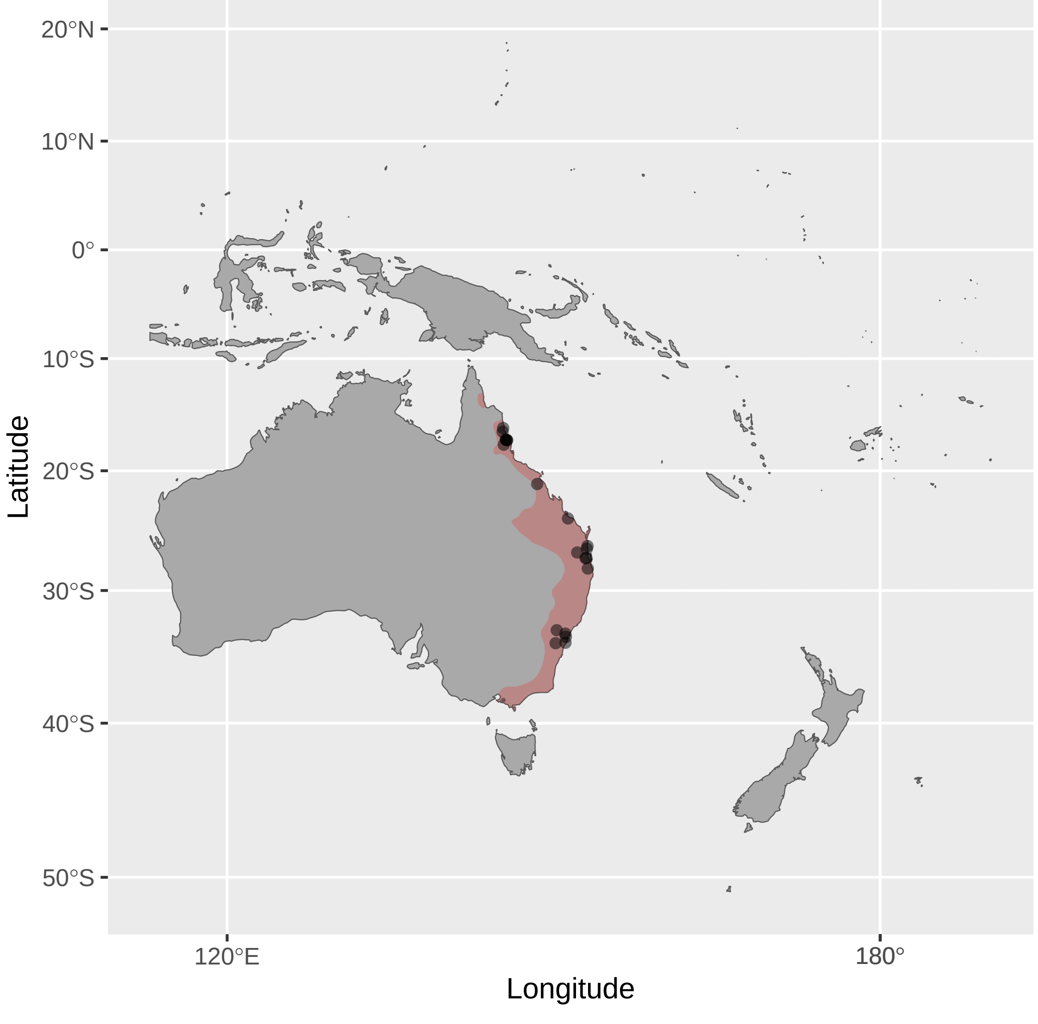


Melipotes_gymnops (n = 1)

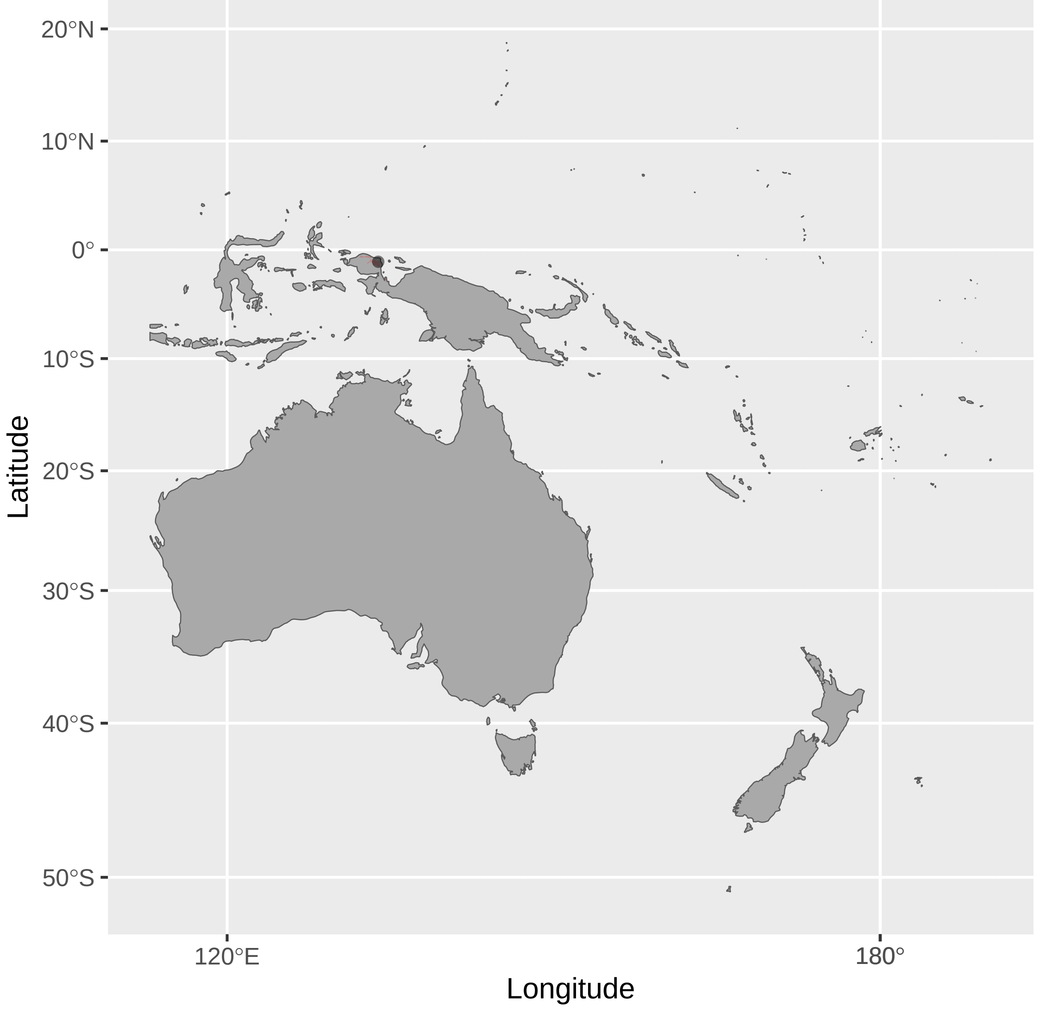

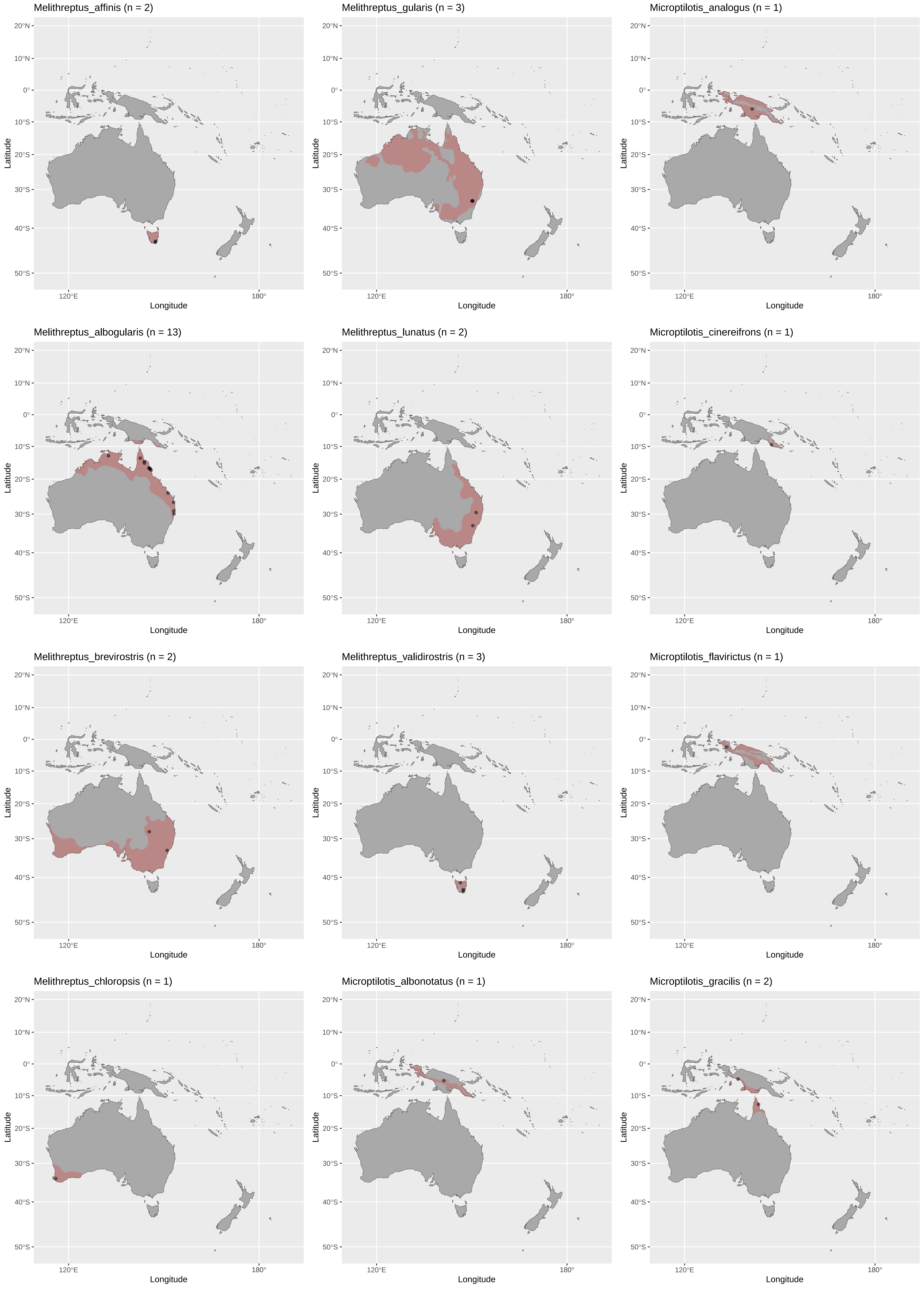
Melithreptus_affinis (n = 2)
Melithreptus_gularis (n = 3)
Microptilotis_analogus (n = 1)
Melithreptus_albogularis (n = 13)
Melithreptus_lunatus (n = 2)
Microptilotis_cinereifrons (n = 1)
Melithreptus_brevirostris (n = 2)
Melithreptus_validirostris (n = 3)
Microptilotis_flavirictus (n = 1)
Melithreptus_chloropsis (n = 1)
Microptilotis_albonotatus (n = 1)
Microptilotis_gracilis (n = 2)
20°N
10°N
0°
10°S
20°S
30°S
40°S
50°S
Latitude
120°E
180°
Longitude

Microptilotis_imitatrix (n = 1)

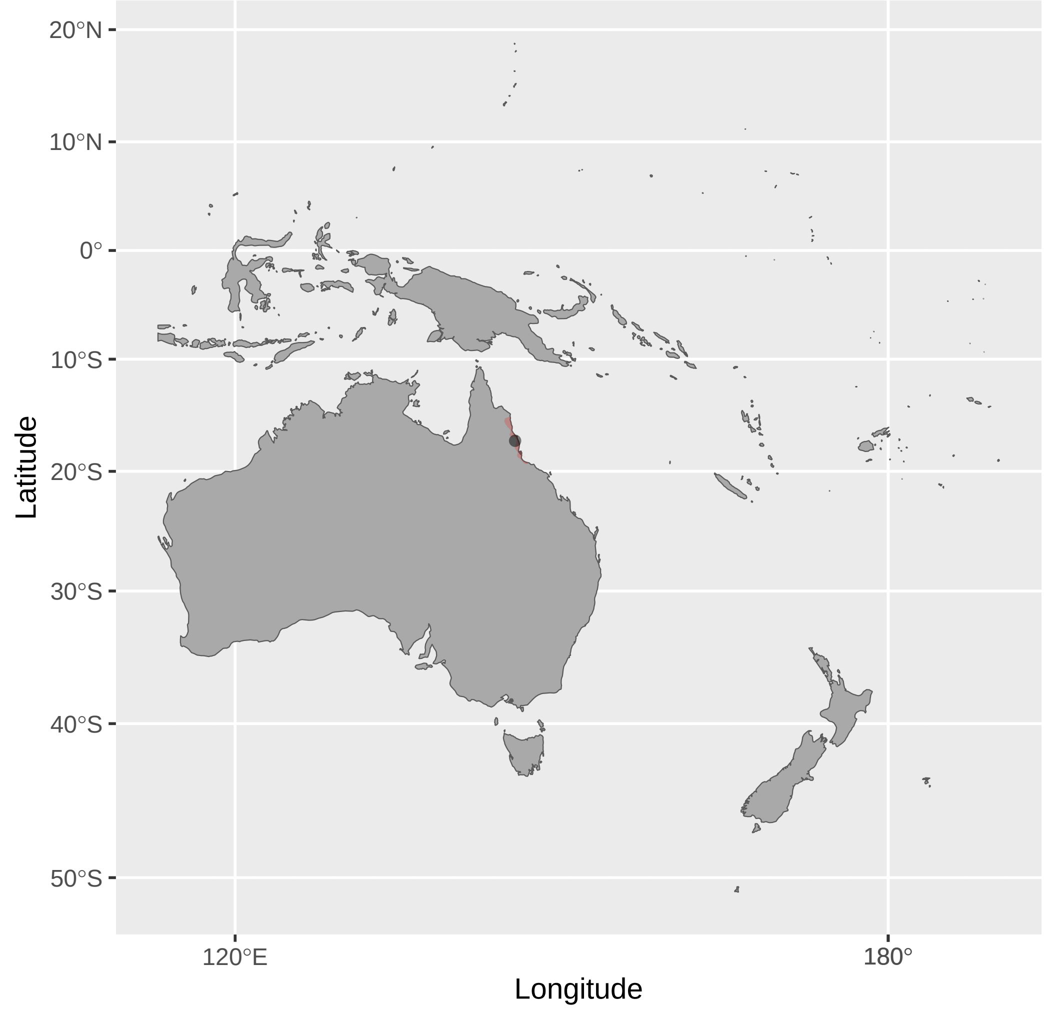


Myzomela_adolphinae (n = 1)

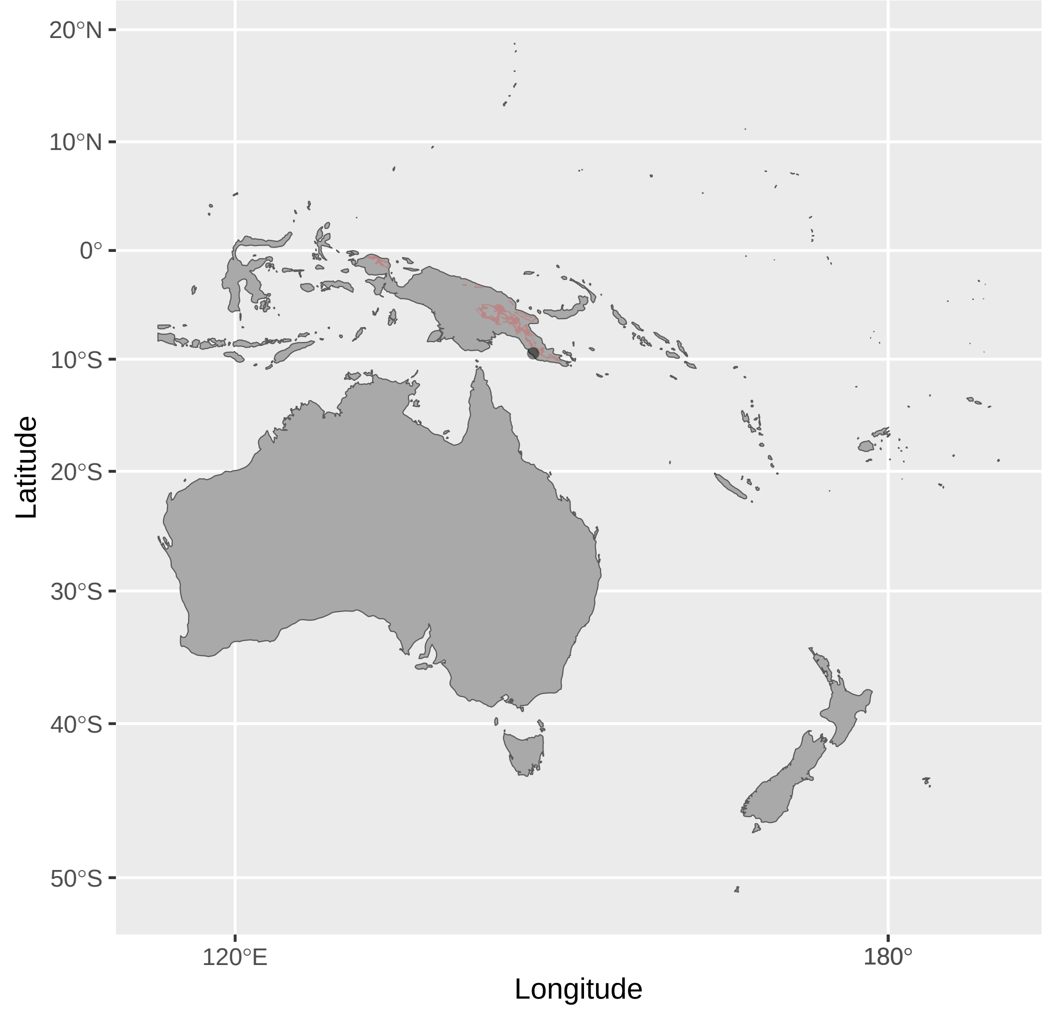


Myzomela_caledonica (n = 1)

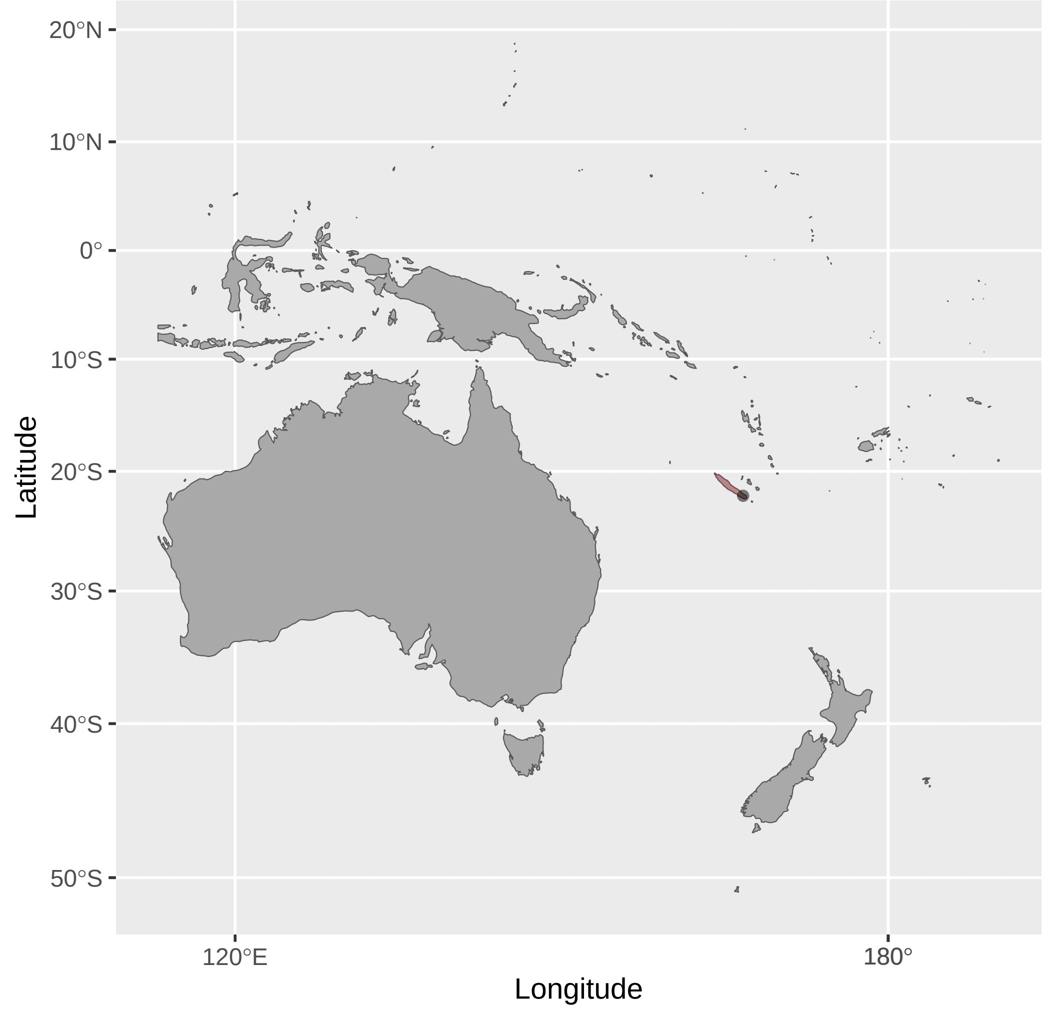


Microptilotis_orientalis (n = 1)

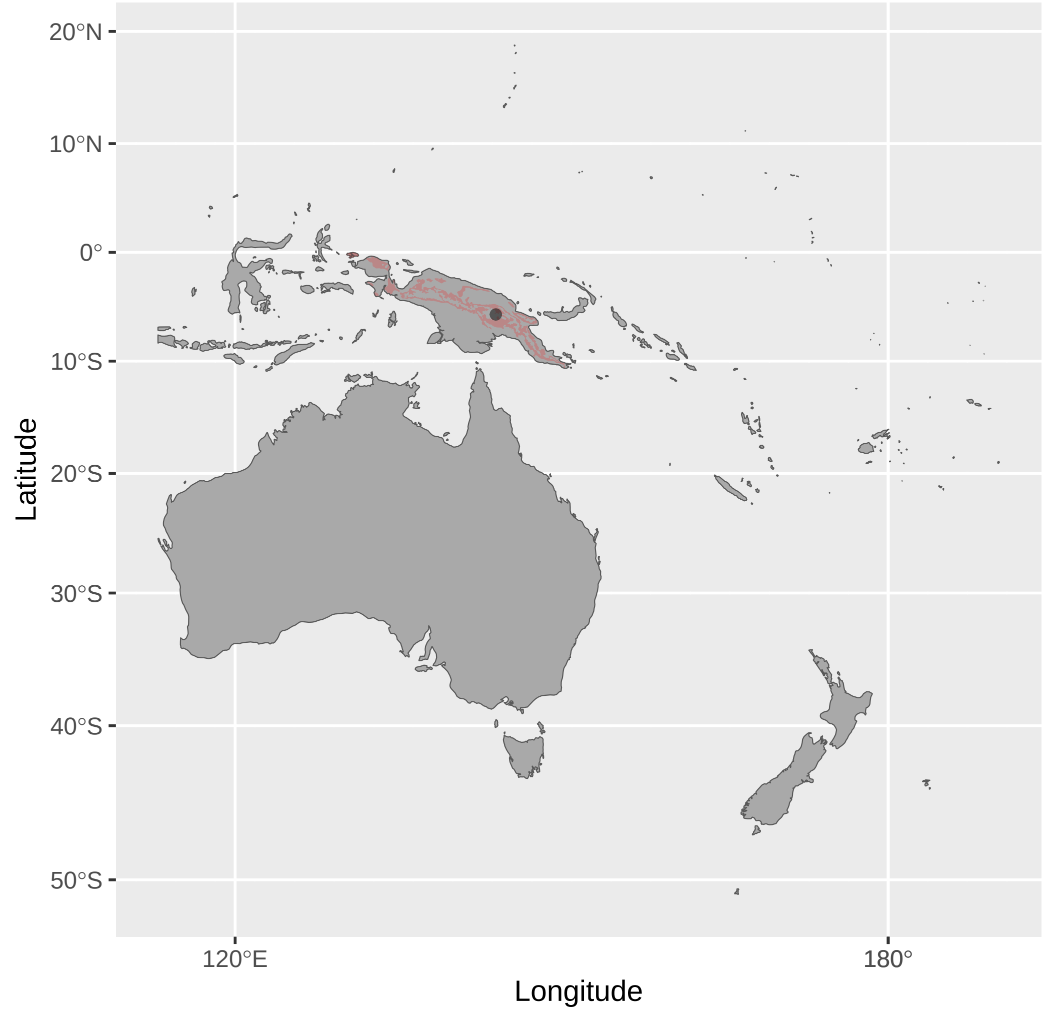


Myzomela_albigula (n = 1)

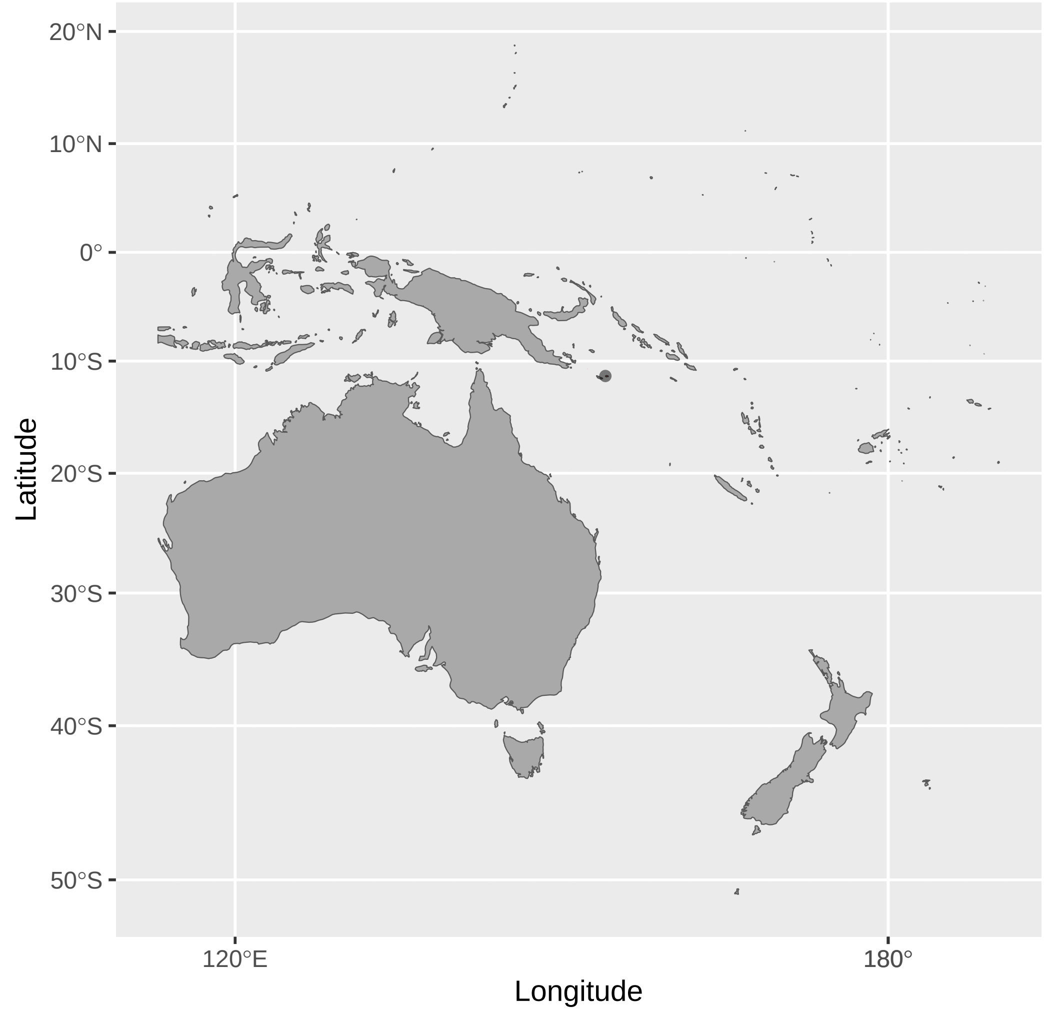


Myzomela_cardinalis (n = 3)

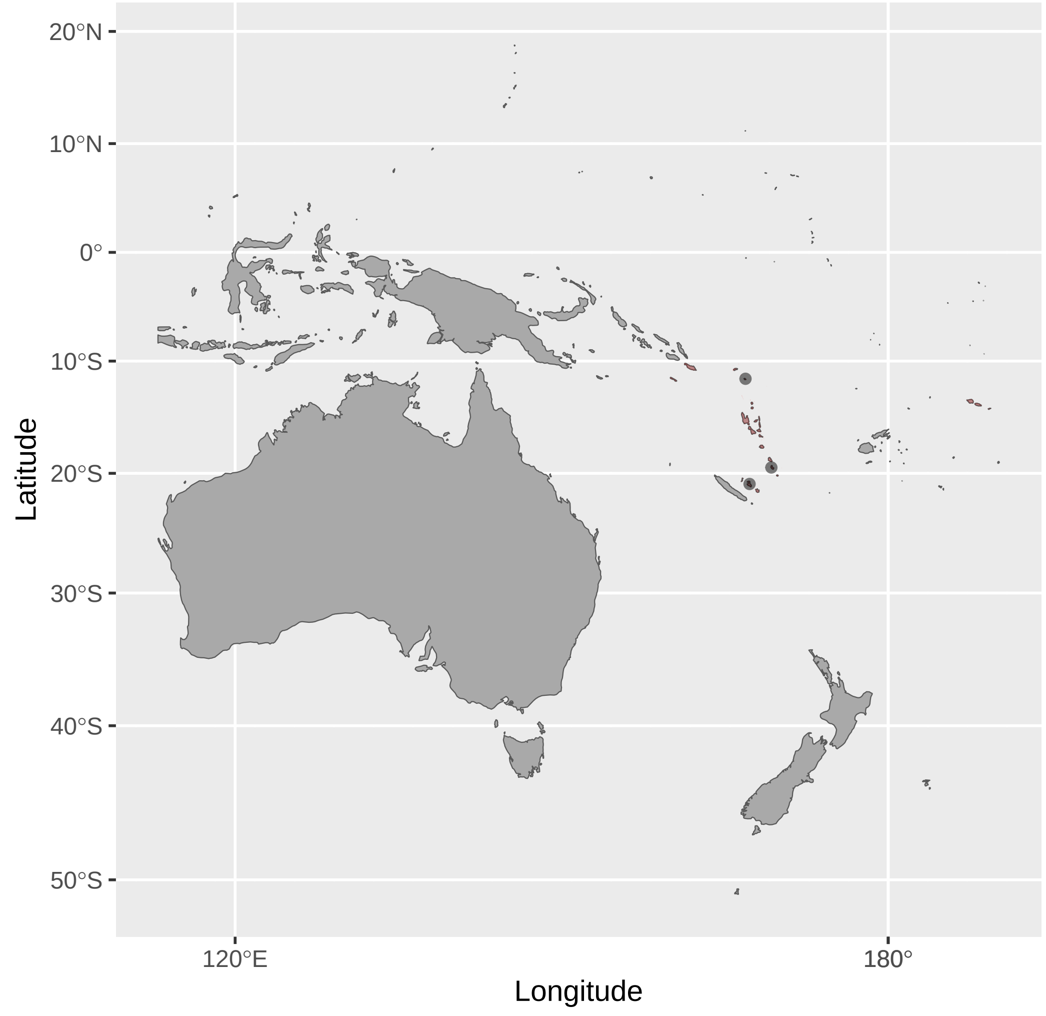


Microptilotis_vicina (n = 1)

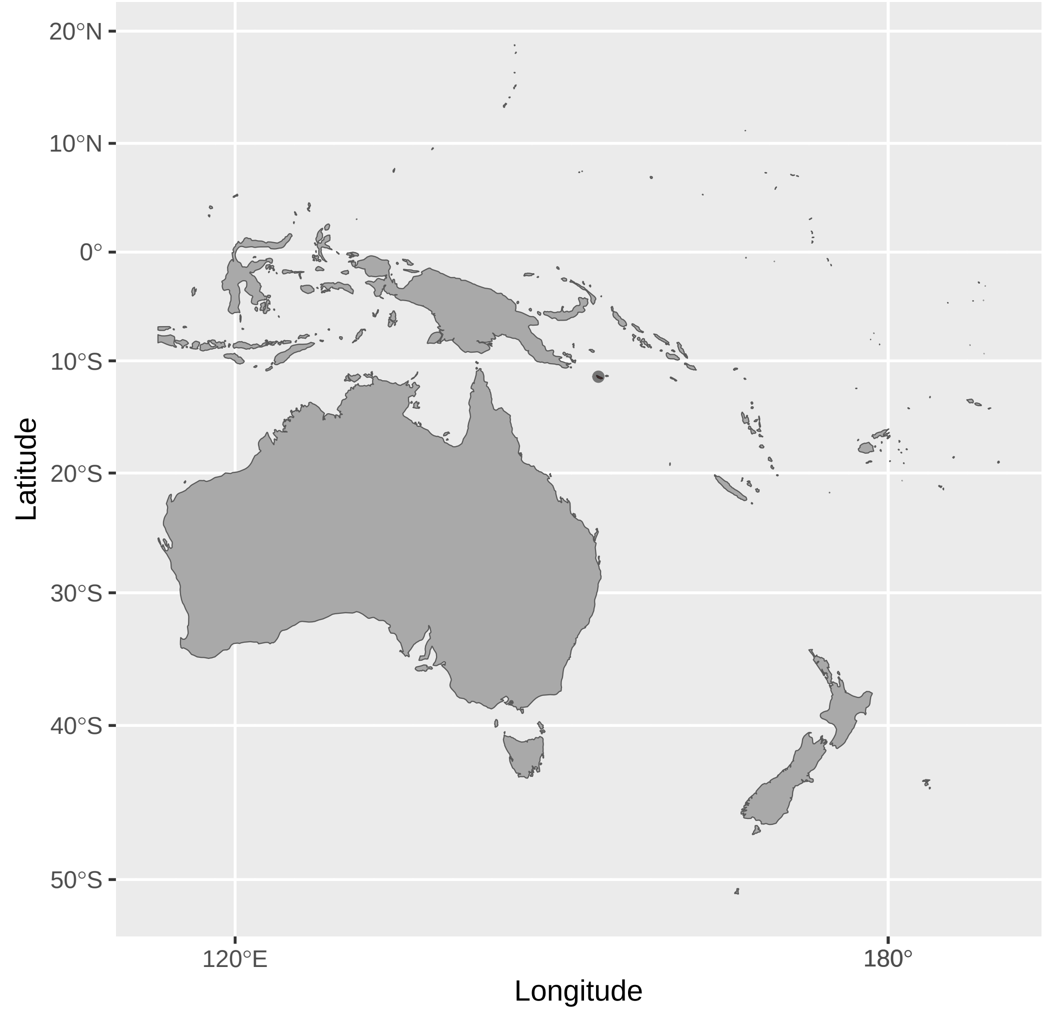


Myzomela_blasii (n = 1)

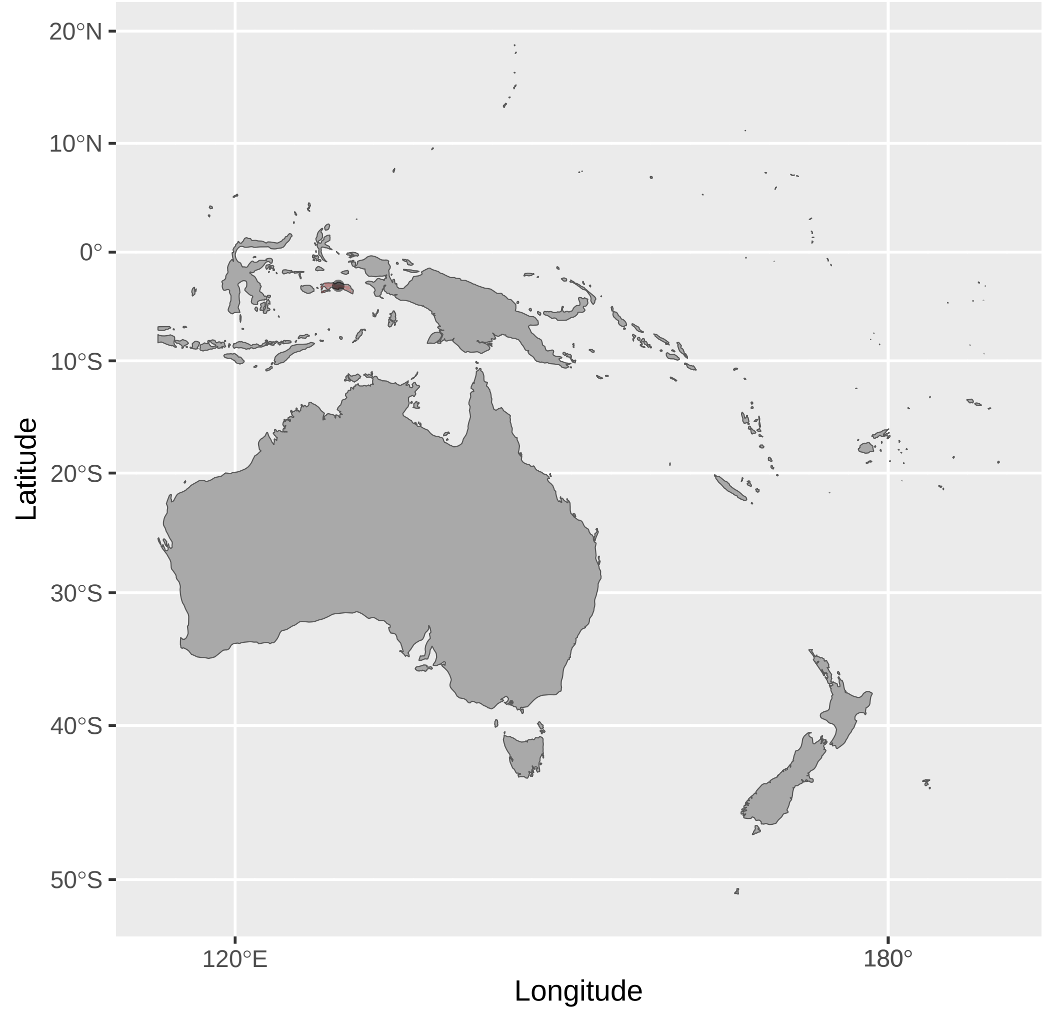


Myzomela_chloroptera (n = 4)

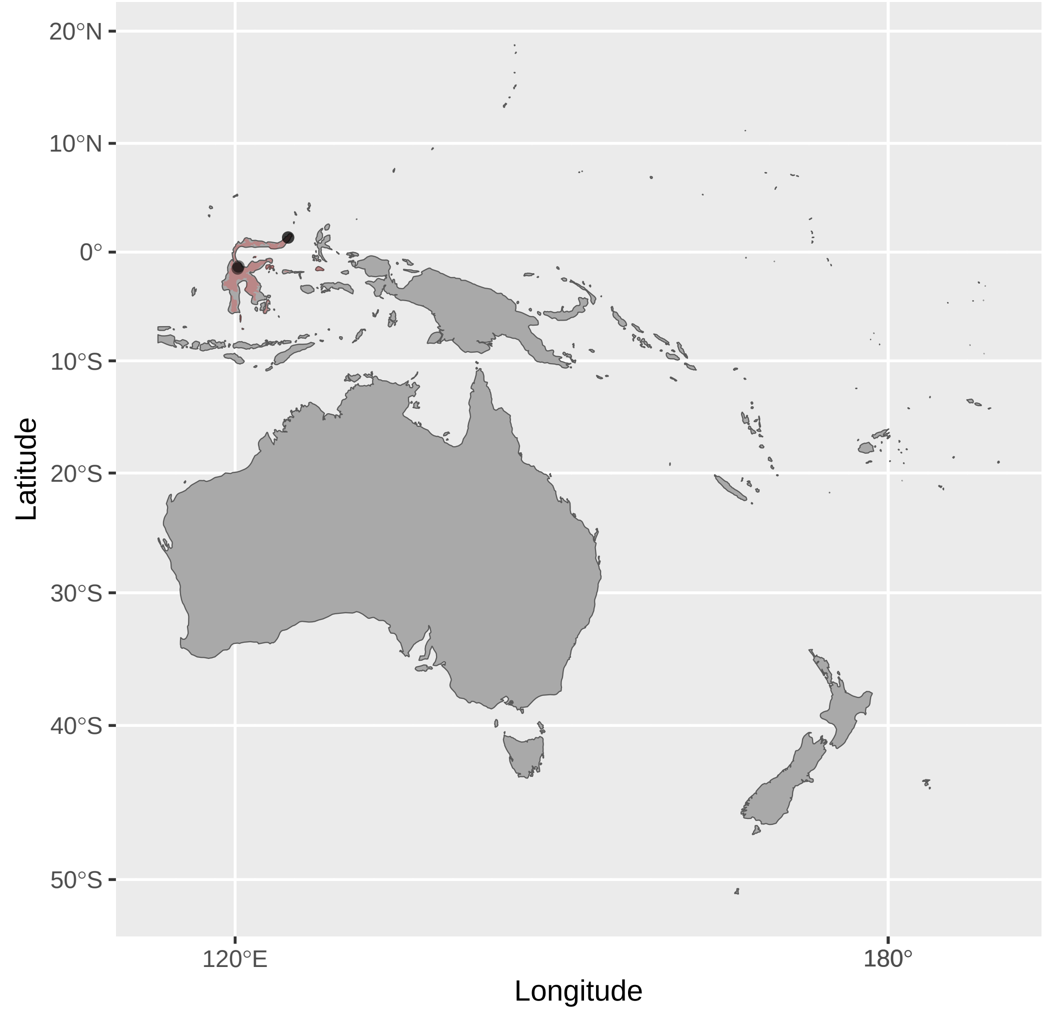


Myza_sarasinorum (n = 1)

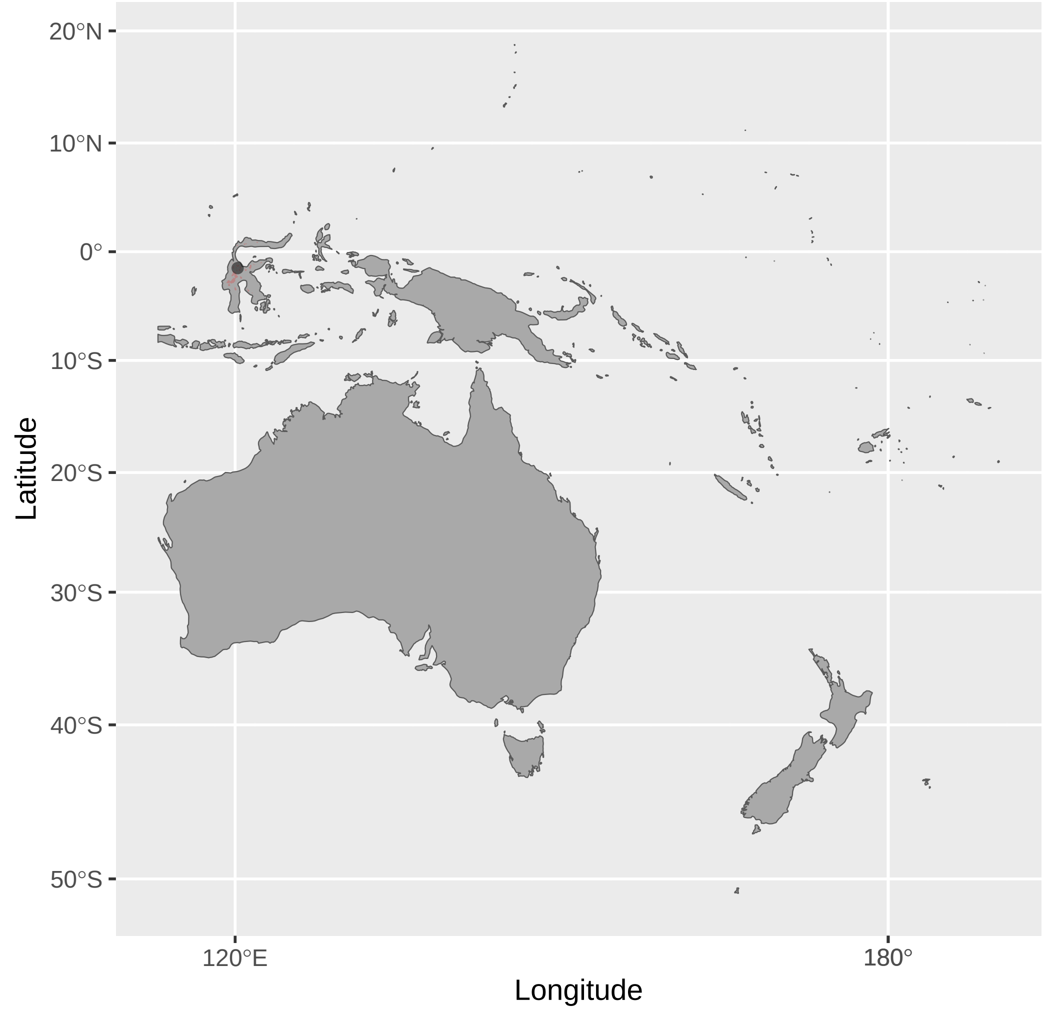


Myzomela_boiei (n = 2)

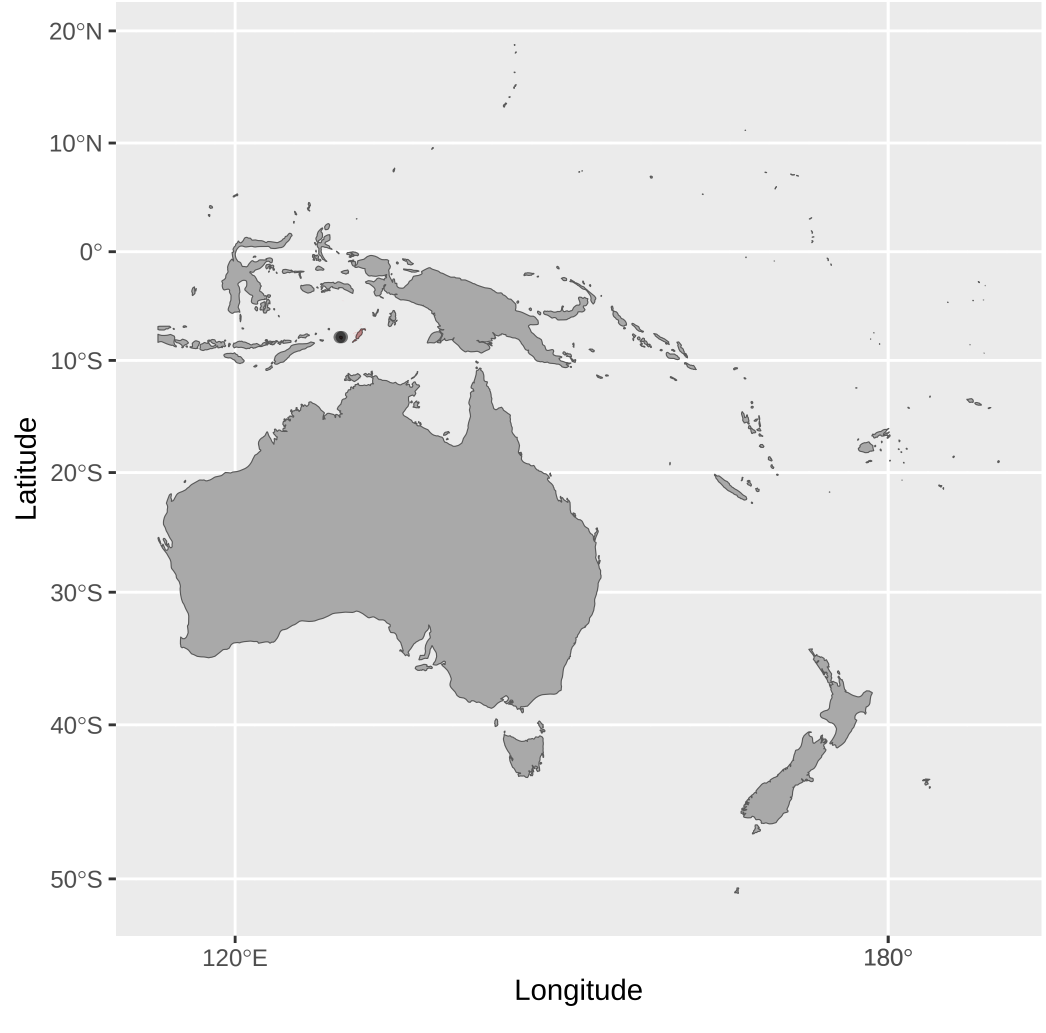


Myzomela_dammermani (n = 3)

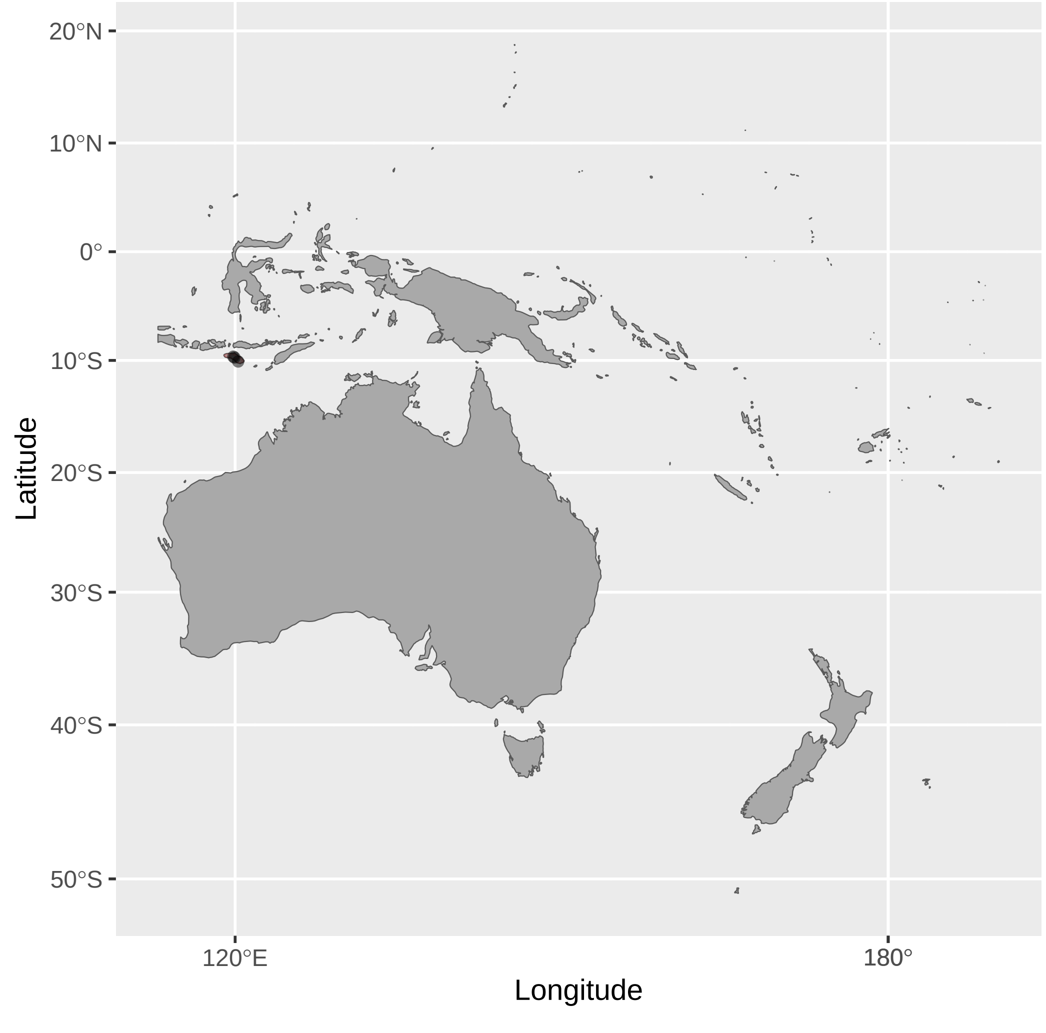

Myzomela_eichhorni (n = 1)

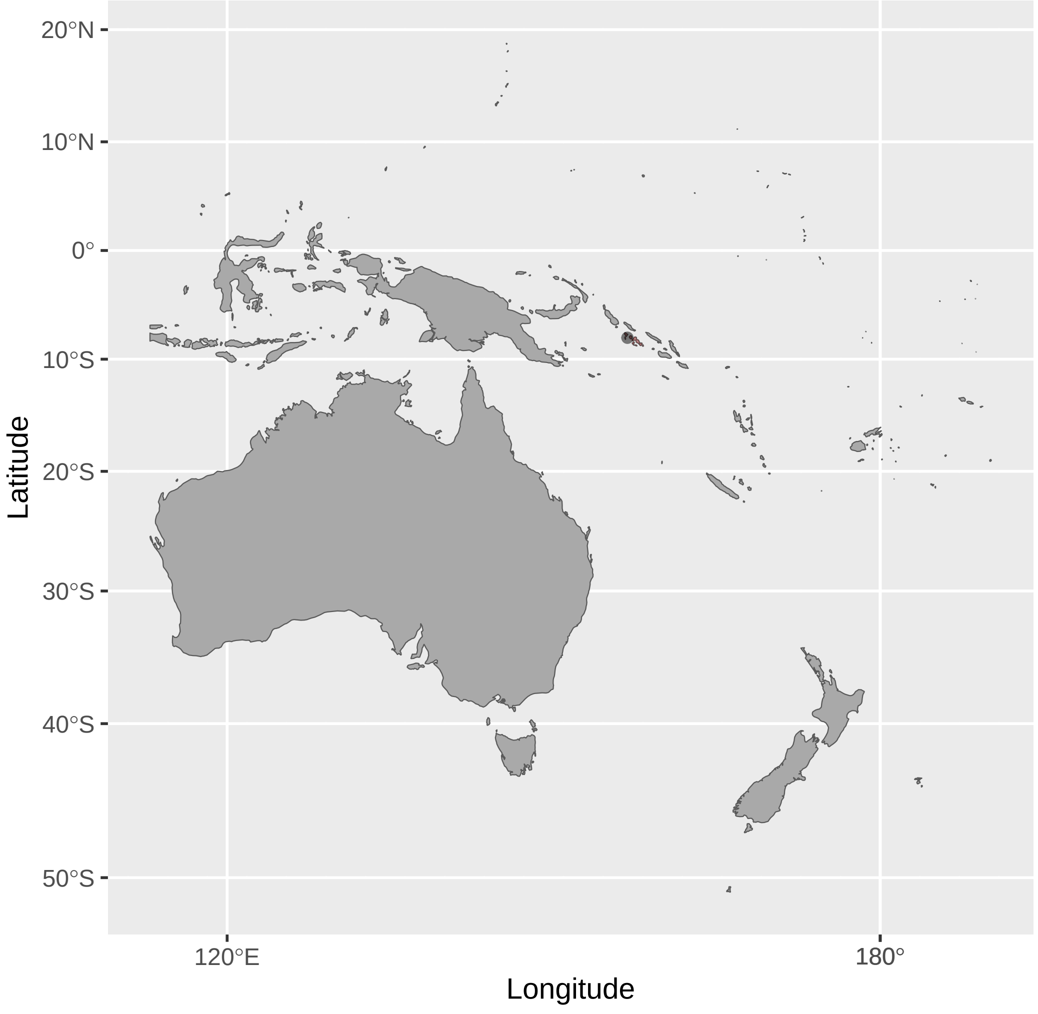


Myzomela_kuehni (n = 1)

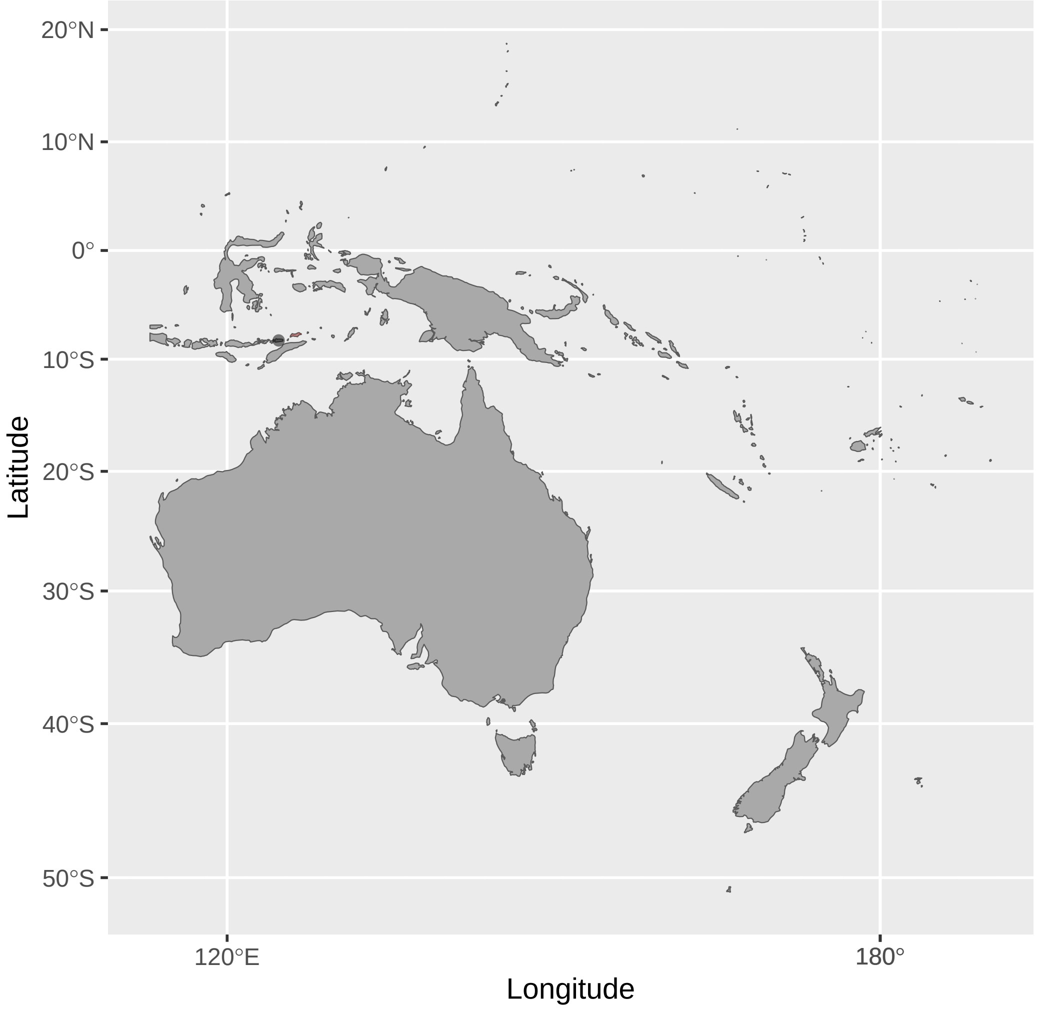


Myzomela_pammelaena (n = 1)

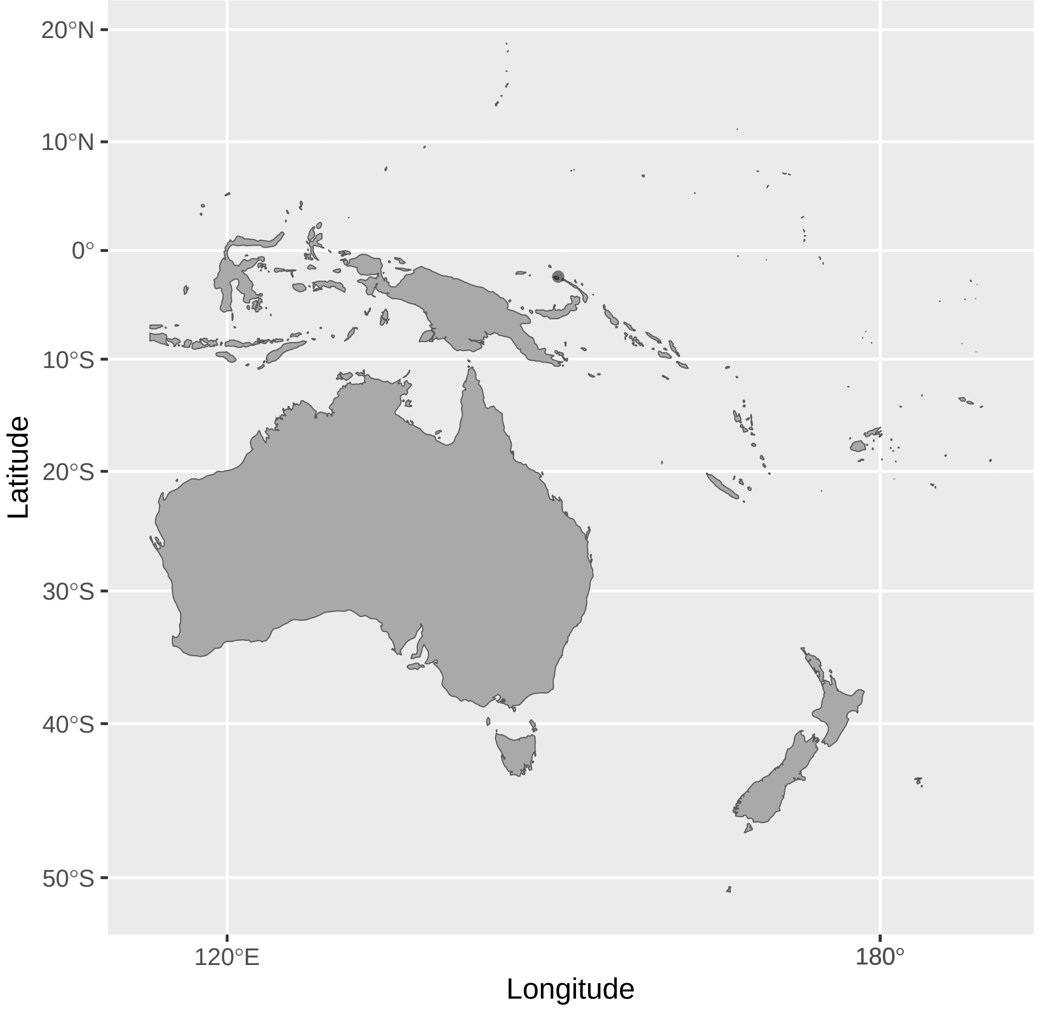


Myzomela_erythrocephala (n = 2)

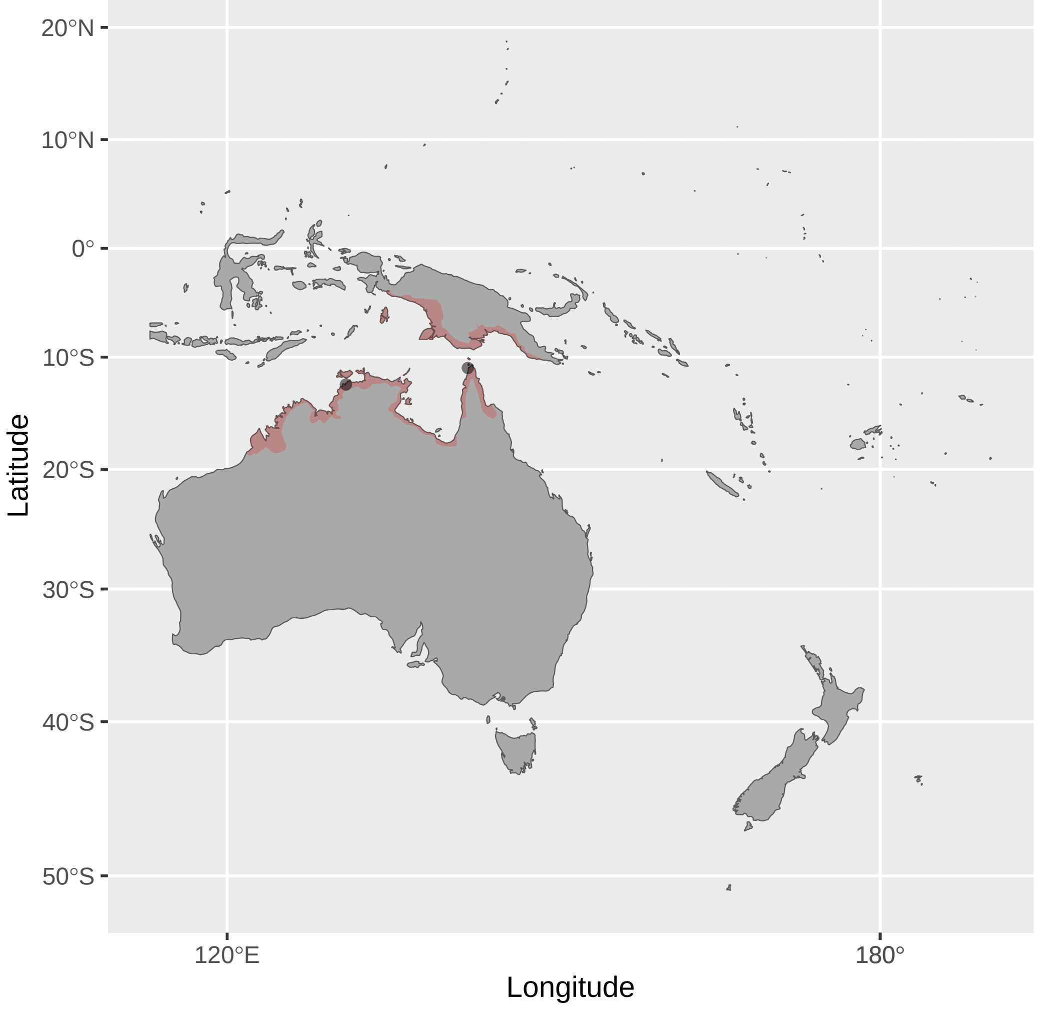


Myzomela_lafargei (n = 1)

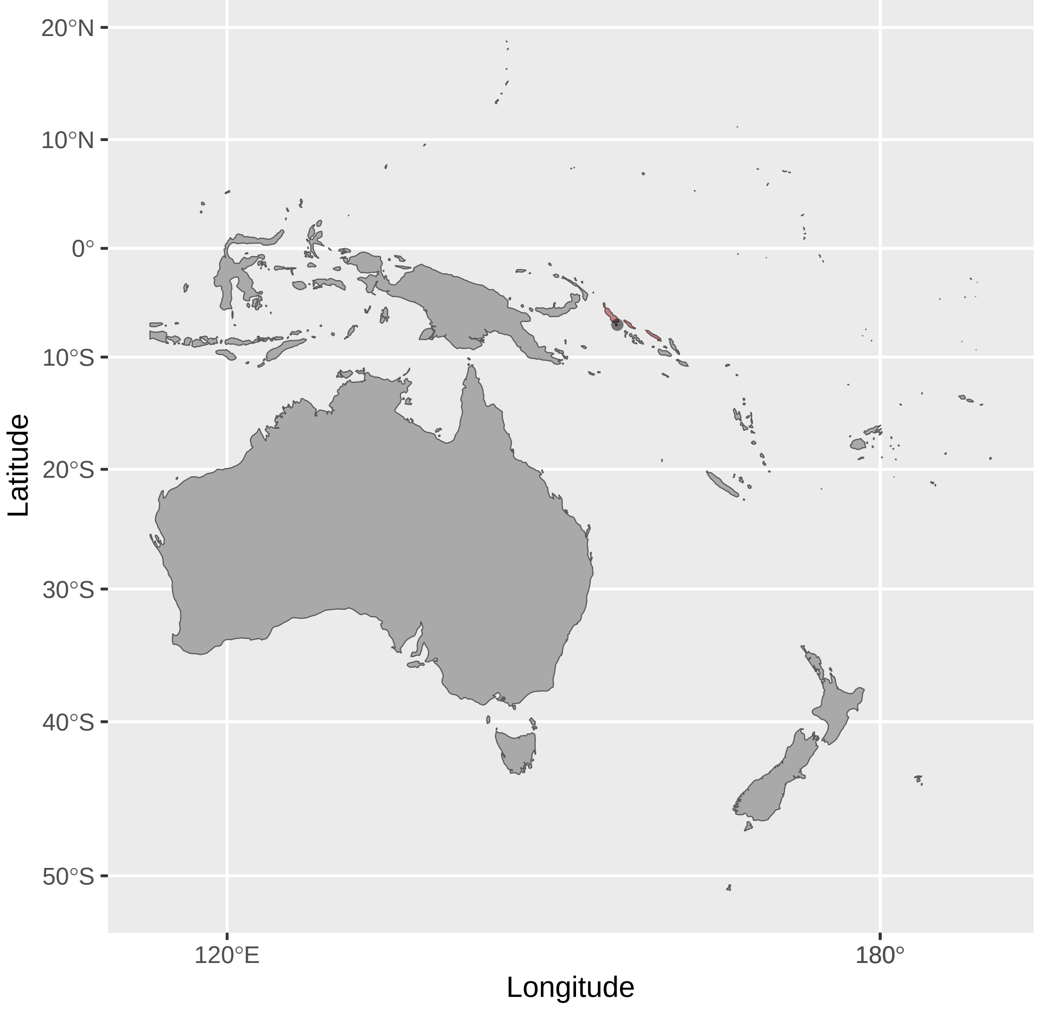


Myzomela_prawiradilagae (n = 5)

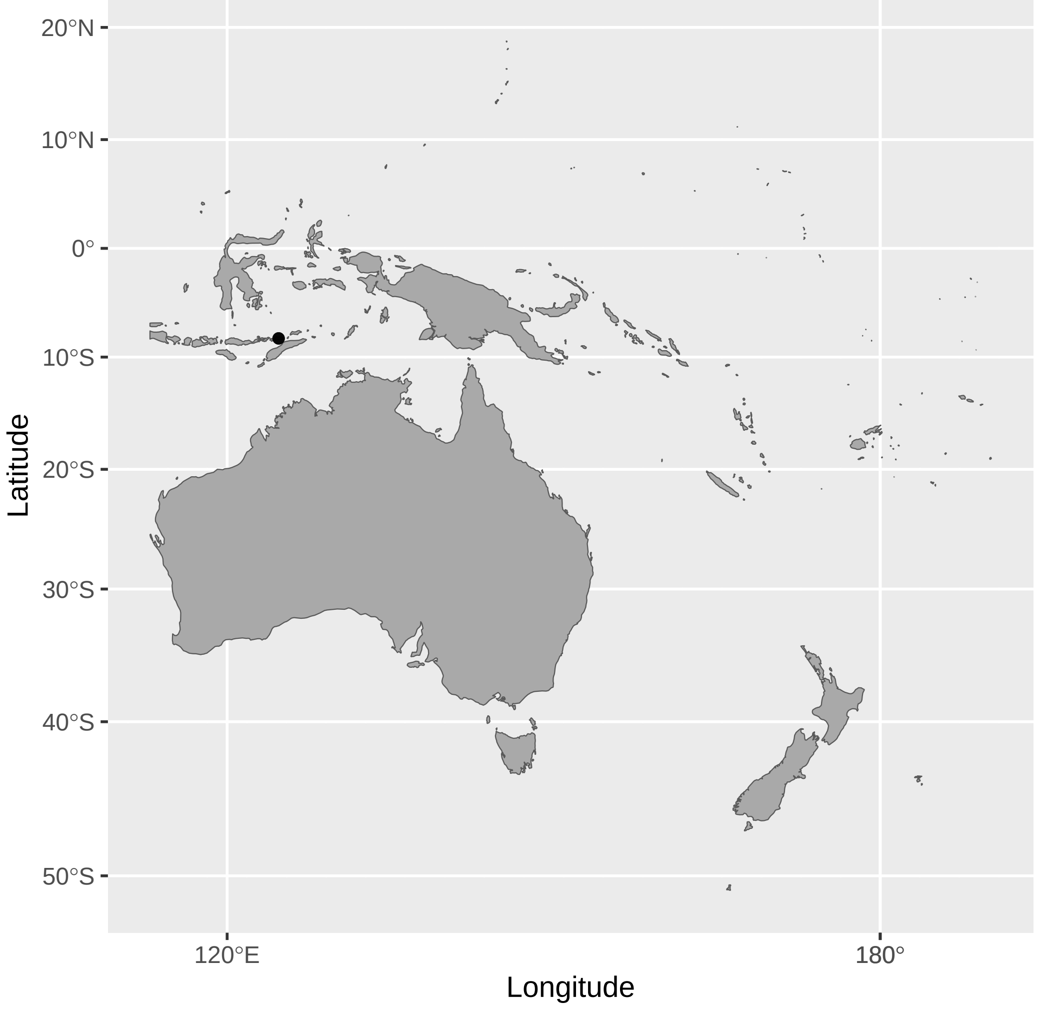


Myzomela_irianawidodoae (n = 3)

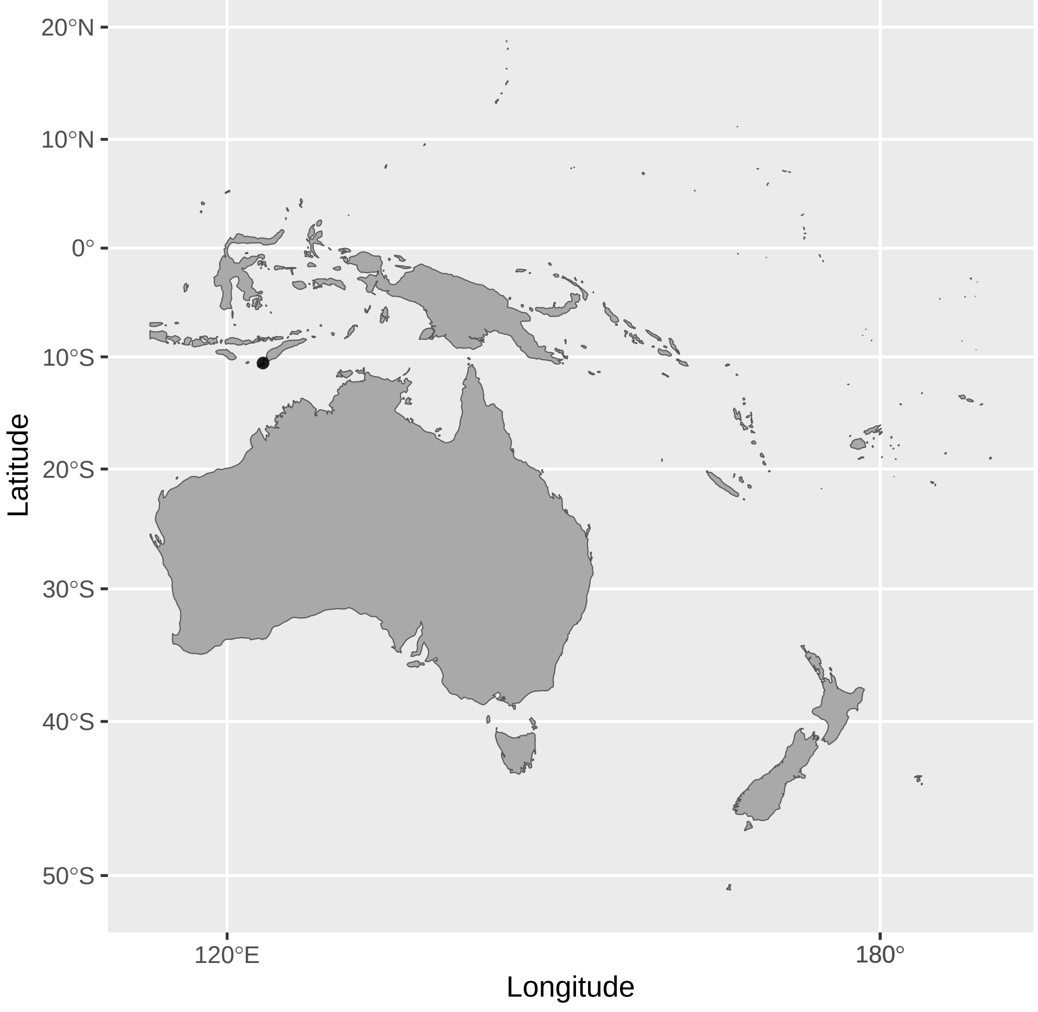


Myzomela_melanocephala (n = 1)

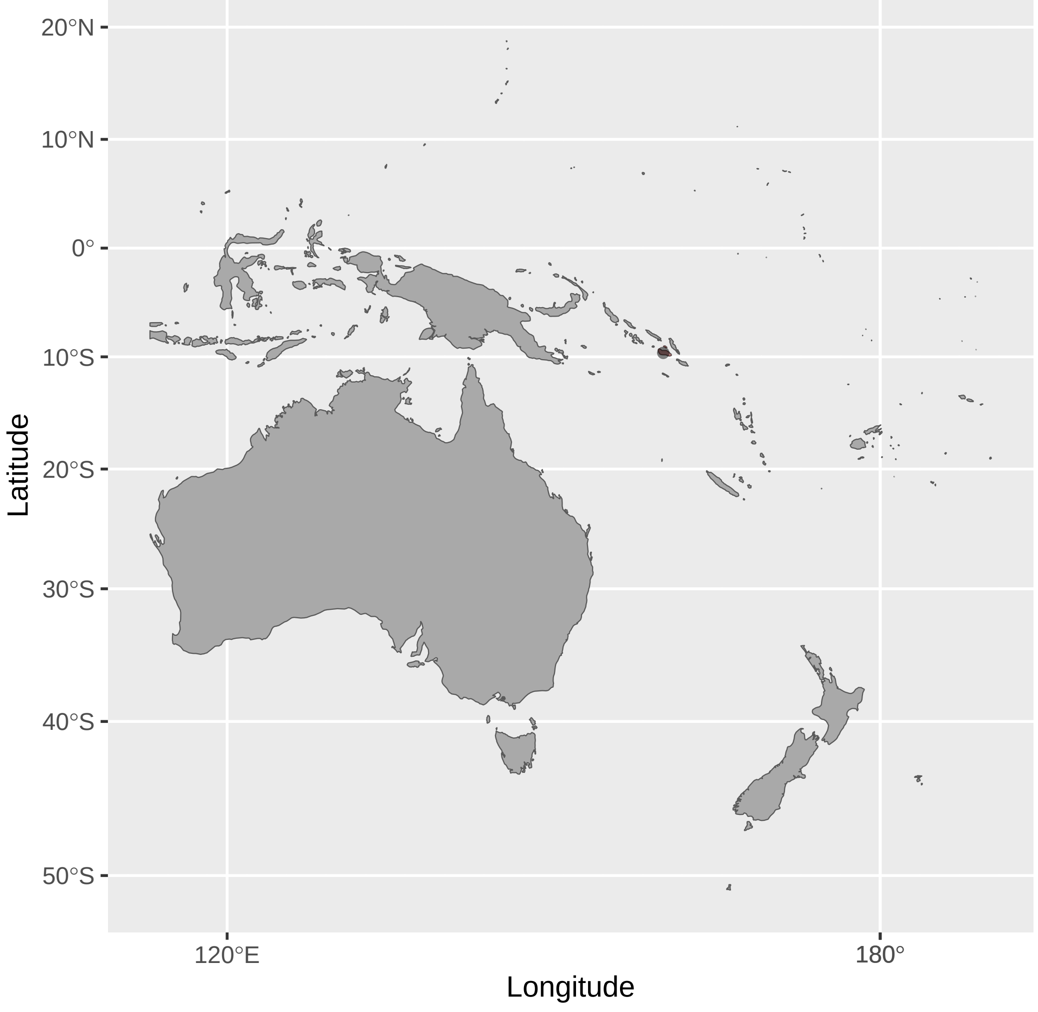


Myzomela_rosenbergii (n = 6)

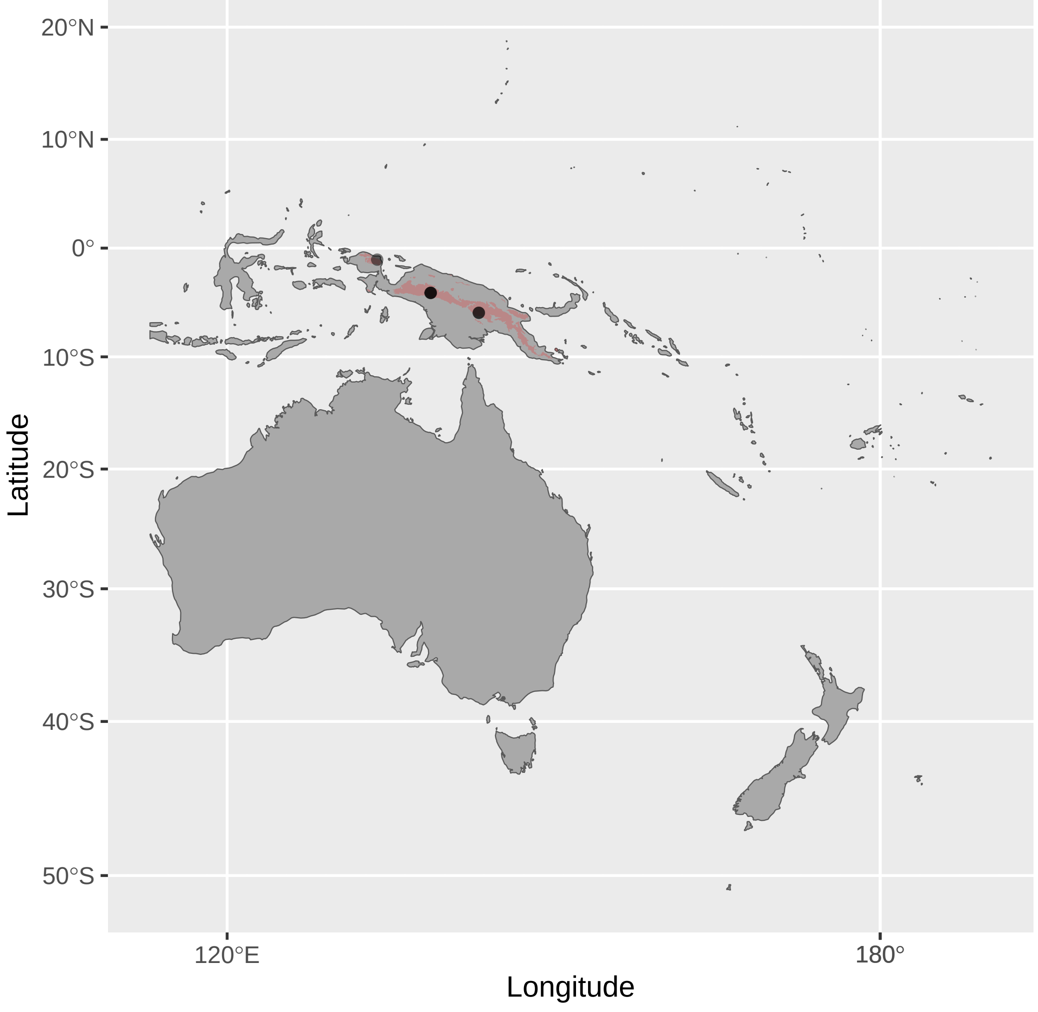


Myzomela_jugularis (n = 2)

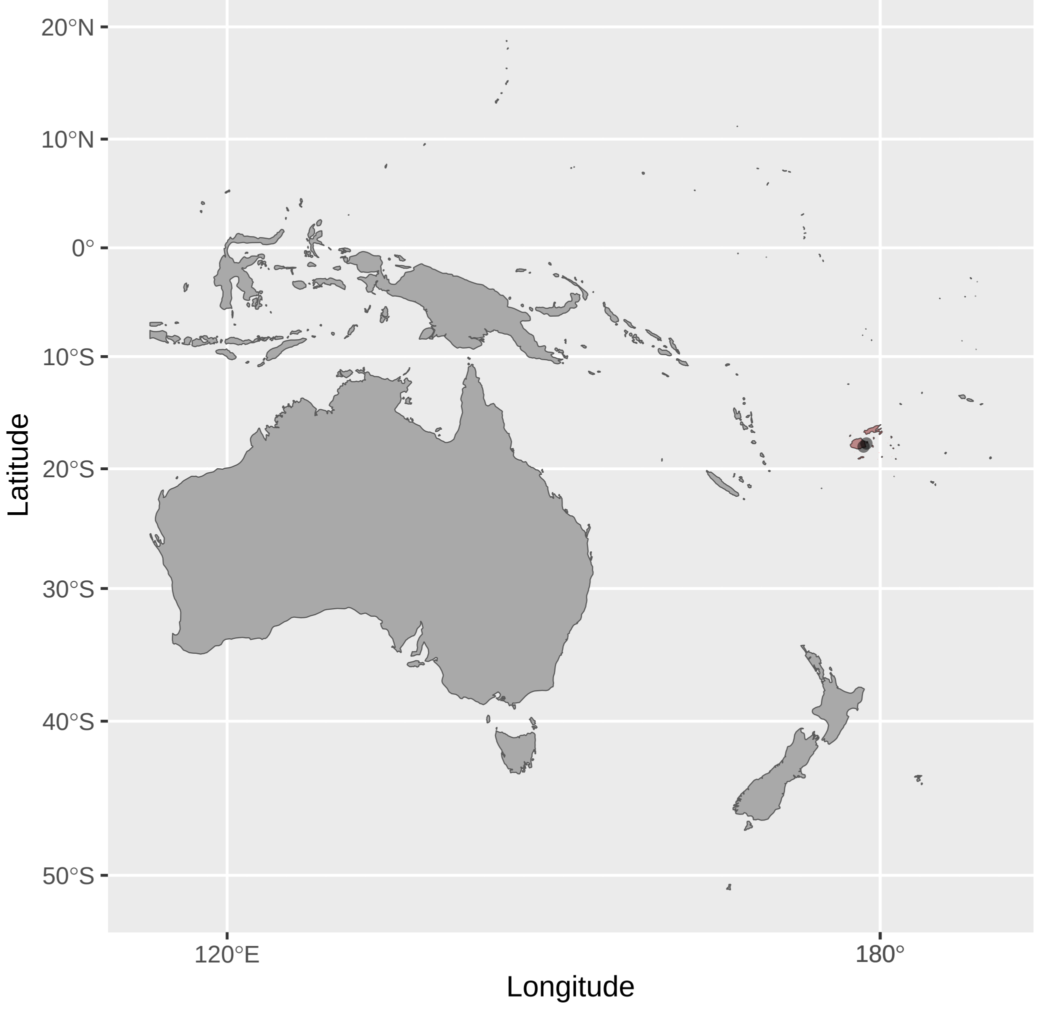


Myzomela_obscura (n = 11)

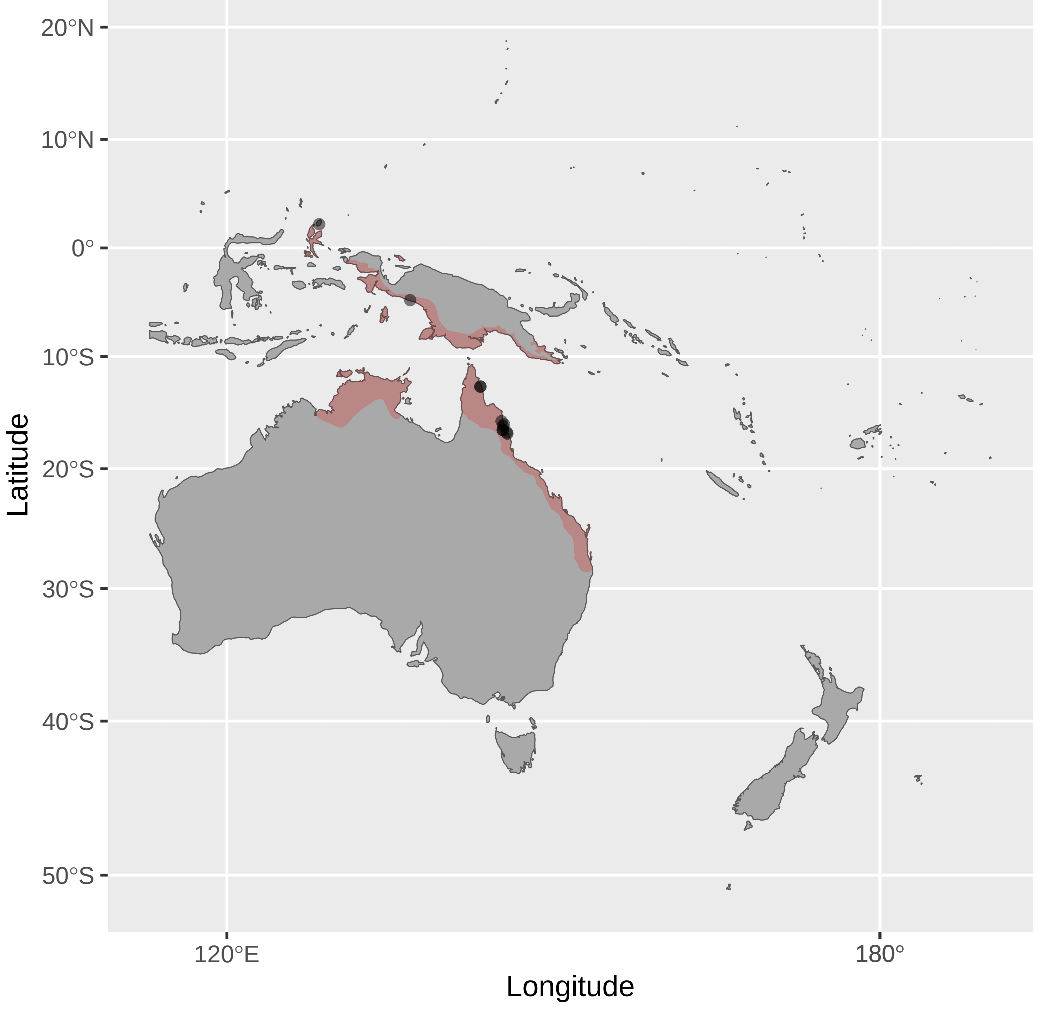


Myzomela_rubratra (n = 3)

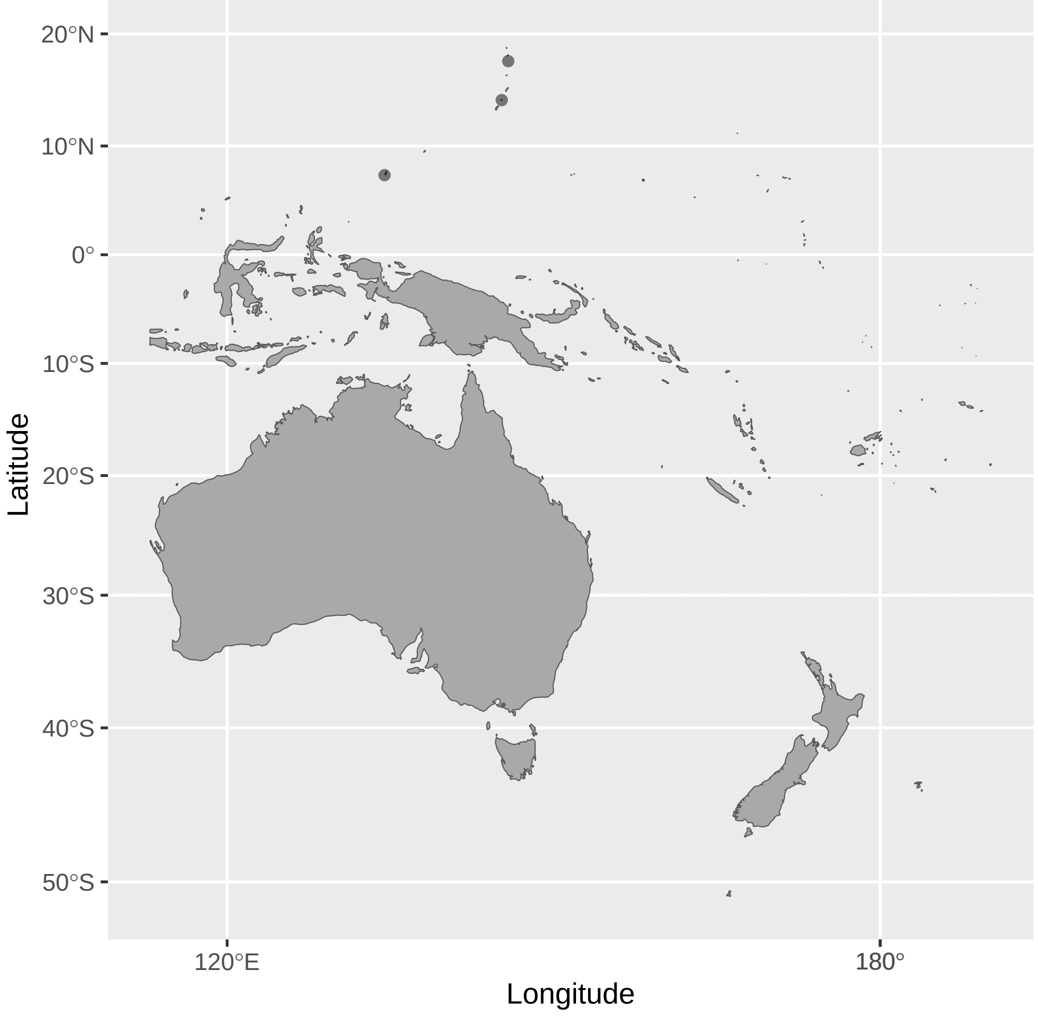

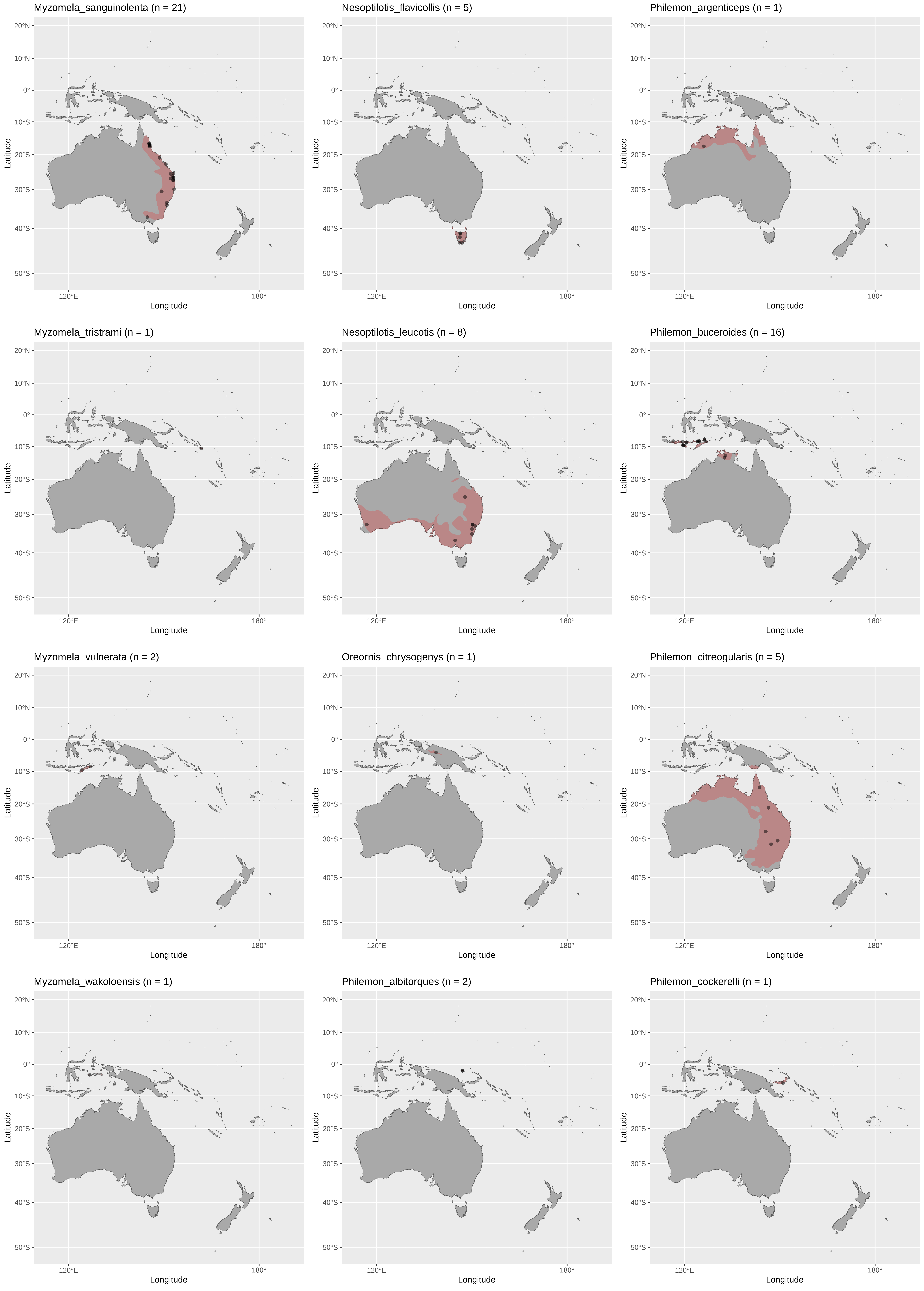
Myzomela_sanguinolenta (n = 21)
Nesoptilotis_flavicollis (n = 5)
Philemon_argenticeps (n = 1)
Myzomela_tristrami (n = 1)
Nesoptilotis_leucotis (n = 8)
Philemon_buceroides (n = 16)
Myzomela_vulnerata (n = 2)
Oreornis_chrysogenys (n = 1)
Philemon_citreogularis (n = 5)
Myzomela_wakoloensis (n = 1)
Philemon_albitorques (n = 2)
Philemon_cockerelli (n = 1)
Latitude
Longitude
20°N
10°N
0°
10°S
20°S
30°S
40°S
50°S
120°E
180°

Philemon_corniculatus (n = 11)

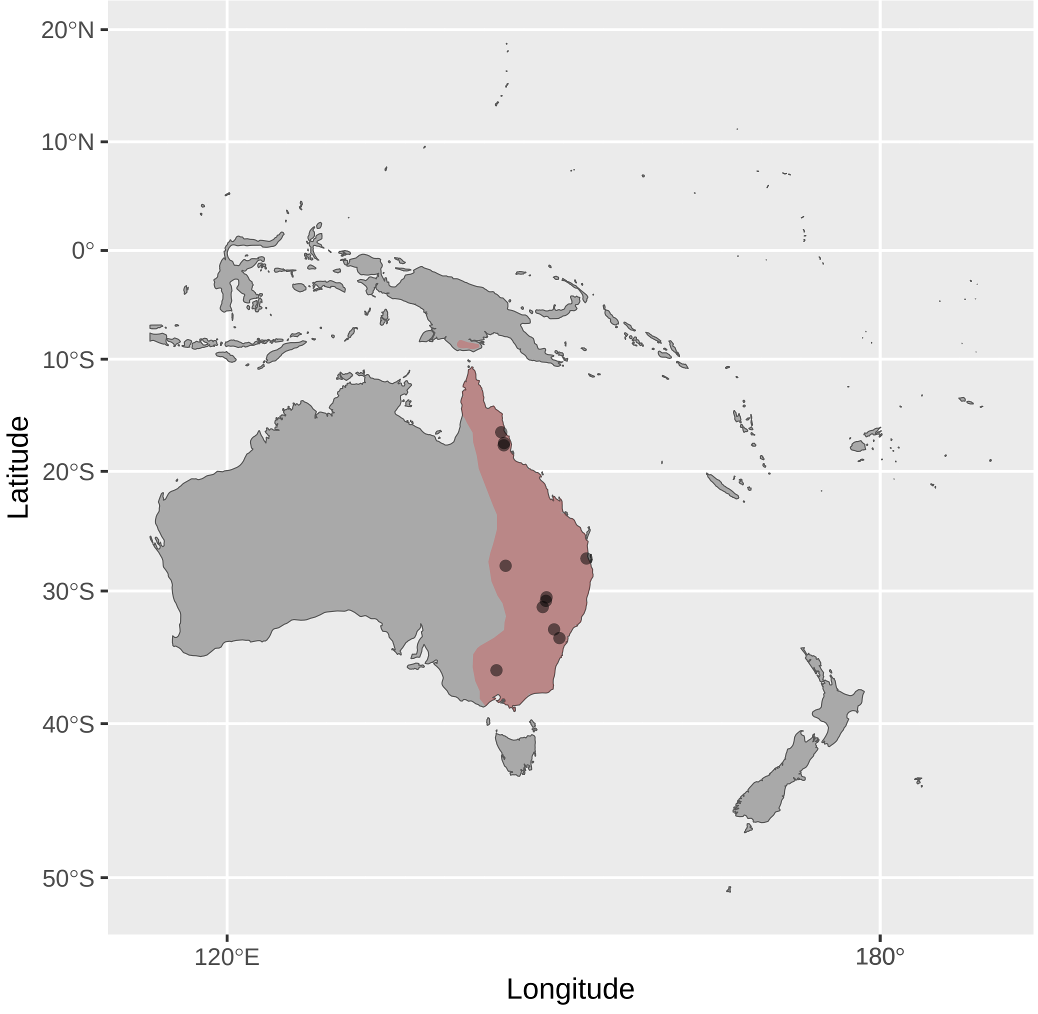


Philemon_kisserensis (n = 1)

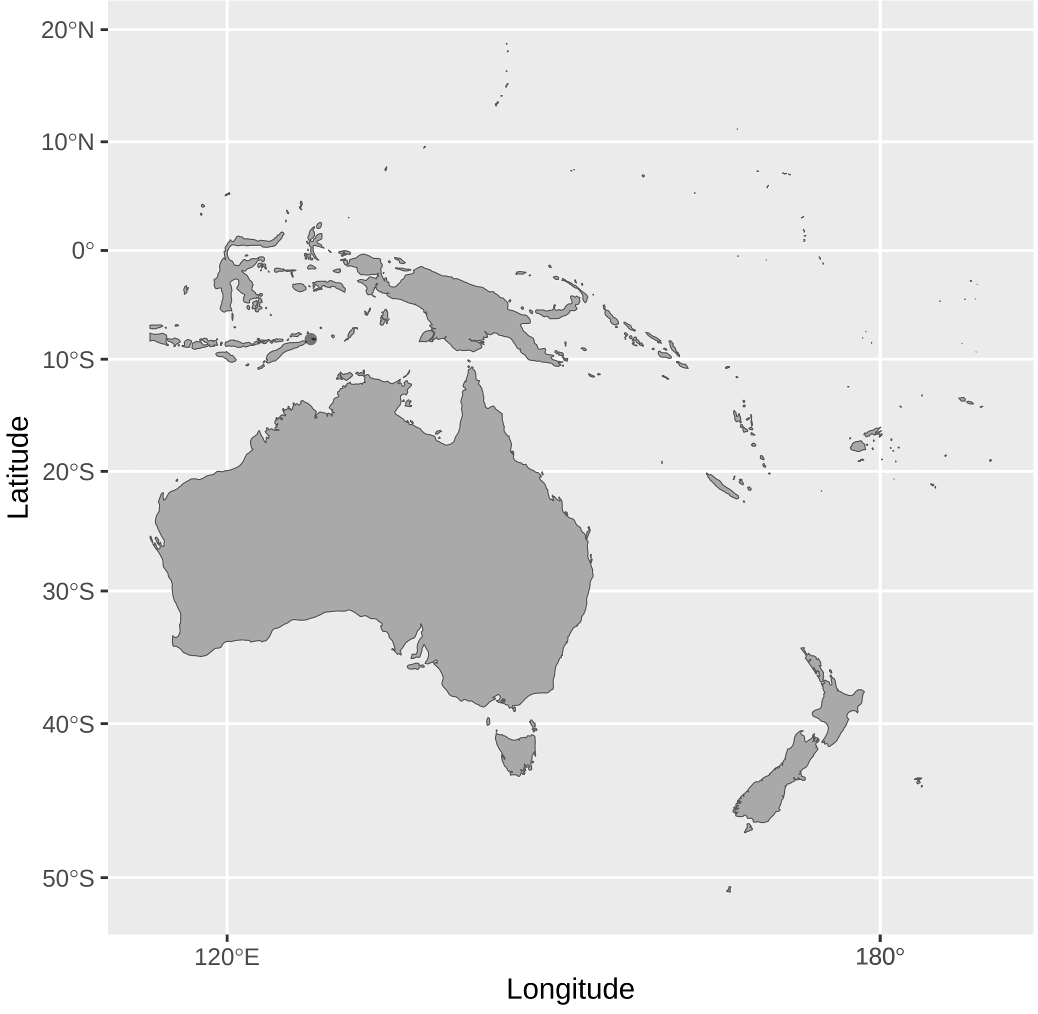


Philemon_plumigenis (n = 3)

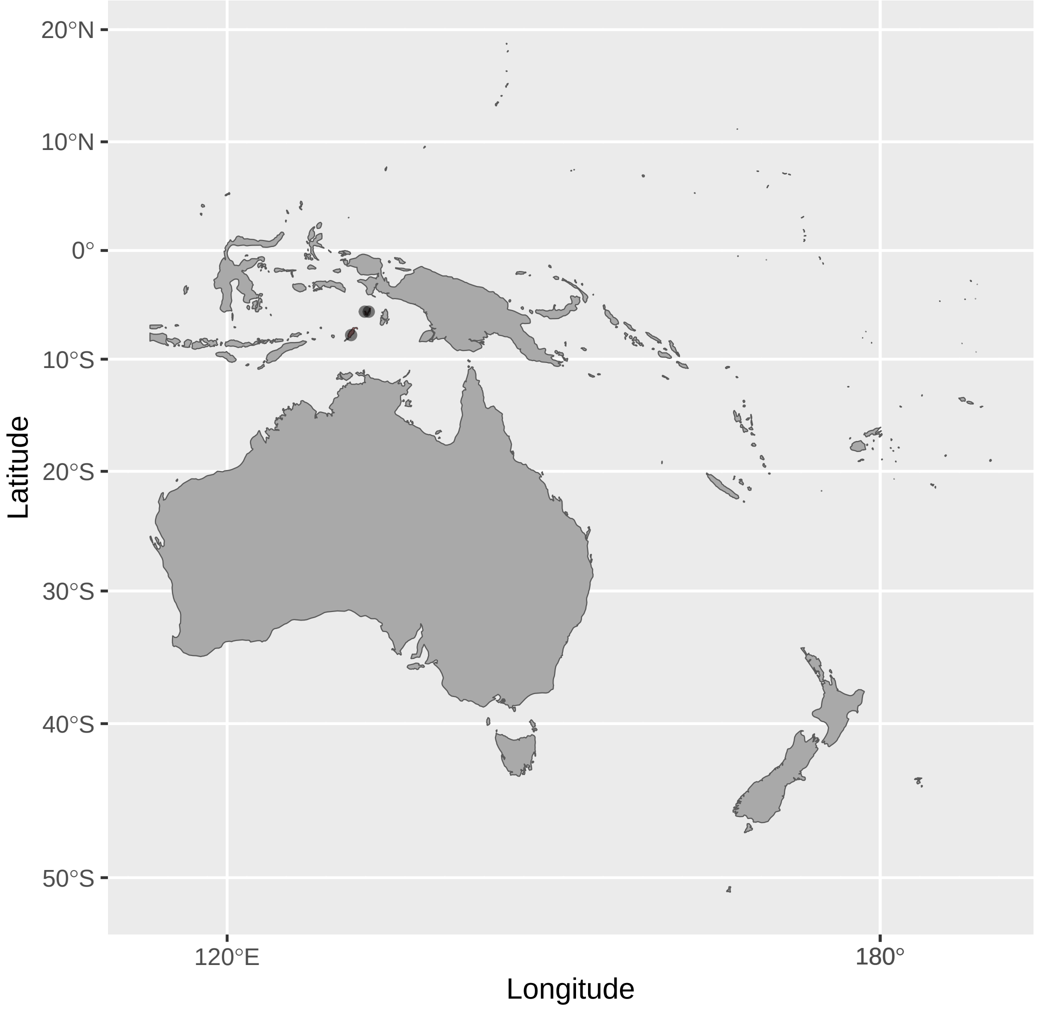


Philemon_diemenensis (n = 1)

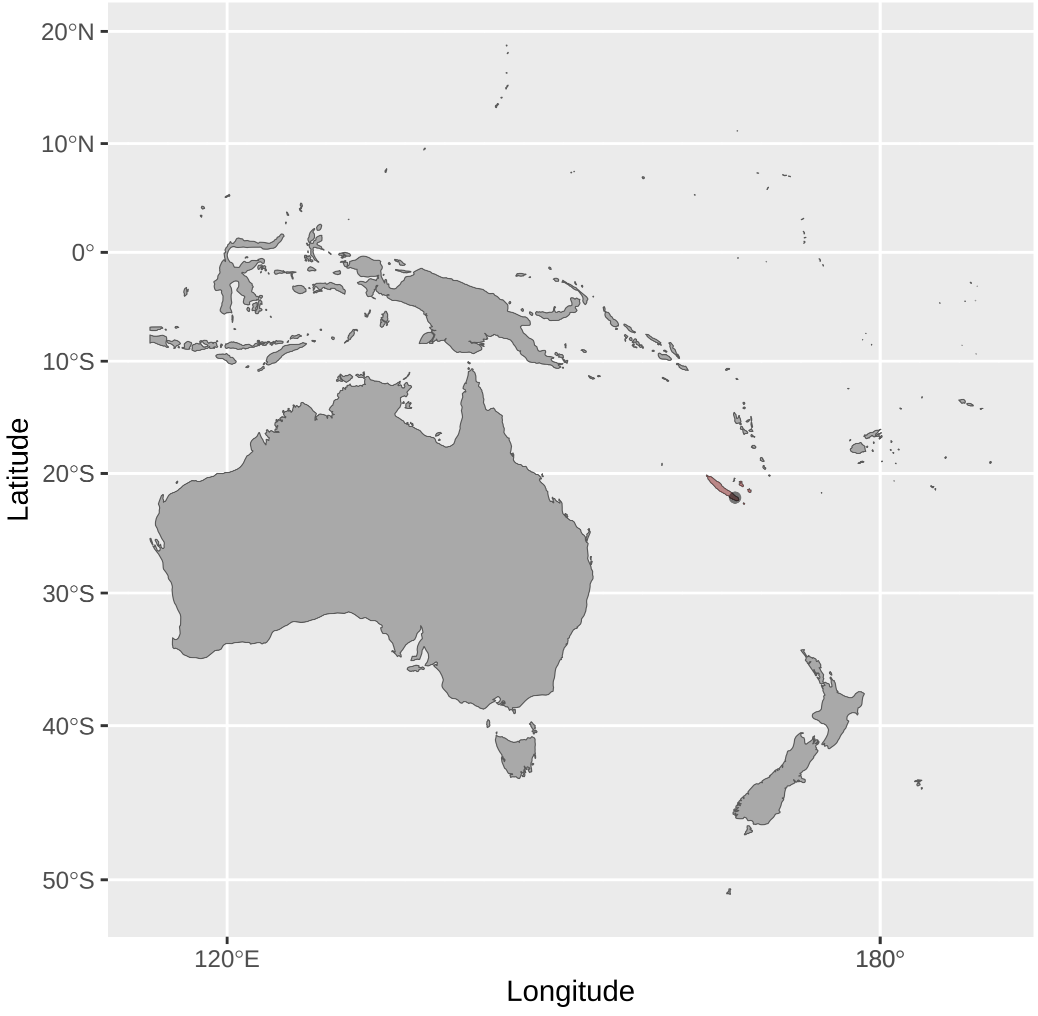


Philemon_meyeri (n = 6)

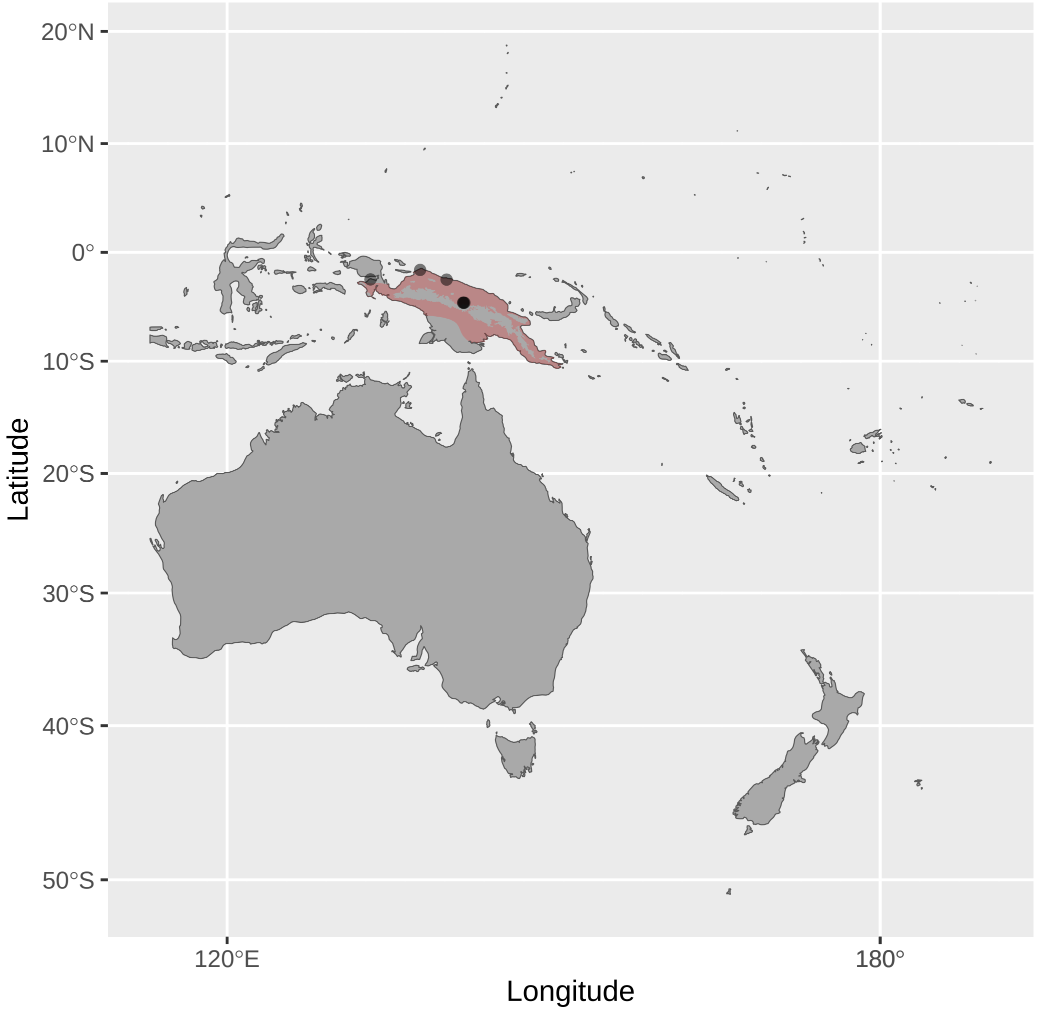


Philemon_subcorniculatus (n = 1)

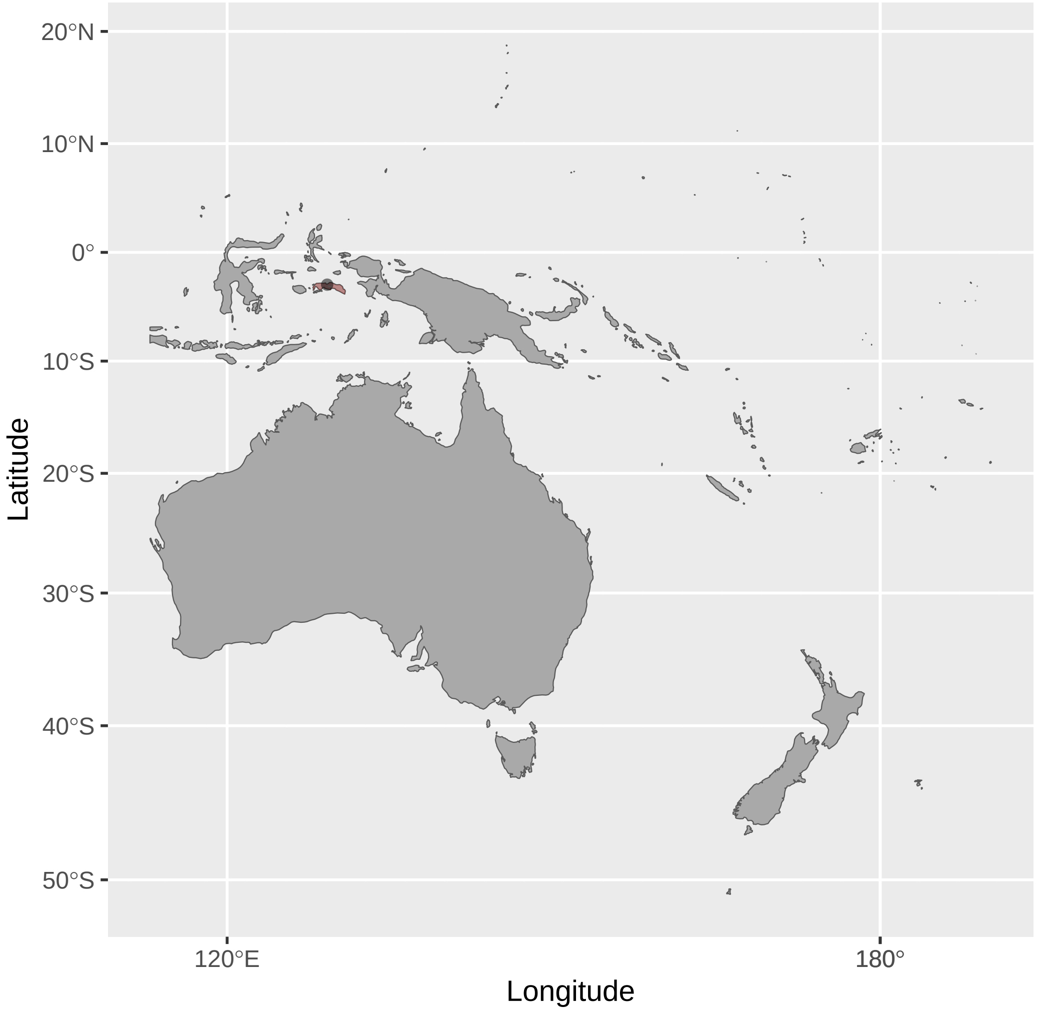


Philemon_fuscicapillus (n = 5)

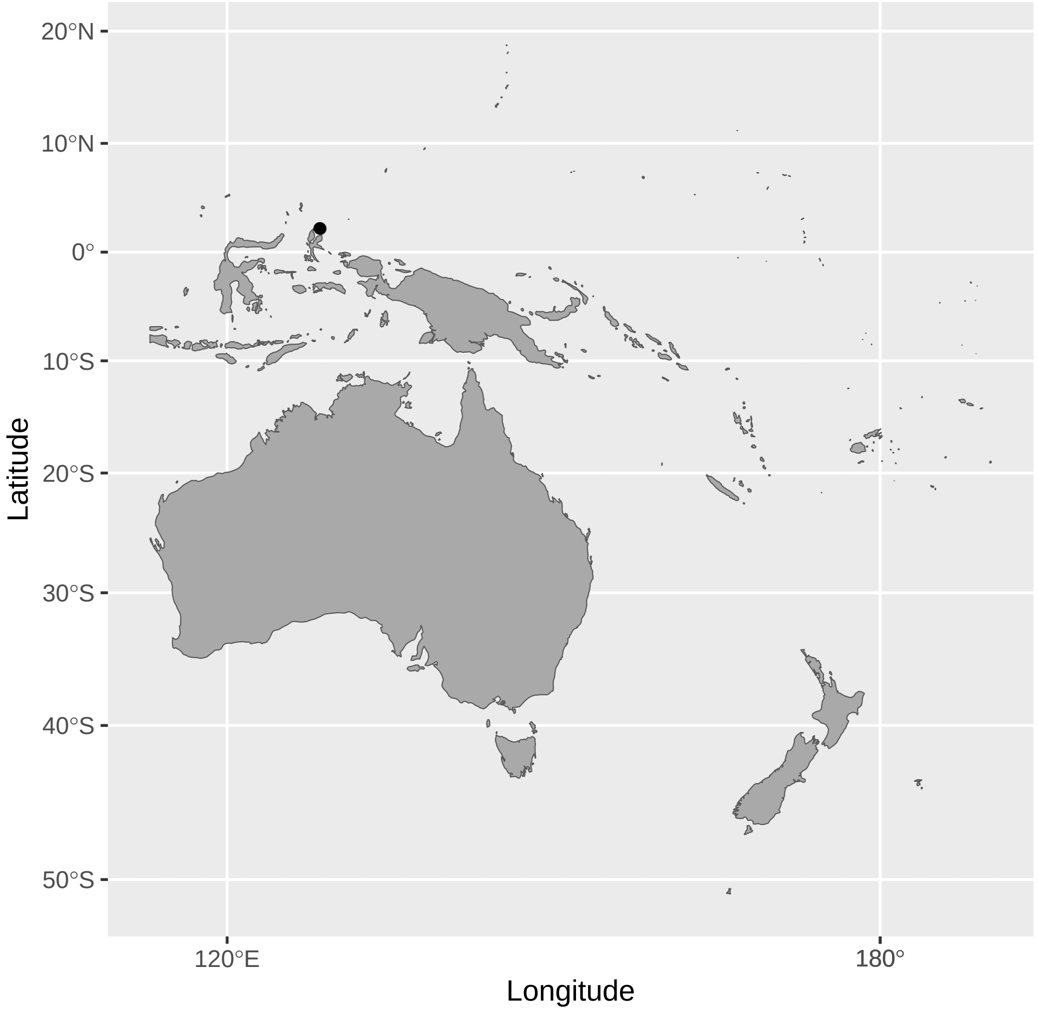


Philemon_moluccensis (n = 1)

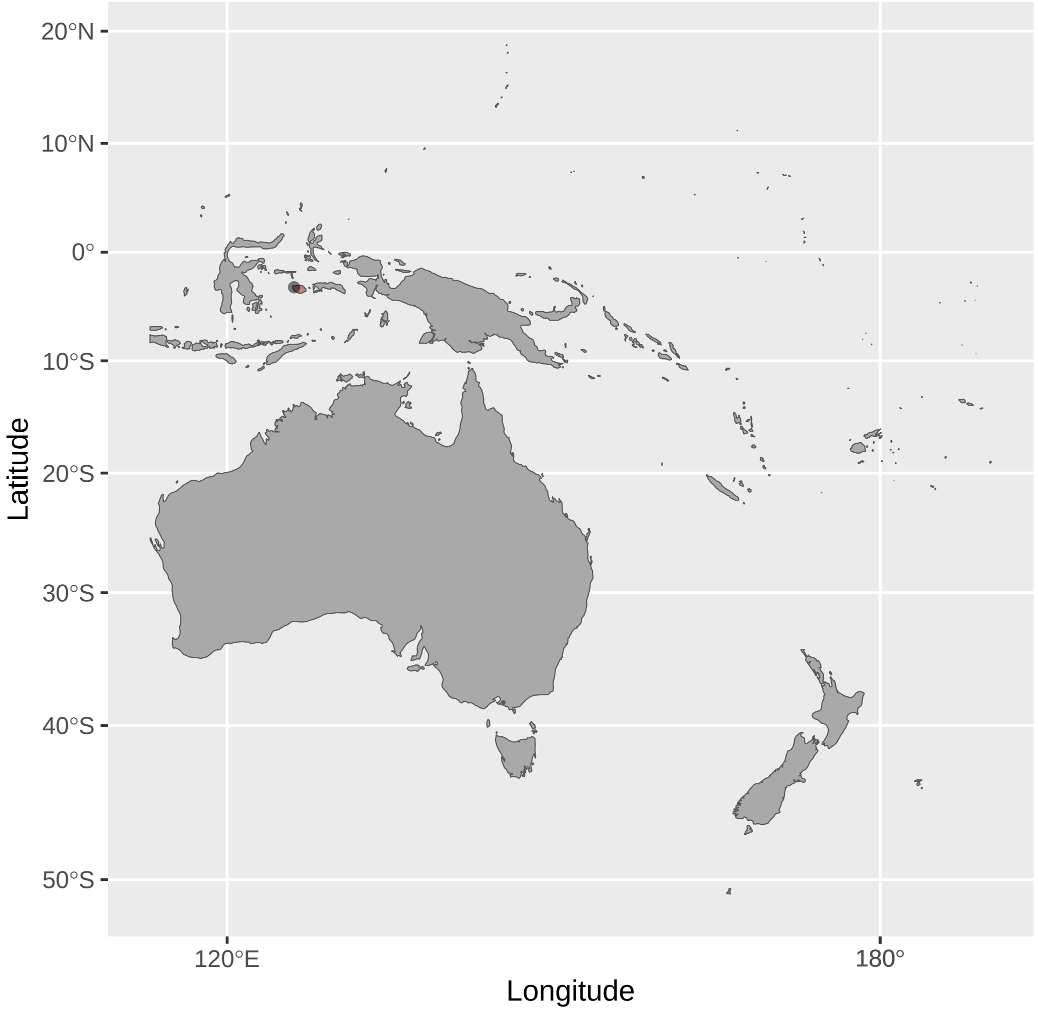


Philemon_yorki (n = 5)

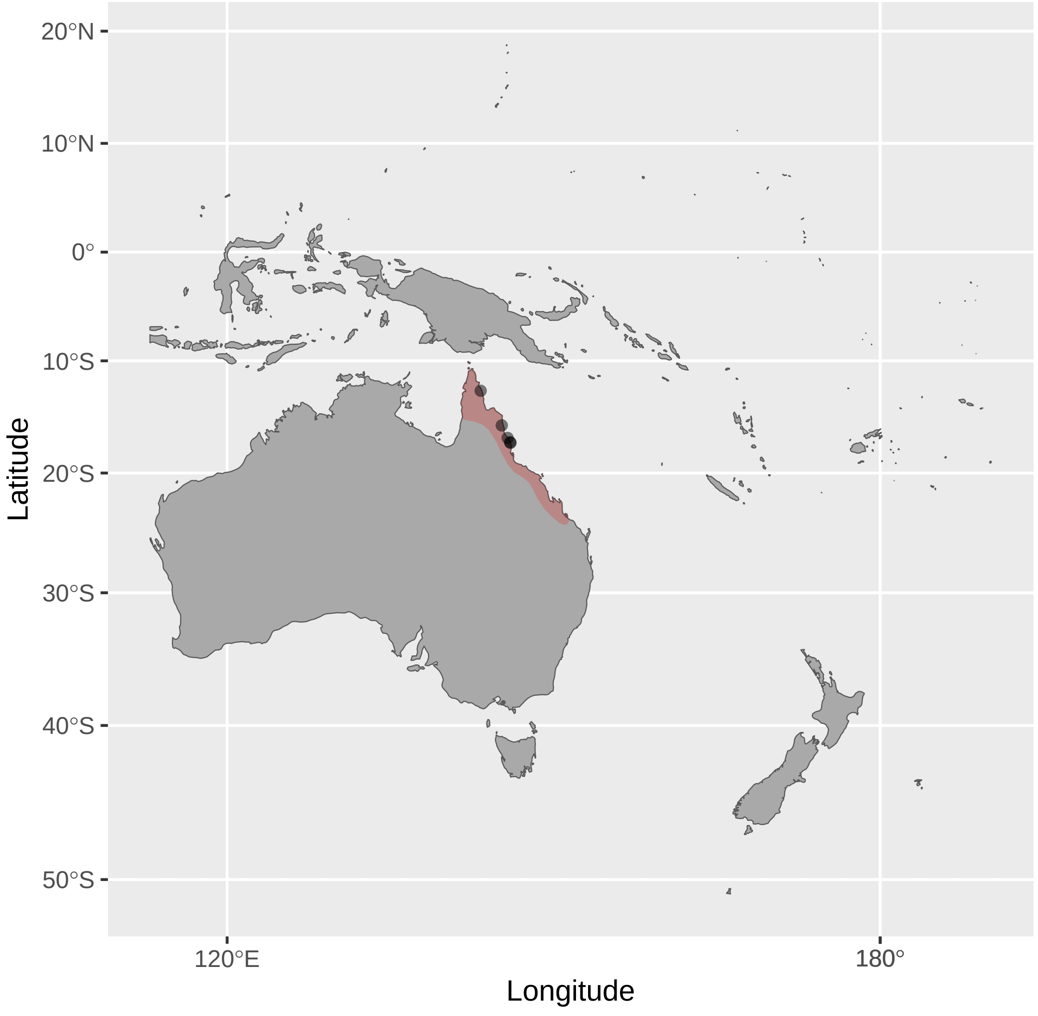


Philemon_inornatus (n = 3)

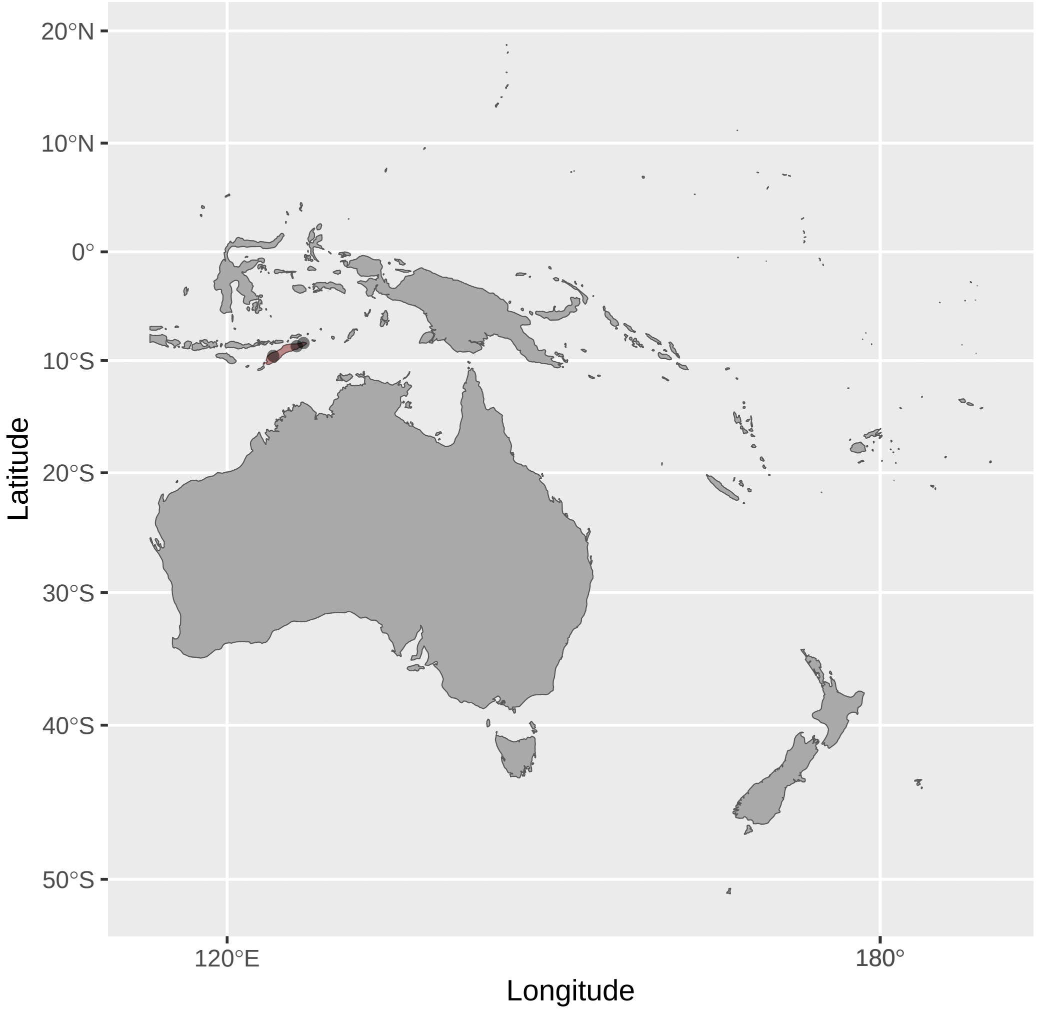


Philemon_novaeguineae (n = 12)

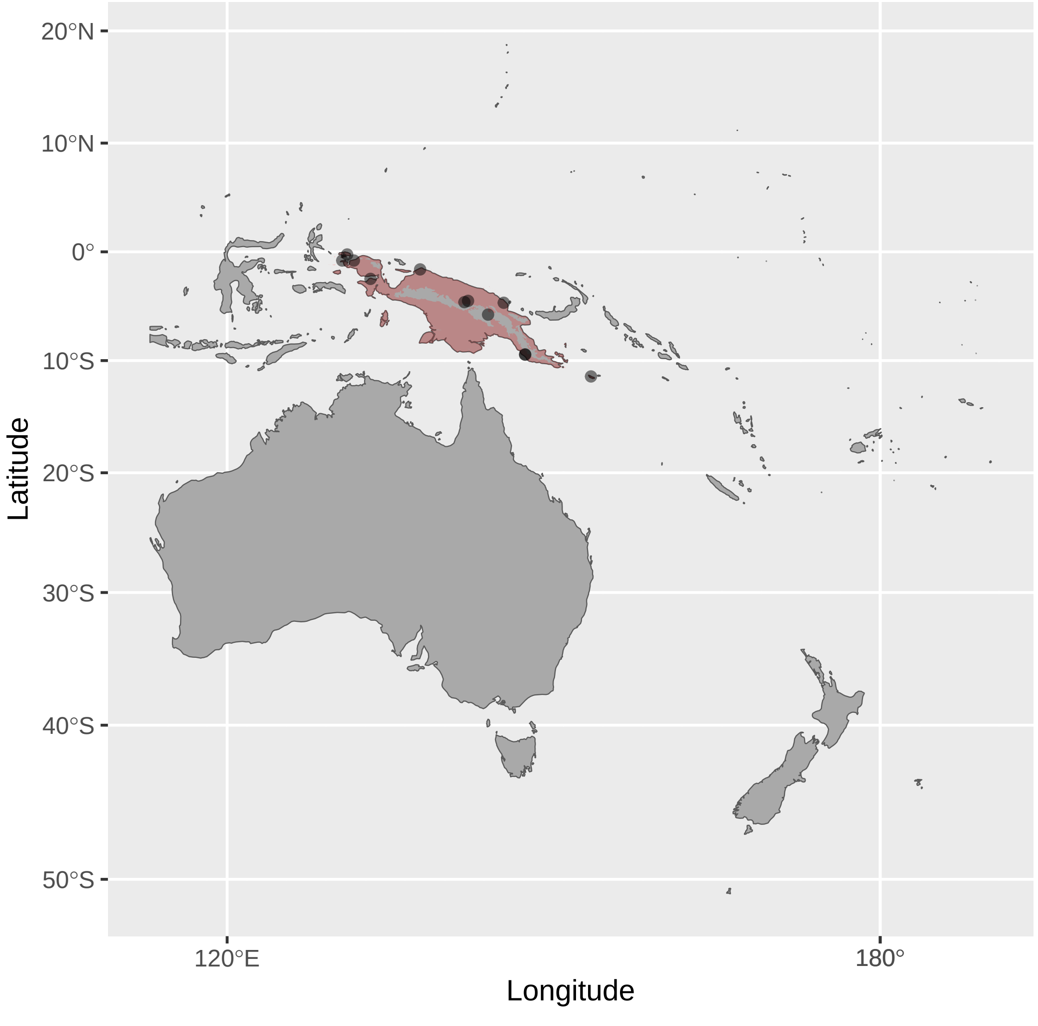


Phylidonyris_niger (n = 2)

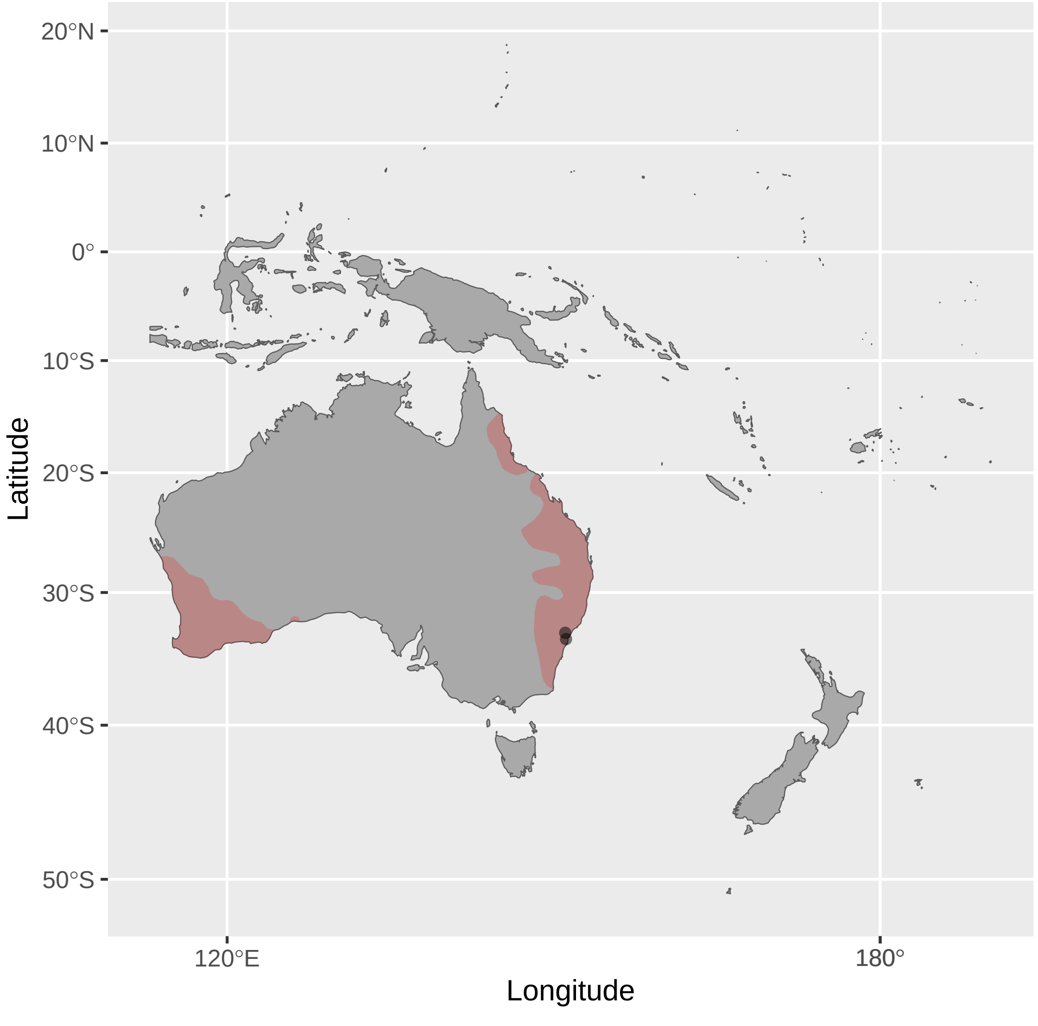

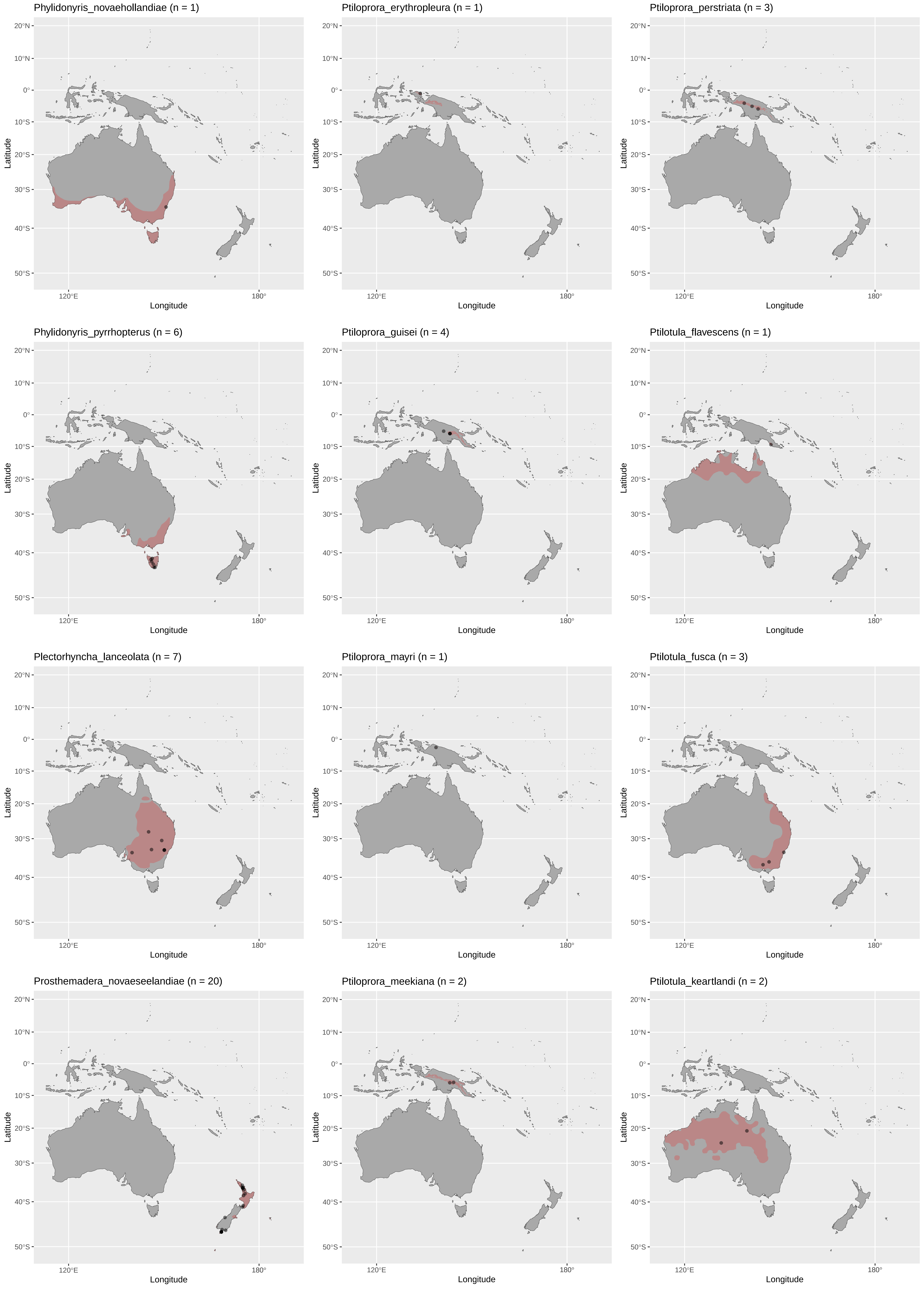
Phylidonyris_novaehollandiae (n = 1)
Ptiloprora_erythropleura (n = 1)
Ptiloprora_perstriata (n = 3)
Phylidonyris_pyrrhopterus (n = 6)
Ptiloprora_guisei (n = 4)
Ptilotula_flavescens (n = 1)
Plectorhyncha_lanceolata (n = 7)
Ptiloprora_mayri (n = 1)
Ptilotula_fusca (n = 3)
Prosthemadera_novaeseelandiae (n = 20)
Ptiloprora_meekiana (n = 2)
Ptilotula_keartlandi (n = 2)
Latitude
Longitude
20°N
10°N
0°
10°S
20°S
30°S
40°S
50°S
120°E
180°

Ptilotula_ornata (n = 2)

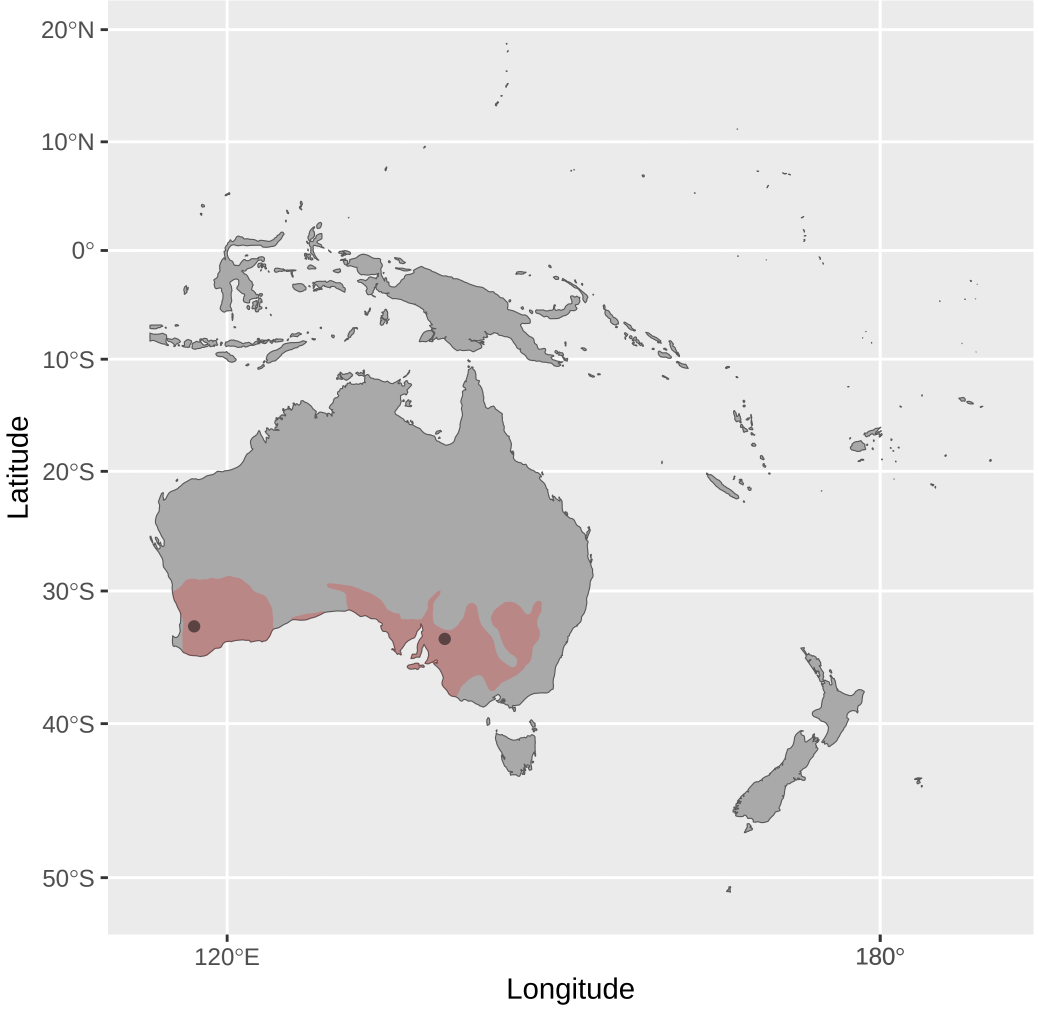


Pycnopygius_cinereus (n = 1)

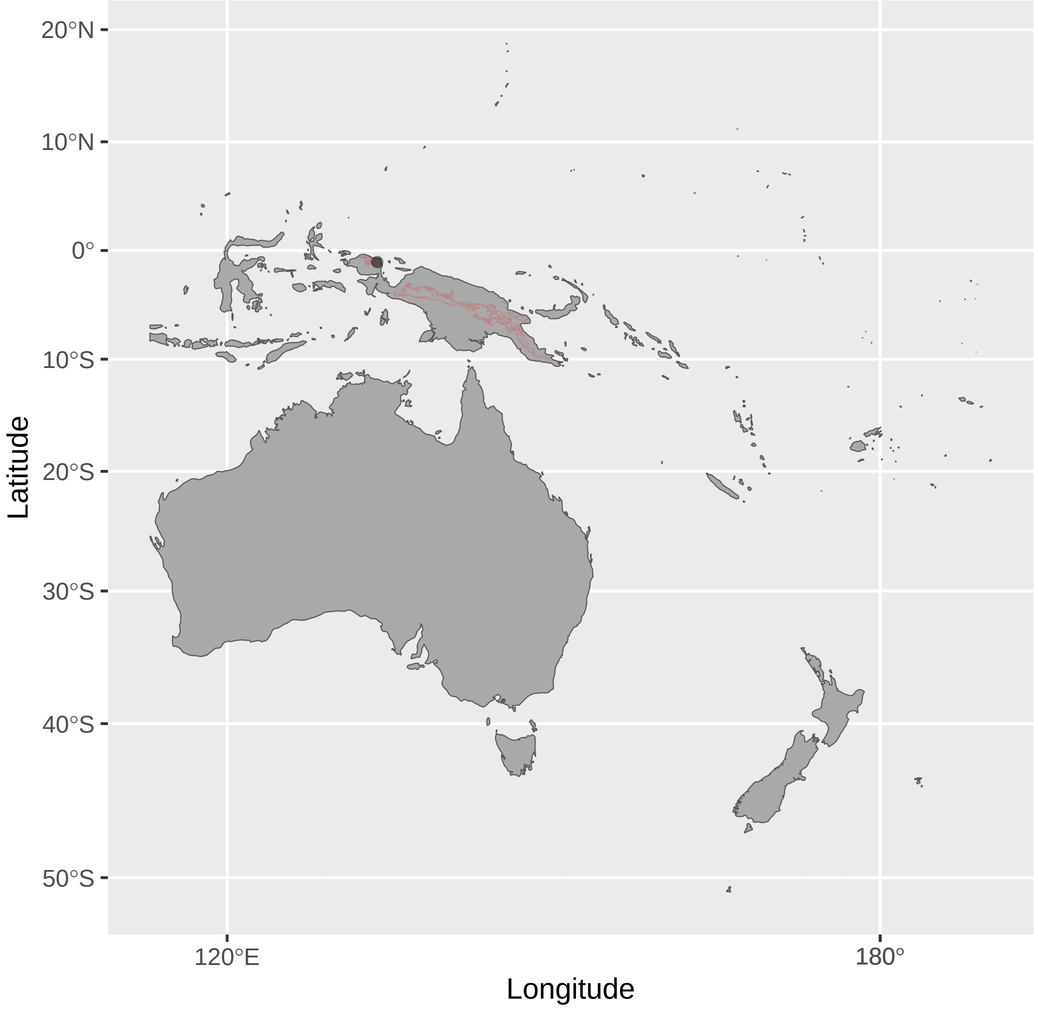


Ramsayornis_modestus (n = 4)

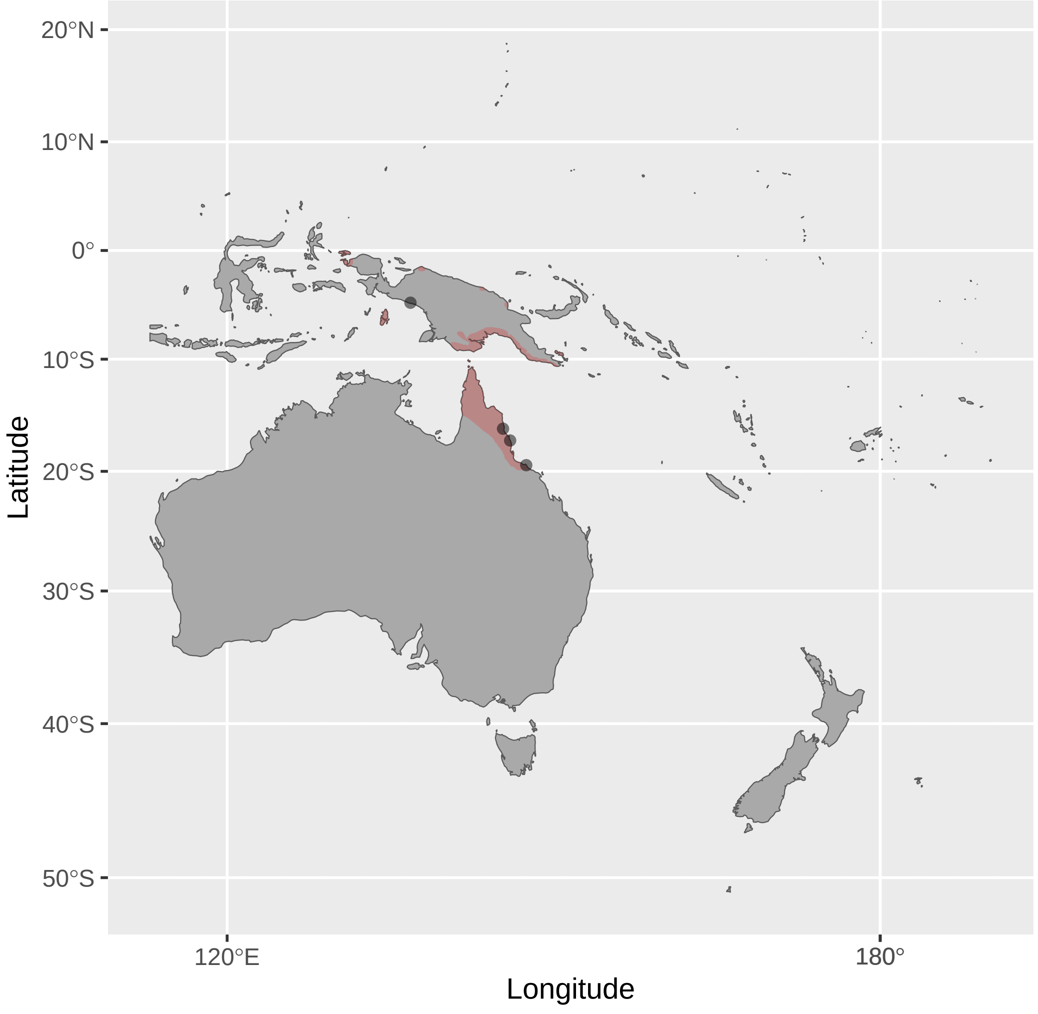


Ptilotula_penicillata (n = 4)

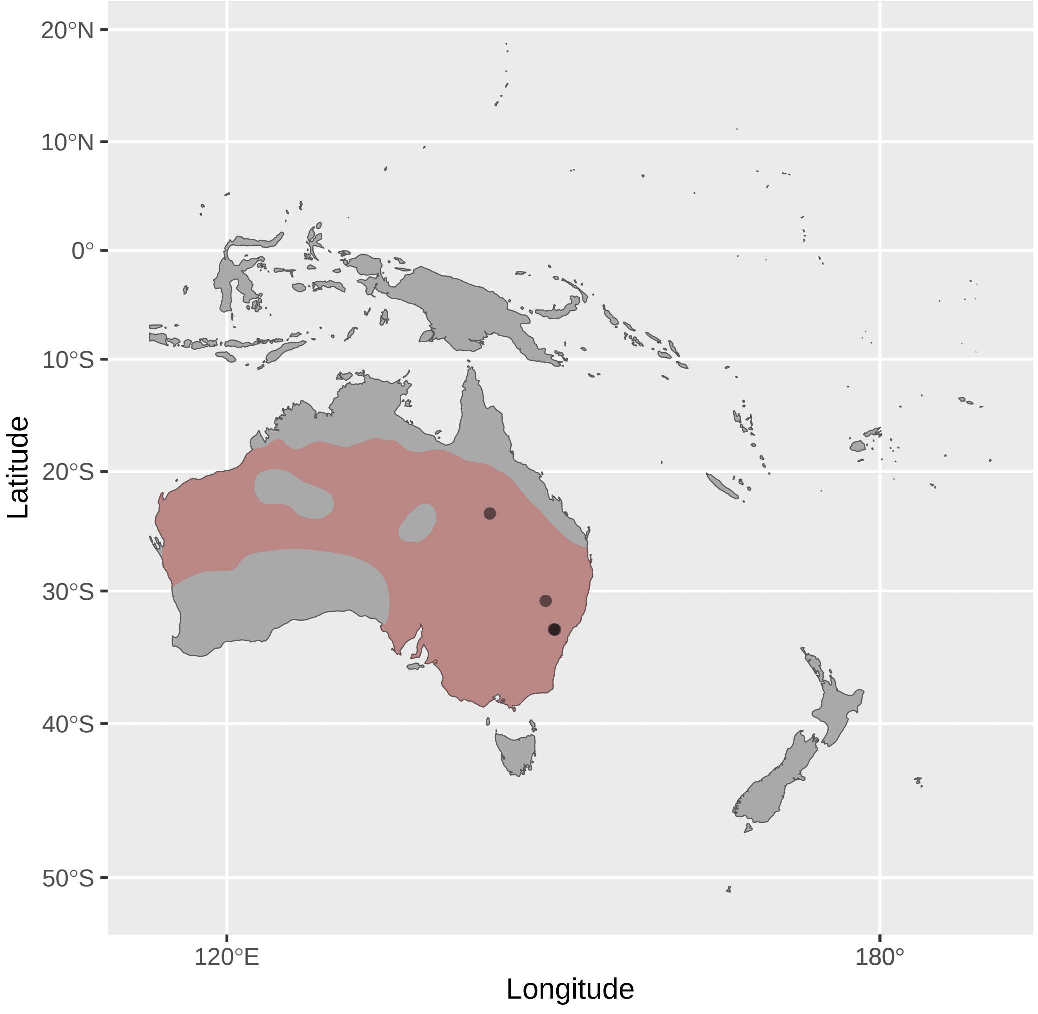


Pycnopygius_ixoides (n = 1)

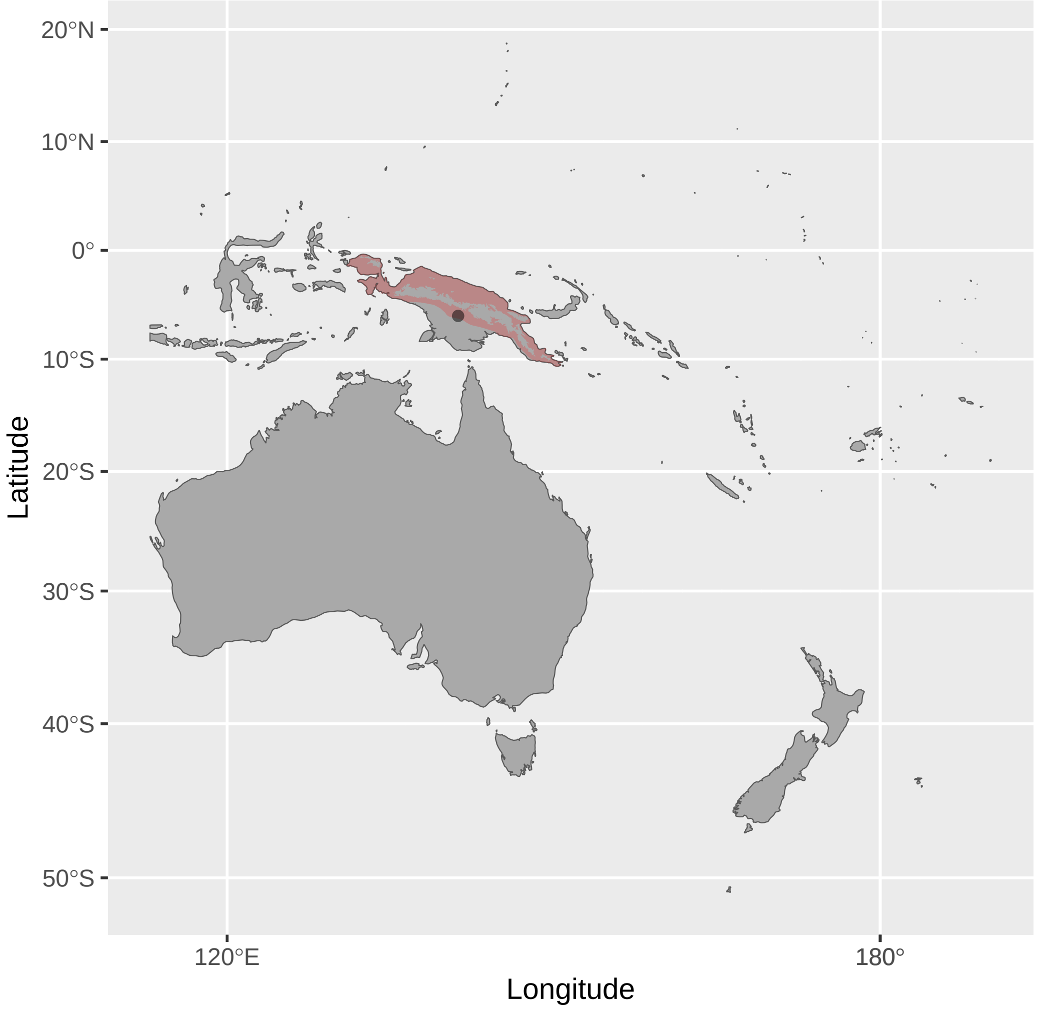


Stomiopera_flava (n = 5)

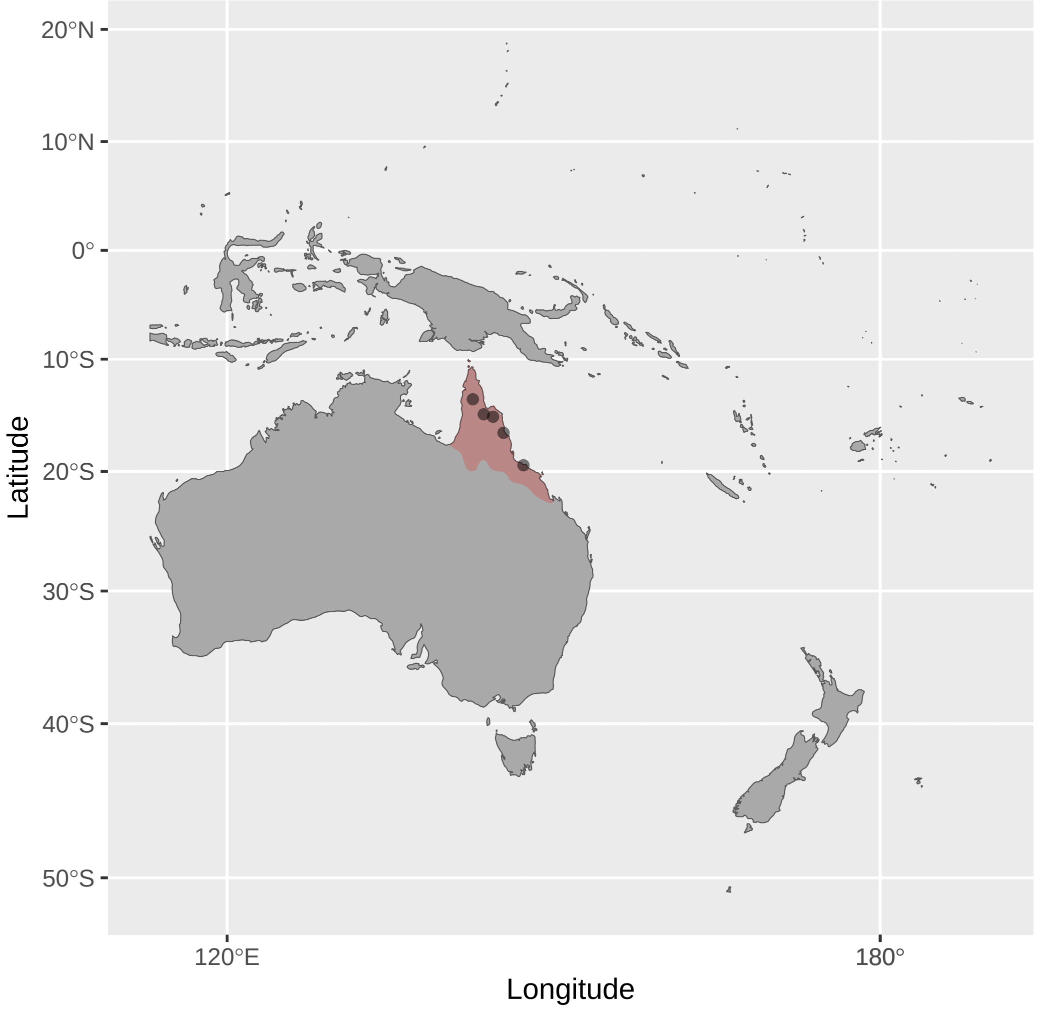


Ptilotula_plumula (n = 1)

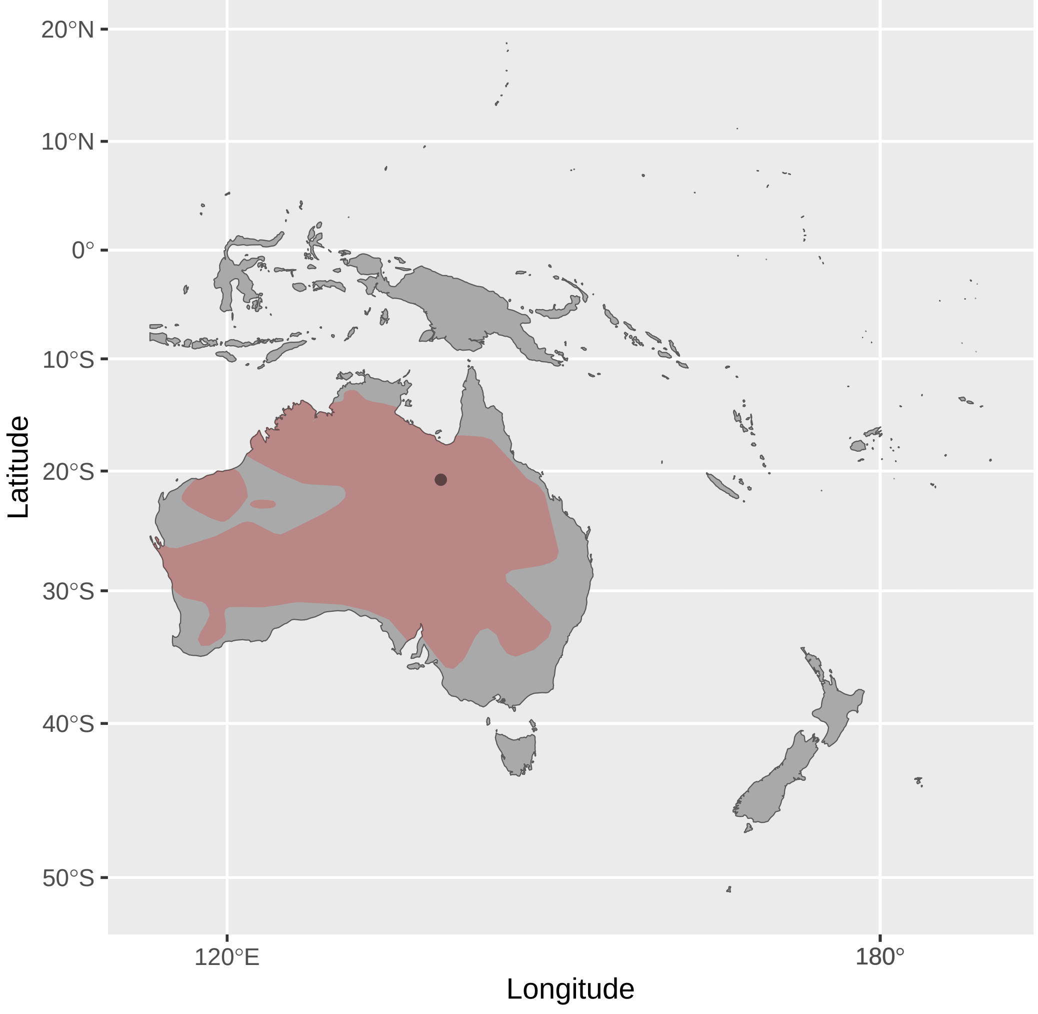


Pycnopygius_stictocephalus (n = 6)

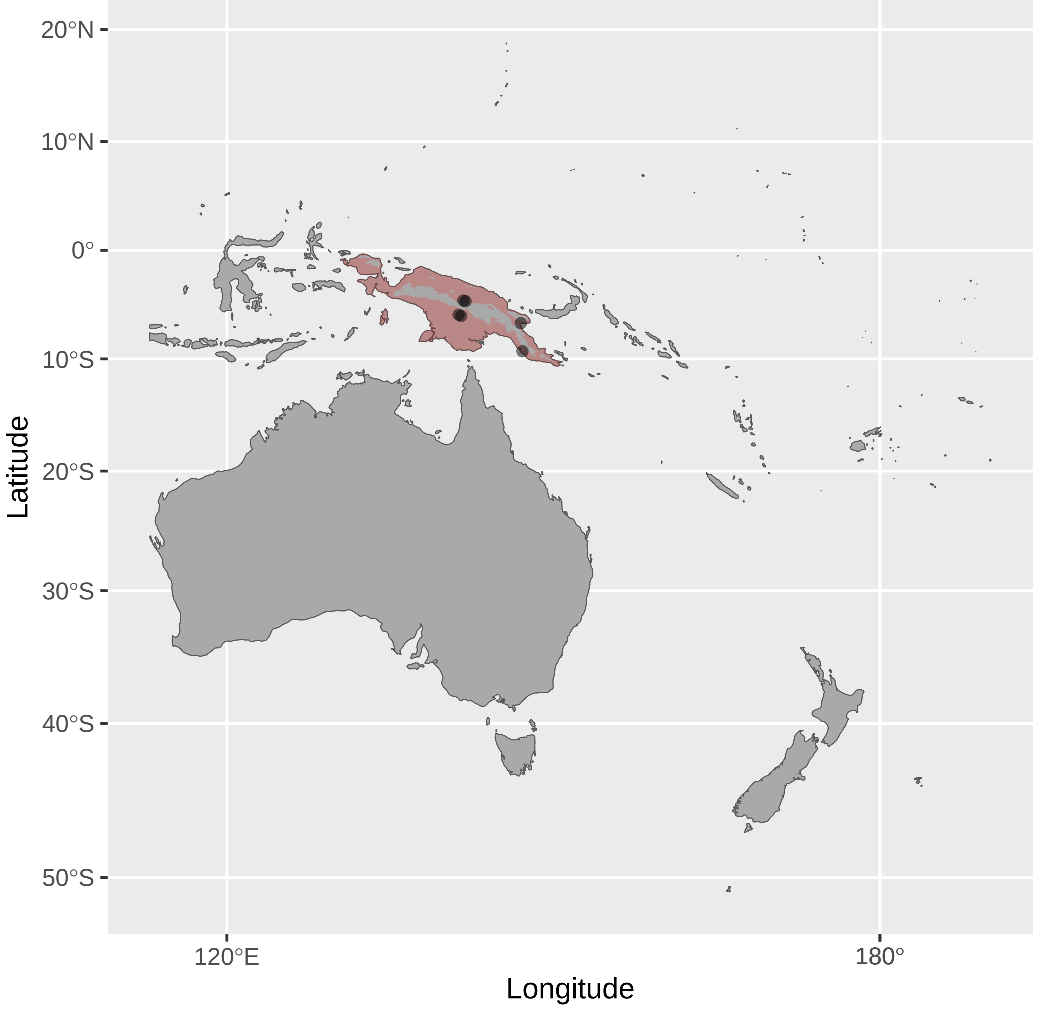


Stomiopera_unicolor (n = 3)

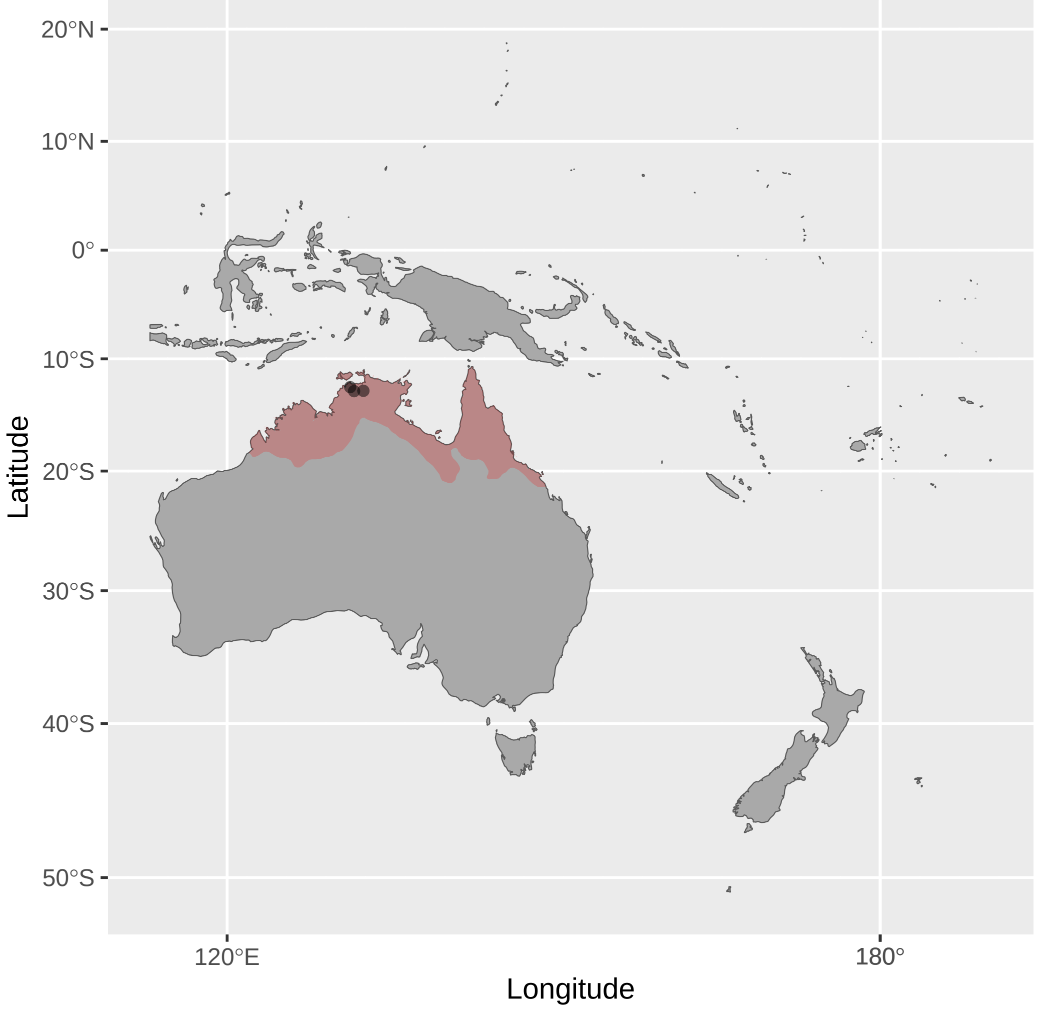


Purnella_albifrons (n = 5)

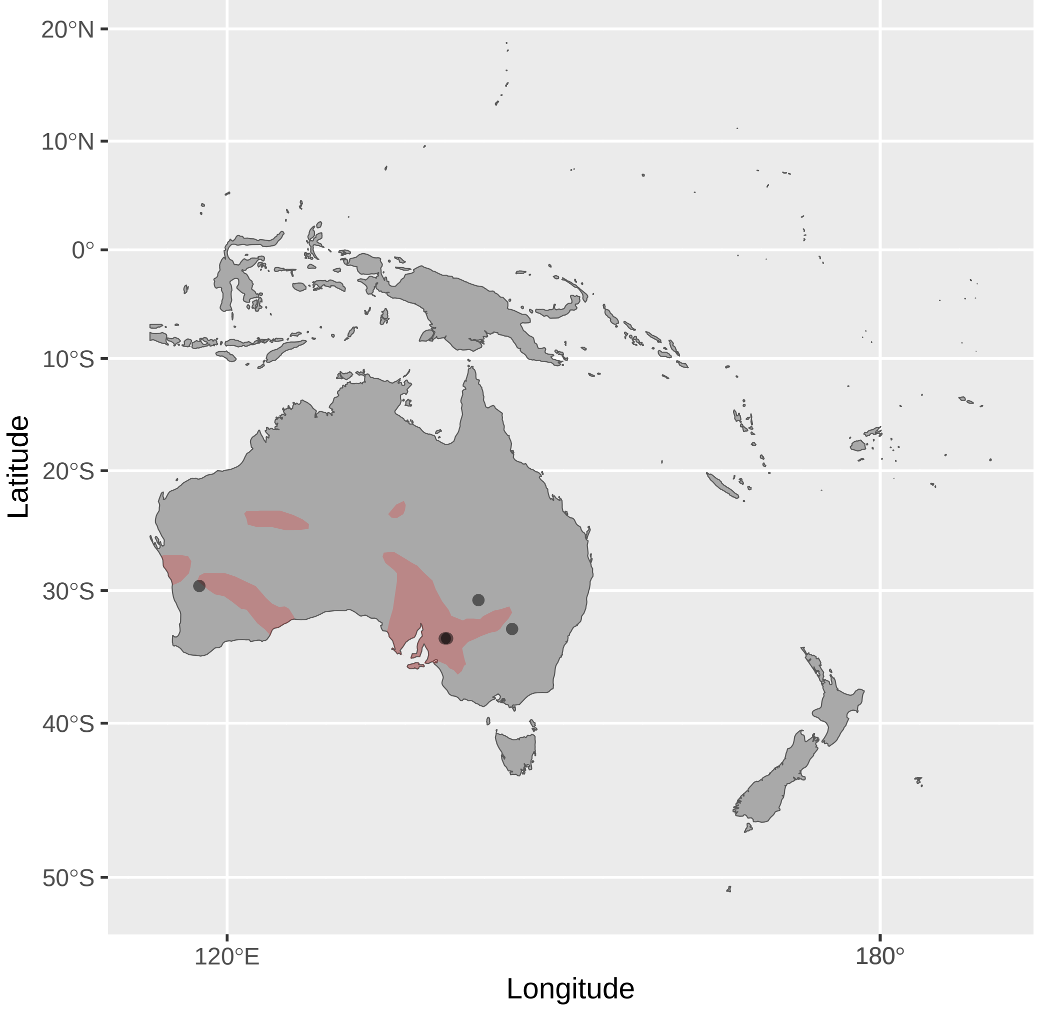


Ramsayornis_fasciatus (n = 1)

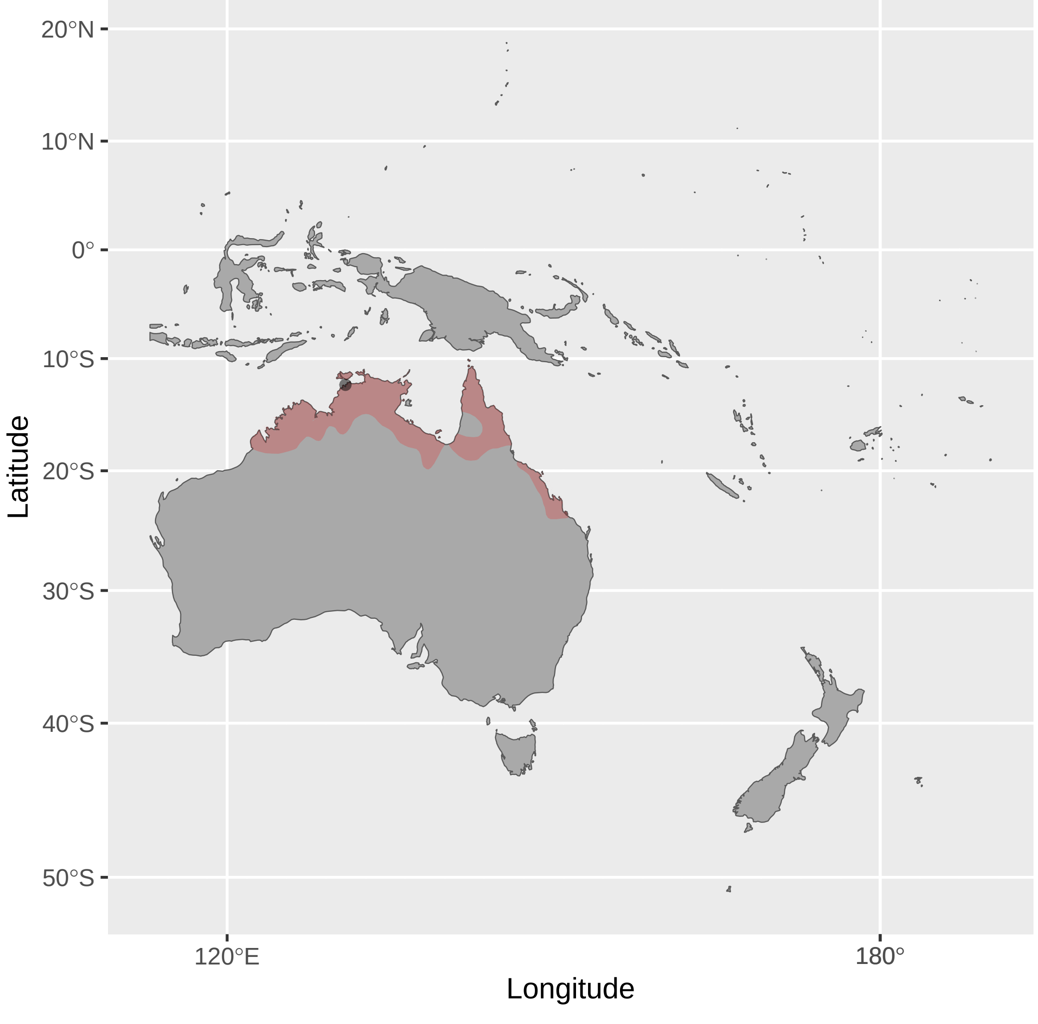


Stresemannia_bougainvillei (n = 1)

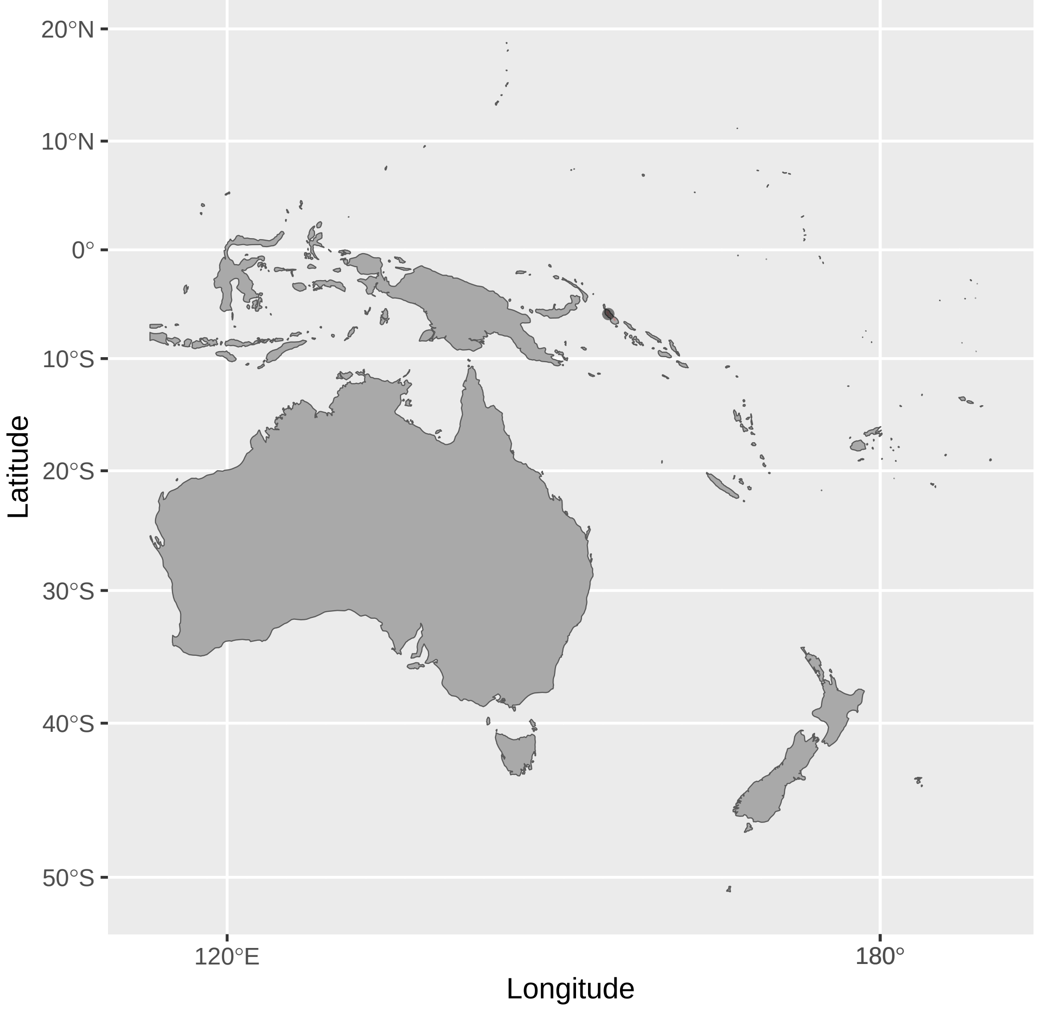

Sugomel_niger (n = 3)

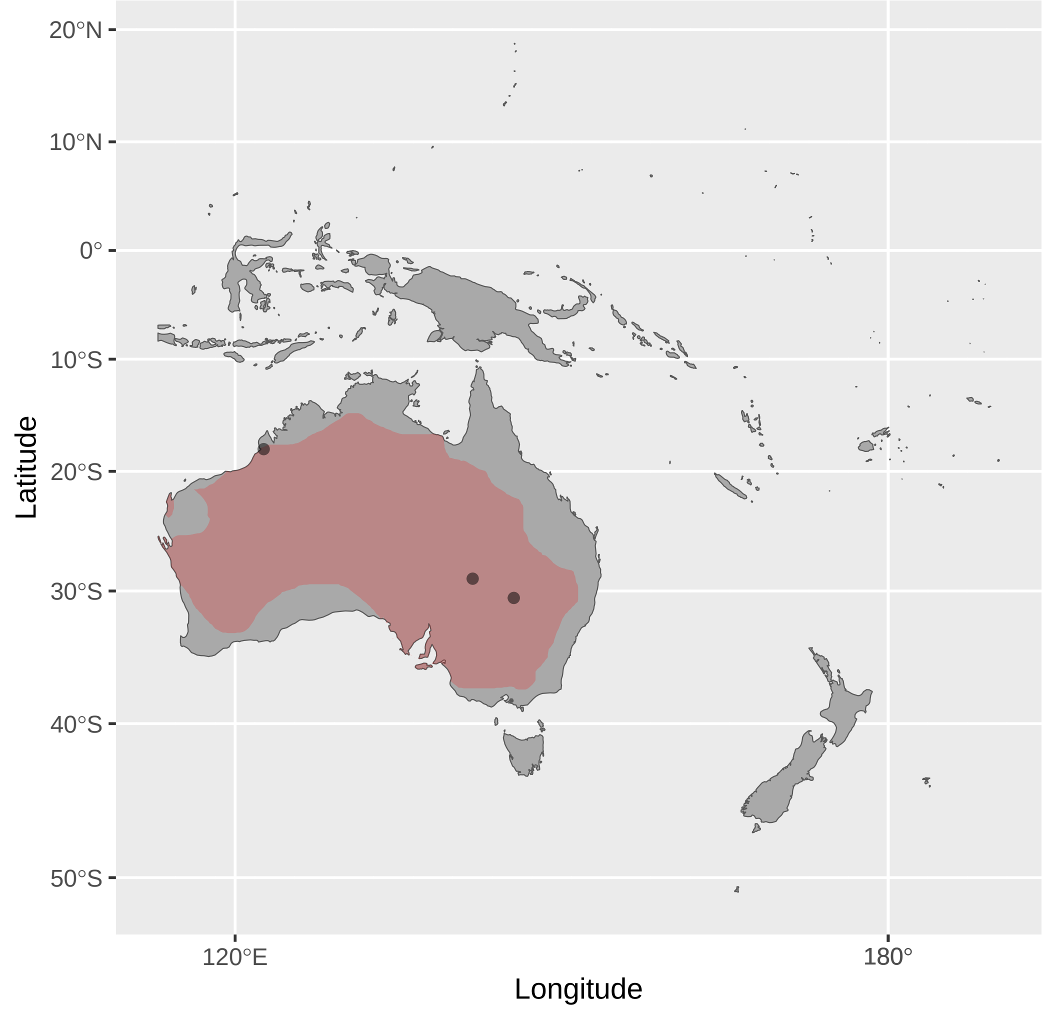


Trichodere_cockerelli (n = 1)

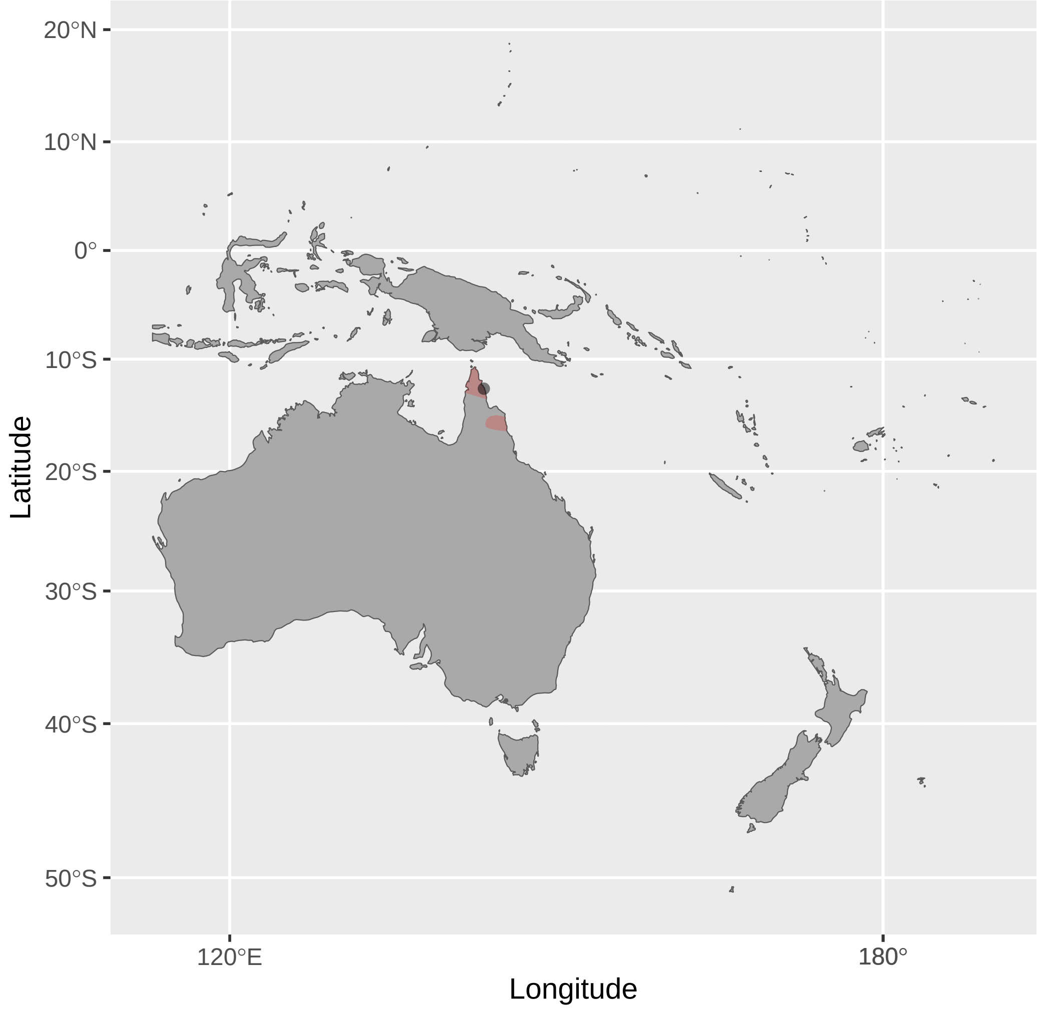


Territornis_albilineata (n = 2)

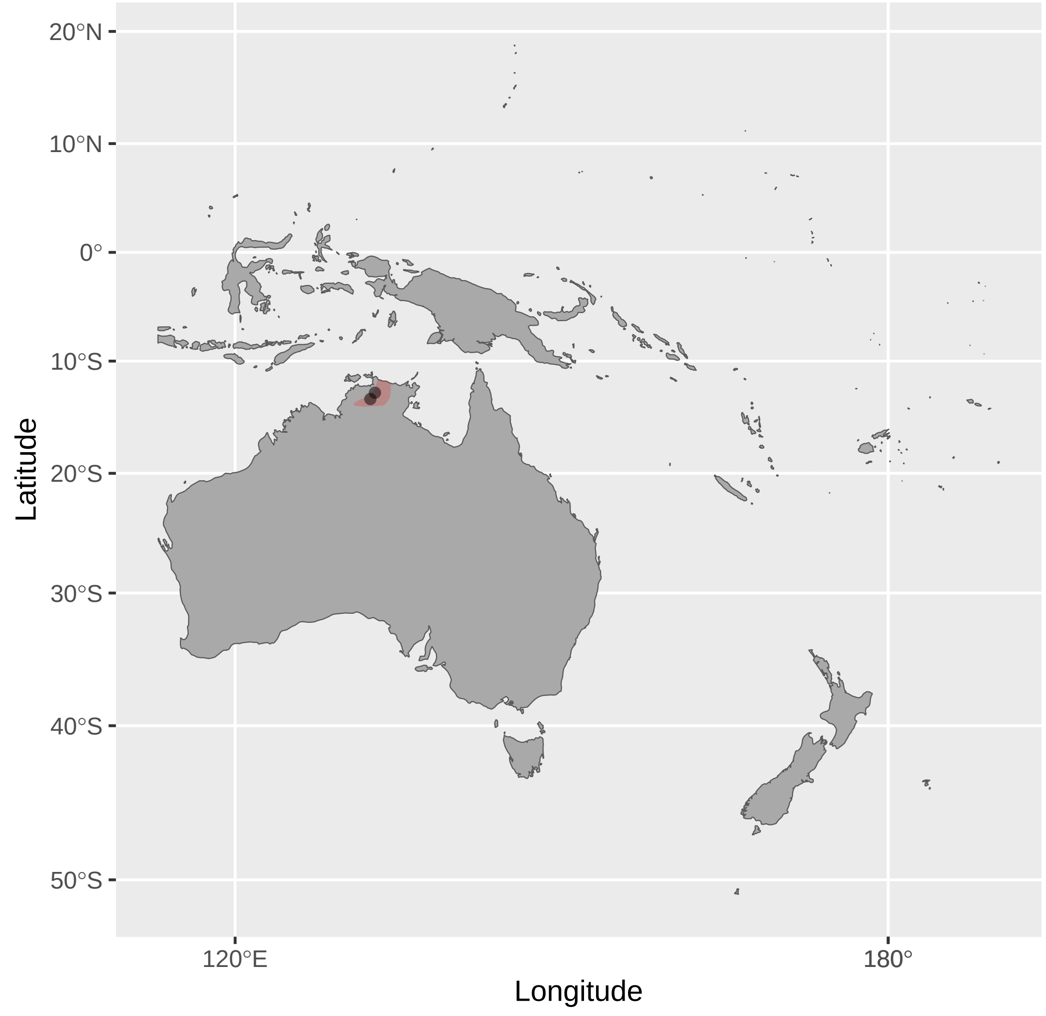


Xanthotis_flaviventer (n = 4)

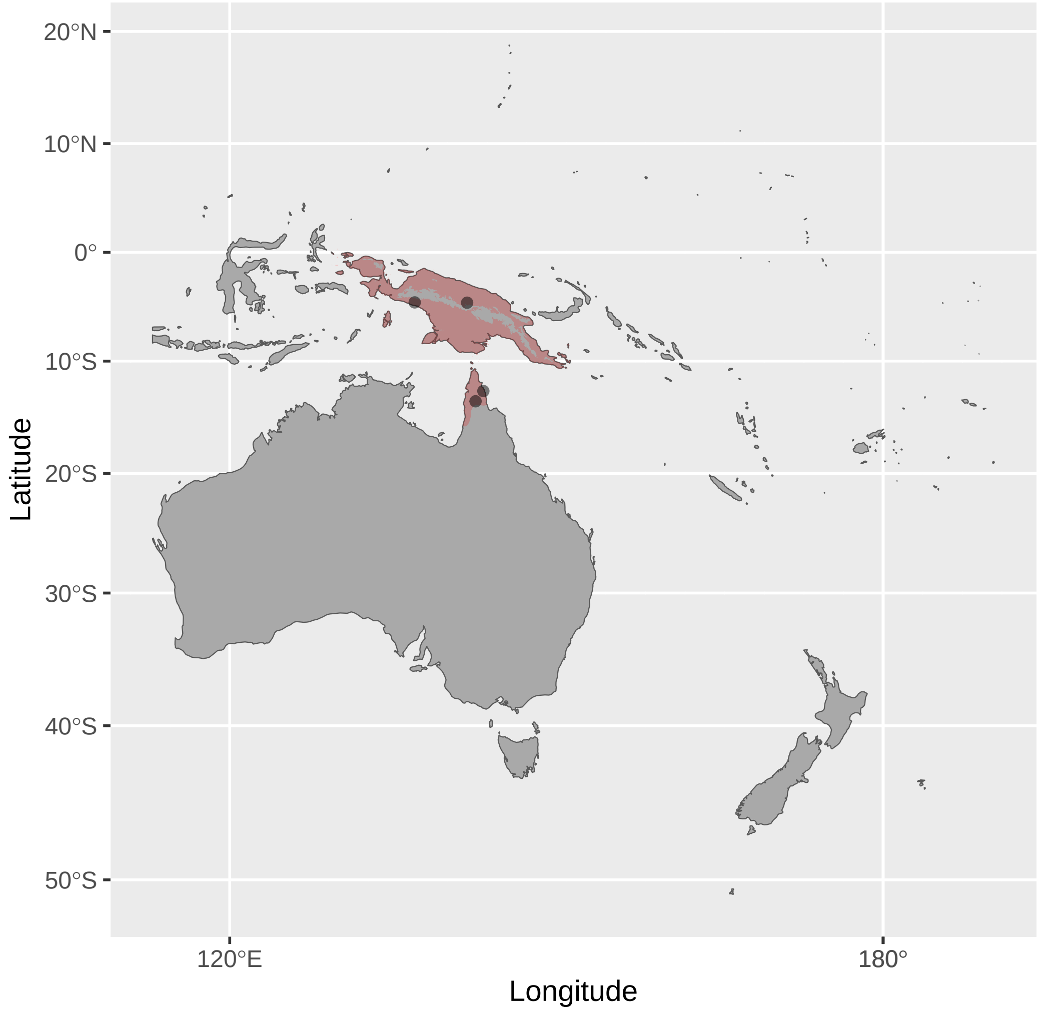


Territornis_fordiana (n = 1)

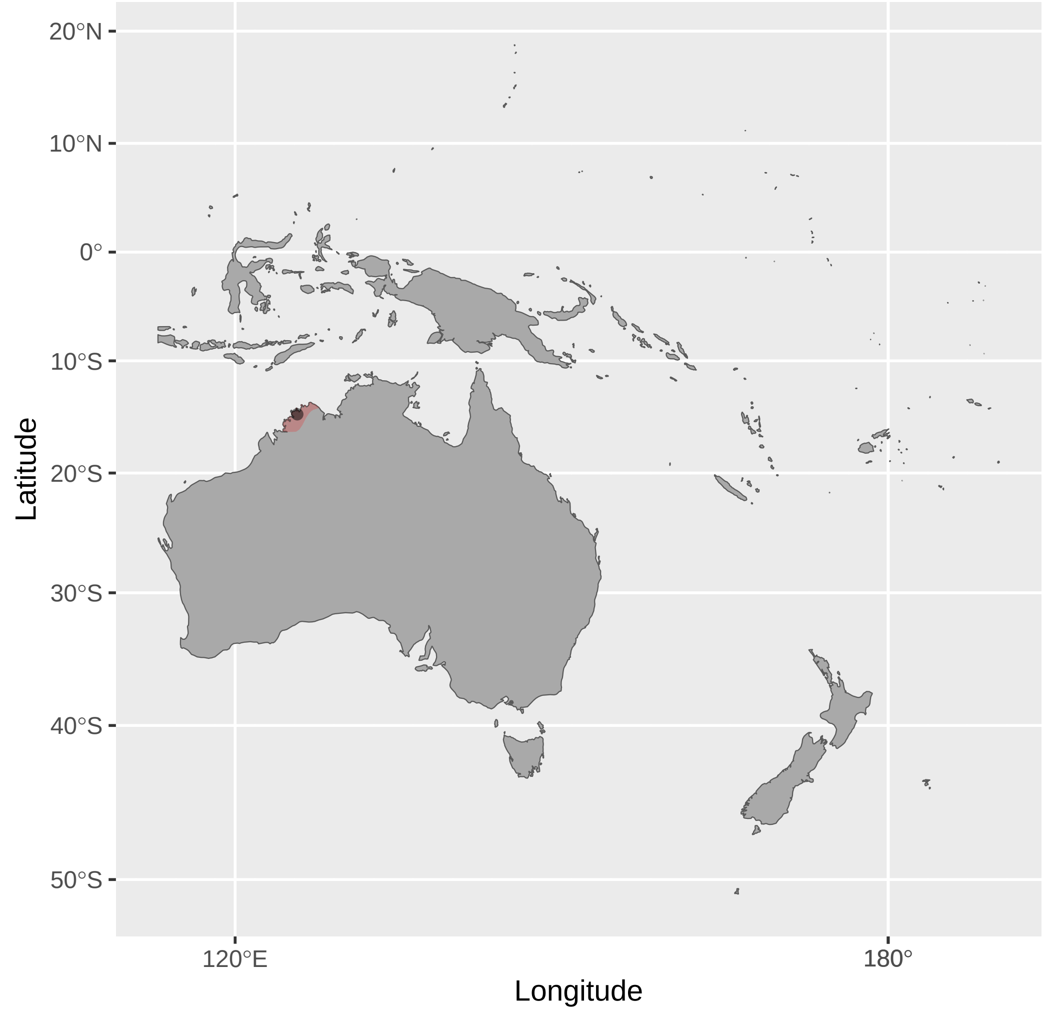


Xanthotis_macleayanus (n = 8)

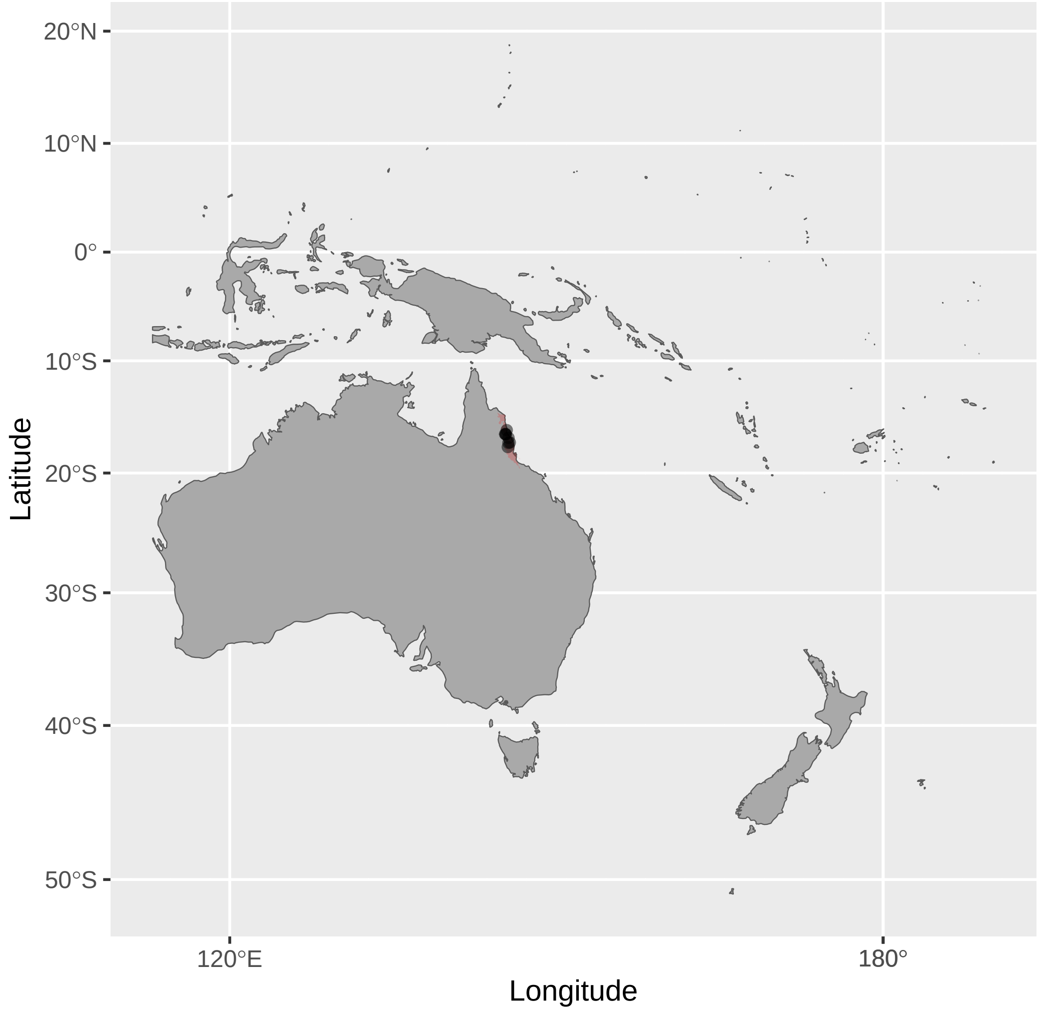


Territornis_reticulata (n = 3)

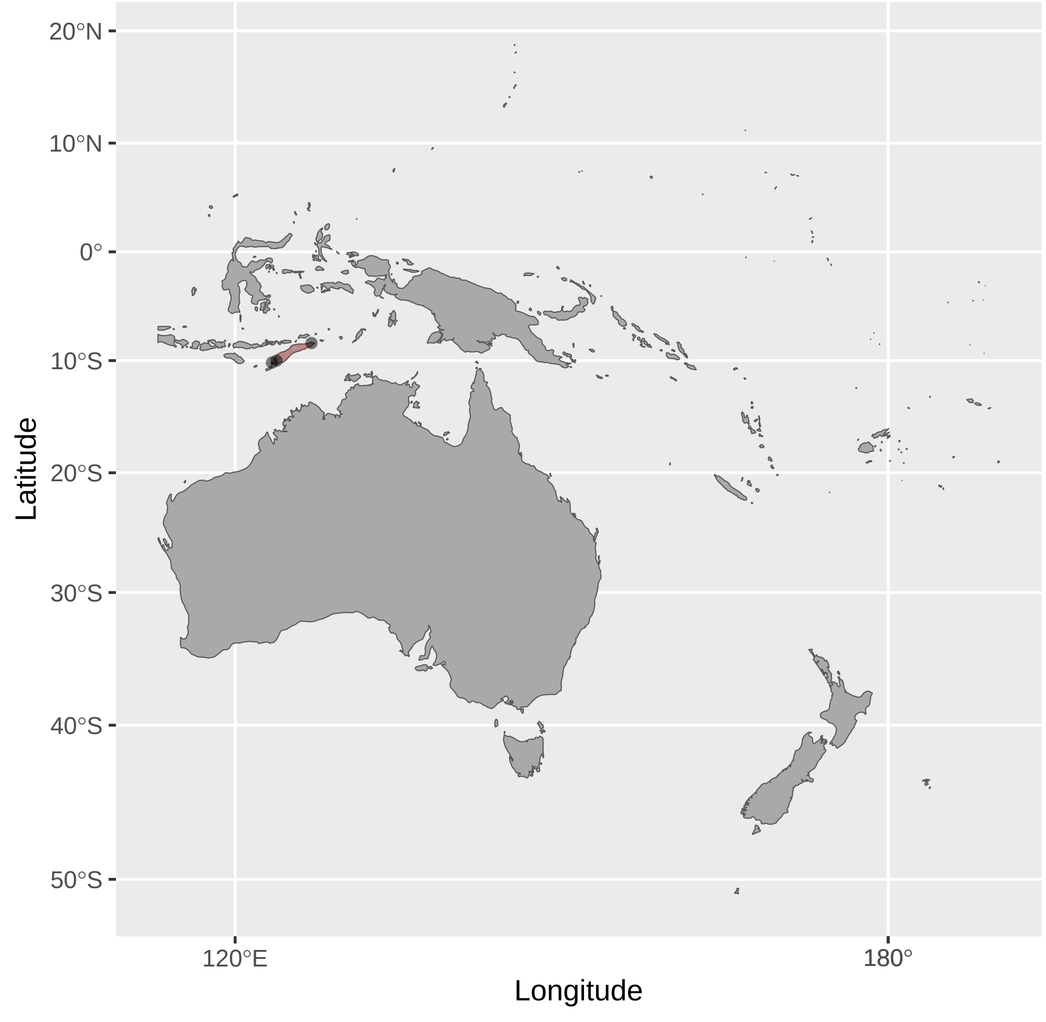
